# Azetidines kill *Mycobacterium tuberculosis* without detectable resistance by blocking mycolate assembly

**DOI:** 10.1101/2020.09.03.281170

**Authors:** Alice Lanne, Yixin Cui, Edward Browne, Philip G. E. Craven, Nicholas J. Cundy, Nicholas J. Coltman, Katie Dale, Antonio Feula, Jon Frampton, Aaron Goff, Mariwan A. Hama Salih, Xingfen Lang, Xingjian Li, Christopher W. Moon, Michael Morton, Jordan Pascoe, Xudan Peng, Vanessa Portman, Cara Press, Timothy Schulz-Utermoehl, Micky Tortorella, Zhengchao Tu, Zoe E. Underwood, Changwei Wang, Akina Yoshizawa, Tianyu Zhang, Simon J Waddell, Joanna Bacon, Cleopatra Neagoie, John S. Fossey, Luke J. Alderwick.

## Abstract

Tuberculosis (TB) is the leading cause of global morbidity and mortality resulting from infectious disease, with over 10 million new cases and 1.5 million deaths in 2019. This global emergency is exacerbated by the emergence of multi-drug-resistant MDR-TB and extensively-drug-resistant XDR-TB, therefore new drugs and new drug targets are urgently required. From a whole-cell phenotypic screen a series of azetidines derivatives termed BGAz, that elicit potent bactericidal activity with MIC_99_ values <10 μM against drug-sensitive *Mycobacterium tuberculosis* and MDR-TB were identified. These compounds demonstrate no detectable drug resistance. Mode of action and target deconvolution studies suggest that these compounds inhibit mycobacterial growth by interfering with cell envelope biogenesis, specifically late-stage mycolic acid biosynthesis. Transcriptomic analysis demonstrates that the BGAz compounds tested display a mode of action distinct from existing mycobacterial cell-wall inhibitors. In addition, the compounds tested exhibit toxicological and PK/PD profiles that pave the way for their development as anti-tubercular chemotherapies.

## Introduction

Tuberculosis (TB) is the principal infectious disease cause of death worldwide, accounting for 1.5 million deaths in 2019. One-third of the world’s population is currently infected with latent TB, and over ten million new cases of active TB are recognised per annum.^1,2^ Patients suffering with TB are treated with a cocktail of four drugs over a six-month period. Whilst cure rates can be as high as 90–95%,^3^ a combination of poor patient compliance and pharmacokinetic variability has led to the emergence of multi-drug-resistant (MDR) and extensively drug-resistant (XDR) TB.^4,5^ The alarming increase in MDR-TB (500,000 new cases in 2018),^1^ coupled with the fact that the last novel frontline anti-TB drug, Rifampicin, was discovered over 40 years ago^6^ suggests development and implementation of new control measures are essential for the future abatement of TB.^7^ Herein, the identity and anti-mycobacterial activity of azetidine derivatives with MIC_99_ values <10 μM against *Mycobacterium tuberculosis* are disclosed. These compounds did not give rise to emerging specific resistance in mycobacterial model organism *Mycobacterium smegmatis* and *Mycobacterium bovis* BCG. Mode of action and target deconvolution studies suggest mycobacterial growth inhibition is conferred by a hitherto uncharacterised mechanism that arrests late-stage mycolic acid biosynthesis. DMPK and toxicology profiles confirm that the azetidine derivatives identified display relevant and acceptable profiles for translation.

## Results

### Identification and development of azetidine derivatives with anti-mycobacterial activity

A bespoke compound library of novel lead-like small molecules^8^ for activity screening, that displayed a high fraction of sp^3^ (Fsp^3^, an indication of complexity and *3D-character*) atoms,^9–11^ that were free from pan-assay interference compounds (PAINS)^12,13^ and are synthetically tractable allowing for hit-to-lead scaffold elaboration^14^ were sought. Unrelated synthetic chemistry methodology studies^15^ proved to be an ideal untapped source of such compounds. ^16–19^ Compounds were fed into an open-ended anti-mycobacterial compound screen at the *University of Birmingham Drug Discovery Facility*.^20^ From which, an azetidine derivative **BGAz-001** (general formula Figure 1: R^1^ = R^2^ = H, R^3^ = Br, R^4^ and R^5^ = pyrrolidine fragment) that displayed promising anti-mycobacterial activity against both *Mycobacterium smegmatis* and *Mycobacterium bovis* BCG, with MICs of 30.5 μM and 64.5 μM, respectively (Table 1, entry 1) was identified and further azetidine derivatives were synthesised at *Guangzhou Institutes of Biomedical Health* (GIBH).^21^ Analysis of calculated medicinal chemistry properties and appraisal of preliminary screening against BCG at 2 μM and 20 μM in an end point REMA assay for anti-mycobacterial activity, alongside secondary MIC determination against BCG (MIC refers to MIC_99_ unless otherwise stated), was sufficient to rule out the majority of ancillary compounds for further study.^22^ Thus four additional azetidine-analogues with good activity against model organisms, were retained for further evaluation of anti-TB activity (Figure 1 and Table 1, entries 2-5: **BGAz-002**: R^1^ = OCF_3_, R^2^ = H, R^3^ = OCF_3_ and R^4^ & R^5^ = pyrrolidine fragment; **BGAz-003**: R^1^ = H, R^2^ = CF_3_, R^3^ = Br and R^4^ & R^5^ = pyrrolidine fragment; **BGAz-004**: R^1^ = H, R^2^ = Br, R^3^ = Br and R^4^ & R^5^ = pyrrolidine fragment; and **BGAz-005** where R^1^ = OCF_3_, R^2^ = H, R^3^ = OCF_3_, R^4^ =methyl and R^5^ = H, Table 1, entries 2-5)

**Figure 1.**
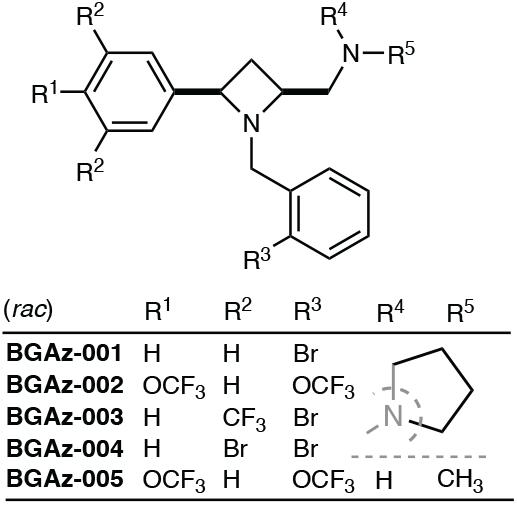
University of **B**irmingham and **G**uangzhou Institutes of Biomedicine and Health co-developed **Az**etidines (**BGAz**) with antitubercular activity.

**Table 1.**
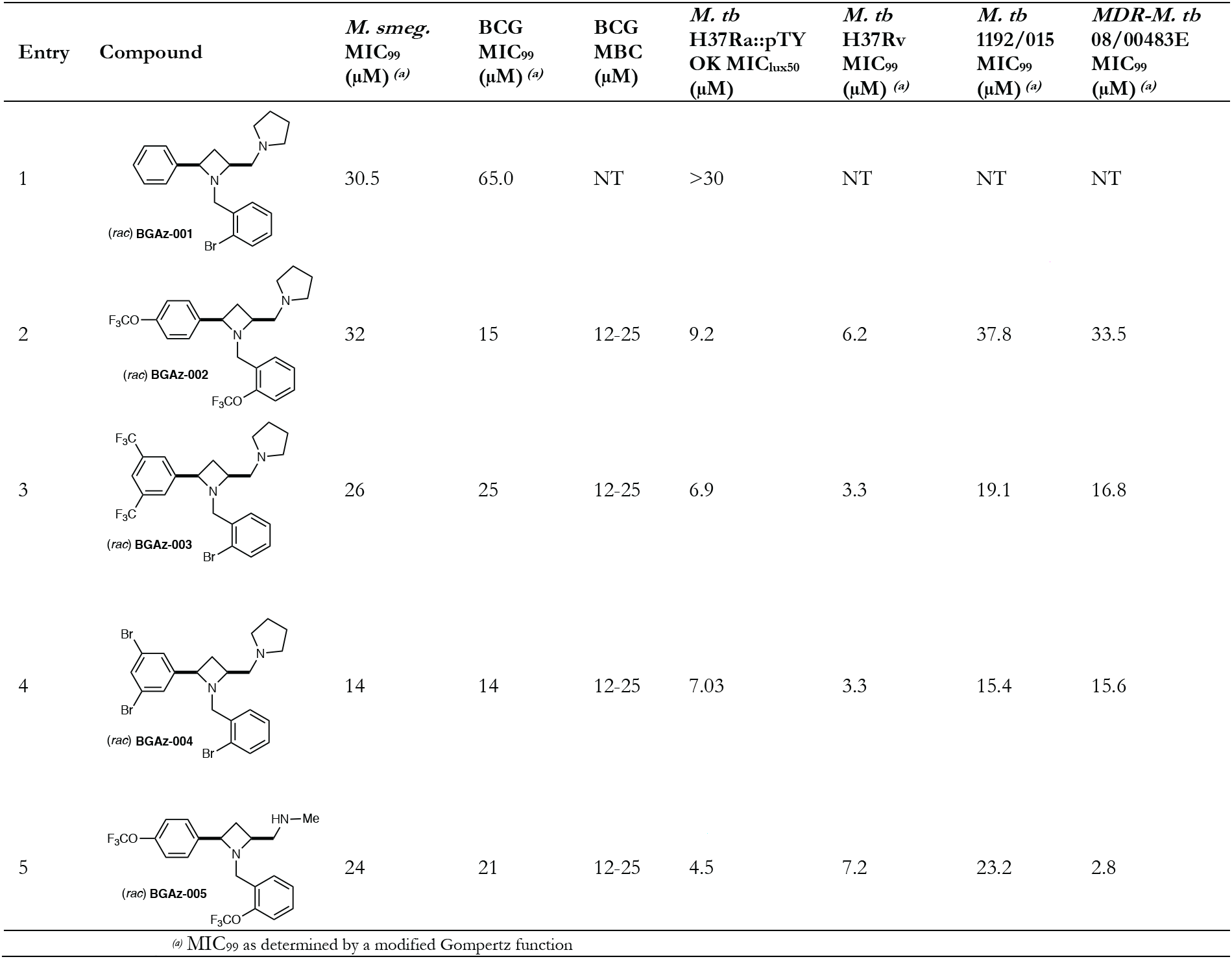
MIC and MBC values of the **BGAz-001** –**BGAz-005** against mycobacterial strains with different drug-susceptibility profiles. The *M. tb* H37Ra::pTYOK is an autoluminescent strain of mycobacteria.^23^ The MIC_99_ values of **BGAz-001** –**BGAz-005** against *M. smegmatis* and *M. bovis* BCG, drug-sensitive *M. tuberculosis* H37Rv (reference strain) and *M. tuberculosis* 1192/015 (clinical isolate), and multi-drug-resistant *M. tuberculosis* 08/00483E (clinical isolate resistant to INH, RIF, PZA and EMB). MIC values were determined from three biological replicates using a resazurin endpoint assay. The minimum bactericidal concentration (MBC) was determined for **BGAz-002** –**BGAz-005** against *M. bovis* BCG. NT = not tested.

| Entry | Compound | *M. smeg.*<br>MIC<sub>99</sub><br>(μM) <sup>(a)</sup> | BCG<br>MIC<sub>99</sub><br>(μM) <sup>(a)</sup> | BCG<br>MBC<br>(μM) | *M. tb*<br>H37Ra::pTY<br>OK MIC<sub>lux50</sub><br>(μM) | *M. tb*<br>H37Rv<br>MIC<sub>99</sub><br>(μM) <sup>(a)</sup> | *M. tb*<br>1192/015<br>MIC<sub>99</sub><br>(μM) <sup>(a)</sup> | MDR-*M. tb*<br>08/00483E<br>MIC<sub>99</sub><br>(μM) <sup>(a)</sup> |
| --- | --- | --- | --- | --- | --- | --- | --- | --- |
| 1 | (rac) **BGAz-001** | 30.5 | 65.0 | NT | >30 | NT | NT | NT |
| 2 | (rac) **BGAz-002** | 32 | 15 | 12-25 | 9.2 | 6.2 | 37.8 | 33.5 |
| 3 | (rac) **BGAz-003** | 26 | 25 | 12-25 | 6.9 | 3.3 | 19.1 | 16.8 |
| 4 | (rac) **BGAz-004** | 14 | 14 | 12-25 | 7.03 | 3.3 | 15.4 | 15.6 |
| 5 | (rac) **BGAz-005** | 24 | 21 | 12-25 | 4.5 | 7.2 | 23.2 | 2.8 |
| <sup>(a)</sup> MIC<sub>99</sub> as determined by a modified Gompertz function |  |  |  |  |  |  |  |  |

### Anti-tubercular activity of azetidine derivatives (BGAz-001–BGAz-005)

Compounds **BGAz002** –**BGAz005** displayed antitubercular activity against *M. tuberculosis* strains that include reference strains H37Ra::pTYOK and H37Rv, and two clinical isolates *M. tuberculosis* (Beijing/W lineage 1192/015) and *M. tuberculosis* (Beijing 08/00483E) that are drug-sensitive or multi-drug-resistant to isoniazid, rifampicin, pyrazinamide and ethambutol (Table 1).

Compounds **BGAz-002** –**BGAz-005** elicited anti-tubercular activities ranging from 4.5–9.2 μM MIC_lux50_ using a recently reported autoluminescent avirulent strain of *M. tuberculosis* H37Ra (Table 1, entries 2–5). Both **BGAz-003** and **BGAz-004** inhibit *M. tuberculosis* (H37Rv) at an MIC of 3.3 μM (Table 1, entries 3 and 4), with **BGAz-002** and **BGAz-005** inhibiting growth at MICs of 6.2 μM and 7.2 μM, respectively (Table 1, entries 2 and 5). The MICs determined for **BGAz-002** –**BGAz-005** are higher in the drug-sensitive *M. tuberculosis* Beijing/W (1192/015) clinical isolate compared to H37Rv. When comparing MICs between drug-sensitive and drug MDR clinical strains, no significant differences were observed for **BGAz-002** –**BGAz-004**, suggesting that the acquisition of mutations conferring front-line drug resistance does not impact upon the antitubercular activity of these compounds. However, it is noteworthy that when **BGAz-005** was tested against *M. tuberculosis* 08/00483E (a clinical isolate that is resistant to isoniazid, rifampicin, pyrazinamide and ethambutol),^24^ this compound displayed an eight-fold lower MIC (2.8 μM, Table 1, entry 5) in comparison to the drug-sensitive isolate. The absence of cross-resistance of these new anti-TB agents with the current frontline TB drugs is an important consideration in development of new TB therapies with distinct modes of action.^25^ Evaluation of the Minimal Bactericidal Concentrations (MBCs) for **BGAz-002** –**BGAz-005** against BCG demonstrates that these compounds exhibit bactericidal activity, since both MIC and MBC values overlap (Table 1).

### Physiochemical and Toxicological properties of BGAz-002–BGAz-005

**BGAz-002** –**BGAz-005** were subjected to *in vitro* DMPK testing; **BGAz-001** was excluded from further study due to its comparatively poor anti-mycobacterial activity. The poorer kinetic solubility **BGAz-002** –**BGAz-004** (9 to 57 μM) in aqueous buffered solution in comparison to the higher solubility of **BGAz-005** (117 μM) can be attributed to the presence of a secondary *versus* tertiary amine functionality (Table 2). Metabolic stability of the compounds was evaluated through measuring the intrinsic clearance (CL_int_) by mouse liver microsomes and by liver hepatocytes. Compounds **BGAz-002** –**BGAz004** all exhibited a CL_int_. of >150 μL/min/mg in the microsomal stability assay indicating a rapid clearance (Table 2, entries 1 to 3), with **BGAz-005** giving the lowest rate of microsomal clearance (36 μL/min/mg, Table 2, entry 4). Experiments were repeated using mouse liver hepatocytes and all four compounds afforded CL_int_ values of <60 μL/min/mg, indicating good overall metabolic stability (Table 2). Caco-2 permeability assays were conducted to predict both intestinal permeability and drug efflux. Compounds **BGA-002** –**BGAz-004** exhibited poor efflux ratios whilst **BGAz-005** continued to perform well with an efflux ratio of less than 1.0.

**Table 2.**
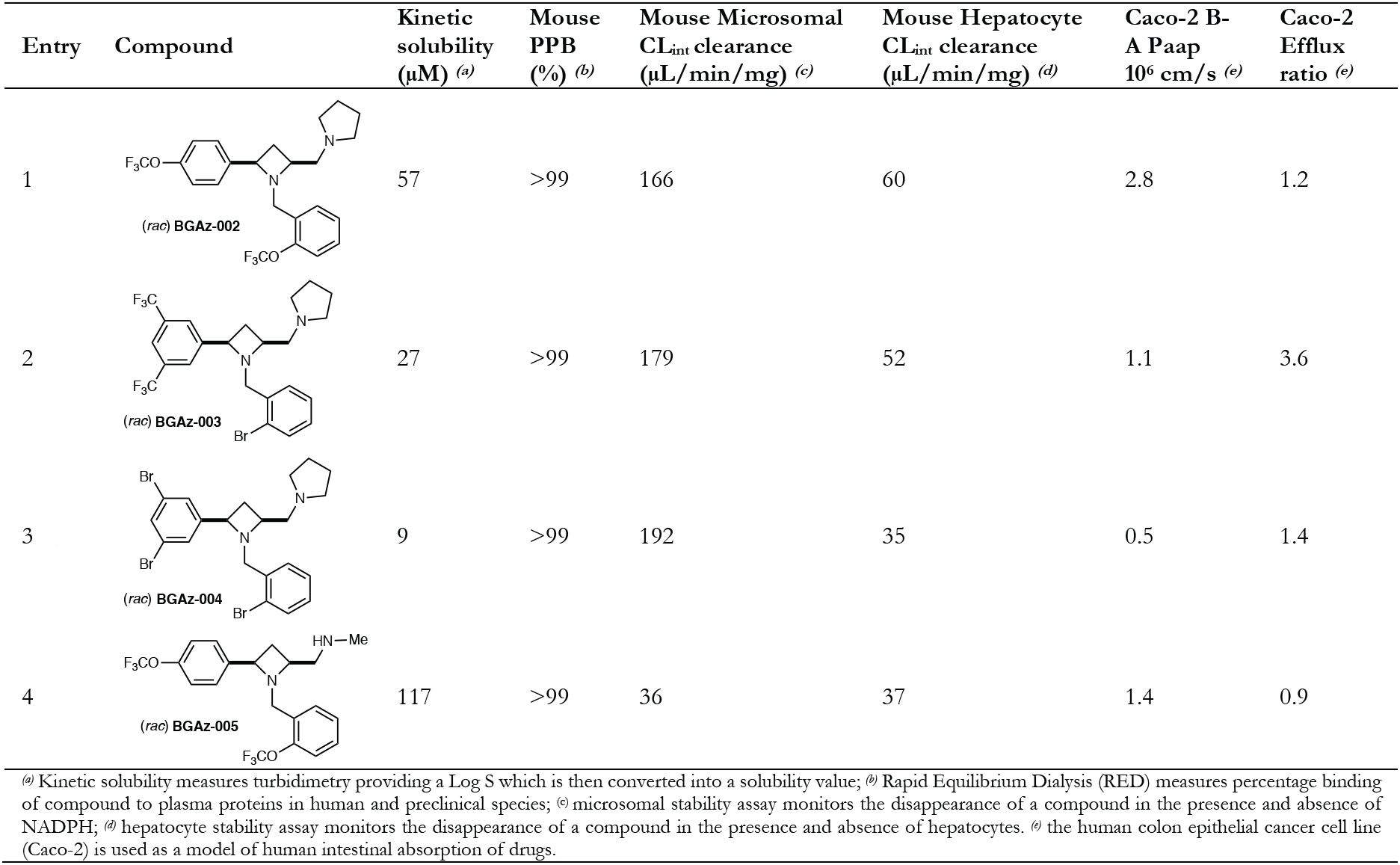
Physiochemical and toxicological properties of **BGAz-002** –**BGAz-005**.

| Entry | Compound | Kinetic solubility (μM) <sup>(a)</sup> | Mouse PPB (%) <sup>(b)</sup> | Mouse Microsomal CL<sub>int</sub> clearance (μL/min/mg) <sup>(c)</sup> | Mouse Hepatocyte CL<sub>int</sub> clearance (μL/min/mg) <sup>(d)</sup> | Caco-2 B-A Paap 10<sup>6</sup> cm/s <sup>(e)</sup> | Caco-2 Efflux ratio <sup>(e)</sup> |
| --- | --- | --- | --- | --- | --- | --- | --- |
| 1 | (rac) **BGAz-002** | 57 | >99 | 166 | 60 | 2.8 | 1.2 |
| 2 | (rac) **BGAz-003** | 27 | >99 | 179 | 52 | 1.1 | 3.6 |
| 3 | (rac) **BGAz-004** | 9 | >99 | 192 | 35 | 0.5 | 1.4 |
| 4 | (rac) **BGAz-005** | 117 | >99 | 36 | 37 | 1.4 | 0.9 |

<sup>(a)</sup> Kinetic solubility measures turbidimetry providing a Log S which is then converted into a solubility value; <sup>(b)</sup> Rapid Equilibrium Dialysis (RED) measures percentage binding of compound to plasma proteins in human and preclinical species; <sup>(c)</sup> microsomal stability assay monitors the disappearance of a compound in the presence and absence of NADPH; <sup>(d)</sup> hepatocyte stability assay monitors the disappearance of a compound in the presence and absence of hepatocytes. <sup>(e)</sup> the human colon epithelial cancer cell line (Caco-2) is used as a model of human intestinal absorption of drugs.

The pharmacokinetic (PK) parameters of **BGAz-001** –**BGAz-005** in a mouse model were investigated by cassette (combined) dosing at 5 mg/kg PO, 1 mg/kg IV and IP (Supplementary PK Data file: Tables 1 to 5). For the oral dosing, at 5 mg/kg, **BGAz-001**, **BGAz-002** and **BGAz-003** gave peak serum concentrations (C_max_) of 54.7, 53.9 and 56.0 μg/L respectively. **BGAz-004** gave the highest C_max_ whereas **BGAz-005** gave the lowest, 87.6 and 43.1 μg/L respectively (Table 3, entries 2 and 3). The plasma half-lives (*T*_1/2_) revealed **BGAz001** to be 1.5 h, **BGAz002** and **BGAz003** to be 24.9 and 28.0 h respectively. **BGAz004** has a half-life of 11.4 h with **BGAz005** having the longest half-life of 35.7 h. **BGAz001–003** were excluded from further study due to poor PK/PD parameters. Mice were dosed multiple times with **BGAz-004** and **BGAz-005** (four and three times respectively) at 30 mg/kg (PO) in order to provide evidence of compound tolerability and refinement of measured parameters. **BGAz-004** and **BGAz-005** gave C_max_ values of 363.0 and 1712.5 μg/L (respectively); *T*_1/2_ values of 8.1 and 80.4 hours were calculated respectively (Table 3, entries 2 and 3). The total body exposure from multiple dosing at 30 mg/kg (PO) of **BGAz-005** (82247.7 ng/mL*h) was significantly greater than **BGAz-004** (5420.99 ng/mL*h), indicating a superior overall pharmacokinetic profile for **BGAz-005** (Table 3, entries 2 and 3). **BGAz-002**, **BGAz-004** and **BGAz-005** were tested for cytochrome P450 (CYP450) metabolic activity by measuring the inhibition each of specific enzymes in human liver microsomes. All three compounds exhibited no discernible inhibition of CYP1A2, an enzyme known to metabolise aromatic/heterocyclic amine-containing drugs (Table 3). The CYP2C9 enzyme is a relatively abundant CYP450 in the liver that dominates CYP450-mediated drug oxidation. In this regard, only minimal CYP2C9 inhibition when **BGAz-004** was pre-incubated for 30 min prior to the addition of NADPH to initiate catalysis was observed. Known to metabolise a wide range of drug molecules, CYP2C19 is an essential member of the CYP450 superfamily as it contributes ~16% of total hepatic content in humans. While **BGAz-002** and **BGAz-005** displayed only negligible inhibition of this enzyme, **BGAz-004** displays strong inhibitory activity when pre-incubated for 30 min prior to initiation of catalysis (Table 3). CYP2D6 is widely implicated in the metabolism of drugs that contain amine functional groups, such as monoamine oxidase inhibitors and serotonin reuptake inhibitors. CYP2D6 is responsible for the second highest number of drugs metabolised by the CYP450s, as demonstrated by the significant inhibition of this enzyme by **BGAz-002**, **BGAz-004** and **BGAz-005**. All four compounds were evaluated for mitochondrial dysfunction by measuring IC_50_ values against HepG2 cells cultured in media containing either glucose or galactose, which serves to direct cellular metabolic activity towards glycolysis or oxidative phosphorylation, respectively. Whilst **BGAz-004** exhibited a negligible effect both **BGAz-002** and **BGAz-005** demonstrated cytotoxicity with respective IC_50_ values of 38 μM and 21 μM, respectively (Table 3). In order to enhance cellular susceptibility to mitochondrial toxicants, assays were repeated in the presence of galactose which resulted in Glu/Gal ratios of <1, confirming no mitochondrial toxicity. Compounds **BGAz-002**, **BGAz-004** and **BGAz-005** were assayed for hERG inhibition using *IonWorks* patch clamp electrophysiology. An eight-point concentration response curve was generated from three-fold serial dilution of a top compound concentration of 167 μM. Compound **BGAz-002** was the best-performing compound, displaying a hERG liability IC_50_ of 173 μM, (i.e. inhibition of less than 50% at top 167 μM test concentration). In comparison, compounds **BGAz-004** and **BGAz-005** performed significantly worse with hERG liability IC_50_ of 25 μM and 12.7 μM, respectively. Overall, the BGAz compounds investigated in this study display an encouraging toxicological and PK/PD profile to enable further exploration and development towards the clinic.

**Table 3.**
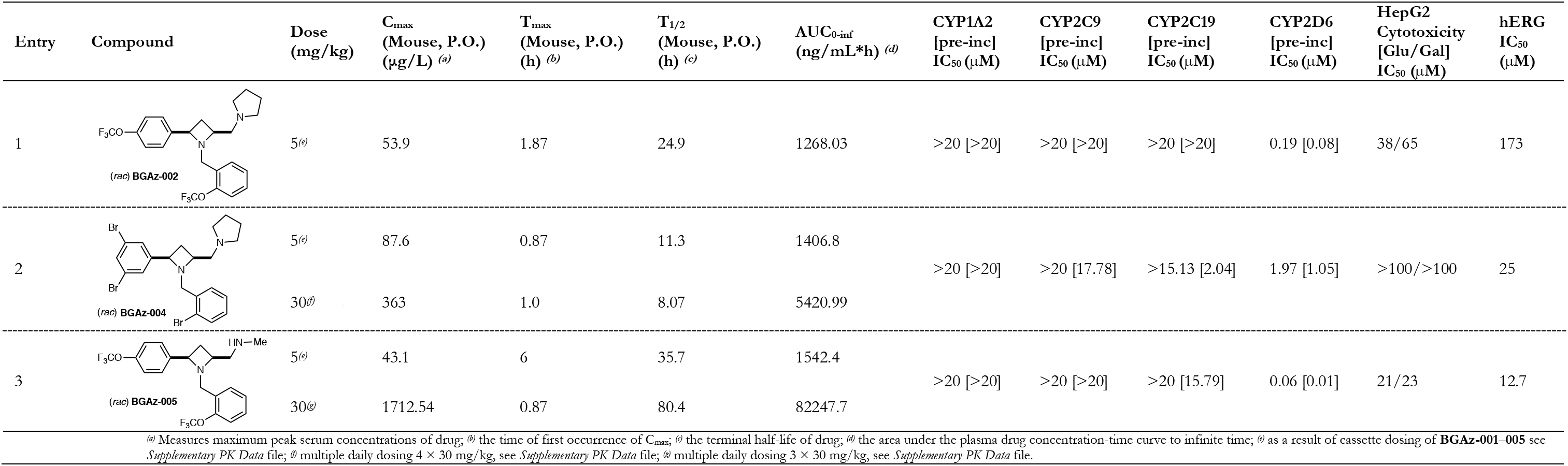
Pharmacokinetic profiles, CYP450 activities, mitochondrial dysfunction and hERG liabilities of **BGAz-002**, **BGAz-004** and **BGAz-005**.

### BGAz compounds kill M. tuberculosis with bactericidal activity

The bactericidal activity of **BGAz-004** and **BGAz-005** against *M. tuberculosis* H37Rv was further assessed by exposing the bacilli to a range of concentrations of **BGAz004** and **BGAz-005** over a time-course of 14 days and total viable counts (CFUmL^−1^) enumerated on solid medium. Both **BGAz-004** and **BGAz-005** were active against *M. tuberculosis* H37Rv (Figure 2, panels A and B) with **BGAz-005** demonstrating statistically significantly greater early bactericidal and concentration-dependent activity than **BGAz-004** at day six (*P* = 0.046) and day ten (*P* =0.049) (Figure 2D). Exposure of *M. tuberculosis* H37Rv to **BGAz-005** resulted in a greater bactericidal effect with more pronounced activity earlier in the time-course, with a reduction of 3.28 ± 1.00 log_10_ CFUmL^−1^ after six days of exposure and reduction of 3.99 + 0.60 log_10_ CFUmL^−1^ after 14 days’ exposure, at a concentration of 96 μM (Figure 2B). Equivalent activity was not observed by **BGAz-004** early in the time-course and showed delayed activity at all concentrations, only achieving a decrease in 0.79 ± 2.20 log_10_ CFUmL^−1^ by day six and 1.90 ± 1.28 log_10_ CFUmL^−1^ reduction after 14 days at 96 μM (Figure 2A). The profile for **BGA_Z_-004** is commensurate with antibiotics that exhibit bacteriostatic activity at lower concentrations. Isoniazid, demonstrated a higher rate of bactericidal activity compared to both **BGAz-004** and **BGAz-005** by achieving a reduction of >4.52 ± 0.60 log_10_ CFUmL^−1^, to a limit of detection (100 CFUmL^−1^), by day ten, at a lower concentration of 29 μM (Figure 2C).

**Figure 2.**
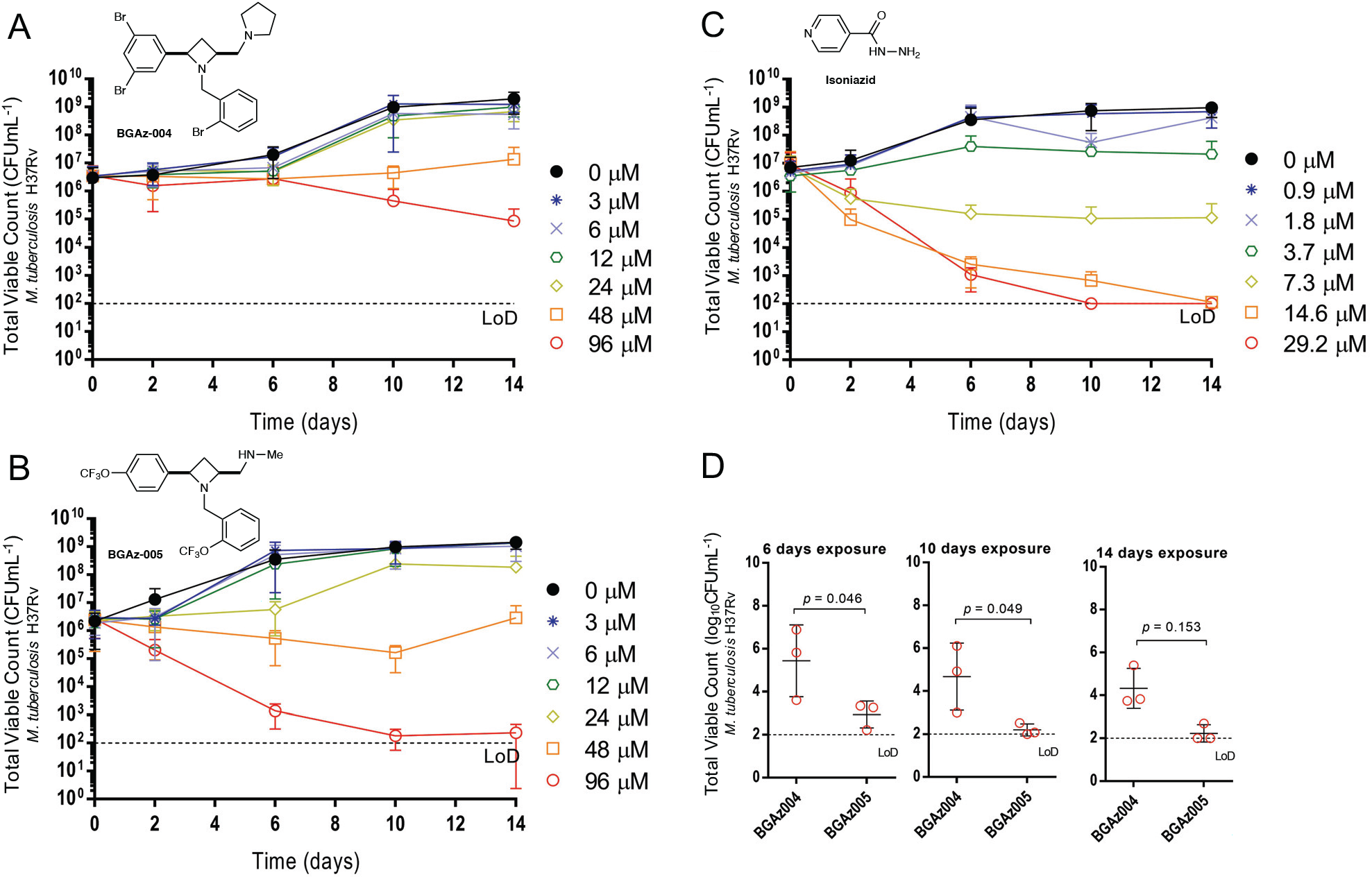
Assessment of bactericidal activity of **BGAz-004** and **BGAz-005** against *M. tuberculosis* H37Rv. Average total viable counts (CFUmL-1) of *M. tuberculosis* cultures exposed to either **BGAz-004** (**Panel A**) or **BGAz-005** (**Panel B**) at concentrations: 0 μM (0.1% DMSO) (circle, closed), 3 μM, 6 μM, 12 μM, 24 μM, 48 μM and 96 μM, or isoniazid (**Panel C**) at concentrations 0 μM (0.1% DMSO), 0.9 μM, 1.8 μM, 3.7 μM, 7.3 μM, 14.6 μM and 29.2 μM over a 14-day time-course. Samples were taken after 0, 2, 6, 10, and 14 days of antibiotic exposure, serially diluted, and plated by the method of Miles *et al*.^26^ Statistical comparisons were performed at 6, 10 and 14 days of antibiotic exposure at 96 μM **BGAz-004** and **BGAz-005** using factorial ANOVA and post-hoc Tukey’s honestly significant difference test (**Panel D**). Data represents three biological repeats ± standard deviation.

In addition to the enumeration of viable bacilli on agar, the activity of **BGAz-004** and **BGAz-005** was determined using flow cytometry. This approach allows for a direct assessment of whether BGAz compounds are able to kill *M. tuberculosis* in a dose dependent manner and whether the killing profile was similar between these compounds and to that observed for isoniazid, which would provide insights about their mode of action.^27^ Culture samples were taken at each time-point and dual-stained using Calcein Violet with an acetoxy-methyl ester group (CV-AM) that is a correlate of metabolic activity, and Sytox Green (SG) that enables measurement of cell-wall permeability (a proxy for cell death). Single bacilli were identified by forward scattered light area and height using flow cytometry analyses. Gated single cells were further differentiated based upon the presence and absence of CV-AM and SG staining using a quadrant gating approach. The percentages of the population that are unstained or stained with each dye (or both dyes) is represented in four gates P1-P4 (P1: CV-AM^−^/SG^−^, P2: CV-AM^+^/SG^−^, P3: CV-AM^+^/SG^+^, P4: CV-AM^−^/SG^+^) (Figure 3). The CV-AM staining profiles (metabolic activity) for these compounds were reflective of the total viable counts (Figure 3); the decrease in CV-AM staining over the time-course at 96 μM for **BGAz-005** was statistically significant after day six (*P* = 0.043), day ten (*P* = 0.08), and day fourteen (*P* = 0.011) compared to the decrease in the CV-AM staining for **BGAz-004**, at the same concentration (Figure 3A and B; P2). A similar difference in activity was observed at 48 μM, (*P* = 0.067, 0.065, 0.066 for days 6, 10 and 14, respectively). The SG-staining profiles showed that both compounds possessed equivalent killing activity at high concentrations of 96 μM (Figure 3A and B; P4); however, **BGAz-005** shows higher levels of kill at days six and fourteen with a lower concentration of 48 μM (*P* = 0.027 and 0.068, respectively). Both **BGAz-004** and **BGAz-005** show a similar staining profiles to isoniazid (Figure 3C), which targets the mycobacterial cell-wall.^27^

**Figure 3.**
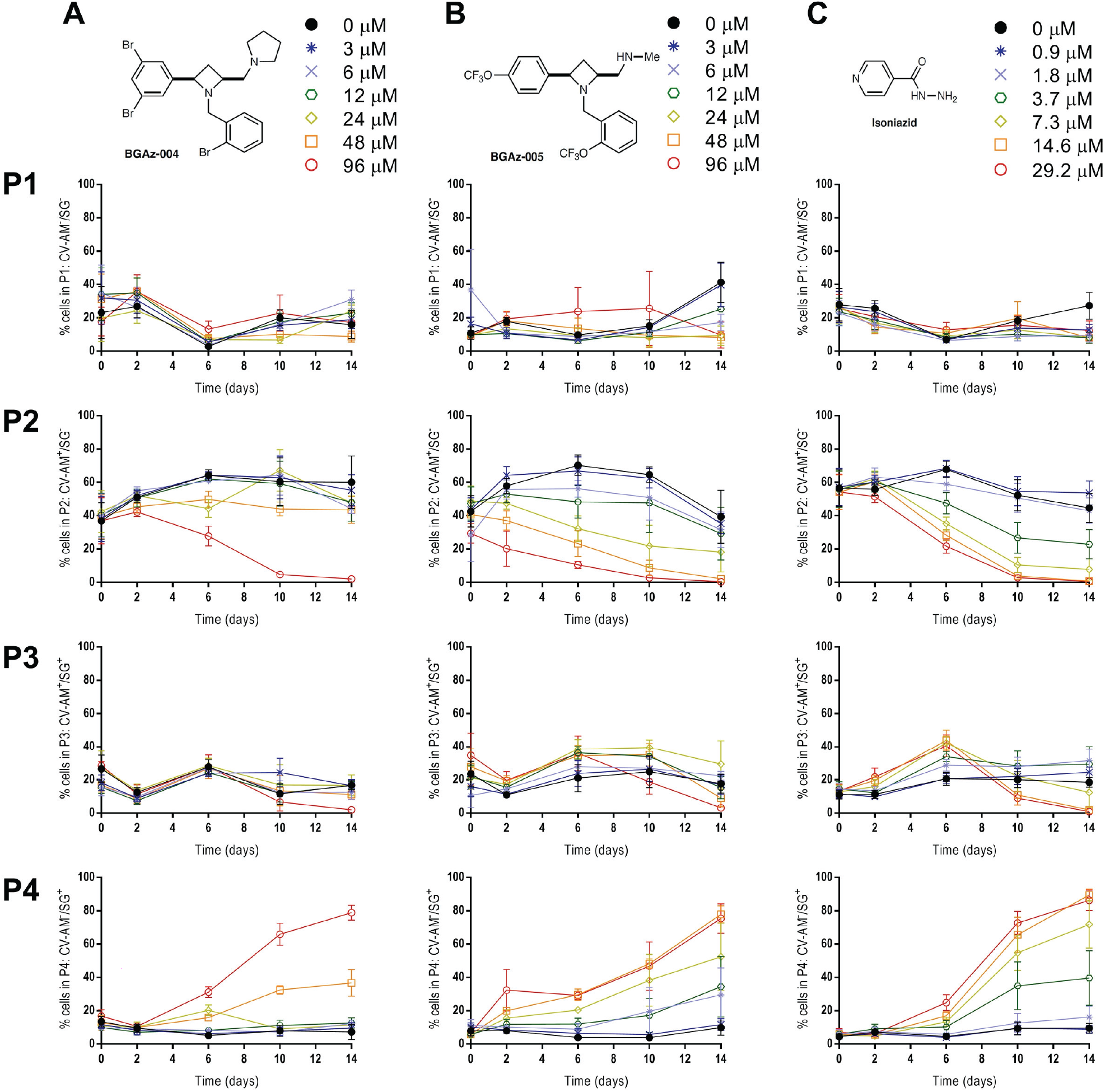
Assessment of bactericidal activity of BGAz-004 and BGAz-005 against M. tuberculosis H37Rv. Quantitation of Calcien-Violet-AM (CV-AM) and Sytox-green (SG) fluorescence of M. tuberculosis H37Rv, using flow cytometry, after exposure to BGAz-004 (column A), BGAz-005 (column B) at concentrations: 0 μM (0.1% DMSO), 3 μM, 6 μM, 12 μM, 24 μM, 48 μM and 96 μM or (column C) isoniazid at concentrations 0 μM (0.1% DMSO), 0.9 μM, 1.8 μM, 3.7 μM, 7.3 μM, 14.6 μM and 29.2 μM over a 14-day time-course. The percentages of the population that are unstained or stained with each dye (or both dyes) is represented in four gates (rows P1–P4). Row P1: unstained population (CV-AM^−^/SG^−^); row P2: CV-stained population (CV-AM^+^/SG^−^); row P3: dual-stained population (CV-AM^+^/SG^+^); and row P4: SG-stained population (CV-AM^−^/SG^+^). Data represents three biological repeats ± standard deviation. Statistical comparisons were made using factorial ANOVA and *post-hoc* Tukey’s honestly significant difference test.

### BGAz-004 and BGAz-005 inhibit the incorporation of mycobacterial cell-wall precursors and display no detectable resistance

Whole genome sequencing (WGS) of laboratory-generated mutants that are resistant to TB drugs is a widely used approach to determine the mode of action of novel antibacterial compounds.^28–30^ Multiple attempts (>5 biological repeats) to generate drug-resistant mutants of **BGAz-002–BGAz-005** in *M. smegmatis* and *M. bovis* BCG (including a strain of BCG devoid of *recG* which has a higher mutational frequency)^31^ were unsuccessful, implying an undetectably low frequency of resistance for these compounds (Supplementary Information). This advantageous property is a double-edged sword. The discovery of the BGAz series as novel antitubercular compounds with low frequencies of resistance is attractive in terms of drug development, especially in the context of MDR-TB; however, the inability to generate resistant mutants against the most active compounds suggests that this series of compounds may elicit pleiotropic activity or have non-specific modes of action, or non-protein target(s). Therefore, in order to investigate the mode of action of **BGAz-005**, the most active of the compounds tested against mycobacteria, biosynthetic inhibition of five major macromolecular pathways were evaluated by measuring incorporation of selected radiolabelled precursors during microbial cell culture. Addition of **BGAz-005** up to a concentration of 0.75 × MIC had almost no effect of the incorporation of [^3^H]-thymidine, [^3^H]-uridine and [^3^H]-leucine with only a moderate 20% reduction of incorporation at 1 × MIC, suggesting that **BGAz-005** does not directly inhibit DNA, RNA or protein biosynthesis (Figure 4). In contrast, **BGAz-005** decreased the incorporation of both [^3^H]-DAP and [^14^C]-acetic acid from six hours post-labelling and at 0.5 × and 1 × MIC, caused a titratable decrease in [^14^C]-acetic acid incorporation, exerting a ~50% and ~75% loss of lipid biosynthesis, respectively (Figure 4). These data suggest that the **BGAz-005** acts by inhibiting aspects of mycobacterial cell envelope biosynthesis.

**Figure 4.**
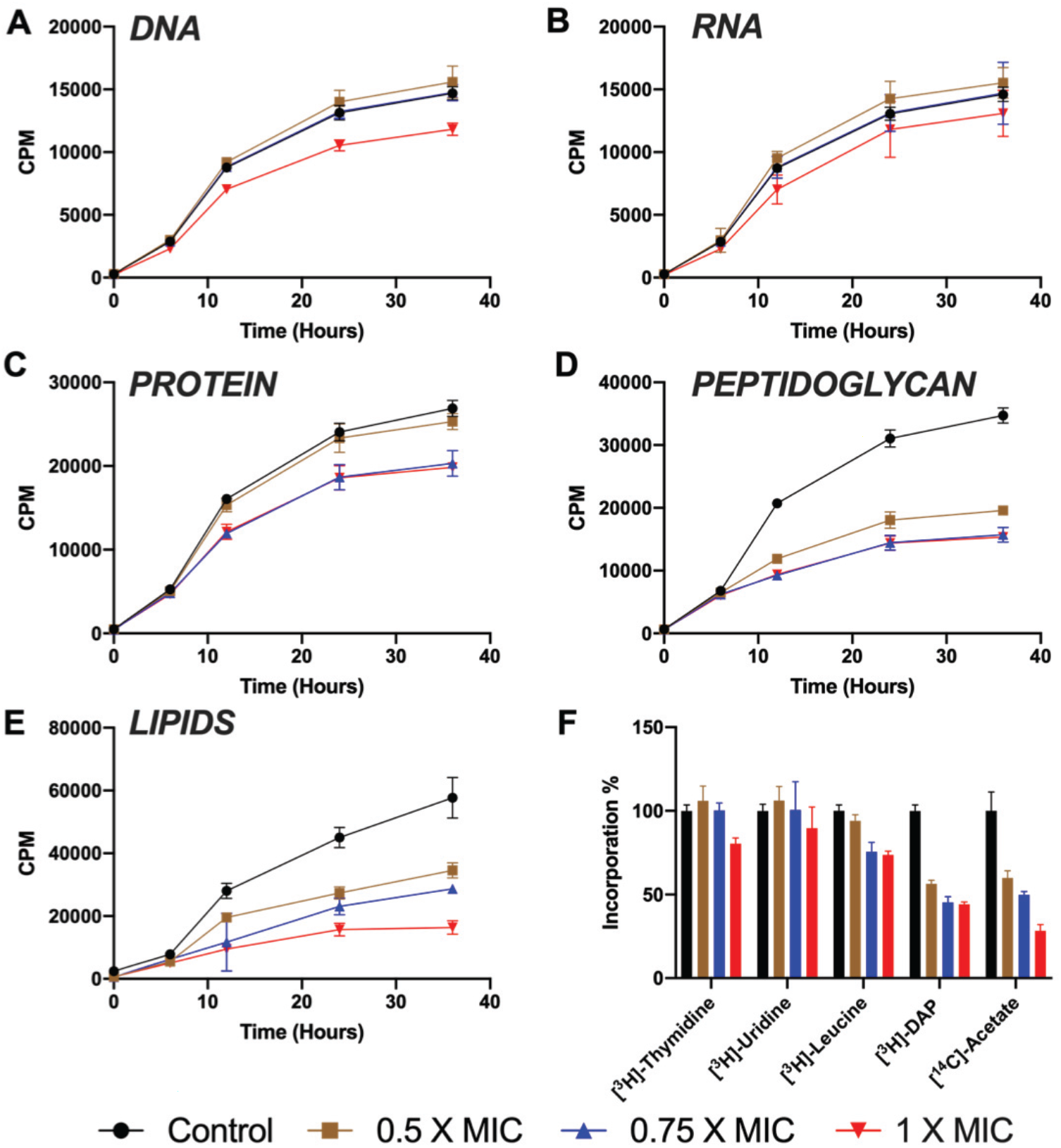
Effect of **BGAz-005** on the incorporation of radiolabelled precursors into the major cellular macromolecules of *M. smegmatis.* The incorporation of (**A**) [*methyl-*^3^H]thymidine (for DNA), (**B**) [5,6-^3^H]uridine (for RNA), (**C**) L-[4,5-^3^H]leucine (for protein), (**D**) [^3^H]*meso-*diaminopimelic acid (for peptidoglycan) and (**E**) [^14^C]acetic acid (for lipids) were measured over a period of 36 hours. The percentage of incorporation measured at 36 hours is represented in panel F. Each plot and error bars represent the average of three independent experiments.

### BGAz-005 dysregulates the expression of cell envelope biosynthetic genes

To explore the mode of action of **BGAz-005** using an unsupervised approach, the *M. bovis* BCG transcriptional response to drug exposure was profiled by RNAseq. A signature consisting of 160 induced and 126 repressed genes was identified after eight hours’ exposure to 1 × MIC **BGAz-005**. The response was comprised of three principal features, namely, inhibition of cell-wall biosynthesis, dysregulation of metal homeostasis and disruption of the respiratory chain (Figure 5). The inhibition of cell-wall synthesis was evidenced by induction of key regulators of cell-wall stress *sigE* and *mprAB*, alongside significant upregulation of their regulons (hypergeometric p value of *sigE* regulon enrichment 2.47×10^−17^;^32^ *mprAB* hgp 5.29×10^−11^).^33^ In contrast to isoniazid and ethambutol where FasII genes are induced by drug exposure, FasII genes (*hadA*, *fabG1*, *inhA*, *acpM*) alongside mycolic acid synthesis and modification genes (*mmaA2*, *mmaA3*, *fbpA*, *fbpB*, *fbpD*, *desA2*) were repressed by **BGAz-005** treatment, indicating a different mechanism of **BGAz-005** drug action to cell-wall inhibitors currently in use. The functional category (I.H) lipid biosynthesis was significantly repressed by **BGAz-005** (hgp 6.52×10^−5^), and mycolyl-arabinogalactan-peptidoglycan complex biosynthesis was the top pathway dysregulated by **BGAz-005** (pathway perturbation score of 3.4).^34,35^ Genes involved in the synthesis of alternative cell-wall factors, sulfolipids (*mmpL8*, *papA1*, *pks2*) and the oleic acid stearoyl-CoA desaturases that produce phospholipids (*desA3_1*, *desA3_2*, *BCG_3260c/Rv3230c*) were induced.^36^ A series of metal-responsive regulatory systems were upregulated by **BGAz-005** (*cmtR*, *zur*, *ideR*, *tcrYX*) as well as genes encoding the lipid-bound siderophore mycobactin (*mbtB*, *mbtC*, *mbtD*), representing disruption of metal control systems, likely impacted by loss of cell-wall structure. Induction of redox-inducible *clgR* in combination with repression of the *dosR* regulon (hgp 6.40×10^−9^) reflected the impact of **BGAz-005** on the respiratory chain.^37^ However, unlike many drugs that affect respiration, no differential expression of energy metabolism systems (*nuoA-N*, *qcrA-C*, *ctaC-E*, *cydA-D*, *narG-J*) was observed.^38,39^ Systems implicated in the efflux (*mmpL5*, *mmpS5, BCG_0727*/*Rv0678*, *BCG_0728c*/*Rv0679c*) or detoxification (*BCG_3184c*/*Rv3160c*, *BCG_3185c*/*Rv3161c*, *BCG_3186c*/*Rv3162c*) of antimicrobial drugs were also induced by **BGAz-005**.^40^ Significantly, the efflux pump *efpA*, highly induced by cell-wall targeting drugs isoniazid, ethambutol and benzothiazinone, was not induced by **BGAz-005** exposure.

**Figure 5.**
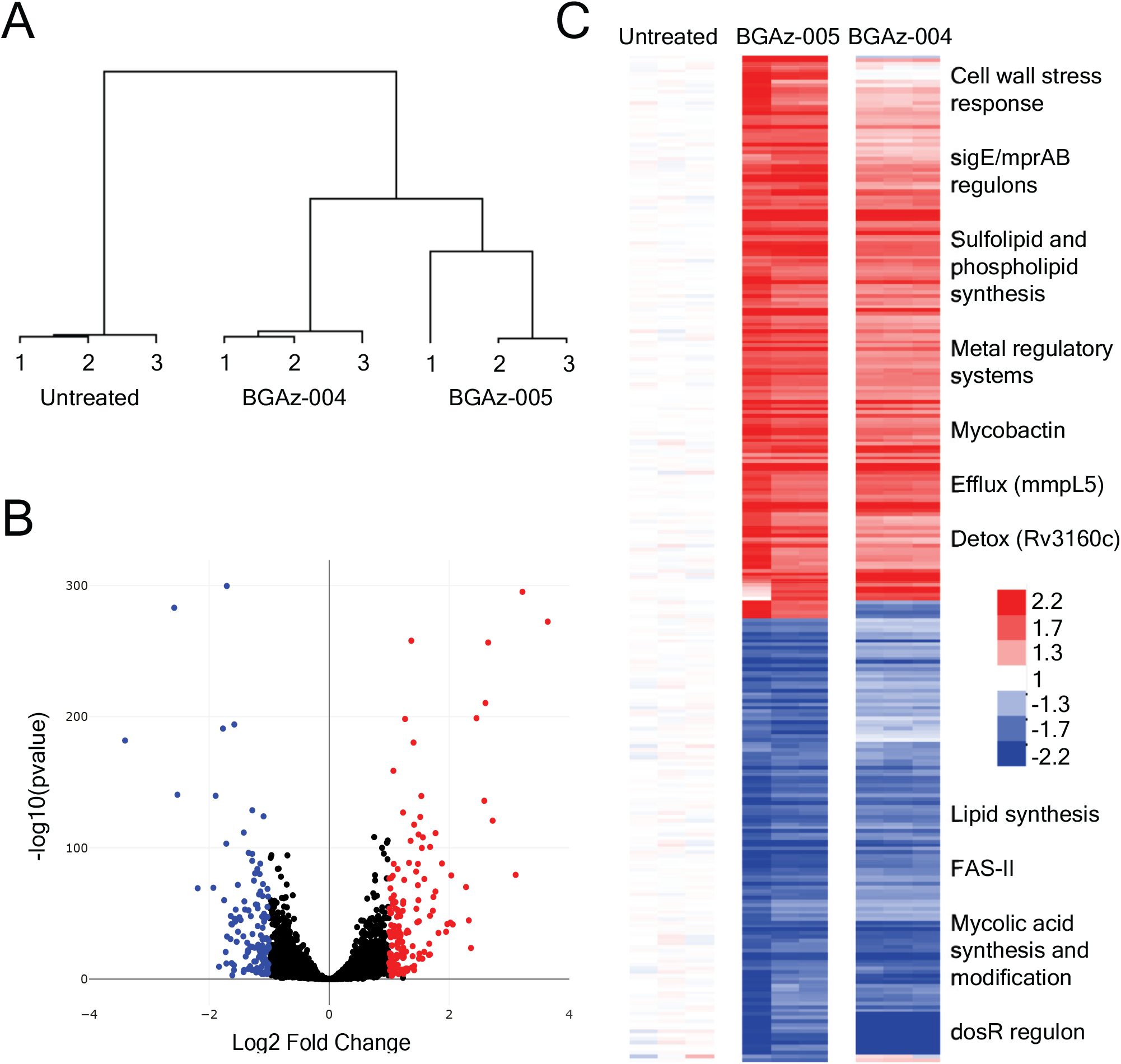
The transcriptional response to **BGAz-005** exposure demonstrating inhibition of mycobacterial cell envelope biosynthesis. (**A**) Cluster diagram of all genes showing similarity of biological replicates and separation of drug-treated from carrier control samples. (**B**) Volcano plot of *M. bovis* BCG response to **BGAz-005**, highlighting genes significantly differentially expressed. (**C**) Heatmap of 286 gene **BGAz-005** signature relative to carrier control. Conditions as columns, genes as rows; red colouring highlighting induced genes, blue repressed genes. The **BGAz-004** signature is clustered alongside indicating a similar mode of drug action.

Mapping the drug-responsive gene clusters identified by Boshoff and co-workers revealed significant enrichment of GC-27 and GC-82 representing cell-wall inhibition,^41^ alongside GC-39 (*dosR* regulon) and GC-108 (iron scavenging). The most similar drug signatures were the phenothiazines, chlorpromazine and thioridazine (hgp 2.83×10^−14^), disrupting the cell-wall and electron transfer chain,^42^ alongside analogues of ethambutol (hgp 4.98×10^−13^)^43^ and benzothiazinone (hgp 8.00×10^−8^) targeting arabinose biosynthesis in the mycobacterial cell-wall.^44^ Thus, **BGAz-004** and **BGAz-005** elicit a transcriptomic response representing major abrogation of normal cell envelope function.

### BGAz-004 and BGAz-005 significantly alter mycobacterial cell envelope composition

The results of transcriptomic profiling and whole cell phenotyping support the hypothesis that **BGAz-004** and **BGAz-005** inhibit aspects of cell envelope biosynthesis. To further investigate the mechanism by which mycobacterial cell envelope lipid composition is affected, actively growing cultures of BCG were exposed to increasing concentrations of **BGAz-005** followed by metabolic labelling using [^14^C]-acetic acid. Autoradiographaphs of cell envelope lipids separated by thin layer chromatography (TLC) revealed that treatment of BCG with **BGAz-005** at 0.5 × MIC caused a significant reduction in trehalose monomycolate (TMM) and trehalose dimycolate (TDM) and a complete loss of TMM and TDM at concentrations beyond the MIC (Figure 6A). The formation of cytoplasmic membrane phospholipids (PIMs and CL) remain unaffected (Figure 6A), The analysis of lipids loaded and separated by TLCs that had been normalised for total lipids extracted, revealed an altered lipid profile highlighting the accumulation of an unidentified lipid species that resolves to a relatively high Rf (Lipid species X, Figure 6B). The analysis of mycolic acid methyl esters (MAMES) reveals that both alpha and keto mycolates bound to the cell-wall arabinogalactan (AG) are gradually depleted as BCG is exposed to increasing concentrations of **BGAz-005** during active cell culture (Figure 6C). Quantification of the relative abundance of each lipid species highlights the significant depletion of mycolates (either conjugated to trehalose in the form of TMM/TDM or AG) when **BGAz-005** is used at a half MIC, whilst other lipids including PI and PIMS remain largely unaffected (Figure 6D). **BGAz-004** affects mycobacterial cell envelope lipid biosynthesis in an almost identical manner (Supplementary Figure 3). The immediate and specific arrest in the biosynthesis of TMM and TDM, and as a result, loss of esterified mycolates to AG, strongly supports the hypothesis that **BGAz-004** and **BGAz-005** inhibit mycobacteria by targeting mycolate biosynthesis.

**Figure 6.**
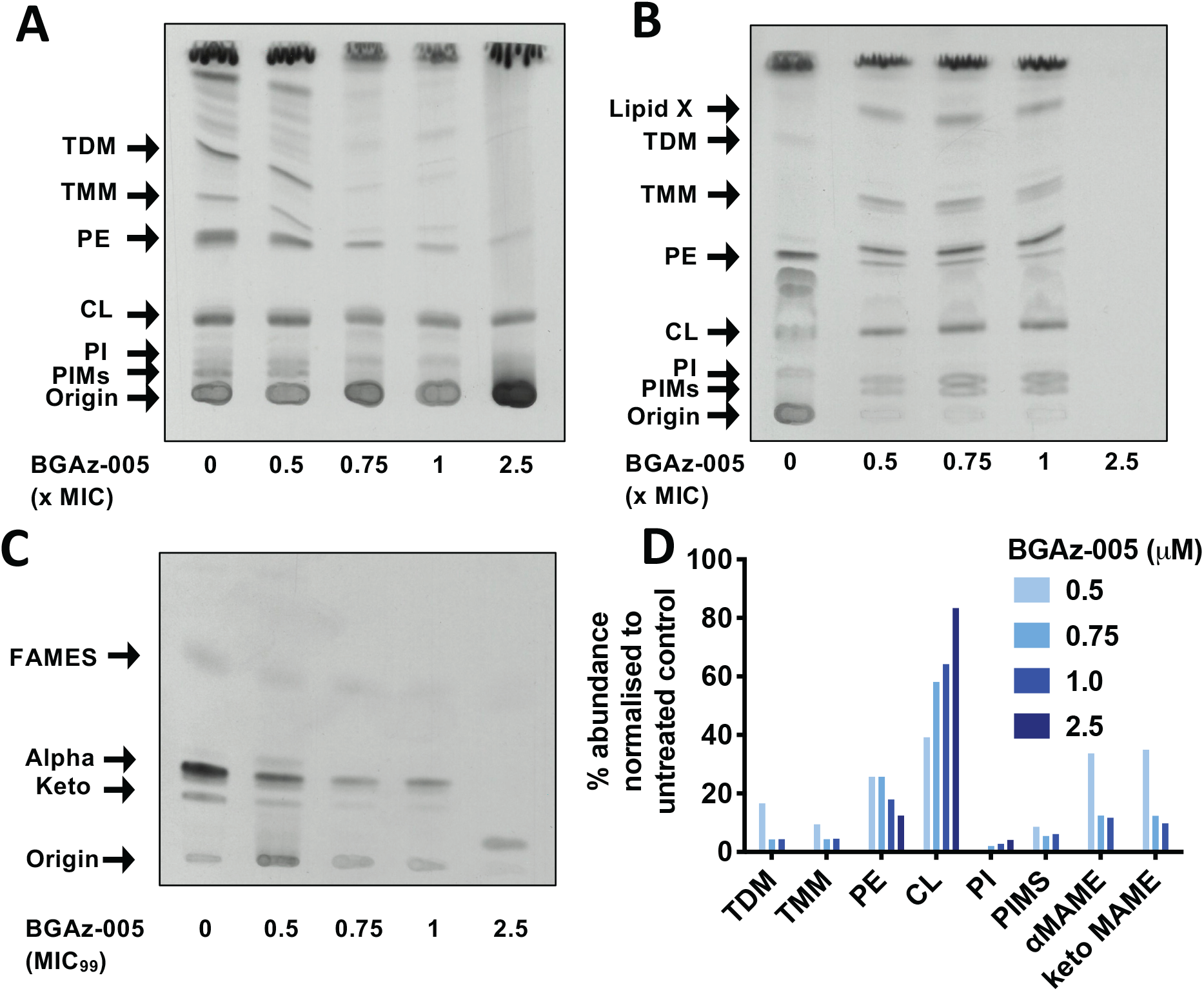
BCG cell envelope lipid analysis upon exposure to **BGAz-005**. BCG were cultured in 7H9 broth and exposed to increasing concentrations of **BGAz-005**. Lipids were selectively labelled with [^14^C]-acetic acid for 12 hours and cell envelope lipids were selectively removed by solvent extraction, separated by TLC (chloroform/methanol/water, 80:20:2, v/v/v), and visualised by autoradiography. **A** : equal volumes of lipids loaded adjusted for BCG growth; **B** : equal counts of lipids (25,000 cpm) loaded; **C** : Mycolic acid methyl ester (MAME) analysis of cell-wall bound mycolates released by 5% TBAH and separated by TLC (petroleum ether/acetone, 95:5, v/v); **D** : quantification of BCG lipids from panels A-C by densitometry.

To investigate the effect of **BGAz-005** on mycobacterial cell envelope composition and to identify the composition of Lipid-X, actively growing cultures of *M. smegmatis* were exposed to a range of **BGAz-005** concentrations, which resulted in a titratable-dependent reduction in the formation of TMM and TDM as observed by staining with MPA and α-naphthol (Figure 7A/B), consistent with [^14^C]-labelling experiments performed when **BGAz-005** was exposed to BCG (Figure 6). *M. smegmatis* exposed to the highest concentration of **BGAz-005**, resulted in a significant increase in the relative abundance of free mycolic acid (MA) within the cell envelope (Figure 7C); **BGAz004** affects mycobacterial cell envelope lipid biosynthesis in an almost identical manner (Supplementary Figure 4. The gradual reduction of TMM and TDM abundance in *M. smegmatis* and *M. bovis* BCG is a distinct observable phenotype that occurs upon exposure to **BGAz-005** (Figure 6 and Figure 7; and Supplementary Figures 3 and 4). The separation of solvent extractable lipids by two-dimensional TLC provides further confirmatory evidence that the increasing abundance of Lipid-X can be directly attributed to free MA.^45^ This large increase in free mycolic acid, paralleled with the loss of TMM, TDM and arabinogalactan linked mycolates illustrates that **BGAz-005** (and **BGAz-004**) compounds kill mycobacteria by arresting the final stages of mycolate biosynthesis.

**Figure 7.**
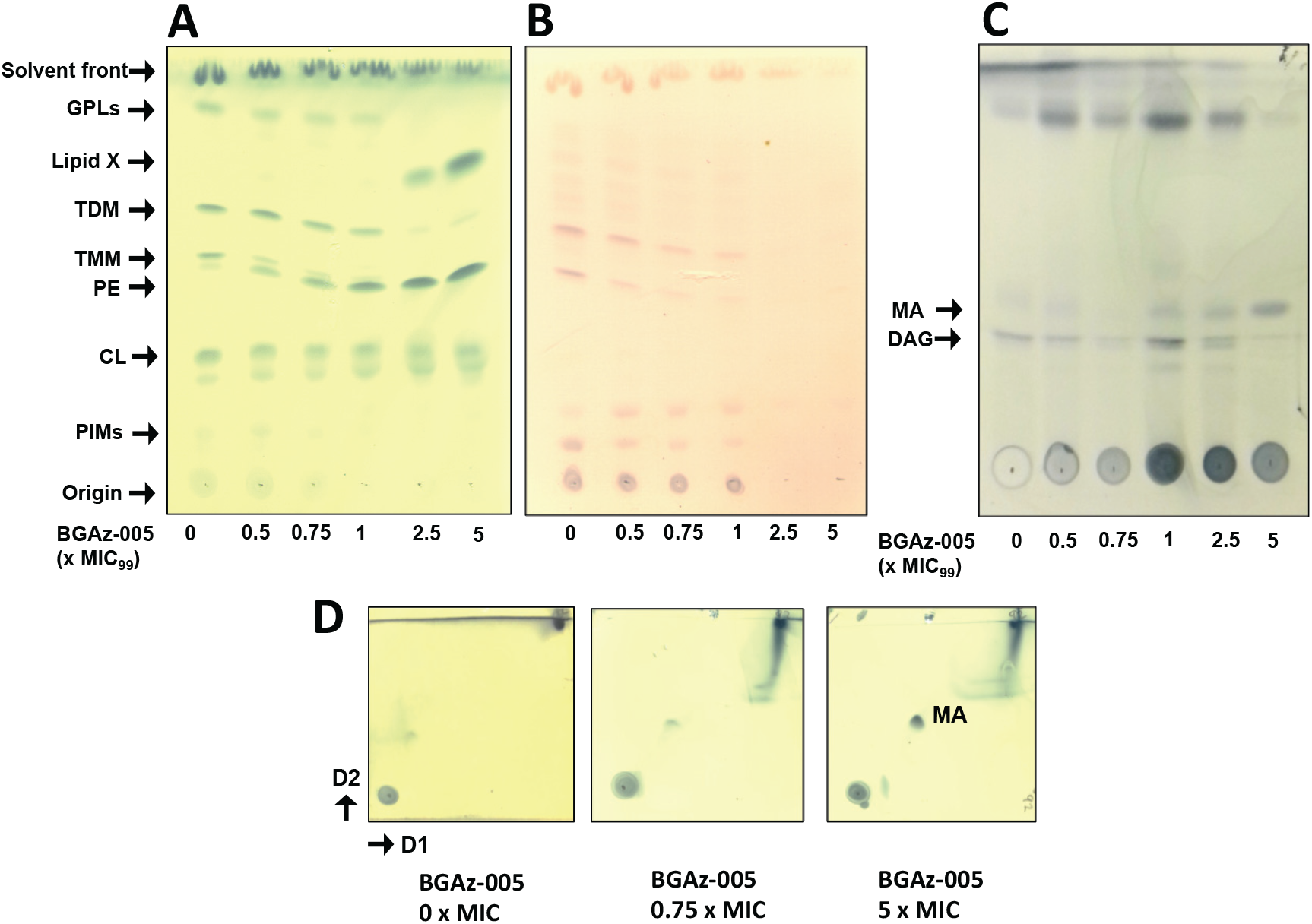
*M. smegmatis* cell envelope lipid analysis upon exposure to **BGAz-005**. *M. smegmatis* were cultured in 7H9 broth, exposed to increasing concentrations of **BGAz-005** for six hours and the cell envelope lipids selectively removed by solvent extraction. Equal volumes of lipid adjusted by bacterial growth were separated by TLC (chloroform/methanol/water, 80:20:2, v/v/v), and stained with MPA (**A**) or alpha-naphthol (**B**). Equal volumes of lipid adjusted by bacterial growth were separated by TLC (hexane/diethyl ether/acetic acid), 70:30:1, v/v/v and stained with MPA (**C**). Equal volumes of lipid adjusted by bacterial growth were separated by 2D-TLC (direction 1 chloroform/methanol 96:4, v/v, direction 2 toluene/acetone 80:20, v/v) and stained with MPA (**D**).

### BGAz-004 and BGAz-005 target late-stage mycolic acid biosynthesis enzymes

The inhibition of mycolate incorporation into the mycobacterial cell-wall, supported by transcriptomic profiling suggests that **BGAz-004** and **BGAz-005** act by targeting late-stage mycolate biosynthesis (Figure 5, Figure 6 and Figure 7). The accumulation of free mycolic acid upon exposure of **BGAz-004** and **BGAz-005** at the highest concentrations, suggests that mycolates are being formed but not deposited into the cell-wall (Figure 7 and Supplementary Figure 4). Pks13, MmpL3 and the Ag85 complex (FbpA, FbpB and FbpC) represent a selection of putative enzyme targets of mycolate biosynthesis that could be inhibited by **BGAz-004** and **BGAz-005**. Pks13 catalyses the last condensation reaction in mycolate biosynthesis, condensing two fatty acids to form mycolic acids.^38^ It also plays a role in TMM formation through acylation of trehalose. ^39^

Although essential in mycobacteria, *Corynebacterium glutamicum* can survive without mycolates,^38^ and so a Pks13 deletion mutant of *C. glutamicum* was utilised in this study (Supplementary Information). **BGAz-005** inhibits corynemycolic acid biosynthesis in *C. glutamicum* similarly to mycobacteria (Supplementary Figure 6. Both **BGAz-002** –**BGAz-005** retained similar levels of activity in *C. glutamicum*Δ*pks13* as in the wild-type strain, implying that Pks13 is not the target (Supplementary Table 2). MmpL3 is the essential membrane transporter responsible for translocating TMM across the cytoplasmic membrane.^37,40^ Treatment of *M. bovis* BCG harbouring an MmpL3 overexpression vector (pMV261A-*mmpL3* ^41^) with **BGAz-002** –**BGAz-005** resulted in no shift in the MIC_99_ compared to the empty vector (pMV261A) control (Supplementary Table 2). This negates MmpL3 as a potential target of the active BGAz compounds in this study, as overexpression of *mmpL3* should result in an increase in MIC as a result of increased copy number and target abundance. The Ag85 complex consists of three essential enzymes with mycolyltransferase activity, responsible for the formation of TMM, TDM, and the covalent attachment of mycolic acids to arabinogalactan.^42^ BCG harbouring the plasmids pTIC6-*fbpA*, pTIC6-*fbpB*, and pTIC6-*fbpC* overexpressing Ag85A, Ag85B and Ag85C, respectively, were treated with **BGAz-004** and **BGAz-005**. A statistically significantly increase in MIC was seen with FbpB and FbpC, but not FbpA (Figure 8). A two-fold increase in MIC was seen for both **BGAz-004** and **BGAz-005** upon overexpression of FbpB and FbpC, compared to the three-fold increase seen with known covalent inhibitor Ebselen.^46^ No shift in MIC was seen for FbpA upon Ebselen treatment. Furthermore, *M. smegmatis* cultured in the presence of Ebselen also resulted in a significant increase of free mycolic acid (MA) within the cell envelope and a concomitant loss of TMM and TDM (Supplementary Figure 5), mirroring the lipid profiles induced by **BGAz-004** and **BGAz-005** (Figure 7). Collectively, these findings point toward the antigen 85 enzymes as a possible target of the active BGAz compounds of this study.

**Figure 8.**
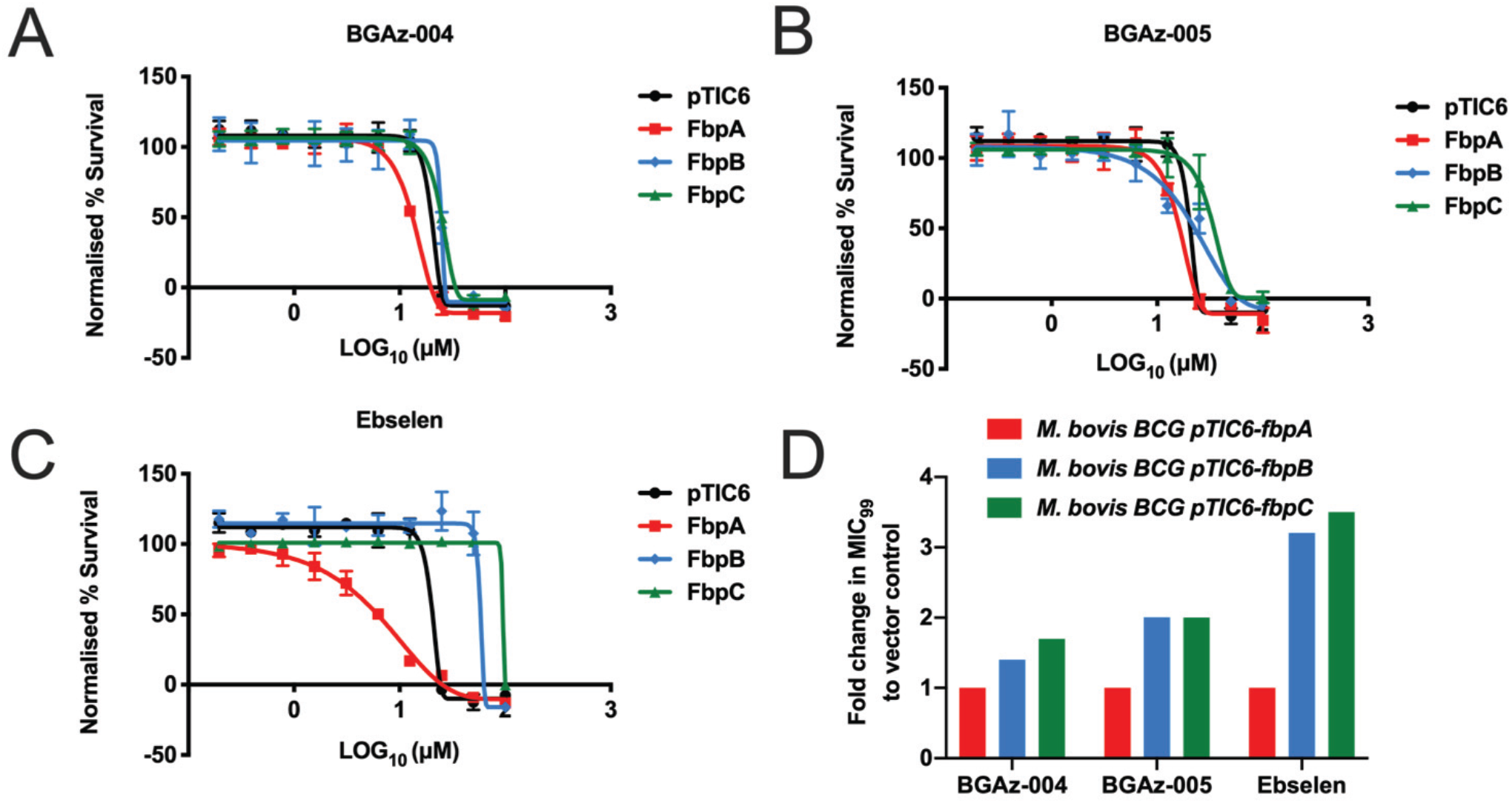
Assessing the MIC shift of **BGAz-004** and **BGAz-005** against the AG85 complex. MIC values of **BGAz-004**, **BGAz-005** and Ebselen were determined against BCG harbouring overexpression vectors and compared to empty vector controls (pTIC6) in order to identify a shift in MIC against fbpA (A), fbpB (B), fbpC (C). Fold change in MIC shift (D). The MIC_99_ was calculated using an endpoint resazurin assay and the Gomperz equation for MIC determination (GraphPad Prism). Data is of triplicate repeats.

## Discussion

The emergence of multi-drug-resistant TB means that new drugs to treat this disease are desperately required. Any new therapy should meet a number of parameters: it should be effective against MDR-TB; it should be rapidly bactericidal; it should show a novel mechanism of action; and it should possess ADME properties suitable for once a day oral dosing and co-administration with the current TB therapies and anti-HIV agents.^1^ **BGAz-002** –**BGAz-005** display potent inhibitory activity against different mycobacterial species (including virulent and avirulent *M. tuberculosis* reference strains), as well as drug-sensitive and drug-resistant (resistant to isoniazid, rifampicin, pyrazinamide and ethambutol) clinical isolates of *M. tuberculosis.* It is promising that **BGAz-002** –**BGAz-005** retain similar levels of activity between drug-sensitive and drug-resistant clinical isolates of *Mtb*; not only do these compounds target MDR-TB, but this also suggests that there is no cross-resistance with the current frontline drugs, indicative of a distinct mode of action. In addition to targeting an MDR strain of *Mtb*, the active BGAz compounds tested display no detectable resistance against mycobacteria in the laboratory, suggesting that development of clinical resistance to this class of compounds will be slow to occur. Similarly, teixobactin is a cyclic undecapeptide antibiotic that elicits bactericidal activity towards clinically relevant Gram-positive pathogens, also displaying an undetectable frequency of resistance. Teixobactin has a unique mode of action; by binding simultaneously to the cell-wall biosynthetic precursors lipid II and Lipid III, this antibiotic inhibits the biosynthesis of peptidoglycan and cell-wall teichoic acids, respectively.^47^

For each of **BGAz-002** –**BGAz-005**, the MBC is within four-fold of the MIC_99_, demonstrating their bactericidal nature in BCG. Further assessment in *M. tuberculosis* by time-kill viable counts and flow cytometry confirmed the bactericidal activity of **BGAz-004** and **BGAz-005**, with **BGAz-005** being significantly bactericidal after six days. The significantly earlier bactericidal activity of **BGAz-005** compared to **BGAz-004** could be attributed to the increased kinetic solubility of **BGAz-005**, or it could suggest that **BGAz-005** has additional targets. Whilst neither **BGAz-004** nor **BGAz-005** displayed bactericidal activity as rapidly as the current frontline drug isoniazid,^48^ the slower killing induced by the BGAz compounds tested may prove advantageous in preventing bacterial re-growth.^49^ Previous studies have shown that the rapid, early bactericidal activity of isoniazid results in bacterial regrowth, compared to no regrowth seen with the slower acting bactericidal drugs rifampicin and pyrazinamide.^2^ The longer-acting bactericidal activity of the BGAz compounds tested, combined with the absence of any detectable generation of resistance, suggests that they may be superior to isoniazid, as the potential for tolerance and resistance are very low.

Comprehensive mode of action studies support the hypothesis that the BGAz compounds target mycobacterial cell-wall biosynthesis. Radiolabelled precursor incorporation studies demonstrate that **BGAz-005** specifically arrests peptidoglycan and lipid biosynthesis, and transcriptome signatures of **BGAz-004-** and **BGAz-005**-treated *M. bovis* BCG reveal significant alterations in cell-wall biosynthetic genes. The mycobacterial cell-wall is a well validated and commonly occurring target amongst anti-tubercular drugs, including frontline drugs isoniazid^3^ and ethambutol,^4^ as well as ethionamide,^3^ SQ109,^5^ and D-cylcoserine.^6^ Whilst the BGAz compounds tested in this study also inhibit cell-wall biosynthesis, they do so without displaying target redundancy against current front line drugs, such as isoniazid. Transcriptomic analysis revealed marked differences between the **BGAz-004**, **BGAz-005**, isoniazid and ethambutol, specifically the downregulation of mycolic acid synthesis genes of the FasII system and mycolic acid synthesis and modification genes, and the lack of induction of the *efpA* efflux pump. Compared to other genome-wide transcriptional studies of anti-tubercular drugs,^50^ the observed differences induced by BGAz compounds suggest that they possess a unique mode of action compared to current chemotherapeutic agents.

To further probe the mechanisms by which the BGAz compounds perturb the mycobacterial cell envelope, the lipid profiles of mycobacteria exposed to increasing concentrations of **BGAz-005** were examined. A specific and rapid depletion in TMM and TDM was seen upon BGAz treatment of both *M. smegmatis* and *M. bovis* BCG, whilst other lipids such as cardiolipin and PIMs remained constant. The effects are more pronounced in BCG due to its increased sensitivity of radiolabelling, but in both instances, there is an almost complete arrest in TMM and TDM production by 1 × MIC of **BGAz-005**. Analysis of the cell-wall bound mycolates revealed a concurrent depletion in MAMEs. The specific loss in mycolates, both non-covalently (TMM and TDM) and covalently (MAMEs) associated, indicates that the BGAz compounds tested target mycolic acid biosynthesis. Mycolates are an essential component of the mycobacterial cell envelope and are targeted by the current drugs isoniazid and ethionamide.^51^ That the BGAz compounds tested target the same biosynthetic pathway as isoniazid corroborates with the flow cytometry analysis, where the staining profiles of **BGAz-004** and **BGAz-005** were like those of isoniazid, suggesting a similar target. Further cell envelope analysis revealed a pronounced increase in free mycolic acid correlating with increasing concentration of **BGAz-005**. Typically, the relative abundance of free MA in the cell envelope of planktonically cultured mycobacteria is extremely low. However, previous studies have demonstrated that free MA levels increase significantly when mycobacteria are cultured as a pellicle biofilms^52^ or as non-replicating populations induced by gradual nutrient starvation.^53^ This simultaneous depletion of mycolates conjugated to trehalose and AG, alongside an accumulation of free mycolic acid implies that the mycolates are being synthesised (demonstrated by the accumulation of free mycolate), but are not incorporated into the cell-wall (demonstrated by the loss of TMM, TDM and MAMEs). Thus, unlike isoniazid, the BGAz compounds tested target late-stage mycolic acid biosynthesis and have a mode of action distinct to that of isoniazid and Ethionamide, which target the early stages of mycolate production.^3^ There are several enzyme candidates involved in the latter stages of mycolic acid biosynthesis and incorporation into the cell envelope. Specifically, These encode the polyketide synthase (Pks13) responsible for the last condensation reaction in mycolate biosynthesis,^7^ the essential membrane transporter responsible for translocating TMM across the cytoplasmic membrane (MmpL3),^8,9^ and the mycolyltransferase responsible for the formation of TMM, TDM, and the covalent attachment of mycolic acids to arabinogalactan by the antigen 85 complex enzymes (FbpA, FbpB and FbpC),^10^ respectively. Target engagement over-expression studies ruled out MmpL3 as a BGAz target, while experiments conducted in *C. glutamicum* demonstrate that Pks13 is equally not inhibited by these compounds. Guided by evidence obtained from transcriptomic profiling and cell envelope lipid analysis, further target engagement over-expression studies revealed that FbpB and FpbC afforded moderate protection to BCG exposed to **BGAz-004** or **BGAz-005**. Previous studies have identified FbpA, FbpB and FpbC as druggable targets for the development of new anti-tubercular agents,^46,54–56^ however many of these agents display unfavourable toxicological and PK/PD profiles. In this regard, the BGAz compound series demonstrates an encouraging overall toxicological profile with good absorption, low mitochondrial toxicity and rapid clearance from hepatocytes. In order to reduce the potential of compound-related risk factors a selection of mechanistic screening assays was utilised to identify hazardous and undesirable chemistry in this study. A critical example of such compound liabilities is the blocking of hERG potassium channel. The hERG channel is a voltage-gated potassium channel that is expressed in a variety of human tissues such as the brain, thymus, adrenal gland, retina and in cardiac tissue. Any significant blocking of hERG channels by potential drug candidates could have serious off-target effects due to the dysregulation of action potential repolarisation. Furthermore, **BGAz-005** displays a remarkable pharmacokinetic profile in mice, with a blood plasma half-life exceeding three days that exceeds the C_max_ required to reach a therapeutic dose for MDR-TB. Overall, the results presented here demonstrate that the BGAz compound series represents promising novel anti-tubercular agents for further development towards the clinic.

## Methods

### Synthesis of BGAz-001–BGAz-005

Novel racemic 2,4-*cis*-amino-azetidine derivatives (**BGAz-001** –**BGAz-005**) were prepared and purified according to procedures reported for the synthesis of related compounds, for details see supplementary material (Supplementary Scheme 1).^16–19^ Briefly, commercially available aldehydes (**S1**) and amines (**S2**) were dissolved in methanol and heated at reflux to afford the corresponding imines (**S3**).^57–59^ Imines (**S3**) were isolated and subsequently reacted with *in situ* prepared allyl zinc reagent to afford homoallyl amine derivatives thereof (**S4**). The homoallyl amine derivatives (**S4**) thus obtained were dissolved in acetonitrile and treated with iodine (3 equiv.) and sodium bicarbonate (5 equiv.) at temperatures not exceeding 20 °C resulting in cyclisation to the corresponding 2-iodomethyl azetidine derivatives (**S5**). Displacement of iodine in derivatives **S5** by the appropriate primary or secondary amines delivered the BGAz series of compounds in four linear steps.

### Bacterial strains and growth conditions

*M. smegmatis* MC^2^155 was cultured at 37 °C, 180 rpm in Middlebrook 7H9 media supplemented with 0.05% Tween-80, or grown on LB agar. *M. bovis* BCG (strain Pasteur) was cultured at 37 °C and 5% CO_2_, static, in Middlebrook 7H9 media supplemented with 0.05% Tween-80 and 10% (v/v) BBL Middlebrook OADC enrichment or grown on Middlebrook 7H11 agar supplemented with 10% (v/v) BBL Middlebrook OADC enrichment.

### Determination of MIC and MBC

The minimum inhibitory concentration (MIC_99_) was determined in 96-well flat bottom, black polystyrene microtiter plates (Greiner) in a final volume of 200 μL Compounds were two-fold serially diluted in neat DMSO and added to the microtiter plate at a final concentration of 1% DMSO. DMSO (1% in 7H9) was used as a positive control and rifampicin as a negative. The inoculum was standardised at OD_600_ 0.05 in Middlebrook 7H9 medium and added to the plate, which was then incubated without shaking at 37 °C for 24 hours (*M. smegmatis*) or seven days (*M. bovis* BCG). Following incubation, 42 μL of resazurin (0.02% v/v in dH_2_O) was added to each well and incubated for a further two hours (*M. smegmatis*) or 24 h (*M. bovis* BCG). Fluorescence was measured (Polar star omega plate reader ex 544 nm, em 590 nm) and the data normalised using equation one. The concentration of drug required to inhibit cell growth by 99% was calculated by non-linear regression (Gomperz equation for MIC determination, GraphPad Prism).

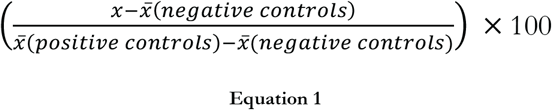

To determine the minimum bactericidal concentration (MBC), *M. smegmatis* and *M. bovis* BCG were grown in the presence of a two-fold serial dilution of compound as above. After a 24-hour (*M. smegmatis*) or seven-day (*M. bovis* BCG) incubation, the cells were pelleted and washed with phosphate buffered saline pH 7.2 (PBS). The washed cells were plated onto agar, devoid of compound, and incubated for four days (*M. smegmatis*) or 21 days *M. bovis* (BCG). The MBC was defined as the lowest concentration of compound for which there was no bacterial growth.

### Determination of MIC_lux50_ against autoluminescent M.tb H37Ra

AlRa^60^ (*Mtb* H37Ra::pTYOK) was homogenised with sterile glass beads in a 50 mL tube containing Middlebook 7H9 medium (5 mL) plus 0.05% Tween 80, 10% v/v oleic acid albumin dextrose catalase (OADC) supplement (7H9-OADC-Tw). When OD_600_ reached 0.3-0.5, relative light unit (RLU) count was determined by placing culture (200 μL) on the detection hole of the luminometer. When the RLU reached 2 million/mL, the activities of compounds were assessed over a range of three-fold increasing from 0.000001 μg/mL to 10 μg/mL prepared in 25 μL AlRa broth culture (RLU diluted to 2000-4000/25 μL) grown in 7H9 broth without Tween 80. DMSO was used as negative control and isoniazid (INH, 10 μg/mL, 1 μg/mL and 0.1 μg/mL) and rifampicin (RIF, 10 μg/mL, 1 μg/mL and 0.1 μg/mL) were used as positive controls. RLU counts were determined daily, for four times (daily, i.e. day 0, 1, 2 and 3). The MIClux50 was defined as determined as the lowest concentration that can inhibit > 50% RLUs compared with that from the untreated controls at day three.^61^

### Determination of MIC_99_ for clinical M.tb strains

The previously described resazurin microtiter assay (REMA) plate method was used.^62,63^ Compounds were two-fold serially diluted in CAMR Mycobacterium Medium MOD2 (CMM MOD2).^64^ Individual wells, in a 96-well plate, were inoculated with 1 × 10^6^ CFUmL^−1^ bacilli and incubated for seven days at 37 °C with agitation (200 rpm). Following this, resazurin solution was added to wells (0.02% (w/v) in PBS pH 7.4, supplemented with 5% Tween 80).^65^ The 96-well plates were incubated at room temperature for six hours. The OD_570nm_ of each well was recorded using a Tecan Sunrise plate reader. The minimum inhibitory concentration (MIC_99_) was calculated using a modified Gompertz function.^66^ The optical density measurements for each drug concentration was compared to vehicle control to determine the percentage reduction in bacterial optical density.

### Physiochemical and toxicological analysis

#### Kinetic Solubility

Compounds were solubilised (10 mM in DMSO) and diluted in phosphate buffered saline (pH 7.4) into a seven-point curve (0.2–100 μM) and incubated for five minutes at 25 °C with shaking (final DMSO concentration 1%). The turbidimetry was assessed at each of the seven concentrations using UV spectrophotometry at 620 nm, and the LogS was converted into the estimated solubility (S) using the equation S = 10^LogS^. Nicardipine hydrochloride was used as a control compound. All experiments were performed in triplicate.

#### Mouse PPB

Compounds were solubilised (10 mM in DMSO) and Rapid Equilibrium Dialysis (RED) was used to measure the percentage binding to mouse plasma protein of the BGAz compounds at a final concentration of 5 μM. The BGAz compound was incubated in 100% mouse plasma and dialysed against buffer in a RED device for four hours at 37 °C in a 5% CO_2_ incubator, with continuous shaking at 200 rpm. Samples were matrix matched and analysed by LC-MS/MS against a six-point standard curve prepared with 100% plasma. All experiments were performed in triplicate.

#### Mouse Microsomal Clearance

Compounds were solubilised (10 mM in DMSO). 1 μM of BGAz compound was incubated with 0.5 mg/mL mouse microsomes in the presence or absence of the Phase 1 cofactor NADPH (1 mM) at 37 °C for 0, 5, 10, 15, and 30 minutes. The disappearance of the BGAz compound was assessed by LC-MS/MS. All experiments were performed with two replicates per compound and were validated by the inclusion of up to three species-specific control compounds. Data output consists of mean intrinsic clearance (CL_int_) and half-life (t_1/2_) measurements.

#### Mouse Hepatocyte Clearance

Compounds were solubilised (10 mM in DMSO). 1 μM of BGAz compound was incubated with 0.5 × 10^6^ cells/mL mouse hepatocytes at 37 °C for 0, 10, 20, 30, 45 and 60 minutes. The disappearance of the BGAz compound in the presence and absence of hepatocytes was assessed using LC-MS/MS. All experiments were performed with two replicates per compound and were validated by the inclusion of up to three species-specific control compounds. Data output consists of mean intrinsic clearance (CL_int_) and half-life (t_1/2_) measurements.

#### Caco-2 and efflux

Compounds were solubilised (10 mM in DMSO) and the CacoReady™ Kit from ReadyCell S.L. (Barcelona, Spain) was used to determine compound permeability. Differentiated and polarized Caco-2 cells (21-day system) were plated on a 96-transwell permeable system as a single monolayer to allow for automated high throughput screening of compounds, and 10 μM BGAz compound was added to the system in HBSS buffer (pH 7.4) and incubated for two hours at 37 °C in a CO_2_ incubator. Lucifer yellow was used as a cell monolayer integrity marker. Drug transport was assessed in both directions (apical to basolateral (A-B) and basolateral to apical (B-A)) across the cell monolayer. The buffer used for the assay does not include HEPES, so as to minimise the inhibitory effect on uptake transporters.^67^ The BGAz compound concentrations were quantified using a calibration curve following analysis by LC-MS/MS, and the apparent permeability coefficient (P_app_) and efflux ratio of the compound across the monolayer were calculated. The efflux ratio is used as an indicator of active efflux. The permeability coefficient (*P_app_*) was calculated from the following equation:

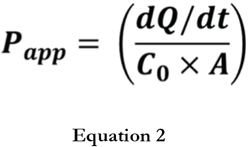

Where dQ/dt is the amount of compound in basal (A-B) or apical (B-A) compartment as a function of time (nmol/s). C_0_ is the initial concentration in the donor (apical or basal) compartment (Mean of T=0) (nmol/mL) and A is the area of the transwell (cm^2^).

The efflux ratio was then calculated as:

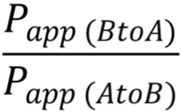

All experiments were performed in triplicate and the MDR1 efflux markers Digoxin, quinidine and propranolol were used as positive controls.

#### Pharmacokinetic studies

Methods for combined single dosing (BGAz001–005) and dosing of single compounds multiple times (BGAz004 and BGAz005) are described in detail in a supplementary file (Supplementary PK Data).

#### Cytochrome P450 activities

Compounds were solubilised to 10 mM in DMSO. The BGAz compounds were incubated at concentrations of 0.003, 0.009, 0.03, 0.08, 0.25, 0.74, 2.2, 6.7 and 20 μM with CYP1A2, CYP2C9, CYP2C19 and CYP2D6 at 37 °C in the presence of the drug-like probe substrate HLM the Phase 1 cofactor NADPH (1 mM). The formation of metabolites of the drug-like probe substrates in the absence and presence of BGAz compound was monitored by LC-MS/MS, and the IC_50_ value determined. All assays had two replicates per compound and included a positive control inhibitor.

#### HepG2 mitochondrial dysfunction

Compounds were solubilised to 30 mM in DMSO. The BGAz compounds were added to the HepG2 cell model in a 96-well microplate in half log dilutions from 100 μM–0.0003 μM (final DMSO concentration 0.3%) using both glucose (DMEM consisting of 25 mM glucose) and galactose (DMEM consisting of 10 mM galactose) media. The compounds were incubated with the cell line or 24 hours at 37 °C in a humidified CO_2_ tissue culture incubator, followed by cell viability staining with MTT (3-(4,5-dimethylthiazol-2-yl)-2.5-diphenyltetrazolium bromide) conversion to Formazan product, determined by absorbance measurement. The cell viability IC_50_ was determined in HepG2 glucose and metabolism modified HepG2 galactose, and the fold change difference between the Glu/Gal IC_50_ was determined. All experiments were performed in duplicate with the mitochondrial toxicity controls rotenone and Antimycin A, and the cytotoxin control tamoxifen.

#### hERG cardiotoxicity function

Compounds were solubilised to 30 mM in DMSO before dilution in PBS to 300 mM. A further three-fold on-board dilution resulted in a final top BGAz compound concentration of 100 mM. Eight-point concentration-response curves were generated using 3.16-fold serial dilutions from the top test concentration. Electrophysiological recordings were made from a Chinese Hamster Ovary cell line stably expressing the full length hERG channel. Single cell ionic currents were measured in the perforated patch clamp configuration (100 μgmL^−1^ amphotericin) at room temperature (21–23 °C) using an IonWorks Quattro instrument (Molecular Devices). The internal solution contained (mM): 140 KCl, 1 MgCl_2_, 1 EGTA, 20 HEPES and was buffered to pH 7.3. The external solution (phosphate-buffered saline, PBS) contained (mM): 138 NaCl, 2.7 KCl, 0.9 CaCl_2_, 0.5 MgCl_2_, 8 Na_2_HPO_4_, 1.5 KH_2_PO_4_ buffered to pH 7.4. Cells were clamped at a holding potential of −70 mV for 30 s and then stepped to +40 mV for one second. This was followed by a hyperpolarising step of one second to −30 mV to evoke the hERG tail current. Currents were measured from the tail step and referenced to the holding current. Compounds were then incubated for 3– 4 minutes prior to a second measurement of the hERG signal using an identical pulsetrain.

### Mycobacterial time kill experiments

**BGAz-004**, **BGAz-005** and isoniazid were two-fold serially diluted in 100 μL CMM MOD2 medium from 96–3 μM, 96–3 μM and 29.2–0.9 μM respectively, vehicle control (0.1% DMSO) was included in all experimentation. Individual wells of a 96-well micro-titre plate were inoculated to a starting bacterial titre of 1 × 10^6^ CFUmL^−1^. Micro-titre plates were incubated for 0, 2, 6, 10 and 14 days at 37 °C with agitation (200 rpm). The bacterial titre at each time point was enumerated *via* outgrowth of bacilli on solid media for three weeks at 37 °C *via* a method adapted from Miles and Misra^26^ where triplicate 20 μL spots of bacterial culture are spotted onto Middlebrook 7H10 agar for each ten-fold serial dilution. Statistical analyses of data were performed using a factorial ANOVA and post-hoc Tukey’s honestly significant difference test.

### Bacterial staining and flow cytometry analyses

The method reported by Hendon-Dunn *et al.* 2016^27^ was used to analyse stained cell populations by flow cytometry 100 μL of *M. tuberculosis* H37Rv from each antibiotic-incubation, at each time-point, was transferred to a micro-titre plate in quadruplicate and incubated in the dark for one hour at 37 °C with either no dye added, 20 μM calcein violet (CV-AM), 20 μM sytox green (SG), or both 20 μM CV-AM and 20 μM SG. These incubations were then spun by centrifugation and the supernatant was removed. The cells were fixed with 4% formaldehyde (v/v in water) for one hour. The stained bacteria were examined using a Cytoflex S (Beckman Coulter) flow cytometer. Lasers with excitatory wavelengths of 488 nm and 405 nm were used. SG fluorescence emission (excitation and emission, 488 nm and 523 nm, respectively) was detected in the FIT-C channel (530/40 BP), and CV-AM fluorescence (excitation and emission, 400 nm and 452 nm, respectively) was detected in the PB-450 channel (450/50 BP). A quadrant gating strategy was used,^27^ briefly, 10,000 single-cells events were gated upon using a two-parameter dot plot of forward scatter height *versus* forward scatter area. From gated single-cells, the percentages of the total cell population residing in each polygonal population gate (P1: CV-AM^−^/SG^−^, P2: CV-AM^+^/SG^−^, P3: CV-AM^+^/SG^+^, P4: CV-AM^−^/SG^+^) were obtained. Statistical analysis of data was performed using a Student’s *t* test.

### Inhibition of Macromolecular Synthesis

Inhibition of macromolecular biosynthesis was assayed by measuring the incorporation of radiolabelled precursors into DNA, RNA, protein, peptidoglycan and fatty acids in the 10% trichloroacetic acid (TCA) extracts of cells exposed to azetidines. *M. smegmatis* were cultured in 7H9 media supplemented with 0.05% Tween-80 and grown to an OD_600_ of 0.4. Cultures (5 mL) were then transferred into sterile glass tubes and preincubated with 0 ×, 0.5 ×, 0.75 × and 1 × MIC_99_ azetidines for one hour at 37 °C with shaking. After preincubation, 10 μL of 500 nCi/μL [*methyl-*^3^H]thymidine, 500 nCi/μL [5,6-^3^H]uridine, 500 nCi/μL L-[4,5-^3^H]leucine, 500 nCi/μL [^3^H]*meso-*diaminopimelic acid and 500 nCi/μL [^14^C]acetic acid were added to cultures to measure synthesis of DNA, RNA, protein, peptidoglycan and lipids, respectively. All cultures were incubated for 36 hours and 100 μL samples were sacrificed at 6 h, 12 h, 24 h and 36 h time points by the addition of 50 μL 30%TCA/70% ethanol in Eppendorf tubes. Tubes were incubated at room temperature for 60 min to allow for precipitation of macromolecular material. Samples were individually vacuum filtered using 0.025 μm membrane filters (VSWP01300, MF-Millipore) that were prewashed with 500 μL 70% ethanol. Samples were washed with 3 × 500 μL 5% TCA followed by 2 × 95% ethanol. Filter papers were dried and combined with 5 mL scintillation fluid before measuring radioactive counts.

### Transcriptomic profiling by RNA-seq analysis

*M. bovis* BCG was cultured to an OD_600_ of 0.4 before exposure to 1 × MIC_99_ concentrations of **BGAz-004** or **BGAz-005** for eight hours in three biological replicates, then compared to carrier control-treated bacilli. Cells were pelleted, flash frozen in liquid nitrogen and stored at −80 °C. Pellets were resuspended in lysozyme (600 μL, 5 mg/mL) and β-mercaptoethanol (7 μL/mL) in TE buffer and lysed by bead beating at 6 m/minute (1 × 45 seconds). Samples were subjected to further bead beating (3 × 45 seconds) following the addition of (60 μL, 10% SDS). Sodium acetate pH 5.2 (3 M, 60 μL) and acidified phenol pH 4.2 (726 μL) were added and the tubes mixed well by inversion. Samples were incubated at 65 °C for five minutes and centrifuged for five minutes at 18,000 ×*g*. The upper aqueous phase was transferred to a fresh tube and an equal volume of acid phenol pH 4.2 added and mixed well by inversion. Following heating (65 °C for two minutes) and centrifugation (five minutes at 18,000 ×*g*), the upper aqueous phase was once more transferred to a fresh tube and an equal volume of chloroform:isoamyl alcohol (24:1 v/v) added. The sample was mixed well by inversion and centrifuged at 18,000 ×*g* for five minutes. The upper aqueous phase was transferred to a fresh tube and a 1/10 volume of sodium acetate (3 M, pH 5.2) and three volumes of 100% ethanol were added. Samples were incubated at −20 °C overnight, centrifuged for ten minutes (4 °C, 14,000 ×*g*) and the supernatant removed. Ethanol (70% in water, 500 μL) was added to the pellet and centrifuged for ten minutes (4 °C, 14,000 ×*g*). The supernatant was removed, the pellet air-dried, and the extracted RNA resuspended in of RNase-free dH_2_O (40 μL). DNase treatment was performed using the TURBO DNA-*free* kit (Invitrogen). Briefly, a 0.1 × volume of 10 × TURBO DNase buffer was added to the RNA, along with TURBO DNase enzyme (1 μL of enzyme stock). The sample was incubated for 30 minutes at 37 °C, an additional TURBO DNase enzyme (1 μL of enzyme) was added, and the sample incubated for another 30 minutes. DNase inactivation reagent (0.2 volumes) were added, and the sample incubated at room temperature for five minutes. Following centrifugation at 10,000 ×*g* for 1.5 minutes, the supernatant containing the RNA was transferred to a fresh tube and stored at −80 °C. The purified RNA was quantified, depleted of rRNA and library-prepped before sequencing by Illumina HiSeq (150 × 2 paired end) by Genewiz Ltd. Adapter sequences and poor-quality reads were removed using Trimmomatic v.0.36, before mapping to the *Mycobacterium bovis* BCG Pasteur 1173P2 genome using Bowtie2 aligner v.2.2.6. Gene expression was quantified using FeatureCounts from the Subread package v.1.5.2. Differentially expressed genes were identified with the DESeq2 R package, normalised by RLE method, and using the Wald test with Benjamini and Hochberg multiple testing correction. Genes with an adjusted p-value <0.05 and log2 fold change >1 were considered to be differentially expressed. Significantly enriched signatures, with updated genome annotation,^68,69^ were identified using the hypergeometric function comparing to published drug responses^40,41^ or mapped to metabolic pathways.^34,35^ Genes significantly differentially expressed in response to **BGAz-004** and **BGAz-005** are detailed in Supplementary Material. Fully annotated RNA-seq data will be deposited in ArrayExpress; accession number to be provided.

### Radioisotope labelling of lipids and analysis

Cells were grown to an OD_600_ 0.5, treated with compound and grown for six hours (*M. smegmatis*) or overnight (*M. bovis* BCG). For radio labelling experiments, 1 μCi mL^−1^ ^14^C acetic acid was then added followed by a 16-hour incubation. Cells were harvested and extracted using chloroform:methanol:water (10:10:3, v/v/v, 2 mL) for two hours at 50 °C. Following centrifugation, the organic extracts were combined with chloroform and water (1.75 mL and 0.75 mL respectively). The lower organic phase containing associated lipids was recovered, washed twice with chloroform:methanol:water (3:47:48, v/v/v, 2 mL) and dried with nitrogen gas. Samples were resuspended in chloroform:methanol (2:1, v/v, 200 μL) and OD adjusted volumes were subjected to thin-layer chromatography (TLC) analysis. Cell-wall associated lipids were visualised by either heating TLC plates after treatment with molybdophosphoric acid (MPA) in ethanol (5% w/v) or alpha-naphthol in ethanol (5% w/v), or by autoradiography by exposure of Kodak BioMax MR film.

Cell-wall bound lipids from the de-lipidated extracts from the above extraction were released by the addition of a solution of tetra-butyl ammonium hydroxide (TBAH) (5% m/v, 2 mL) followed by a 16-hour incubation at 100 °C. Water (2 mL), dichloromethane (4 mL) and iodomethane (200 μL) were then added and mixed thoroughly for 30 min. The organic phase was recovered following centrifugation and washed with water (3 × 4 mL), dried and resuspended in diethyl ether (4 mL). After sonication and centrifugation, the supernatant was dried and resuspended in dichloromethane. Equivalent aliquots of the samples were subjected to TLC in petroleum ether:acetone (95:5, v/v) and visualised by MPA and heat or autoradiography.

### Target gene overexpression studies

The constructs pMV261_*mmpL3* and pVV16_*trpAB*,^70,71^ including the empty pMV261 and pVV16 vectors, were electroporated into *M. bovis* BCG as previously described, and the MIC_99_ determined as described above. The constructs pTIC6_*fbpA,* pTIC6_*fbpB* and pTIC6_*fbpC* were synthesised by GenScript Ltd by inserting the coding regions of the *M. tuberculosis* H37Rv genes into the vector pTIC6, which encodes kanamycin selection. The constructs and empty pTIC6 vector were electroporated into *M. bovis* BCG. Following induction of gene expression with 50 ng/mL of anhydrotetracycline for 24 hours, the MIC_99_ was determined as described above.

## Supplementary material

A supplementary file containing a detailed description of author contributions, general methods, chemical and biological general procedures, synthetic chemistry protocols for the synthesis of **BGAz001** –**BGAz005**, NMR spectrums, details of mass spectrometry analysis and additional corresponding references.^72,73^

## Acknowledgements

LJA and JSF are grateful for the support underpinning much of this study from MRC Confidence in Concepts and EPSRC follow-on fund schemes. The University of Birmingham are thanked for support, including travel funds permitting AY, XL and YC to undertake training placements at GIBH. JSF is grateful to the Royal Society for training provided as a result of a previous Industrial Fellowship and the EPSRC for previous funding (EP/J003220/1). Funding for part of this study was received from Department of Health Grant in Aid. This work was supported by the National Mega-project of China for Innovative Drugs (2019ZX09721001-003-003) and by the Chinese Academy of Sciences Grant (154144KYSB20190005). TZ received a Science and Technology Innovation Leader of Guangdong Province (2016TX03R095) award. SJW and AG thank the National Centre for the Replacement, Refinement and Reduction of Animals in Research (NC3Rs) for grant support (NC/R001669/1). Qiong Pan (GIBH), Jingfang Xiong (GIBH) and Miaoqin She (GIBH) are thanked for conducting aspects of the PK/PD studies of this report. Dr Chi Tsang (UoB), Dr Peter Ashton (UoB) and Jiajia Wei (GIBH) are thanked for helpful discussions and practical support with aspects of mass spectrometry. Dr Cécile S. Le Duff (UoB) and Dr Neil Spencer (UoB) gave advice on aspects of NMR spectroscopy underpinning the preliminary or previously reported, findings. Yingxue Liu (GIBH) is thanked for help with purification of final products by HPLC where required.

## Competing interests

A patent application disclosing aspects of this study has been filed by the University of Birmingham. The views expressed in this publication are those of the authors and not necessarily those of Public Health England, or the Department of Health. The authors declare no other competing interests.

## Author contributions

### Corresponding authors

LJA Biological aspects including mode of action; JSF Synthetic chemistry aspects including azetidine design and synthesis; CN Medicinal chemistry aspects; JB assessments of compound activity against *M. tuberculosis*; SW analysis of RNA seq data and transcriptomic profiling.

### Joint first authors

AL examined biological activity, determined MIC values, probed target/mechanism and elucidated aspects of the mode of action; YC contributed to development of methodology for the synthesis of azetidine derivatives and transferred knowledge between teams. All authors contributed critically to devising and executing aspects of this research. A detailed and comprehensive description of author contributions is defined in the associated supplementary material.

## Notes

### Competing Interest Statement

The authors have declared no competing interest.

