## Supplementary Information for "Azetidines kill *Mycobacterium tuberculosis* without detectable resistance by blocking mycolate assembly"

#### Detailed author contributions

**Contributions in author order: AL** examined anti-TB activity, determined MIC values, probed target/mechanism and elucidated aspects of the mode of action, and wrote sections of the manuscript; **YC** applied and optimised developed methodology for the synthesis of optimised active azetidine derivatives, effectively transferred knowledge across the team *via* secondment visit and co-wrote aspects; **EB** conducted and analysed DMPK experiments and data; **NJCo** contributed and suggested aspects of biological screening around ADME, gave comment and revision of manuscript; **PGEC** contributed to synthetic target selection and suggested experiments; **NJCu** applied developed methodology to synthesise active azetidine derivatives;

**KD** examined biological activity and determined MIC values; **AF** uncovered enabling methodology for the synthesis azetidine derivatives and established protocols underpinning this study; **JF** contribution to the conception of the work, project management and revision of the manuscript; **AG** undertook transcriptomic analysis; **MAHS** applied developed methodology to synthesise active azetidine derivatives; **XLang** contributed to developing methodology applied to the synthesis of azetidine derivatives screened; **XLi** applied developed methodology to synthesise active azetidine derivatives and effectively transferred knowledge across the team *via* secondment visit; **CWM** evaluated activity of azetidine derivatives against MDR-TB and determined MIC values; **MM** devised and conducted hERG liability in vitro screening; **JP** evaluated activity of azetidine derivatives against MDR-TB and determined MIC values; **XP** contributed to developing methodology applied to the synthesis of azetidine derivatives screened; **VP** conducted and analysed DMPK experiments and data; **CP** examined biological activity and determined MIC values; **TS-U** conducted and analysed DMPK experiments and data; **MT** oversight of aspects of the project and commented on progress and decision points; **ZT** biological investigation of activity of a selection of compounds using a fluorescence-based assay; **ZEU** evaluated activity of azetidine derivatives against MDR-TB and determined MIC values; **CW** biological investigation of activity of a selection of compounds using a fluorescence-based assay; **AY** Synthesised first biologically active compound identified as active in this study. Applied and optimised developed methodology for the synthesis of azetidine derivatives and effectively transferred knowledge across the team *via* secondment visit; **TZ** biological investigation of activity of a selection of compounds using a fluorescence-based assay; **SJW** led transcriptomics and co-wrote aspects of the manuscript; **JB** led on evaluation of activity of azetidine derivatives against MDR-TB, determination of MIC values and co-wrote aspects of the manuscript; **CN** responsible for delivery of some synthetic aspects, suggested experiments, contributed to interpreting findings, supervised aspects of the research and co-wrote aspects of the manuscript; **JSF** led chemistry aspects, suggested critical experiments, interpreted key findings, supervised aspects of the research and co-wrote aspects of the manuscript; **LJA** led biology aspects, suggested critical experiments, interpreted key findings, supervised aspects of the research, co-wrote much of the manuscript.

#### Further general biological details

*C. glutamicum* (ATCC 13032) wild type and *C. glutamicum△pks13* were cultured at 30 °C, 180 rpm in brain heart infusion (BHI) media. ($MIC_{99}$) was determined in 96-well flat bottom, black polystyrene microtiter plates (Greiner) in a final volume of 200 μL. Compounds were two-fold serially diluted in neat DMSO and added to the microtiter plate at a final concentration of 1% DMSO. DMSO (1% in 7H9) was used as a positive control and rifampicin as a negative. The inoculum was standardised at $OD_{600}$ 0.05 in BHI medium and added to the plate, which was then incubated without shaking at 30 °C for 24 hours. Following incubation, 42 μL of resazurin (0.02% v/v in $dH_2O$) was added to each well and incubated for a further two hours. Fluorescence was measured (Polarstar Omega plate reader ex 544 nm, em 590 nm) and the data normalized. The concentration of drug required to inhibit cell growth by 99% was calculated by non-linear regression (Gomperz equation for MIC determination, GraphPad Prism). For lipid analysis of *C. glutamicum* (ATCC 13032) wild type and *C. glutamicum△pks13,* cells were harvested and extracted using chloroform:methanol:water (10:10:3, v/v/v, 2 mL) for two hours at 50 °C. Following centrifugation, the organic extracts were combined with chloroform and water (1.75 mL and 0.75 mL respectively). The lower organic phase containing associated lipids was recovered, washed twice with chloroform:methanol:water (3:47:48, v/v/v, 2 mL) and dried with $N_2$. Samples were resuspended in chloroform:methanol (2:1, v/v, 200 μL) and OD adjusted volumes were subjected to thin-layer chromatography (TLC) analysis. Cell wall associated lipids were visualised by either heating TLC plates after treatment with molybdophosphoric acid (MPA) in ethanol (5% w/v) or alpha-naphthol in ethanol (5% w/v).

#### General chemical synthesis aspects

##### *Chemistry at GIBH*

All commercially available solvents and reagents were purchased and used without further purification. All reactions were monitored by thin layer chromatography (TLC) with silica gel-coated plates and were visualised under UV light at 254 nm or by potassium permanganate solution staining followed by heating. $^{1}$H NMR spectrums were acquired *via* a Bruker AVIII300 or AVIII400 at 300 or 400 MHz respectively at room temperature (21 to 28 °C). $^{13}$C NMR spectrums were recorded *via* a Bruker AVIII400 or AVIII500 at 101 MHz or 126 MHz respectively at room temperature. $^{19}$F NMR spectrums were recorded *via* a Bruker AVIII500 at 471 MHz at room temperature. Chemical shifts (δ) are reported in parts per million relative to residual solvent for $^{1}$H and $^{13}$C NMR spectroscopy. The NMR spectral data collected thus were processed using the MestReNova-12.0.3 software package. Coupling constants (*J*) are reported in Hertz (Hz). Multiplicities of the signals are abbreviated as singlet (s), doublet (d), triplet (t), quartet (q), septet (sept), multiplet (m) and broad (br). Mass spectra were obtained on an API 2000 electrospray mass spectrometer. Infrared spectra were recorded at room temperature using a Bruker Tensor 27 FT-IR spectrometer using KBr pellets. Column chromatography purification was done using Silica Gel 200-300. Thin layer chromatography (TLC) was performed using aluminium-backed, F254-coated analytical TLC plates which were visualised under UV light at 254 nm or by staining phosphomolybdic acid (in ethanol) followed by heating.

***Chemistry at UoB***

All commercially available solvents and reagents were purchased and used without further purification. **NMR spectroscopy:** $^{1}H$ NMR spectrums were acquired *via* a Bruker AVIII300 or AVIII400 at 300 or 400 MHz respectively at room temperature (21 to 28 °C). $^{13}C$ NMR spectrums were recorded *via* a Bruker AVIII400 or AVIII500 at 101 MHz or 126 MHz respectively at room temperature, the JMOD pulse sequence was used in some cases to assist assignment with $CH_3$ and CH designated and reported as (+) and $C_q$ and $CH_2$ designated and reported as (-). $^{19}F$ NMR spectrums were recorded *via* a Bruker AVIII400 at 377 MHz at room temperature. Chemical shifts (δ) are reported in parts per million relative to residual solvent chloroform (7.26 ppm in $CDCl_3$) or tetramethylsilane (TMS, 0.00 ppm) internal standards for $^{1}H$, relative to residual solvent for $^{13}C$ NMR spectroscopy and are indirectly referenced to $CFCl_3$ at 0.00ppm for $^{19}F$ NMR spectroscopy. Coupling constants (*J*) are reported in Hertz (Hz). Multiplicities of the signals were abbreviated as singlet (s), doublet (d), triplet (t), quartet (q), septet (sept), multiplet (m) and broad (br). The NMR spectral data collected thus were processed using the MestReNova-12.0.3 software package. Mass spectra were obtained on a Waters LCT Time-of-Flight (TOF) Mass Spectrometer or a Waters GCT Premier Time-of-Flight Mass Spectrometer (TOF MS), using the ESI+ technique. Infrared spectra were recorded at room temperature nest with the ATR technique using a PerkinElmer 100FT-IR spectrometer. Flash column chromatography was performed using Teledyne ISCO CombiFlash Rf 200i, mobile and stationary phases are described in the general methods or the experimental procedures. Thin layer chromatography (TLC) was performed using aluminium-backed, F254-coated analytical TLC plates which were visualised under UV light at 254 nm or by potassium permanganate solution staining followed by heating.

***General synthetic chemistry procedures***

General procedure A: Imine **S3aa–da** synthesis; Supplementary Scheme 1, step i

To solutions of aldehydes (**S1a–d**, 1.02 equiv.) in methanol, amines (**S2a** or **S2b**, 1.00 equiv., 0.20–0.25 M M) were added and the mixture was heated at reflux for three hours. The mixture was allowed to cool to room temperature and was concentrated *in vacuo* to the corresponding afford imines (**S3aa**, **bb**, **ca** or **da**). The loss of aldehydic (*H*C=O) proton and emergence of a signal consistent with imine (*H*C=N) in respective proton NMR spectrums confirmed formation of product and the materials were used in the next step without further purification or analysis.

General procedure B: Homoallylamine derivatives **S4aa–da** synthesis; Supplementary Scheme 1, step ii

Allyl bromide (2.5 equiv.) was added dropwise into a stirred suspension of freshly activated zinc powder (3.0 equiv.) in anhydrous tetrahydrofuran (0.20–0.25 M) at 0 ºC. After 30 minutes, imine (**S3aa**, **bb**, **ca** or **da**, 1 equiv.) was added to the suspension at room temperature. The resulting mixtures were stirred at room temperature (21–28 °C) for 14 hours, or until reactions were judged complete by TLC (silica, hexane/ethyl acetate 10:1). At which time sodium bicarbonate (saturated aqueous) was added and the resulting mixtures filtered through celite. The filtrates were extracted with ethyl acetate (3 × 30 mL), washed with water (2 × 20 mL), dried over anhydrous magnesium sulfate and concentrated *in vacuo*. The residues thus obtained, were purified by flash chromatography (silica, ethyl acetate/hexane gradient elution 0:100–10:90), to afford the corresponding homoallyl amine (**S4aa**, **S4bb**, **S4ca** or **S4da**).

General procedure C: *cis*-Iodo-azetidine derivatives **S5aa–da** synthesis; Supplementary Scheme 1, step iii

To solutions of homoallyl amines (**S4aa**, **S4bb**, **S4ca** or **S4da**, 0.20–0.25 M, acetonitrile, 30 mL), iodine (3 equiv.) and sodium bicarbonate (5 equiv.) were added. The mixtures were stirred at 16 °C in order to suppress formation of pyrrolidine by-products.[1,2] After the reaction was judged to be complete (by TLC analysis silica, hexane/ethyl acetate 10:1), sodium thiosulfate solution (saturated aqueous) was added in order to facilitate removal of excess iodine. This mixture was extracted with ethyl acetate (3 × 30 mL), washed with water (2 × 20 mL), dried over anhydrous magnesium sulfate and concentrated *in vacuo*. The presence of iodo-azetidine derivatives (**S5aa**, **S5bb**, **S5ca** or **S5da**) were confirmed by proton NMR spectroscopy and used in the next step without further purification.

General procedure 4: Synthesis of *cis*-amino-azetidine derivatives **BGAz-001–005** Supplementary Scheme 1, step iv

Iodo-azetidine derivatives (**S5aa**, **S5bb**, **S5ca** or **S5da**) were dissolved in excess amine (pyrrolidine or propargyl amine, neat or 2.0 M methylamine in THF or DMSO) and stirred at room temperature for 48 h. Excess tetrahydrofuran or amine was removed *in vacuo* and the residues thus obtained were purified by flash column chromatography (ethyl acetate/hexane gradient elution 0:100–40:60), and if required, semi-preparative HPLC (methanol (10–15%)/ammonium hydroxide solution (0.1%))to afford *cis*-amino azetidine derivatives **BGAz-001–005**.

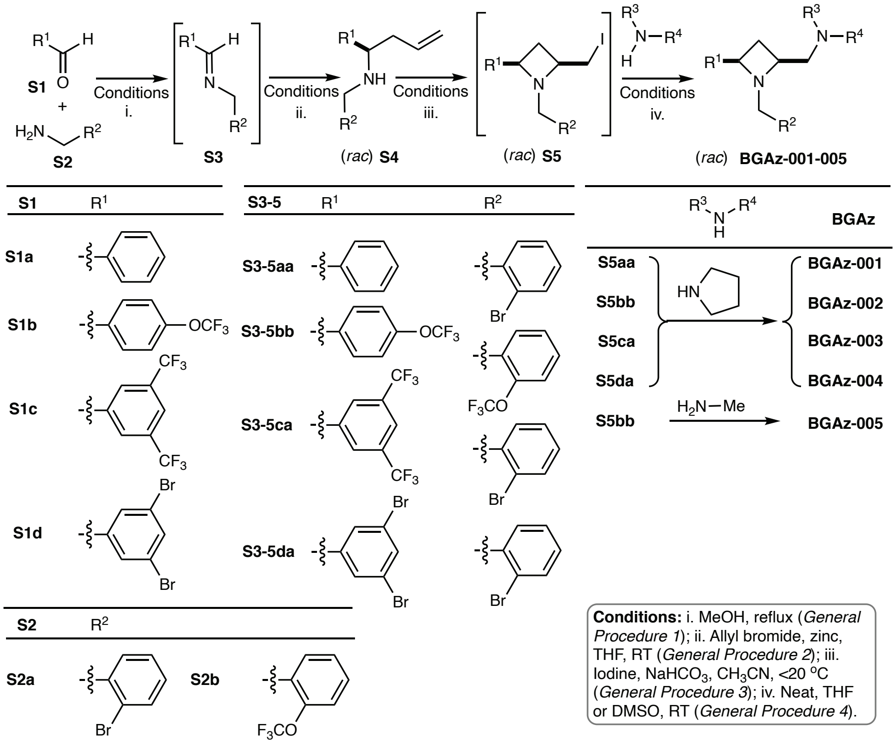

**Supplementary Scheme 1.** Synthesis of **BGAz001–005**.

<u>Synthesis of (*E*)-*N*-(2-bromobenzyl)-1-phenylmethanimine, **S3aa**</u>

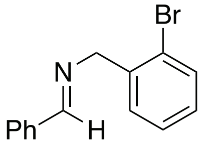

**S3aa**

General procedure A (4.7 mmol scale), benzaldehyde and (2-bromophenyl)methanamine as aldehyde and amine respectively, was employed (**S3aa** was isolated as a pale-yellow oil, quantitative conversion). Imine **S3aa** was used without further purification in the next step. Compound **S3aa** has been previously reported in the literature.[3-5]

<u>Synthesis of (*E*)-*N*-(2-(trifluoromethoxy)benzyl)-1-(4-(trifluoromethoxy)phenyl)methanimine, **S3bb**</u>

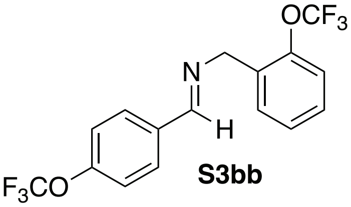

**S3bb**

General procedure A (5.3 mmol scale), 4-(trifluoromethoxy)benzaldehyde and (2-(trifluoromethoxy)phenyl)methanamine as aldehyde and amine respectively, was employed (**S3bb** was isolated as a yellow oil, quantitative conversion). Imine **S3bb** was used without further purification in the next step.

<u>Synthesis of (*E*)-1-(3,5-bis(trifluoromethyl)phenyl)-*N*-(2-bromobenzyl)methanimine, **S3ca**</u>

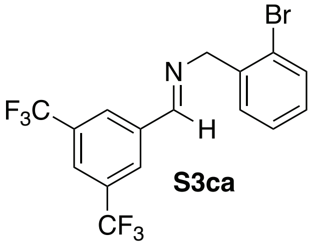

**S3ca**

General procedure A (4.1 mmol scale), 3,5-bis(trifluoromethyl)benzaldehyde and (2-bromophenyl)methanamine as aldehyde and amine respectively, was employed (**S3ca** was isolated as a yellow oil, quantitative conversion). Imine **S3ca** was used without further purification in the next step.

Synthesis of (*E*)-*N*-(2-bromobenzyl)-1-(3,5-dibromophenyl)methanimine, **S3da**

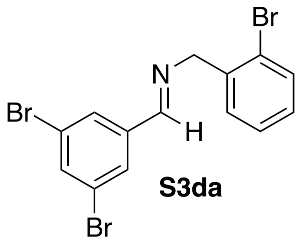

**S3da**

General procedure A (3.8 mmol scale) 3,5-dibromobenzaldehyde and (2-bromophenyl)methanamine as aldehyde and amine respectively, was employed (**S3da** was isolated as a brown solid, quantitative conversion). Imine **S3da** was used without further purification in the next step.

Synthesis of *N*-(2-bromobenzyl)-1-phenylbut-3-en-1-amine, **S4aa**

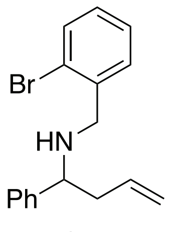

**S4aa** (*rac*)

General procedure B was used employing **S3aa** as the starting imine (1.8 mmol scale, **S4aa** was isolated as a pale-yellow oil, 88% yield). $\boldsymbol{\delta}_{\mathbf{H}}$ (400 MHz, $CDCl_3$) 7.57–7.49 (1 H, m), 7.41–7.32 (4 H, m), 7.30–7.23 (2 H, m), 7.17–7.06 (1 H, m), 5.70 (1 H, dddd, *J* 17.1, 10.1, 8.1 & 5.9), 5.18–4.95 (2 H, m), 3.74 (1 H, d, *J* 13.7), 3.67–3. 64 (1H, m), 3.63 (1 H, d, *J* 13.7), 2.49–2.33 (2 H, m), 1.99 (1 H, s); $\boldsymbol{\delta}_{\mathbf{C}}$ (126 MHz, $CDCl_3$) 143.2, 139.0, 135.2, 132.8, 130.8, 128.6, 128.4, 127.4, 127.3, 127.2, 124.1, 117.7, 61.4, 51.4, 43.0; **IR** $\nu$ ($cm^{-1}$): 3393, 3083 & 1453; **MS** (ES+) found 315.9 [M+H ($^{79}$Br)]$^+$ & 317.9 [M+H ($^{81}$Br)]$^+$.

Synthesis of *N*-(2-(trifluoromethoxy)benzyl)-1-(4-(trifluoromethoxy)phenyl)but-3-en-1-amine, **S4bb**

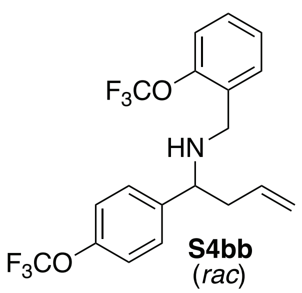

**S4bb** (*rac*)

General procedure B was used employing **S3bb** as the starting imine (1.4 mmol scale, **S4bb** was isolated as a pale-yellow oil, 70% yield). $\boldsymbol{\delta}_{\mathbf{H}}$ (400 MHz, $CDCl_3$) 7.48–7.12 (8 H, m), 5.67 (1 H, dddd, *J* 14.1, 10.6, 8.3, 5.9), 5.14–5.01 (2 H, m), 3.69 - 3.66 (1 H, m) 3.68 (1 H, d, *J* 13.8), 3.61 (1 H, d, *J* 13.8), 2.51–2.25 (2 H, m), 1.8 (1H, s); $\boldsymbol{\delta}_{\mathbf{C}}$ (126 MHz, $CDCl_3$) 148.3 ($^1J_{CF}$ 1.8), 147.7 ($^1J_{CF}$ 1.5), 142.3, 134.7, 132.6, 130.7, 128.5, 128.4, 126.7, 120.9, 120.6 ($^3J_{CF}$ 257.5), 120.6 ($^3J_{CF}$ 257.5) 120.3, 120.3, 118.0, 61.0, 45.9, 43.2; $\boldsymbol{\delta}_{\mathbf{F}}$ (471 MHz, $CDCl_3$) -57.11, -57.87; **IR** $\nu$ ($cm^{-1}$): 3389, 2914 & 1258; **MS** (ES+) 406.5 [M+H]$^+$.

Synthesis of 1-(3,5-bis(trifluoromethyl)phenyl)-*N*-(2-bromobenzyl)but-3-en-1-amine, **S4ca**

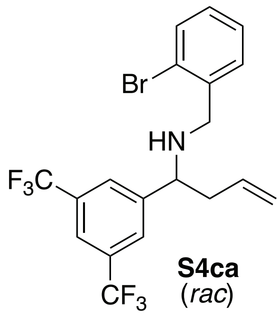

**S4ca** (*rac*)

General procedure B was used employing **S3ca** as the starting imine (1.2 mmol scale, **S4ca** was isolated as a yellow oil, 91% yield). $\boldsymbol{\delta}_{\mathbf{H}}$ (400 MHz, $CDCl_3$) 7.87 (2 H, s), 7.78 (1 H, s), 7.53 (1 H, dd, *J* 7.8, 1.3), 7.25 (1 H, td, *J* 7.4, 1.3), 7.20–7.09 (2 H, m), 5.67 (1 H, dddd, *J* 16.9, 10.4, 8.3, 5.8), 5.21–5.04 (2 H, m), 3.79 (1 H, dd, *J* 8.5, 5.2), 3.71 (1 H, d, *J* 13.5), 3.62 (1 H, d, *J* 13.5), 2.54–2.23 (2 H, m), 2.15 (1 H, s); $\boldsymbol{\delta}_{\mathbf{F}}$ (471 MHz, $CDCl_3$) -62.73; $\boldsymbol{\delta}_{\mathbf{C}}$ (126 MHz, $CDCl_3$) 146.5, 138.4, 133.8, 133.0, 131.6 (q, $^2J_{CF}$ 33.2), 130.7, 129.0, 127.7 (q, $^3J_{CF}$ 2.4), 127.5, 124.1, 123.4 (q, $^1J_{CF}$ 272.5), 121.3 (sept, $^3J_{CF}$ 3.9), 119.0, 60.9, 51.7, 43.1; **IR** $\nu$ ($cm^{-1}$): 3340, 3077, 1279, 1178; **MS** (ES+) 452.4 [M+H ($^{79}$Br)]$^+$ & 454.1 [M+H ($^{81}$Br)]$^+$.

Synthesis of *N*-(2-bromobenzyl)-1-(3,5-dibromophenyl)but-3-en-1-amine, **S4da**

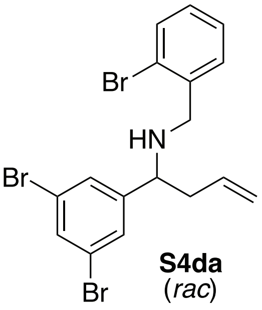
**S4da**
(*rac*)

General procedure B was used employing **S3da** as the starting imine (1.2 mmol scale, **S4da** was isolated as a yellow oil, 93% yield). **δ$_H$** (400 MHz, $CDCl_3$) 7.56 (1 H, t, *J* 1.8), 7.54 (1 H, dd & *J* 7.8, 1.3), 7.48 (2 H, d, *J* 1.8), 7.27 (1 H, td, *J* 7.3 & 1.3), 7.21 (1 H, dd, *J* 7.6 & 2.0), 7.13 (1 H, td, *J* 7.3 & 2.0), 5.65 (1 H, dddd, *J* 16.5, 11.0, 8.3 & 5.8), 5.34–4.69 (2 H, m), 3.72 (1 H, d, *J* 13.6), 3.61 (1 H, d, *J* 13.6), 3.58 (1 H, dd, *J* 8.3 & 5.3) 2.48–2.19 (2 H, m), 2.02 (1 H, s); **δ$_C$** (126 MHz, $CDCl_3$) 148.1, 138.8, 134.2, 132.9, 132.8, 130.6, 129.2, 128.8, 127.4, 124.0, 123.0, 118.5, 60.6, 51.6, 43.0.; **IR** *ν* ($cm^{-1}$): 3385, 2927, 1558, 1882 & 789; **MS** (ES+) 472.1 [M+H$^+$ (3 × $^{79}$Br)]$^+$, 474.3 [M+H (2 × $^{79}$Br, $^{81}$Br)]$^+$, 475.9 [M+H ($^{79}$Br, 2 × $^{81}$Br)]$^+$, 476.9 [M+H ($^{79}$Br, 2 × $^{81}$Br)]$^+$ & 477.9 [M+H (3 × $^{81}$Br)]$^+$.

Synthesis of (*rac*)-1-(((2,4-*cis*)-1-(2-bromobenzyl)-4-phenylazetidin-2-yl)methyl)pyrrolidine, **BGAz-001**

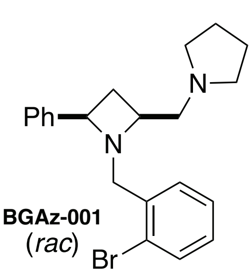
**BGAz-001**
(*rac*)

General procedure C was used employing **S4aa** as the starting homoallyl amine (0.6 mmol scale, **BGAz-001** was isolated as a brown oil, 40% yield). **δ$_H$** (400 MHz, $CDCl_3$): δ 7.48 (1H, dd, *J* 7.9 & 1.5, Ar*H*), 7.45–7.39 (3H, m, Ar*H*), 7.28 (2H, m, Ar*H*), 7.23–7.17 (1H, m, Ar*H*), 7.14 (1H, td, *J* 7.5 & 1.5, Ar*H*), 7.03 (1H, td, *J* 7.5 & 1.9, Ar*H*), 4.10 (1H, t, *J* 8.2, ArC*H*), 3.89 (1H, d, *J* 13.5, ArC*H*HN), 3.83 (1H, d, *J* 13.5, ArCH*H*N), 3.41 (1H, tdd, *J* 8.6, 6.8 & 3.6, $CH_2$C*H*$CH_2$), 2.70–2.40 (7H, m), 1.84 (1H, dt, *J* 10.3 & 8.5, ArCHC*H*H)1.77–1.68 (4H, m); **δ$_C$** (126 MHz, $CDCl_3$): δ 143.5, 137.9, 132.4, 131.6, 128.3, 128.0, 126.9, 126.8, 126.7, 124.3, 66.4, 62.2, 62.1, 60.7, 54.8, 35.0, 23.4; **IR** *ν* ($cm^{-1}$): 2783, 2787, 747 & 697; TOF MS (ES+) found 385.2 [M+H ($^{79}$Br)]$^+$ & 387.2 [M+H$^+$ ($^{81}$Br)]$^+$; **HRMS** (ES+) calcd for $C_{21}H_{26}N_2{}^{79}Br^+$: 385.1274, found 385.1277.

Synthesis of (*rac*)-1-(((2,4-*cis*)-1-(2-(trifluoromethoxy)benzyl)-4-(4-(trifluoromethoxy)phenyl)azetidin-2-yl)methyl)pyrrolidine, **BGAz-002**

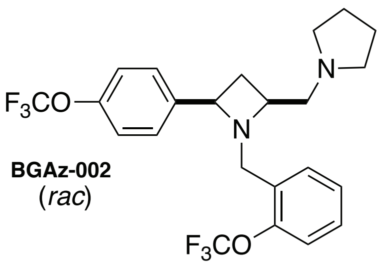
**BGAz-002**
(*rac*)

General procedure C was used employing **S4bb** as the starting homoallyl amine (0.5 mmol scale, **BGAz-002** was isolated as a brown oil, 23% yield). **δ$_H$** (400 MHz, $CDCl_3$) δ 7.37–7.32 (3H, m, Ar*H*), 7.19–7.10 (2H, m, Ar*H*), 7.09–7.03 (3H, m, Ar*H*), 3.98 (1H, t, *J* 8.1, Ar*H*), 3.78 (1H, d, *J* 13.3, ArC*H*HN), 3.72 (1H, d, *J* 13.3, ArCH*H*N), 3.31 (1H, tdd, *J* 8.5, 6.9 & 3.8, $CH_2$C*H*$CH_2$), 2.62–2.51 (2H, m), 2.47–2.38 (5H, m), 1.99–1.40 (5H, m); **δ$_C$** (101 MHz, $CDCl_3$) δ 148.1 (q, $^3J_{CF}$ 1.8), 147.6 (q, $^3J_{CF}$ 1.5), 142.2, 131.9, 130.8, 128.3, 128.0, 126.1, 120.6 (q, $^1J_{CF}$ 257.1), 120.5 (q, $^1J_{CF}$ 256.5), 120.5, 120.1, 65.4, 62.1, 62.1, 54.8, 54.7, 34.9, 23.4; **δ$_F$** (377 MHz, $CDCl_3$) δ -56.99 & -57.96; **IR** *ν* ($cm^{-1}$):

2960, 2790, 1250, 1213 & 1151; **TOF MS** (ES+) found 475.2 $[M+H]^+$ (100%) & 476.2; **HRMS** (ES+) calcd for $C_{23}H_{25}N_2O_2F_6^+$: 475.1815, found 475.1825.

Synthesis of (*rac*)-1-(((2,4-*cis*)-4-(3,5-bis(trifluoromethyl)phenyl)-1-(2-bromobenzyl)azetidin-2-yl)methyl)-pyrrolidine, **BGAz-003**

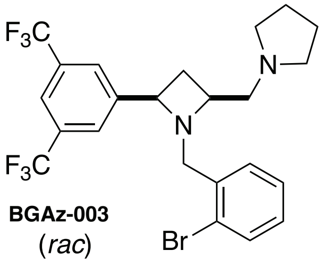

**BGAz-003**
(*rac*)

General procedure C was used employing **S4ca** as the starting homoallyl amine (0.4 mmol scale, **BGAz-003** was isolated as a brown oil, 50% yield). $\delta_H$ (400 MHz, $CDCl_3$) δ 7.73 (2H, s, Ar*H*), 7.59 (1H, s, Ar*H*), 7.39 (1H, dd, *J* 7.9 & 1.3, Ar*H*), 7.22 (1H, dd, *J* 7.5 & 1.7, Ar*H*), 7.04 (1H, td, *J* 7.5 & 1.3, Ar*H*), 6.95 (1H, td, *J* 7.9 & 1.7, Ar*H*), 4.17 (1H, t, *J* 8.1, ArC*H*), 3.95 (1H, d, *J* 12.4, ArC*H*HN), 3.71 (1H, d, *J* 12.4, ArCH*H*N), 3.49–3.42 (1H, m, $CH_2CHCH_2$), 2.79–2.60 (2H, m), 2.51-2.43 (5H, m), 1.96–1.62 (5H, m); $\delta_C$ (126 MHz, $CDCl_3$) δ 146.2, 136.7, 132.8, 131.8, 131.0 (q, $^2J_{CF}$ 32.8), 128.9, 126.9, 126.8 (q, $^3J_{CF}$ 3.6), 124.8, 123.5 (q, $^1J_{CF}$ 272.4), 120.6 (sept, $^3J_{CF}$ 3.8), 65.1, 62.0, 61.7, 60.6, 54.9, 35.0, 23.5; $\delta_F$ (377 MHz, $CDCl_3$) δ -62.81; **IR** *ν* ($cm^{-1}$): 2853, 2794, 1276, 1170 & 1130; **TOF MS** (ES+) found 521.1 $[M+H\ (^{79}Br)]^+$, 523.1 $[M+H\ (^{81}Br)]^+$ & 524.1; **HRMS** (ES+) calcd for $C_{23}H_{24}N_2F_6{}^{79}Br^+$: 521.1022, found 521.1026.

Synthesis of (*rac*)-1-(((2,4-*cis*)-1-(2-bromobenzyl)-4-(3,5-dibromophenyl)azetidin-2-yl)methyl)pyrrolidine, **BGAz-004**

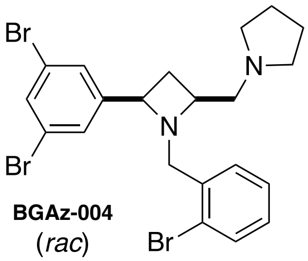

**BGAz-004**
(*rac*)

General procedure C was used employing **S4da** as the starting homoallyl amine (0.4 mmol scale, **BGAz-004** was isolated as a brown oil, 59% yield). $\delta_H$ (400 MHz, $CDCl_3$) δ 7.47 (1H, dd, *J* 7.8 & 1.3, Ar*H*), 7.42–7.38 (3H, m, Ar*H*), 7.28 (1H, dd, *J* 7.6 & 1.7, Ar*H*), 7.13 (1H, td, *J* 7.5 & 1.3, Ar*H*), 7.02 (1H, td, *J* 7.5 & 1.7, Ar*H*), 3.99 (1H, t, *J* 8.1, ArC*H*), 3.88 (1H, dd, *J* 12.9, ArC*H*HN), 3.71 (1H, dd, *J* 12.9, ArCH*H*N), 3.37 (1H, tdd, *J* 8.5, 7.0 & 3.5, $CH_2CHCH_2$), 2.68–2.55 (2H, m), 2.51–2.36 (5H, m), 1.80–1.66 (5H, m); $\delta_C$ (101 MHz, $CDCl_3$) δ 147.8, 137.1, 132.7, 132.2, 131.8, 128.8, 128.5, 126.9, 124.8, 122.5, 64.9, 62.1, 61.9, 60.5, 54.9, 35.0, 23.5; **IR** *ν* ($cm^{-1}$): 2956, 2790, 1584, 1556 & 740; **TOF MS** (ES+) found: 541.0 $[M+H^+\ (3 \times {}^{79}Br)]$, 543.0 $[M+H\ (2 \times {}^{79}Br, {}^{81}Br)]^+$, 545.0 $[M+H\ ({}^{79}Br, 2 \times {}^{81}Br)]^+$ & 547.0 $[M+H^+\ (3 \times {}^{81}Br)]^+$; **HRMS** (ES+) calcd for $C_{21}H_{24}N_2{}^{79}Br_3^+$: 540.9484, found 540.9489.

Synthesis of (*rac*)-*N*-methyl-1-((2,4-*cis*)-1-(2-(trifluoromethoxy)benzyl)-4-(4-(trifluoromethoxy)phenyl)-azetidine-2-yl)-methanamine, **BGAz-005**

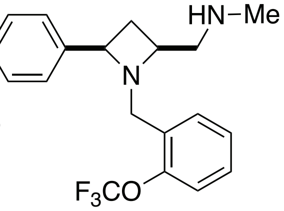

**BGAz-005**
(*rac*)

General procedure C was used employing **S4bb** as the starting homoallyl amine (0.5 mmol scale, **BGAz-005** was isolated as a brown oil, 25% yield). **$\delta_H$** ($CDCl_3$, 400 MHz) δ 7.43–7.31 (3H, m, Ar*H*), 7.24–7.04 (5H, m, Ar*H*), 3.99 (1H, t, *J* 8.1, ArC*H*), 3.75 (1H, d, *J* 13.2, ArC*H*HN), 3.71 (1H, d, *J* 13.2, ArCH*H*N), 3.34 (1H, dddd, *J* 8.7, 7.1, 5.3 & 3.7, $CH_2$C*H*$CH_2$), 2.54 (1H, dd, *J* 12.2 & 3.8, NHC*H*HCH), 2.48–2.36 (2H, m), 2.32 (3H, s, Me), 2.10 (1H, s), 1.95 (1H, dt, *J* 10.2 & 8.6, ArCHCH*H*); **$\delta_C$** (101 MHz, $CDCl_3$) δ 148.2 (q, $^3J_{CF}$ 1.8), 147.7 (q, $^3J_{CF}$ 1.5), 141.9, 131.7, 130.6, 128.6, 128.0, 126.3, 120.6 (q, $^1J_{CF}$ 257.3), 120.6, 120.5 (q, $^1J_{CF}$ 256.6), 120.1 65.0, 62.0, 55.6, 54.8, 36.6, 31.3; **$\delta_F$** (377 MHz, $CDCl_3$) δ -57.03 & -57.96; **TOF MS** (ES+) found 435.2 $[M+H]^+$; **HRMS** (ES+) calcd for $C_{20}H_{21}N_2F_6O_2^+$: 435.1502, found 435.1505; **IR** *ν* ($cm^{-1}$): 2851, 1249, 1212, 1150 & 758.

#### Microbiology

##### ***Extraction of mycobacterial genomic DNA and whole genome sequencing***

*M. smegmatis* and *M. bovis* BCG were plated onto agar containing 5, 10 and 20 × MIC concentrations of **BGAz-002–BGAz-005** in attempts to generate resistant mutants. Treatment of *M. bovis* BCG with 10 × $MIC_{99}$ of **BGAz-002** resulted in the formation of three resistant colonies. Whole-genome sequencing analysis of these three colonies detected six putative non-synonymous single nucleotide polymorphisms (SNPs) compared to the *M. bovis* BCG reference sequence (Supplementary Table 1). Of these six, three were found across all the mutants. However, these three SNPs (in genes *infB, lipN* and BCG_3519) align with SNPs previously reported in the laboratory parental strain of BCG,[6] and are background mutations which have arisen during laboratory storage, and thus cannot be conferring the resistant phenotype. The stochastic nature of the other three SNPs identified means that the genes in which they occur can be assumed not to be the target of the BGAz compounds and are unlikely to confer specific resistance. Indeed, two of these SNPs occur in BCG_0727, a transcriptional regulator of the MmpL5 efflux pump involved in non-specific resistance.[7] Thus, none of the SNPs identified in the three resistant colonies provide insights into the specific gene target of the BGAz compounds tested.

**Supplementary Table 1.** Non-synonymous miscoding single nucleotide polymorphisms of apparent **BGAz-004** resistant mutants of BCG.

| ***M. bovis*** **(BCG) chromosome** | **Codon Change** | **Amino Acid Change** | **Gene** | **Mutant 1** | **Mutant 2** | **Mutant 3** |
|---|---|---|---|---|---|---|
| 810466 | aTg/aGg | M23R | BCG_0727 | - | - | G |
| 810652 | gTc/gGc | V85G | BCG_0727 | - | G | - |
| 3100829 | gCc/gTc | A344V | infB | A | A | A |
| 3278815 | Ctg/Gtg | L351V | lipN | C | C | C |
| 3856853 | gGc/gAc | G294D | BCG_3519 | A | A | A |
| 3907860 | Aac/Gac | N599D | PE_PGRS53 | - | G | - |

#### ***Transcriptomic supporting data***

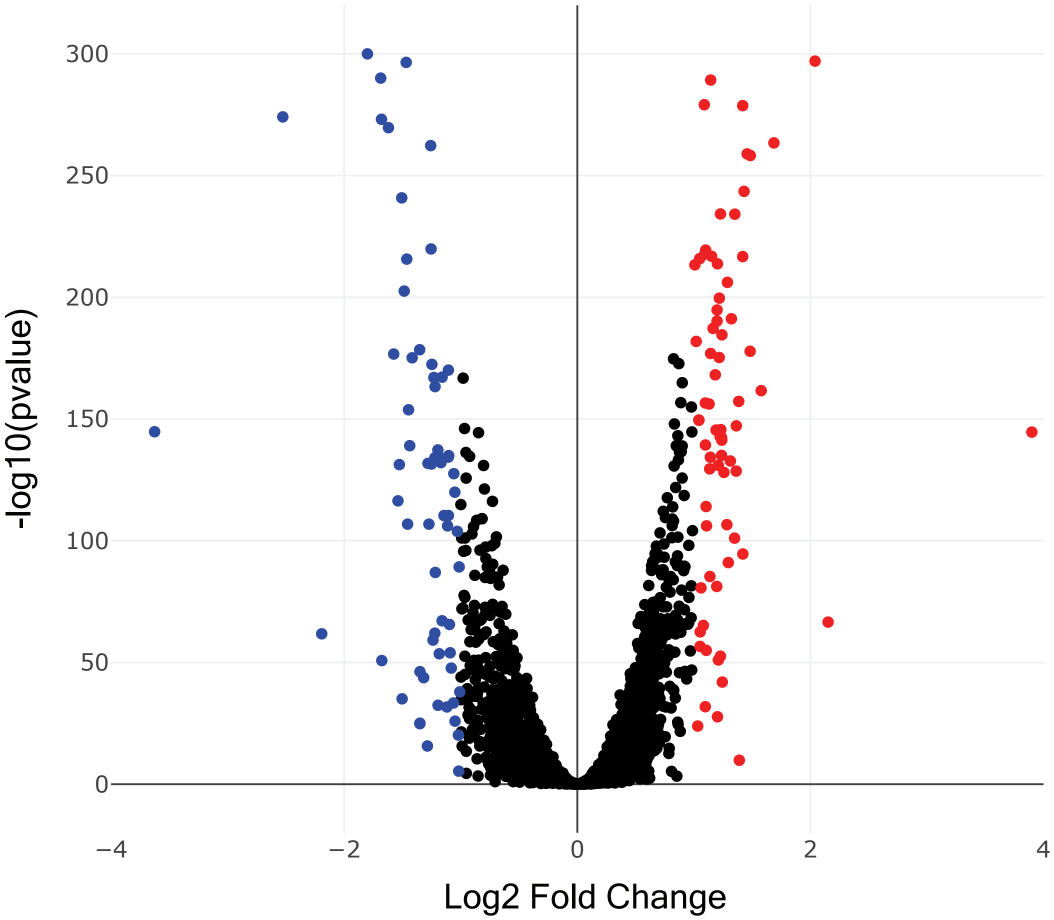

**Supplementary Figure 1.** Volcano plot of the *M. bovis* BCG transcriptional response to **BGAz-004** exposure, highlighting genes significantly differentially expressed relative to carrier control. Red colouring marking significantly induced genes, blue repressed genes.

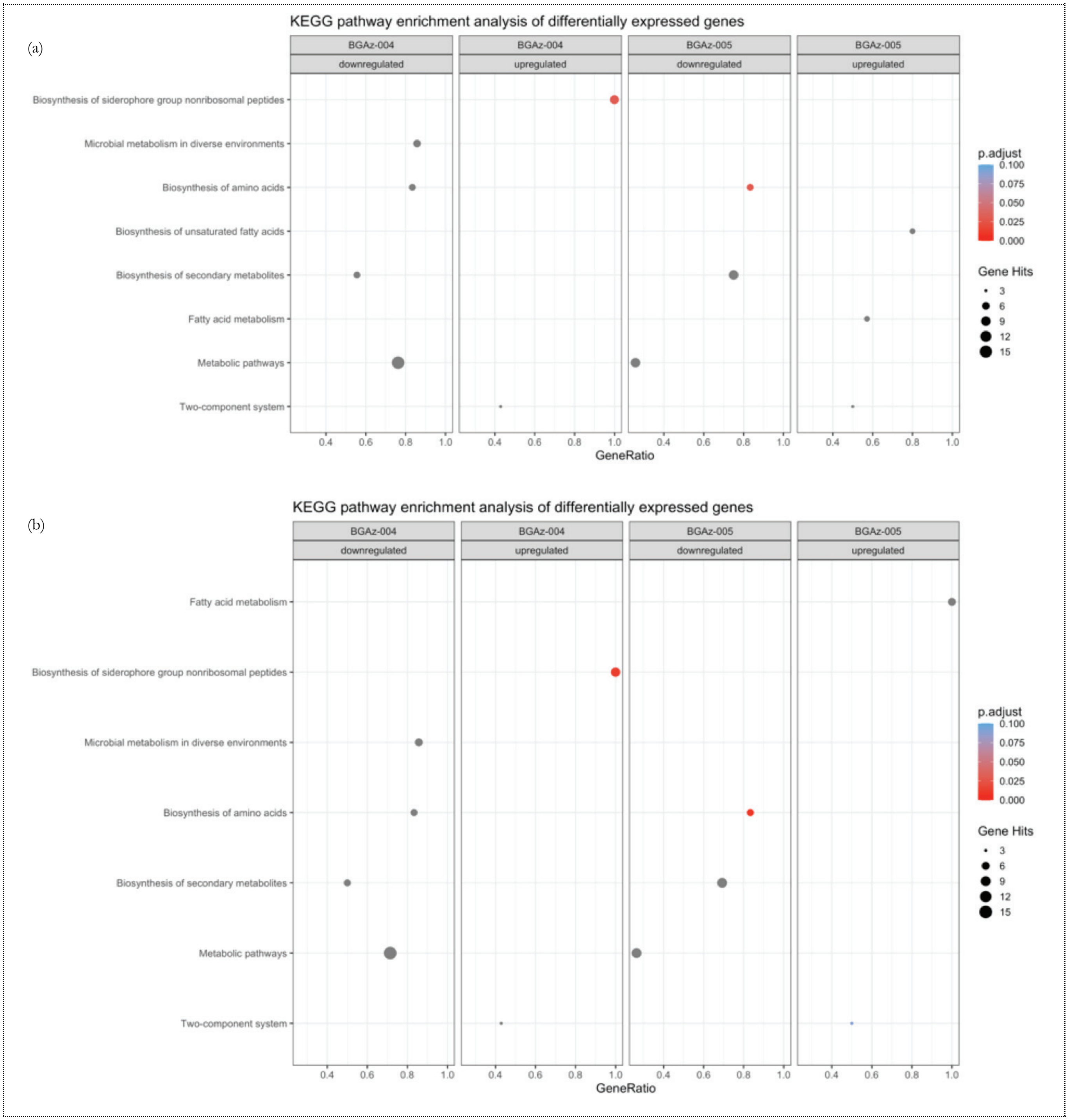

**Supplementary Figure 2.** KEGG Pathway Enrichment Analysis first selects genes with a differential expression of at least log2foldchange of 1. These genes are then clustered into functional groups (using information provided in the KEGG database), up to a P-value of 1. For this analysis, only genes with a 2-fold cutoff of 2 and an adjusted P-value of at least 0.05 were used. All other genes were disregarded. Analysis was completed with both (a) BCG identifiers using Pasteur genome and then (b) Rv identifiers using H37Rv genome.

***BGAz-004* significantly alters mycobacterial cell envelope composition**

To investigate the mechanism by which mycobacterial cell envelope lipid composition is affected by **BGAz-004**, actively growing cultures of BCG were exposed to increasing concentrations of compound followed by metabolic labelling using [$^{14}$C]-acetic acid. Autoradiographs of cell envelope lipids separated by thin layer chromatography (TLC) reveals that treatment of BCG with **BGAz-004** at 0.5 × MIC causes a significant reduction in trehalose monomycolate (TMM) and trehalose dimycolate (TDM) and a complete loss of TMM and TDM at concentrations beyond the MIC (Supplementary Figure 3A). The formation of cytoplasmic membrane phospholipids (PIMs and CL) remain unaffected (Supplementary Figure 3A). The analysis of lipids loaded and separated by TLCs that had been normalised for total lipids extracted, revealed an altered lipid profile highlighting the accumulation of an unidentified lipid species that resolves to a relatively high Rf (Lipid species X, (Supplementary Figure 5B). The analysis of mycolic acid methyl esters (MAMES) reveals that both alpha and keto mycolates bound to the cell wall arabinogalactan (AG) are gradually depleted as BCG is exposed to increasing concentrations of **BGAz-004** during active cell culture (Supplementary Figure 3C). Quantification of the relative abundance of each lipid species highlights the significant depletion of mycolates (either conjugated to trehalose in the form of TMM/TDM or AG) when **BGAz-004** is used at a half MIC, whilst other lipids including PI and PIMS remain largely unaffected (Supplementary Figure 3D).

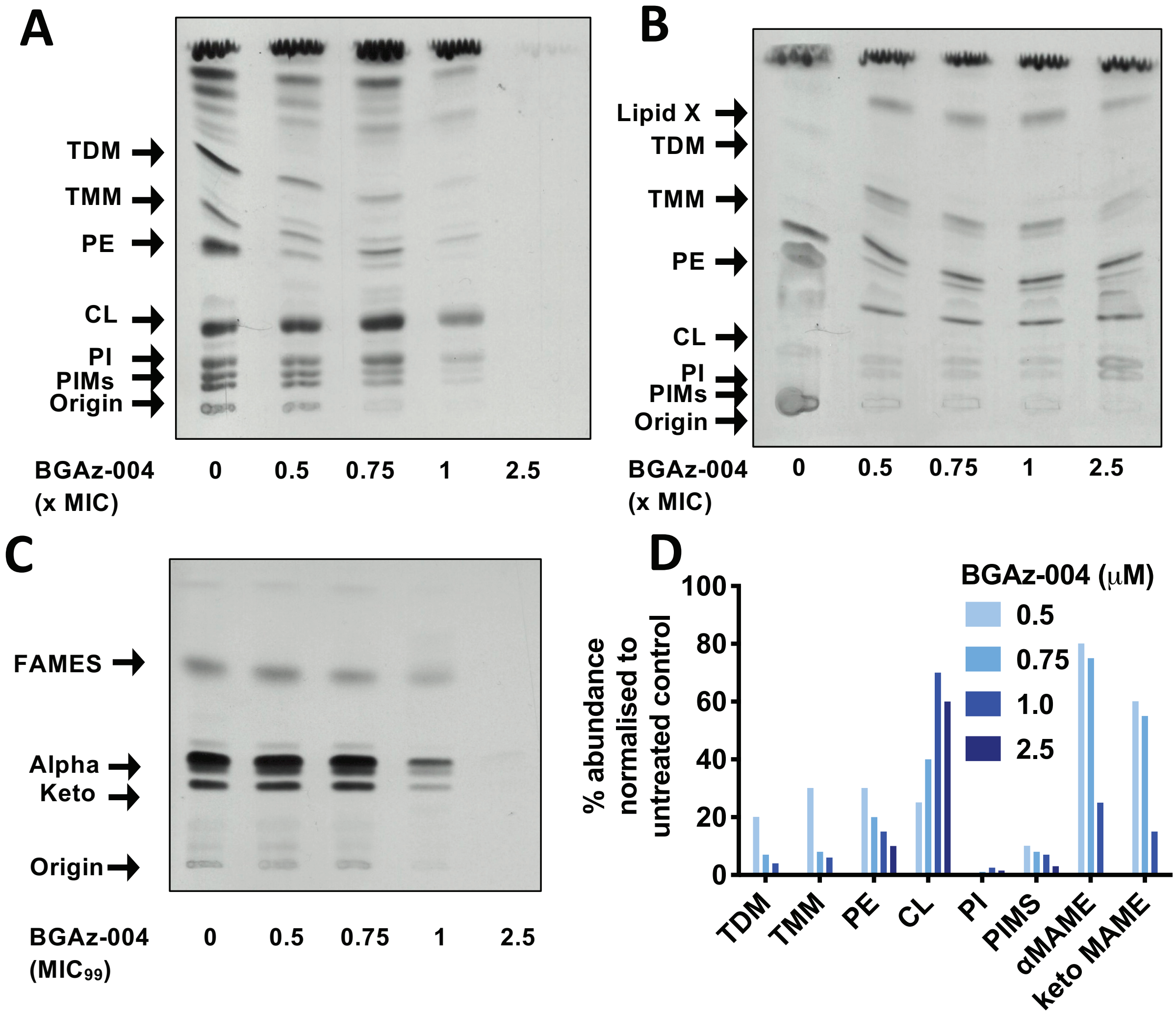

**Supplementary Figure 3.** BCG cell envelope lipid analysis upon exposure to **BGAz-004**. BCG were cultured in 7H9 broth and exposed to increasing concentrations of **BGAz-004**. Lipids were selectively labelled with [$^{14}$C]-acetic acid for 12 hours and cell envelope lipids were selectively removed by solvent extraction, separated by TLC (chloroform/methanol/water, 80:20:2, v/v/v), and visualised by autoradiography. A: equal volumes of lipids loaded adjusted for BCG growth. B: equal counts of lipids (25,000 cpm) loaded. C: Mycolic acid methyl ester (MAME) analysis of cell wall bound mycolates released by 5% TBAH and separated by TLC (petroleum ether/acetone, 95:5, v/v). D: quantification of BCG lipids from panels A-C by densitometry.

To investigate the effect of **BGAz-004** on mycobacterial cell envelope composition and to further identify the composition of lipid-X that appears in mycobacteria, actively growing cultures of *M. smegmatis* were exposed to **BGAz-004** concentrations, which resulted in a titratable-dependent reduction in the formation of TMM and TDM which can be observed by staining with MPA and α-napthol (Supplementary Figure 4A) consistent with [$^{14}$C]-labelling experiments performed when **BGAz-005** was exposed to BCG (Main Article). *M. smegmatis* exposed to the highest concentration of **BGAz-004**, resulted in a significant increase in the relative abundance of free mycolic acid (MA) within the cell envelope (Supplementary Figure 4C and D). This effect can also be observed when *M. smegmatis* is exposed to the known FbpC inhibitor ebselen (Supplementary Figure 5).

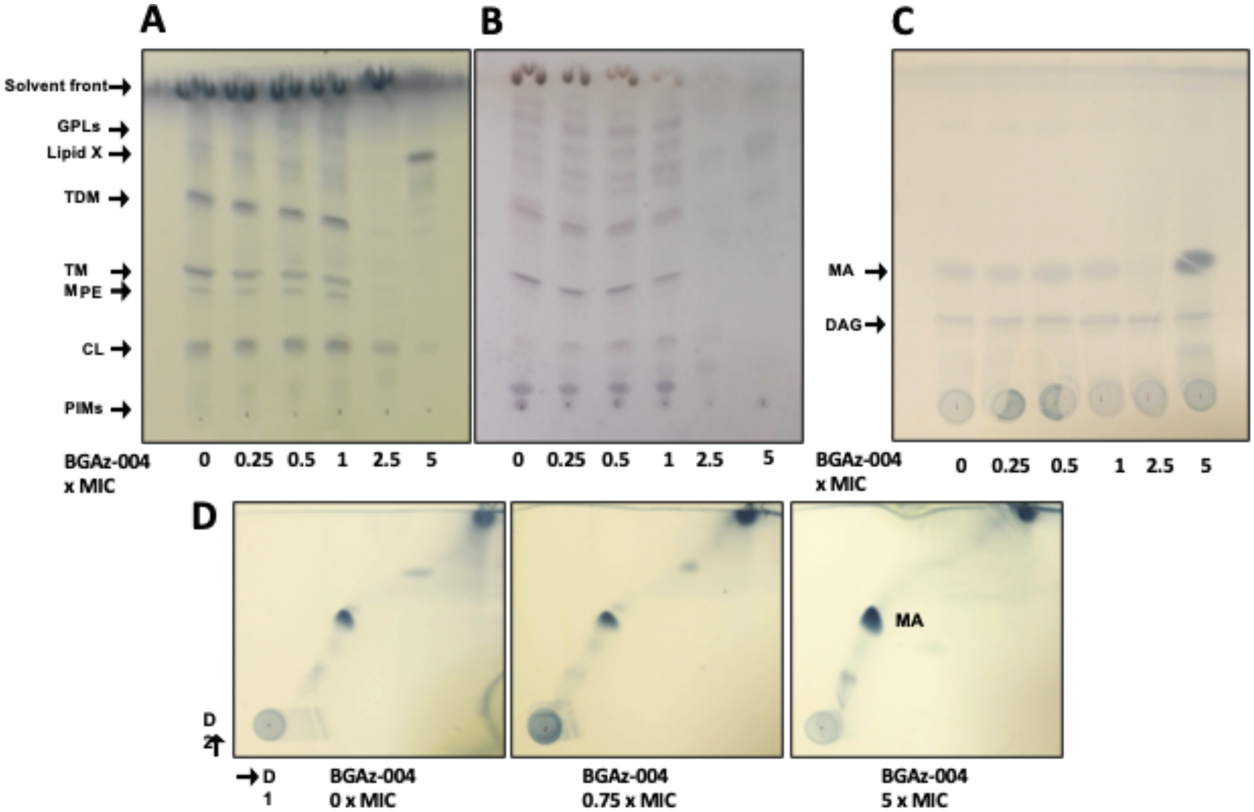

**Supplementary Figure 4.** *M. smegmatis* cell envelope lipid analysis upon exposure to **BGAz-004**. *M. smegmatis* were cultured in 7H9 broth, exposed to increasing concentrations of **BGAz-004** for 6 h and the cell envelope lipids selectively removed by solvent extraction. Equal volumes of lipid adjusted by bacterial growth were separated by TLC (chloroform/methanol/water, 80:20:2, v/v/v), and stained with MPA (A) or alpha-naphthol (B). Equal volumes of lipid adjusted by bacterial growth were separated by TLC (hexane/diethyl ether/acetic acid), 70:30:1, v/v/v and stained with MPA (C). Equal volumes of lipid adjusted by bacterial growth were separated by 2D-TLC (direction 1 chloroform/methanol 96:4, v/v, direction 2 toluene/acetone 80:20, v/v) and stained with MPA (D).

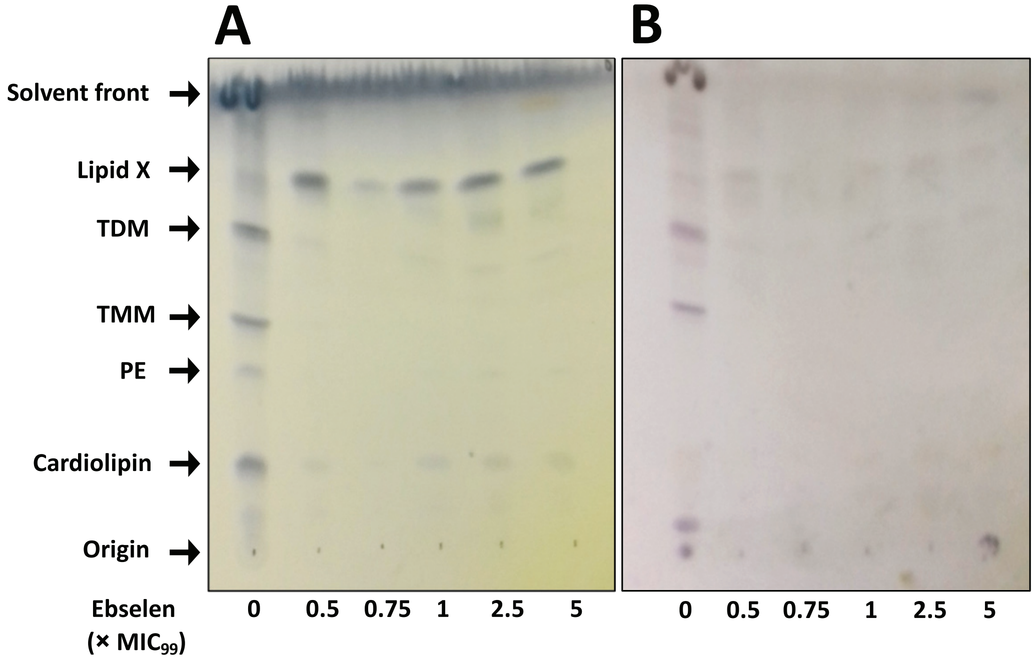

**Supplementary Figure 5.** M. smegmatis cell envelope lipid analysis upon exposure to Ebselen**.** *M. smegmatis* were cultured in 7H9 broth, exposed to increasing concentrations of Ebselen for 6 h and the cell envelope lipids selectively removed by solvent extraction. Equal volumes of lipid adjusted by bacterial growth were separated by TLC (chloroform/methanol/water, 80:20:2, v/v/v), and stained with MPA (A) or alpha-naphthol (B).

#### *BGAz-005* ***targets late-stage mycolic acid biosynthesis in*** **C. glutamicum**

**BGAz-005** inhibits corynemycolic acid biosynthesis in *C. glutamicum* similarly to mycobacteria (Supplementary Figure 6).

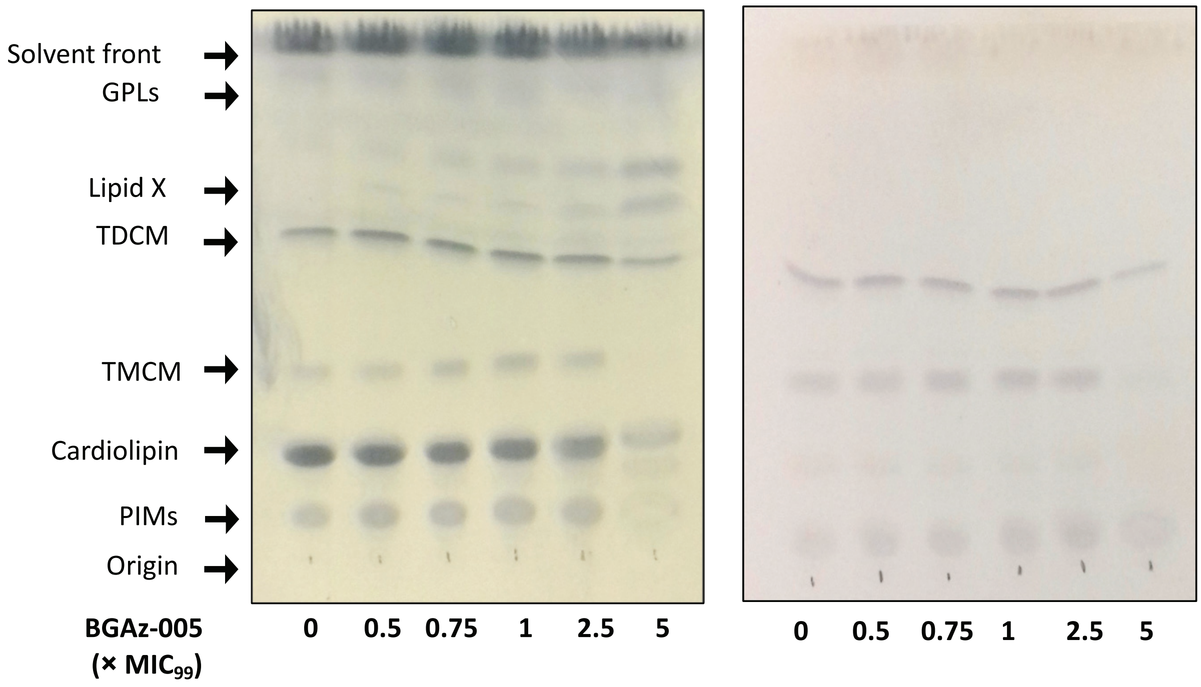

**Supplementary Figure 6.** C. glutamicum cell envelope lipid analysis upon exposure to **BGAz-005.** *M. smegmatis* were cultured in BHI broth, exposed to increasing concentrations of **BGAz-005** for 6 h and the cell envelope lipids selectively removed by solvent extraction. Equal volumes of lipid adjusted by bacterial growth were separated by TLC (chloroform/methanol/water, 60:16:2, v/v/v), and stained with MPA (A) or alpha-naphthol (B).

**Supplementary Table 2.** MIC values of the **BGAz-002–BGAz-005** and ebselen against mycobacterial and corynebacterial strains. NT = not tested.

| Compound | *C. glutamicum* | *C. glutamicum*△*pks13* | BCG pVV16-*mmpL3* | BCG pTIC6-*fbpA* | BCG pTIC6-*fbpB* | BCG pTIC6-*fbpC* |
|---|---|---|---|---|---|---|
| 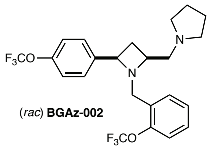 (*rac*) **BGAz-002** | 50 | 50 | 25 | NT | NT | NT |
| 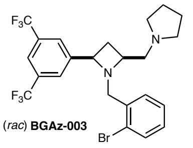 (*rac*) **BGAz-003** | 14 | 10 | 40 | NT | NT | NT |
| 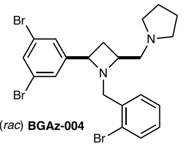 (*rac*) **BGAz-004** | 20 | 10 | 40 | 20 | 28 | 34 |
| 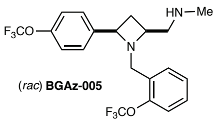 (*rac*) **BGAz-005** | 12 | 10 | NT | 25 | 50 | 50 |
| 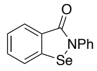 **Ebselen** | NT | NT | NT | 28 | 90 | 98 |

[(a)] MIC as determined by a modified Gompertz function

**S4aa** [1]H NMR Spectrum

S4aa

NAME Feb14-2017
EXPNO 25
PROCNO 1
Date_ 20170214
Time 17.22
INSTRUM spect
PROBHD 5 mm DUL 13C-1
PULPROG zg30
TD 65536
SOLVENT CDCl3
NS 16
DS 2
SWH 8278.146 Hz
FIDRES 0.126314 Hz
AQ 3.9584243 sec
RG 128
DW 60.400 usec
DE 6.50 usec
TE 295.7 K
D1 1.00000000 sec
TD0 1

======== CHANNEL f1 ========
NUC1 1H
P1 12.58 usec
PL1 0.00 dB
PL1W 10.87646866 W
SFO1 400.1324710 MHz
SI 32768
SF 400.1300093 MHz
WDW EM
SSB 0
LB 0.30 Hz
GB 0
PC 1.00

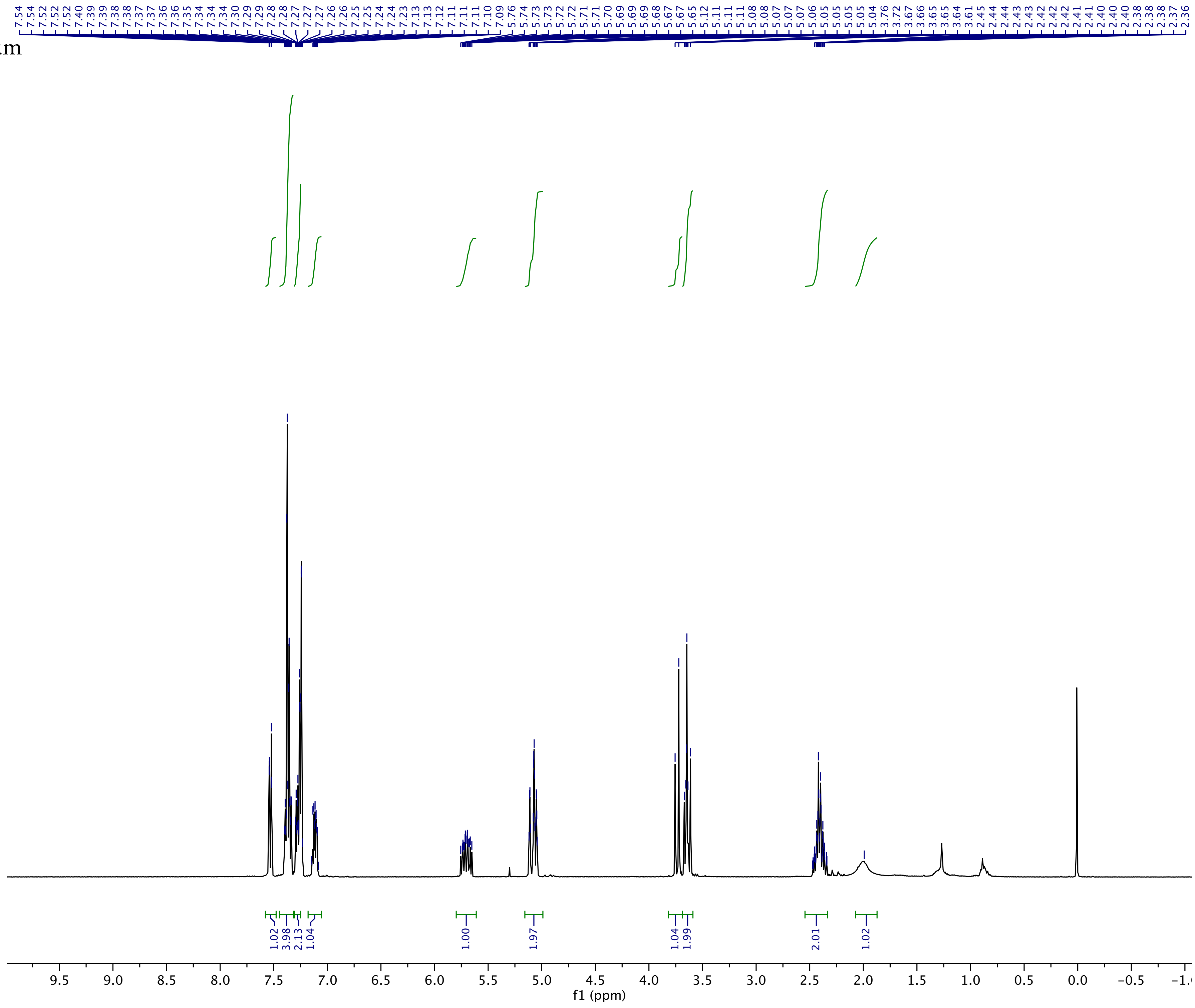

**S4aa** $^{13}$C NMR Spectrum

S4aa

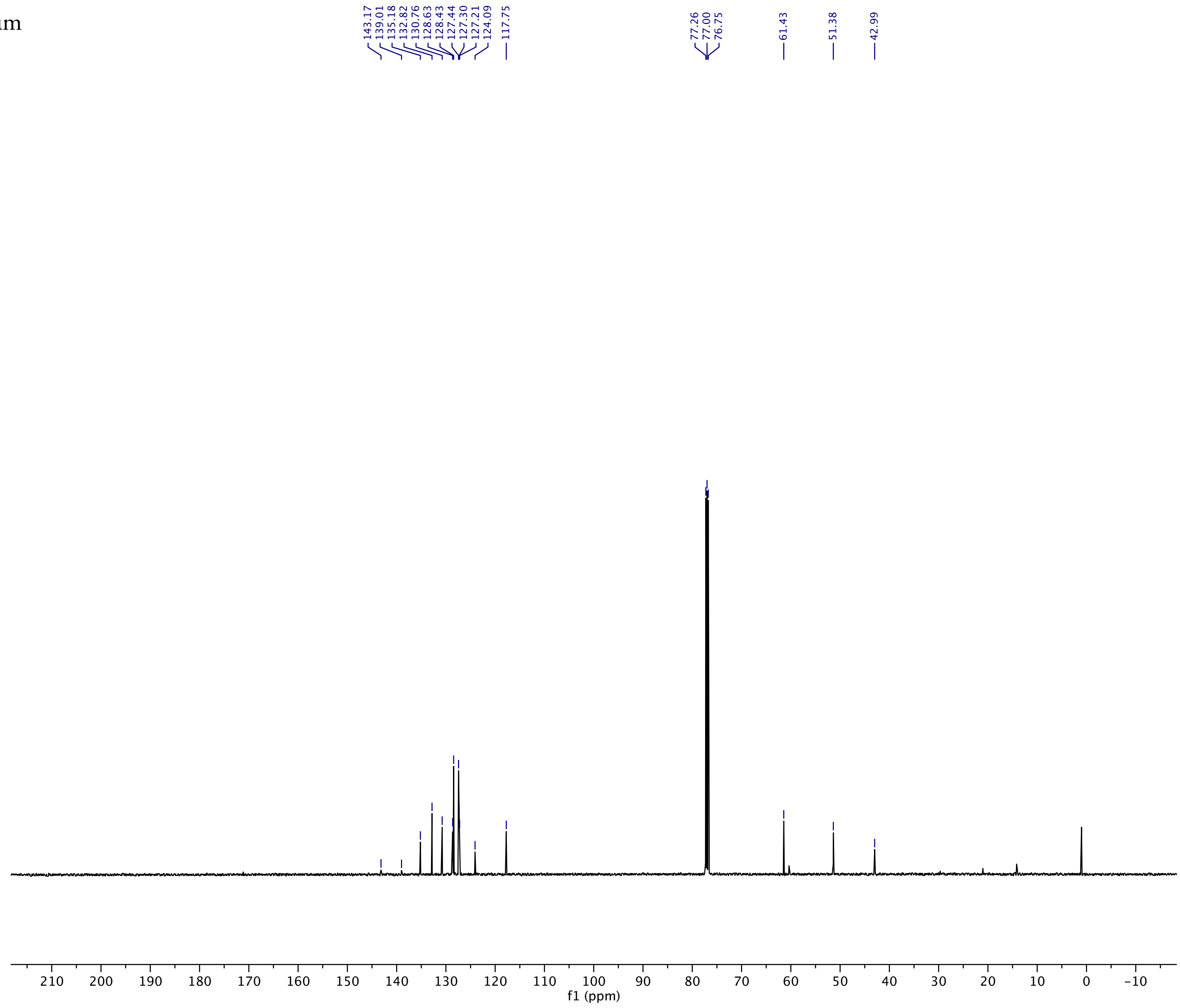

Current Data Parameters
NAME Aug09-2019
EXPNO 2
PROCNO 1

F2 - Acquisition Parameters
Date_ 20190809
Time 18.02 h
INSTRUM Spect
PROBHD Z109128_0101 (
PULPROG zgpg30
TD 65536
SOLVENT CDCl3
NS 2048
DS 4
SWH 29761.904 Hz
FIDRES 0.908261 Hz
AQ 1.1010048 sec
RG 203
DW 16.800 usec
DE 6.50 usec
TE 301.3 K
D1 2.00000000 sec
D11 0.03000000 sec
TD0 1
SFO1 125.7703643 MHz
NUC1 13C
P1 13.50 usec
PLW1 76.00000000 W
SFO2 500.1320005 MHz
NUC2 1H
CPDPRG[2 waltz16
PCPD2 80.00 usec
PLW2 18.00000000 W
PLW12 0.43945000 W
PLW13 0.22104000 W

F2 - Processing parameters
SI 32768
SF 125.7577900 MHz
WDW EM
SSB 0
LB 1.00 Hz
GB 0
PC 1.40

**S4bb** $^{1}$H NMR Spectrum

S4bb

NAME May18-2018
EXPNO 21
PROCNO 1
Date_ 20180518
Time 15.29
INSTRUM spect
PROBHD 5 mm DUL 13C-1
PULPROG zg30
TD 65536
SOLVENT CDCl3
NS 16
DS 2
SWH 8278.146 Hz
FIDRES 0.126314 Hz
AQ 3.9584243 sec
RG 128
DW 60.400 usec
DE 6.50 usec
TE 299.6 K
D1 1.00000000 sec
TD0 1

======== CHANNEL f1 ========
NUC1 1H
P1 12.58 usec
PL1 0.00 dB
PL1W 10.87646866 W
SFO1 400.1324710 MHz
SI 32768
SF 400.1300106 MHz
WDW EM
SSB 0
LB 0.30 Hz
GB 0
PC 1.00

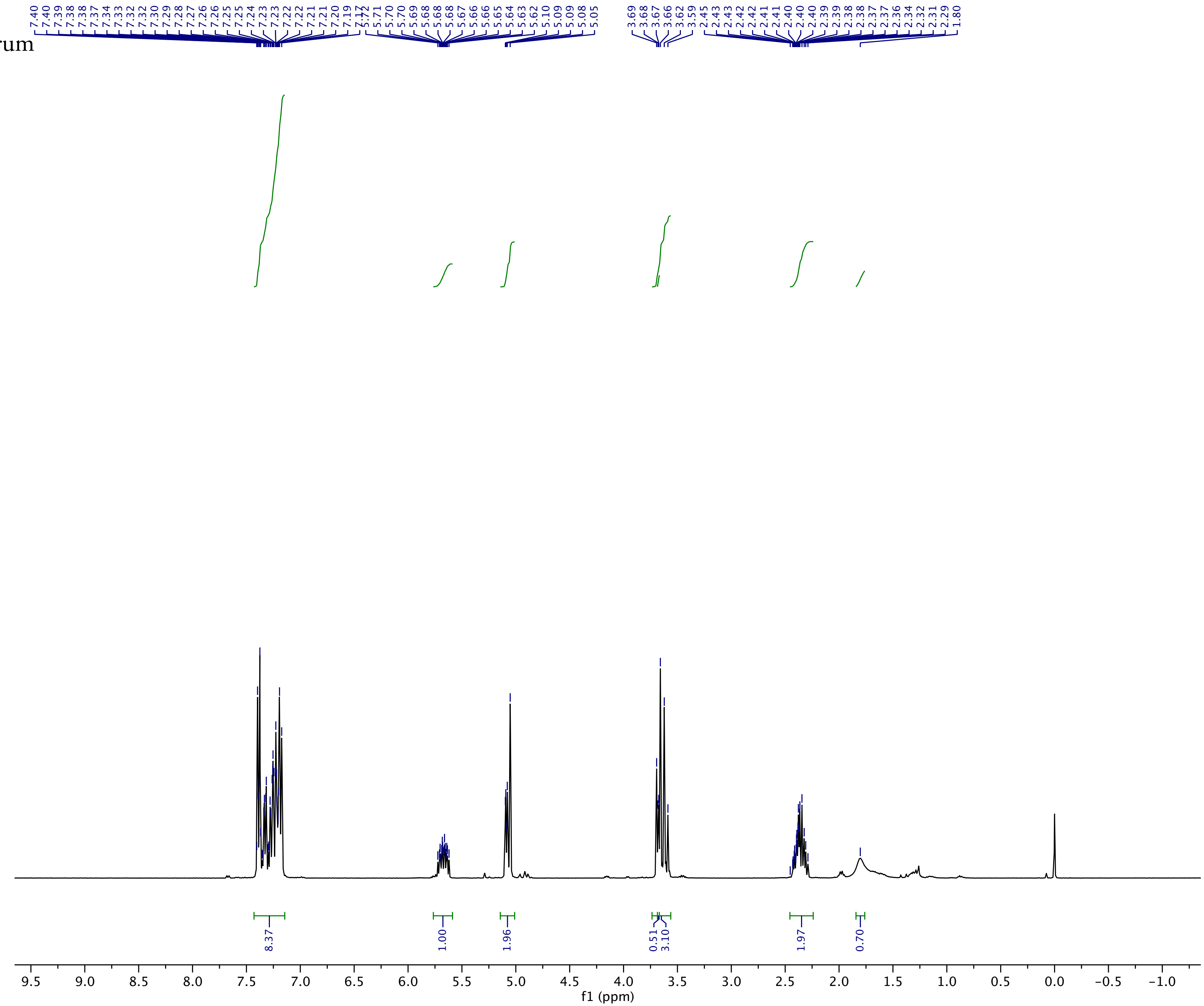

**S4bb** $^{13}$C NMR Spectrum

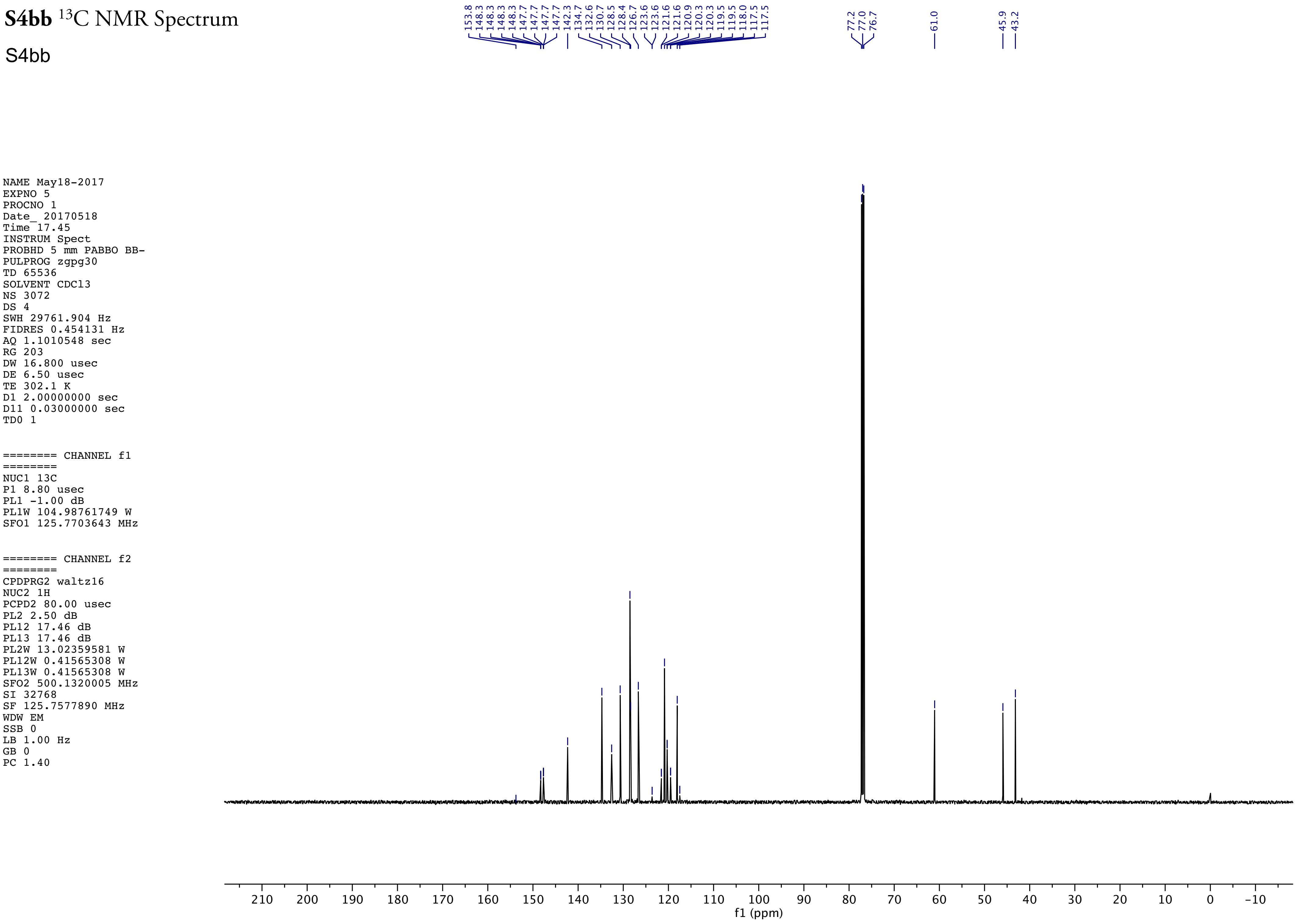

**S4bb** $^{19}$F NMR Spectrum

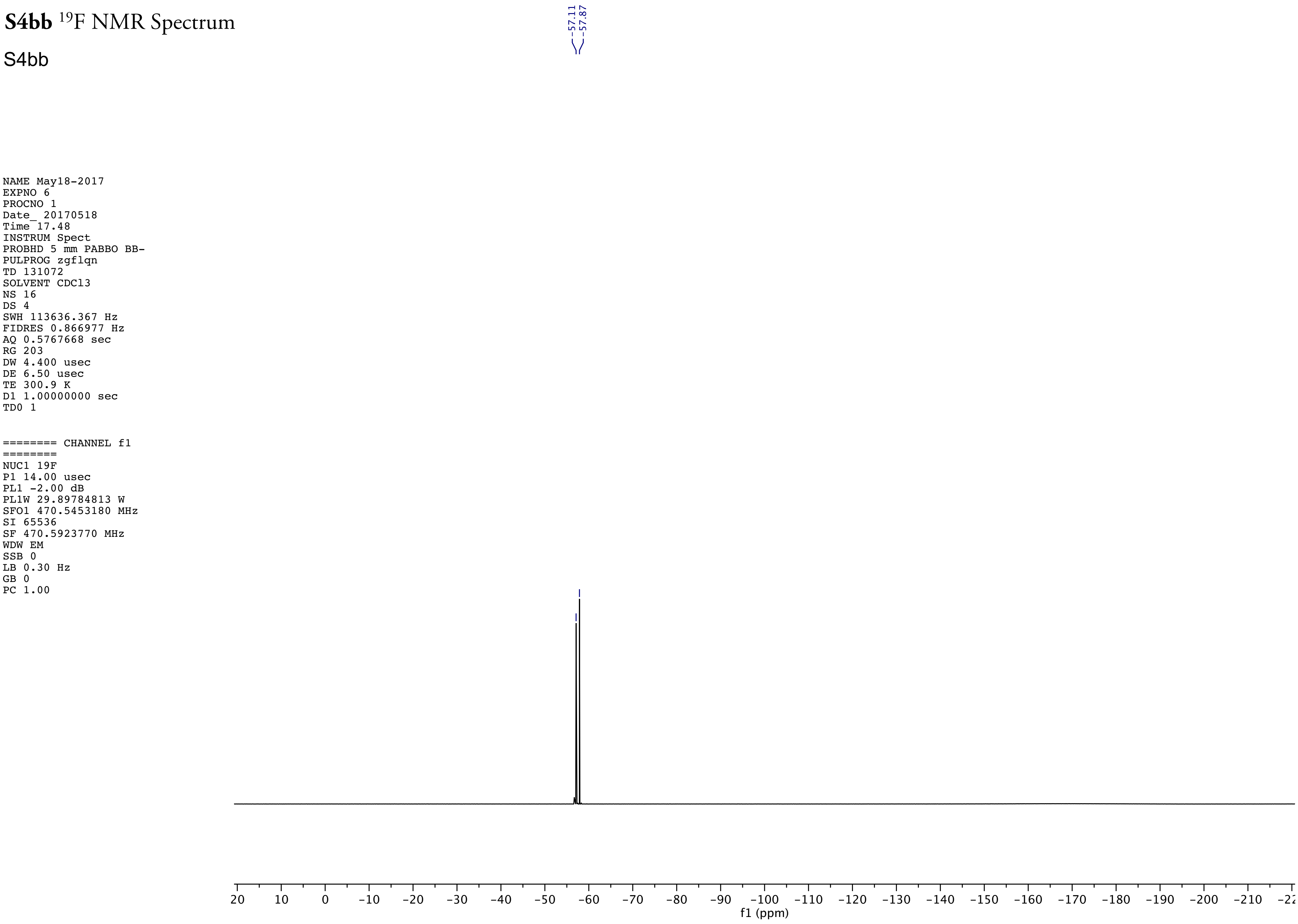

NAME May18-2017
EXPNO 6
PROCNO 1
Date_ 20170518
Time 17.48
INSTRUM Spect
PROBHD 5 mm PABBO BB-
PULPROG zgflqn
TD 131072
SOLVENT CDCl3
NS 16
DS 4
SWH 113636.367 Hz
FIDRES 0.866977 Hz
AQ 0.5767668 sec
RG 203
DW 4.400 usec
DE 6.50 usec
TE 300.9 K
D1 1.00000000 sec
TD0 1

======== CHANNEL f1 ========
NUC1 19F
P1 14.00 usec
PL1 -2.00 dB
PL1W 29.89784813 W
SFO1 470.5453180 MHz
SI 65536
SF 470.5923770 MHz
WDW EM
SSB 0
LB 0.30 Hz
GB 0
PC 1.00

**S4ca** ¹H NMR Spectrum

S4ca

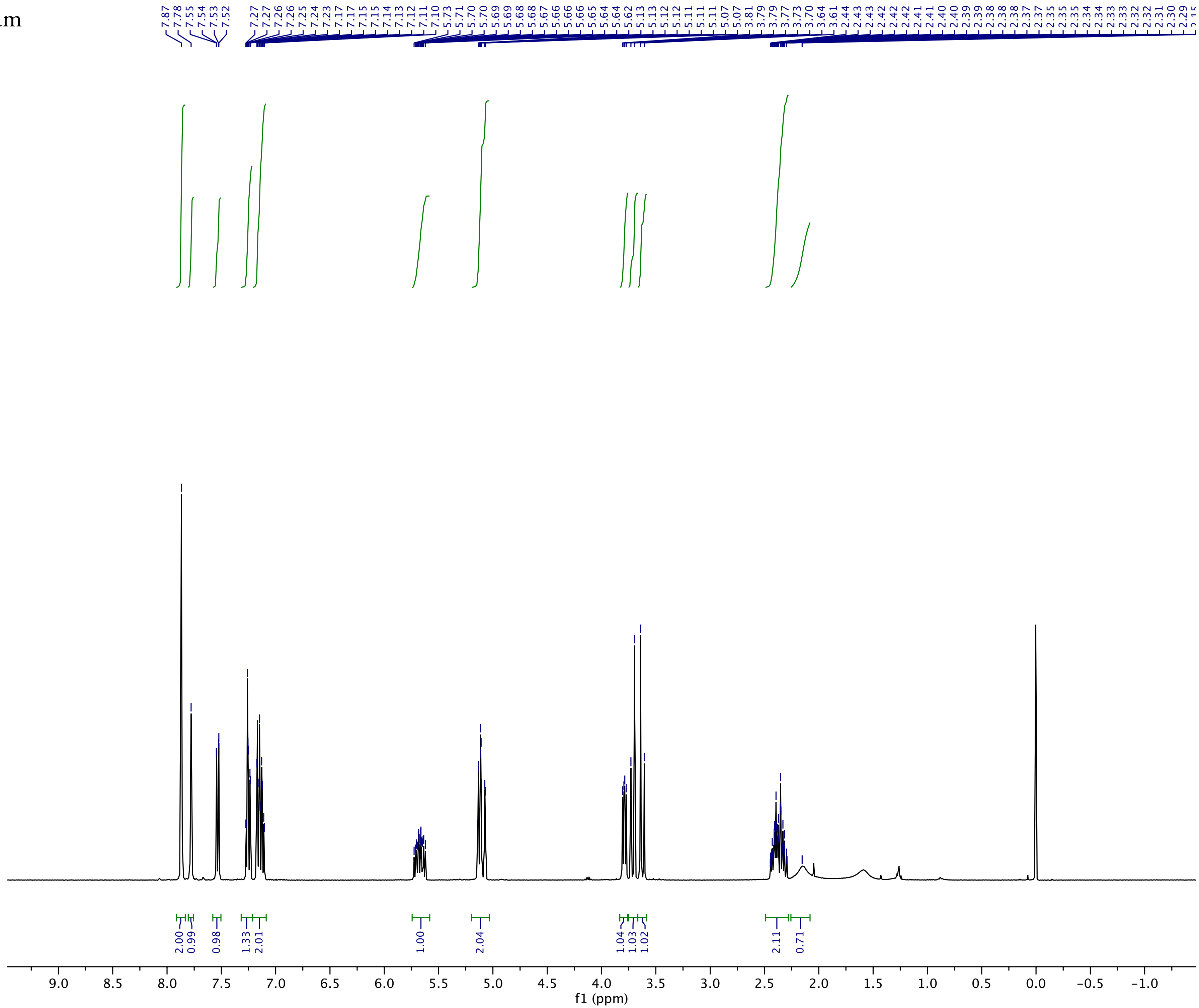

NAME Apr12-2017
EXPNO 41
PROCNO 1
Date_ 20170412
Time 14.27
INSTRUM spect
PROBHD 5 mm DUL 13C-1
PULPROG zg30
TD 65536
SOLVENT CDCl3
NS 16
DS 2
SWH 8278.146 Hz
FIDRES 0.126314 Hz
AQ 3.9584243 sec
RG 161.3
DW 60.400 usec
DE 6.50 usec
TE 297.2 K
D1 1.00000000 sec
TD0 1

======== CHANNEL f1 ========
NUC1 1H
P1 12.58 usec
PL1 0.00 dB
PL1W 10.87646866 W
SFO1 400.1324710 MHz
SI 32768
SF 400.1300085 MHz
WDW EM
SSB 0
LB 0.30 Hz
GB 0
PC 1.00

**S4ca** $^{13}C$ NMR Spectrum

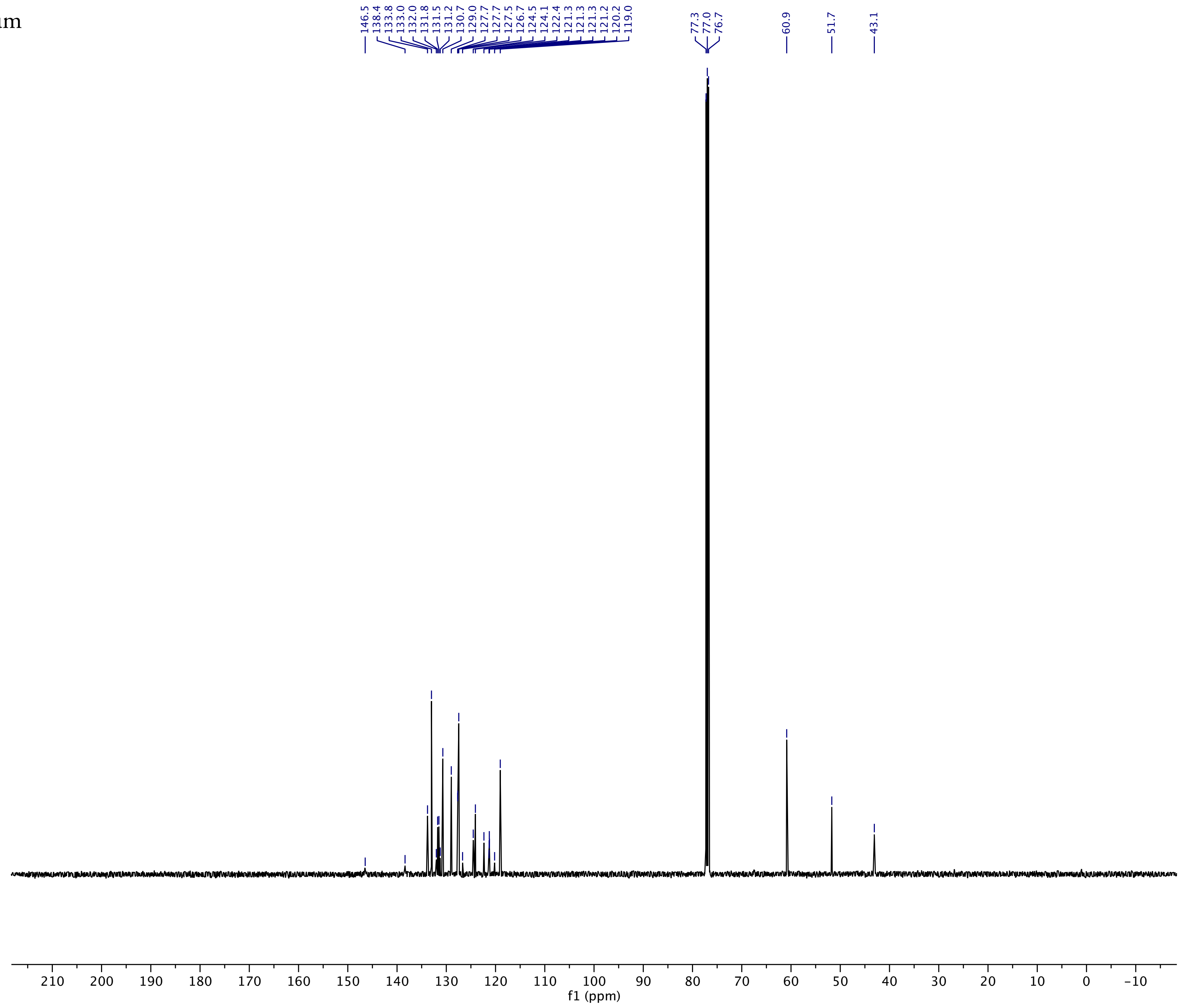

Current Data Parameters
NAME Jul24-2019
EXPNO 46
PROCNO 1

F2 - Acquisition Parameters
Date_ 20190725
Time 0.43 h
INSTRUM Spect
PROBHD Z109128_0101 (
PULPROG zgpg30
TD 65536
SOLVENT CDCl3
NS 1024
DS 4
SWH 29761.904 Hz
FIDRES 0.908261 Hz
AQ 1.1010048 sec
RG 203
DW 16.800 usec
DE 6.50 usec
TE 297.8 K
D1 2.00000000 sec
D11 0.03000000 sec
TD0 1
SFO1 125.7703643 MHz
NUC1 13C
P1 13.50 usec
PLW1 76.00000000 W
SFO2 500.1320005 MHz
NUC2 1H
CPDPRG[2 waltz16
PCPD2 80.00 usec
PLW2 18.00000000 W
PLW12 0.43945000 W
PLW13 0.22104000 W

F2 - Processing parameters
SI 32768
SF 125.7577909 MHz
WDW EM
SSB 0
LB 1.00 Hz
GB 0
PC 1.40

**S4ca** $^{19}$F NMR Spectrum

S4ca

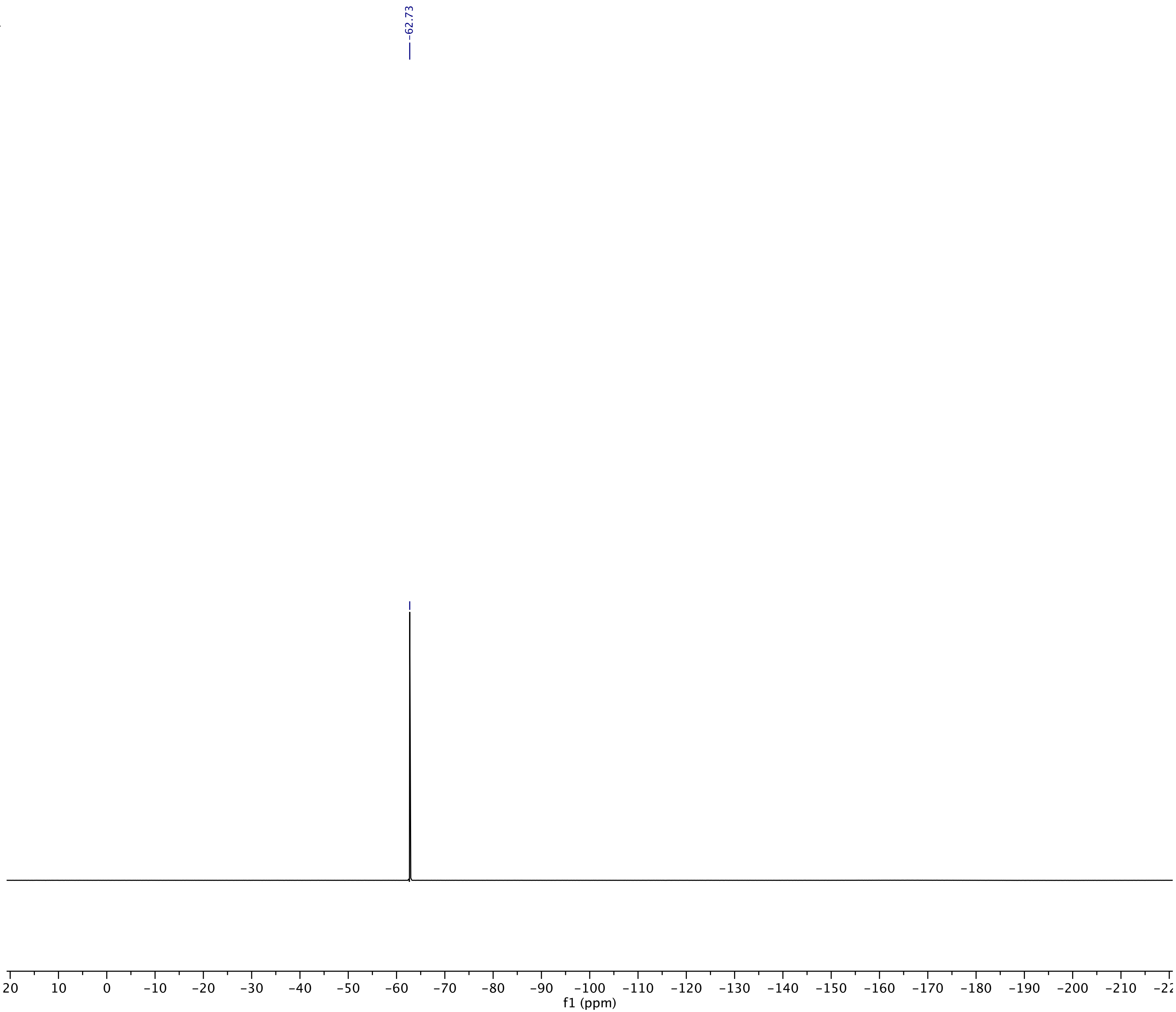

Current Data Parameters
NAME Jul24-2019
EXPNO 47
PROCNO 1

F2 - Acquisition Parameters
Date_ 20190725
Time 0.45 h
INSTRUM Spect
PROBHD Z109128_0101 (
PULPROG zgflqn
TD 131072
SOLVENT CDCl3
NS 16
DS 4
SWH 113636.367 Hz
FIDRES 1.733953 Hz
AQ 0.5767168 sec
RG 203
DW 4.400 usec
DE 6.50 usec
TE 297.1 K
D1 1.00000000 sec
TD0 1
SFO1 470.5453180 MHz
NUC1 19F
P1 18.00 usec
PLW1 20.00000000 W

F2 - Processing parameters
SI 65536
SF 470.5923770 MHz
WDW EM
SSB 0
LB 0.30 Hz
GB 0
PC 1.00

**S4da** ¹H NMR Spectrum

S4da

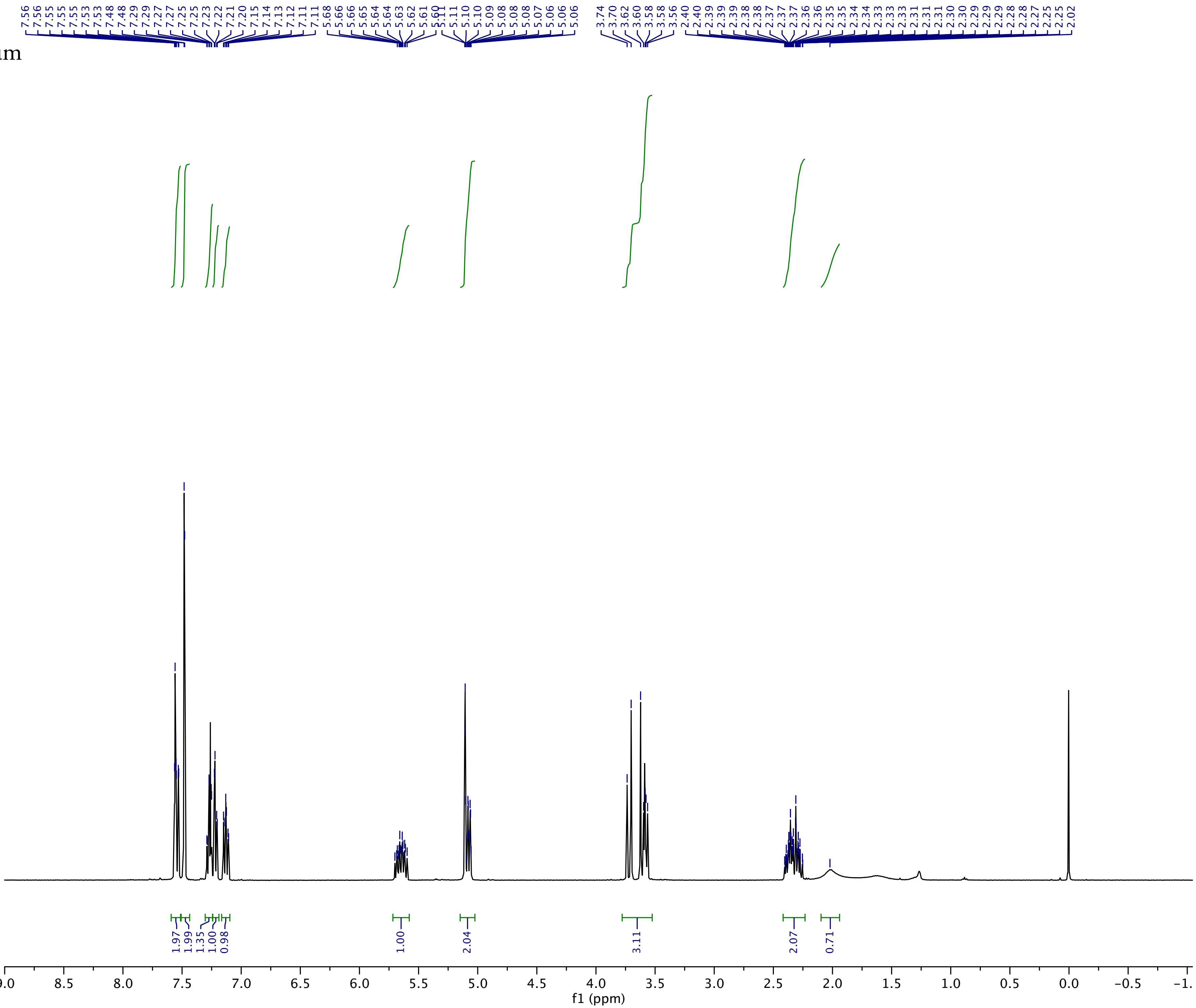

NAME Apr12-2017
EXPNO 42
PROCNO 1
Date_ 20170412
Time 14.33
INSTRUM spect
PROBHD 5 mm DUL 13C-1
PULPROG zg30
TD 65536
SOLVENT CDCl3
NS 16
DS 2
SWH 8278.146 Hz
FIDRES 0.126314 Hz
AQ 3.9584243 sec
RG 161.3
DW 60.400 usec
DE 6.50 usec
TE 297.2 K
D1 1.00000000 sec
TD0 1

======== CHANNEL f1 ========
NUC1 1H
P1 12.58 usec
PL1 0.00 dB
PL1W 10.87646866 W
SFO1 400.1324710 MHz
SI 32768
SF 400.1300083 MHz
WDW EM
SSB 0
LB 0.30 Hz
GB 0
PC 1.00

**S4da** $^{13}C$ NMR Spectrum

S4da

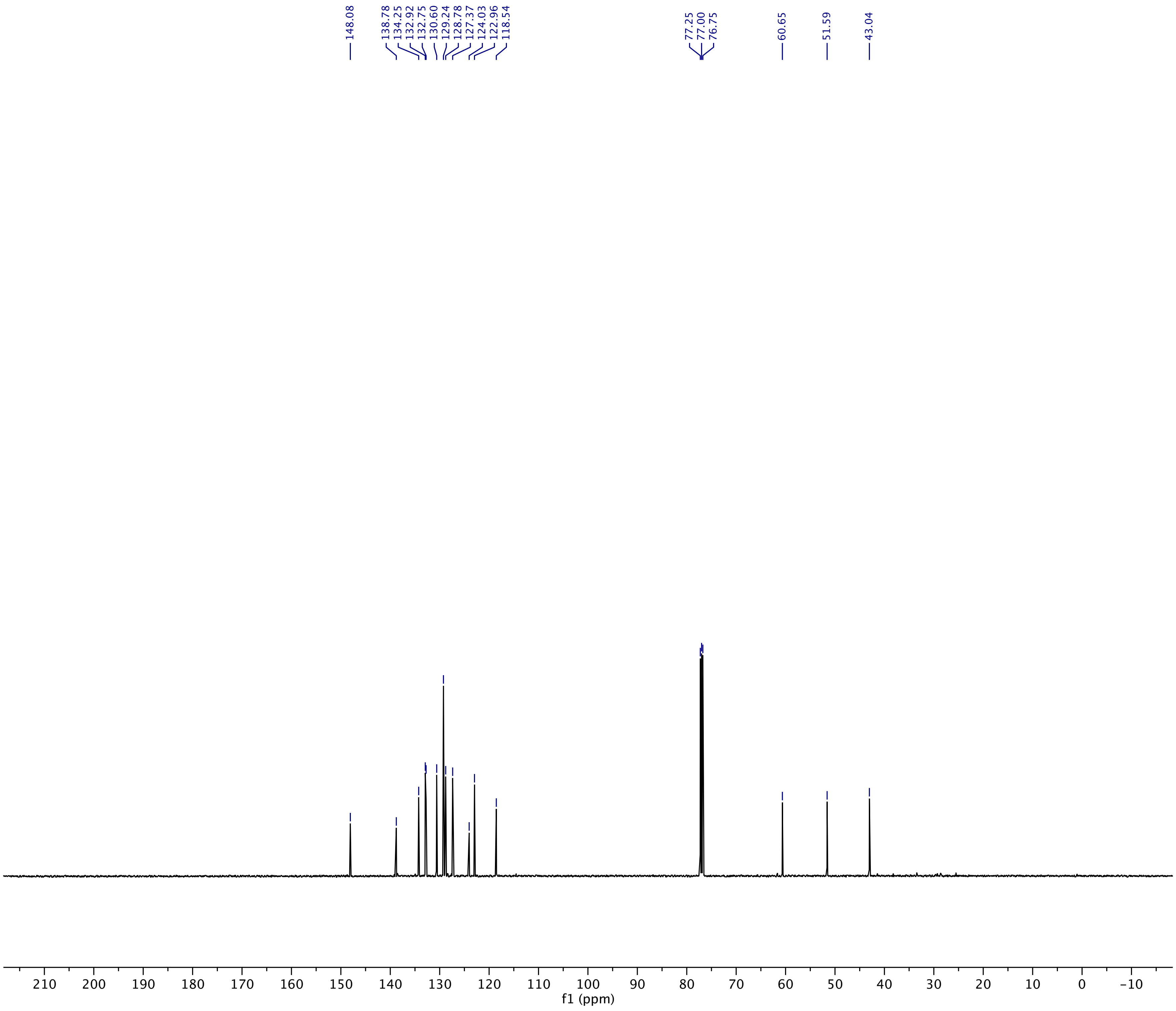

Current Data Parameters
NAME 0086
EXPNO 2
PROCNO 1

F2 - Acquisition Parameters
Date_ 20200726
Time 12.00 h
INSTRUM Spect
PROBHD Z109128_0101 (
PULPROG zgpg30
TD 65536
SOLVENT CDCl3
NS 1024
DS 4
SWH 29761.904 Hz
FIDRES 0.908261 Hz
AQ 1.1010048 sec
RG 203
DW 16.800 usec
DE 6.50 usec
TE 300.2 K
D1 2.00000000 sec
D11 0.03000000 sec
TD0 1
SFO1 125.7703643 MHz
NUC1 13C
P1 13.50 usec
PLW1 76.00000000 W
SFO2 500.1320005 MHz
NUC2 1H
CPDPRG[2 waltz16
PCPD2 80.00 usec
PLW2 18.00000000 W
PLW12 0.43945000 W
PLW13 0.22104000 W

F2 - Processing parameters
SI 32768
SF 125.7577938 MHz
WDW EM
SSB 0
LB 1.00 Hz
GB 0
PC 1.40

**BGAz-001** $^{1}$H NMR spectrum

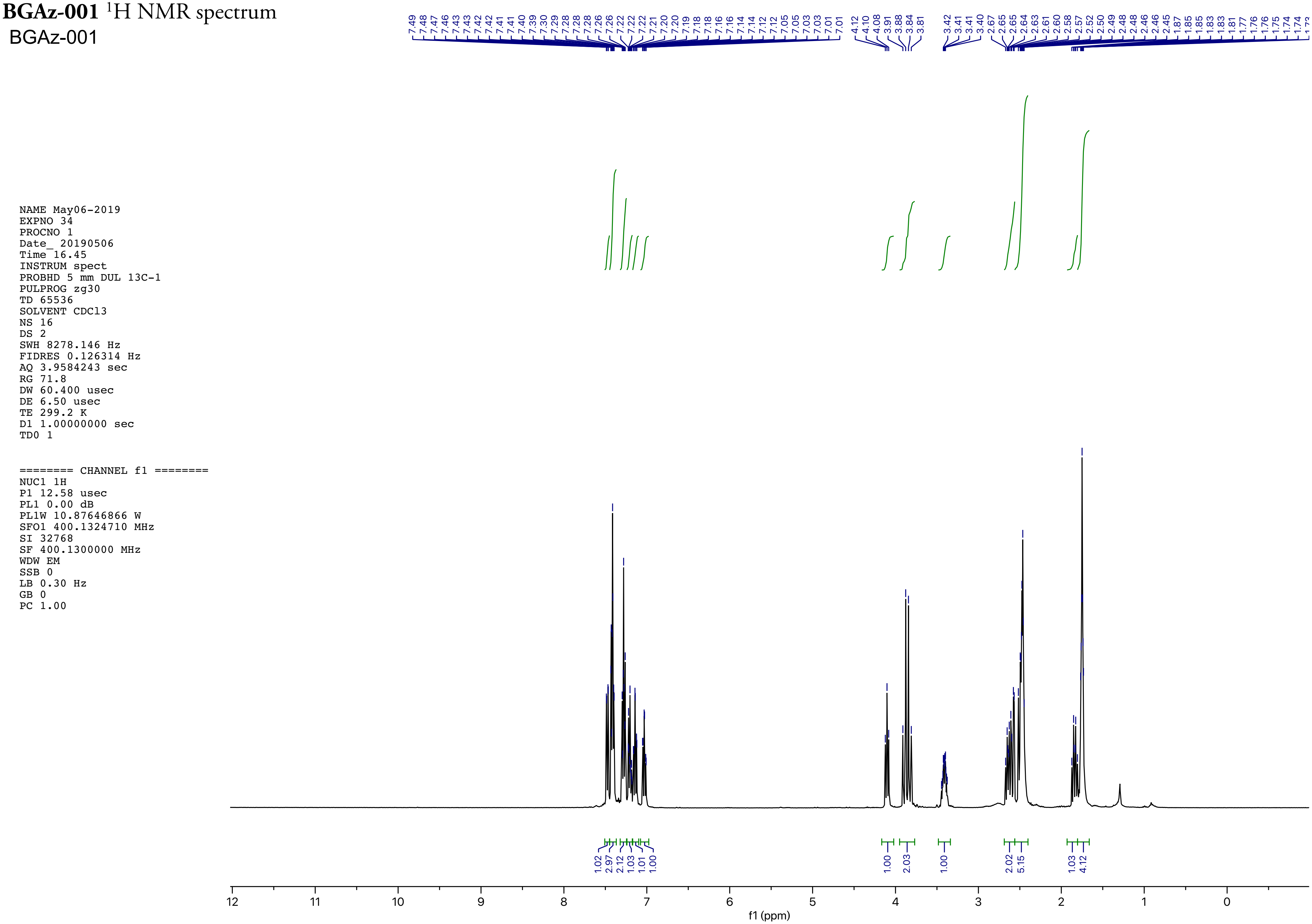

NAME May06-2019
EXPNO 34
PROCNO 1
Date_ 20190506
Time 16.45
INSTRUM spect
PROBHD 5 mm DUL 13C-1
PULPROG zg30
TD 65536
SOLVENT CDCl3
NS 16
DS 2
SWH 8278.146 Hz
FIDRES 0.126314 Hz
AQ 3.9584243 sec
RG 71.8
DW 60.400 usec
DE 6.50 usec
TE 299.2 K
D1 1.00000000 sec
TD0 1

======== CHANNEL f1 ========
NUC1 1H
P1 12.58 usec
PL1 0.00 dB
PL1W 10.87646866 W
SFO1 400.1324710 MHz
SI 32768
SF 400.1300000 MHz
WDW EM
SSB 0
LB 0.30 Hz
GB 0
PC 1.00

**BGAz-001** $^{1}$H NMR spectrum (zoom 1)

BGAz-001

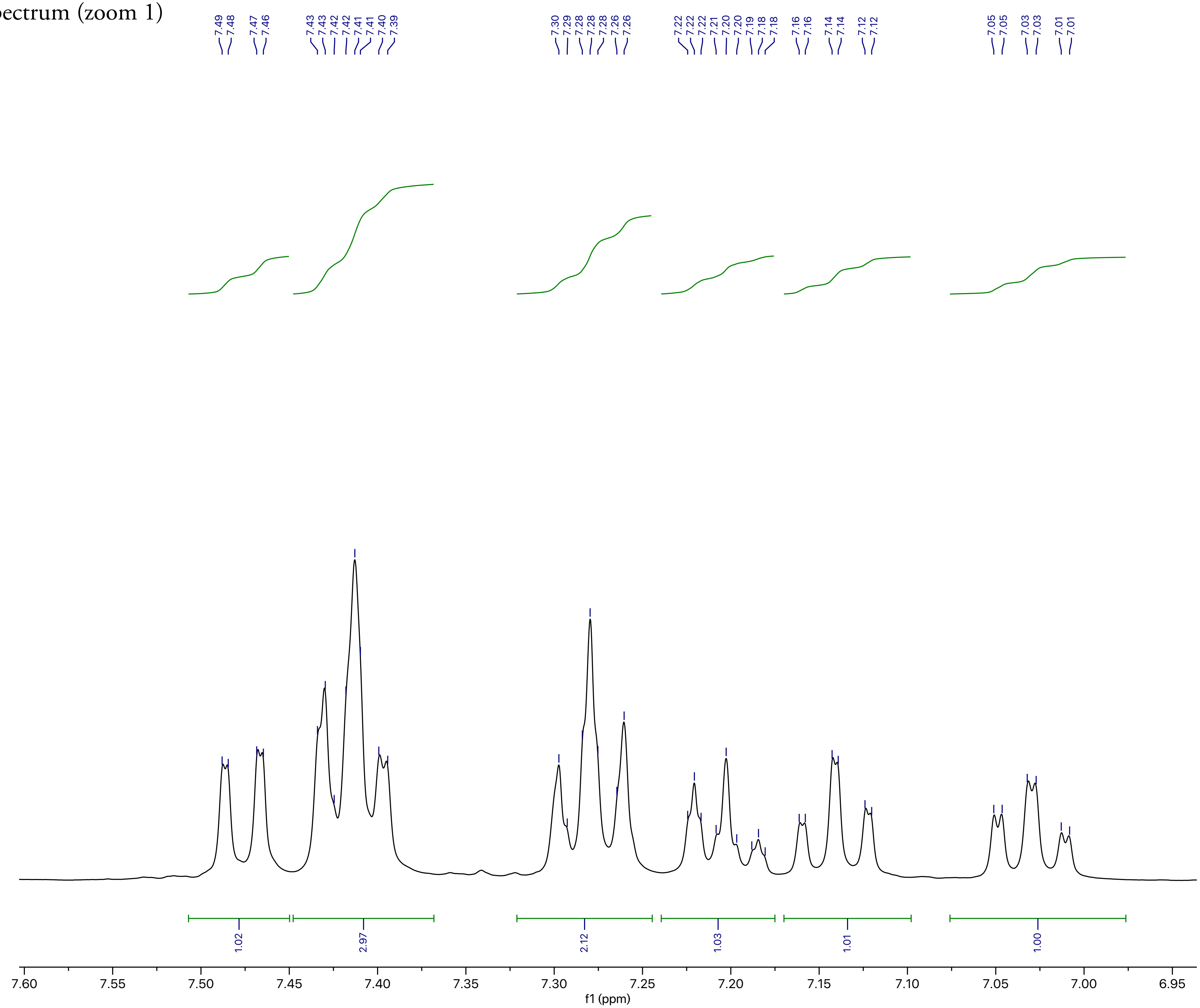

```
NAME May06-2019
EXPNO 34
PROCNO 1
Date_ 20190506
Time 16.45
INSTRUM spect
PROBHD 5 mm DUL 13C-1
PULPROG zg30
TD 65536
SOLVENT CDCl3
NS 16
DS 2
SWH 8278.146 Hz
FIDRES 0.126314 Hz
AQ 3.9584243 sec
RG 71.8
DW 60.400 usec
DE 6.50 usec
TE 299.2 K
D1 1.00000000 sec
TD0 1

======== CHANNEL f1 ========
NUC1 1H
P1 12.58 usec
PL1 0.00 dB
PL1W 10.87646866 W
SFO1 400.1324710 MHz
SI 32768
SF 400.1300000 MHz
WDW EM
SSB 0
LB 0.30 Hz
GB 0
PC 1.00
```

**BGAz-001** $^1$H NMR spectrum (zoom 2)

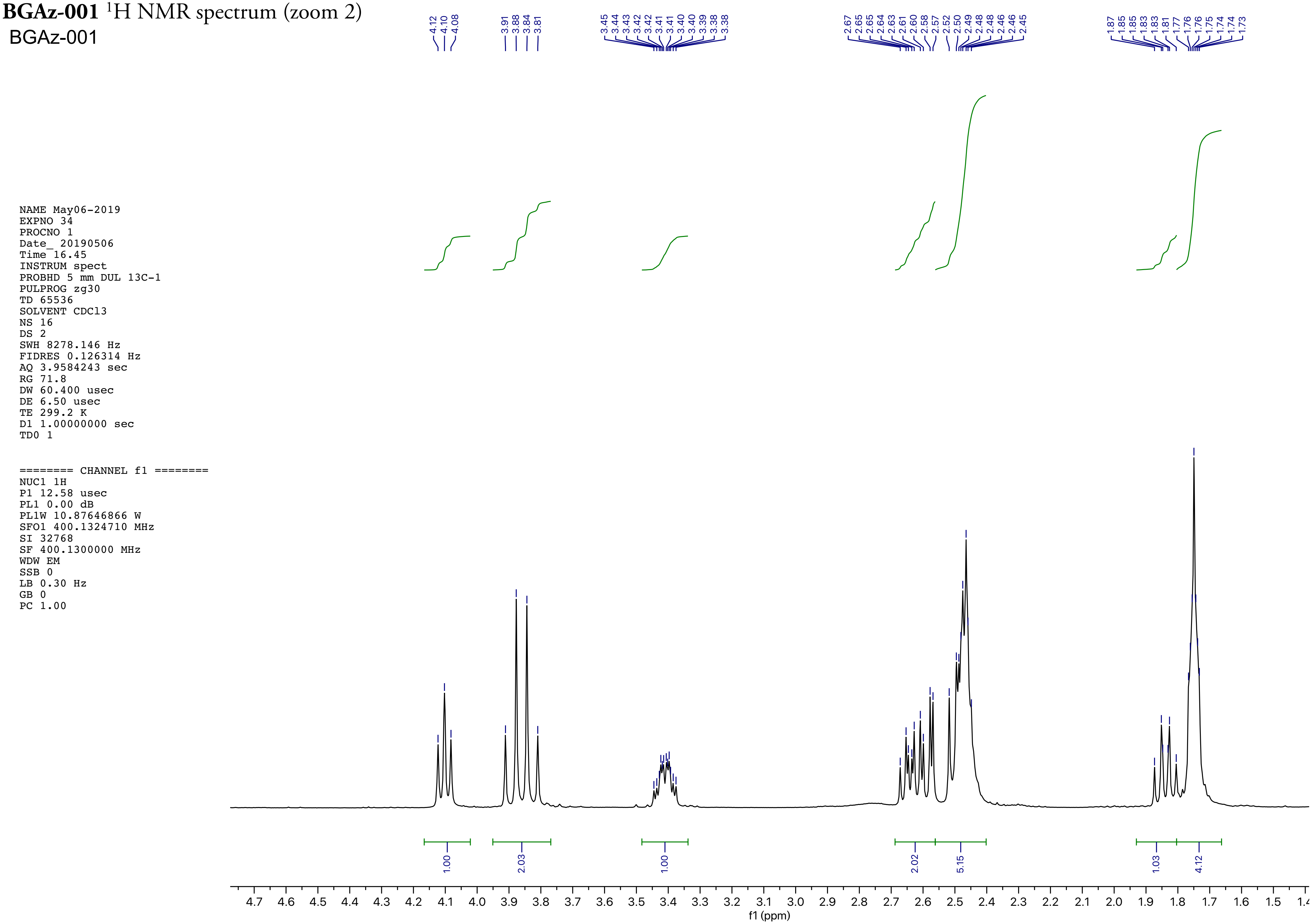

NAME May06-2019
EXPNO 34
PROCNO 1
Date_ 20190506
Time 16.45
INSTRUM spect
PROBHD 5 mm DUL 13C-1
PULPROG zg30
TD 65536
SOLVENT CDCl3
NS 16
DS 2
SWH 8278.146 Hz
FIDRES 0.126314 Hz
AQ 3.9584243 sec
RG 71.8
DW 60.400 usec
DE 6.50 usec
TE 299.2 K
D1 1.00000000 sec
TD0 1

======== CHANNEL f1 ========
NUC1 1H
P1 12.58 usec
PL1 0.00 dB
PL1W 10.87646866 W
SFO1 400.1324710 MHz
SI 32768
SF 400.1300000 MHz
WDW EM
SSB 0
LB 0.30 Hz
GB 0
PC 1.00

**BGAz-001** $^{13}$C NMR spectrum

BGAz-001

Current Data Parameters
NAME May06-2019
EXPNO 7
PROCNO 1

F2 - Acquisition Parameters
Date_ 20190506
Time 20.49 h
INSTRUM Spect
PROBHD Z109128_0101 (
PULPROG zgpg30
TD 65536
SOLVENT CDCl3
NS 4096
DS 4
SWH 29761.904 Hz
FIDRES 0.908261 Hz
AQ 1.1010048 sec
RG 203
DW 16.800 usec
DE 6.50 usec
TE 297.7 K
D1 2.00000000 sec
D11 0.03000000 sec
TD0 1
SFO1 125.7703643 MHz
NUC1 13C
P1 13.50 usec
PLW1 76.00000000 W
SFO2 500.1320005 MHz
NUC2 1H
CPDPRG[2 waltz16
PCPD2 80.00 usec
PLW2 18.00000000 W
PLW12 0.43945000 W
PLW13 0.22104000 W

F2 - Processing parameters
SI 32768
SF 125.7577976 MHz
WDW EM
SSB 0
LB 1.00 Hz
GB 0
PC 1.40

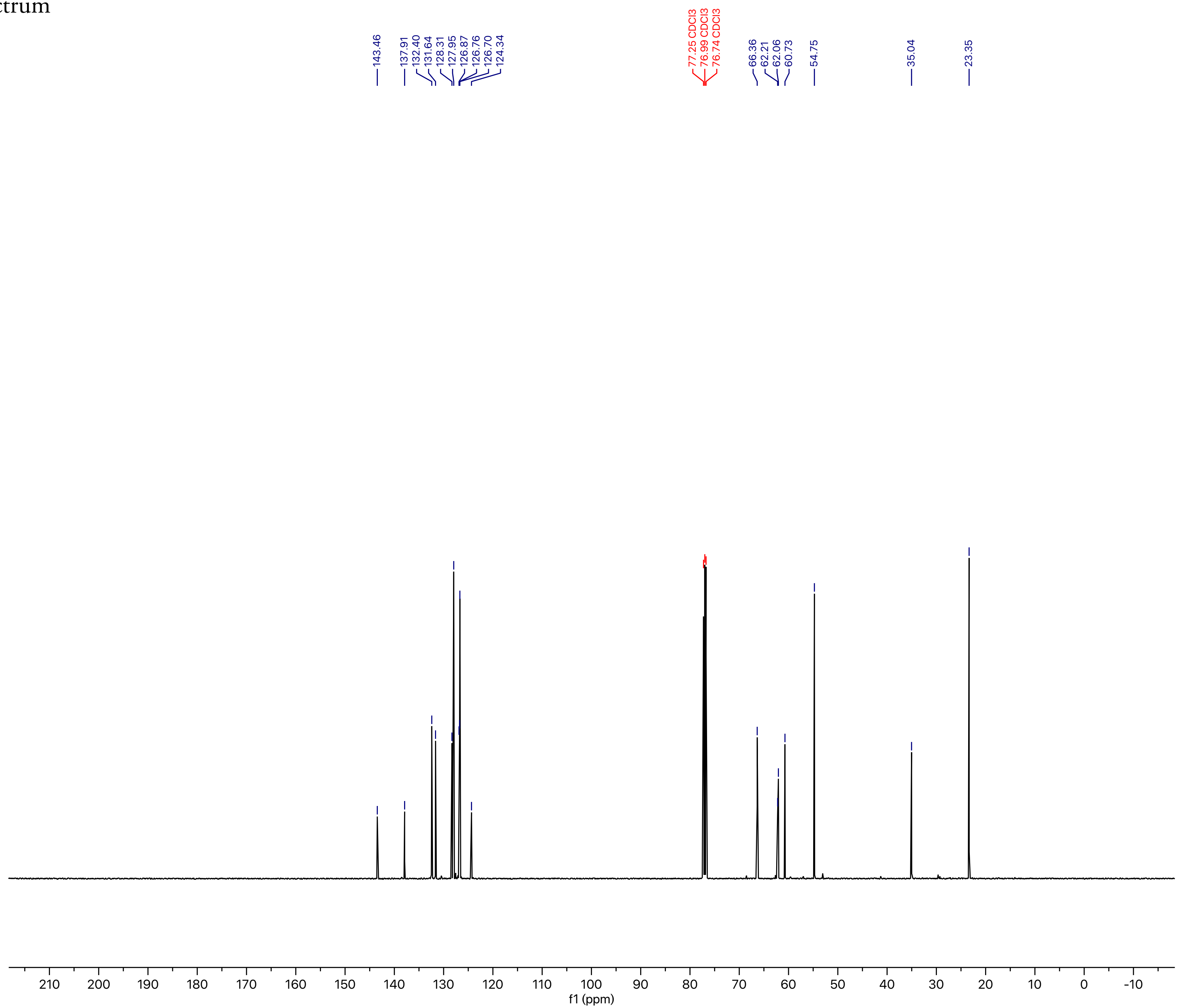

**BGAz-002** ¹H NMR Spectrum

BGAz-002

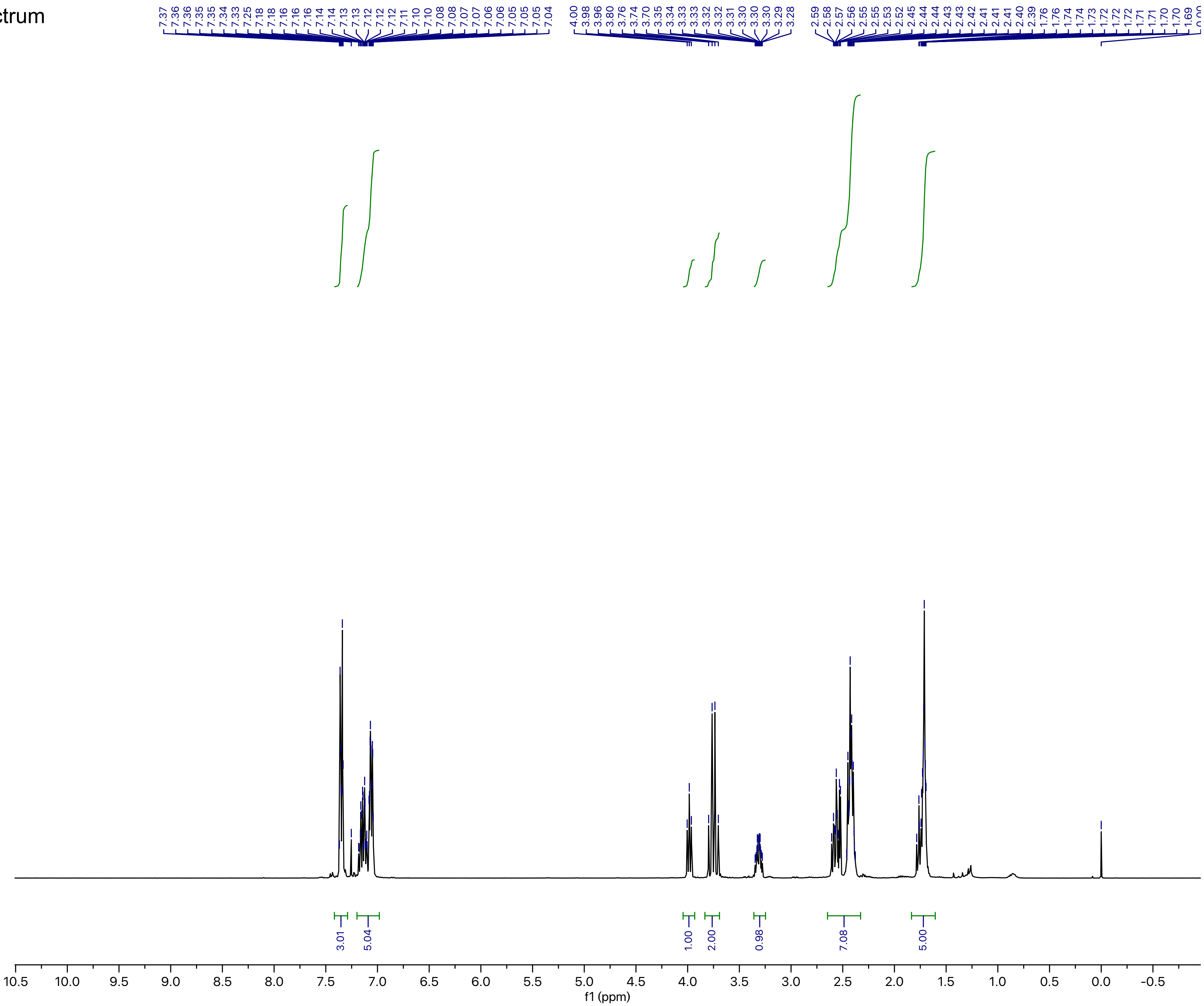

Current Data Parameters
NAME YC-GIBH-A08
EXPNO 10
PROCNO 1

F2 - Acquisition Parameters
Date_ 20190405
Time 8.20 h
INSTRUM AvanceNeo
PROBHD Z116098_0793 (
PULPROG zg30
TD 65536
SOLVENT CDCl3
NS 2
DS 0
SWH 7142.857 Hz
FIDRES 0.217983 Hz
AQ 4.5875201 sec
RG 32
DW 70.000 usec
DE 14.62 usec
TE 298.0 K
D1 2.00000000 sec
TD0 1
SFO1 400.1324008 MHz
NUC1 1H
P0 3.33 usec
P1 10.00 usec
PLW1 18.69700050 W

F2 - Processing parameters
SI 131072
SF 400.1300126 MHz
WDW EM
SSB 0
LB 0.10 Hz
GB 0
PC 1.00

**BGAz-002** [1]H NMR Spectrum (zoom 1)

BGAz-002

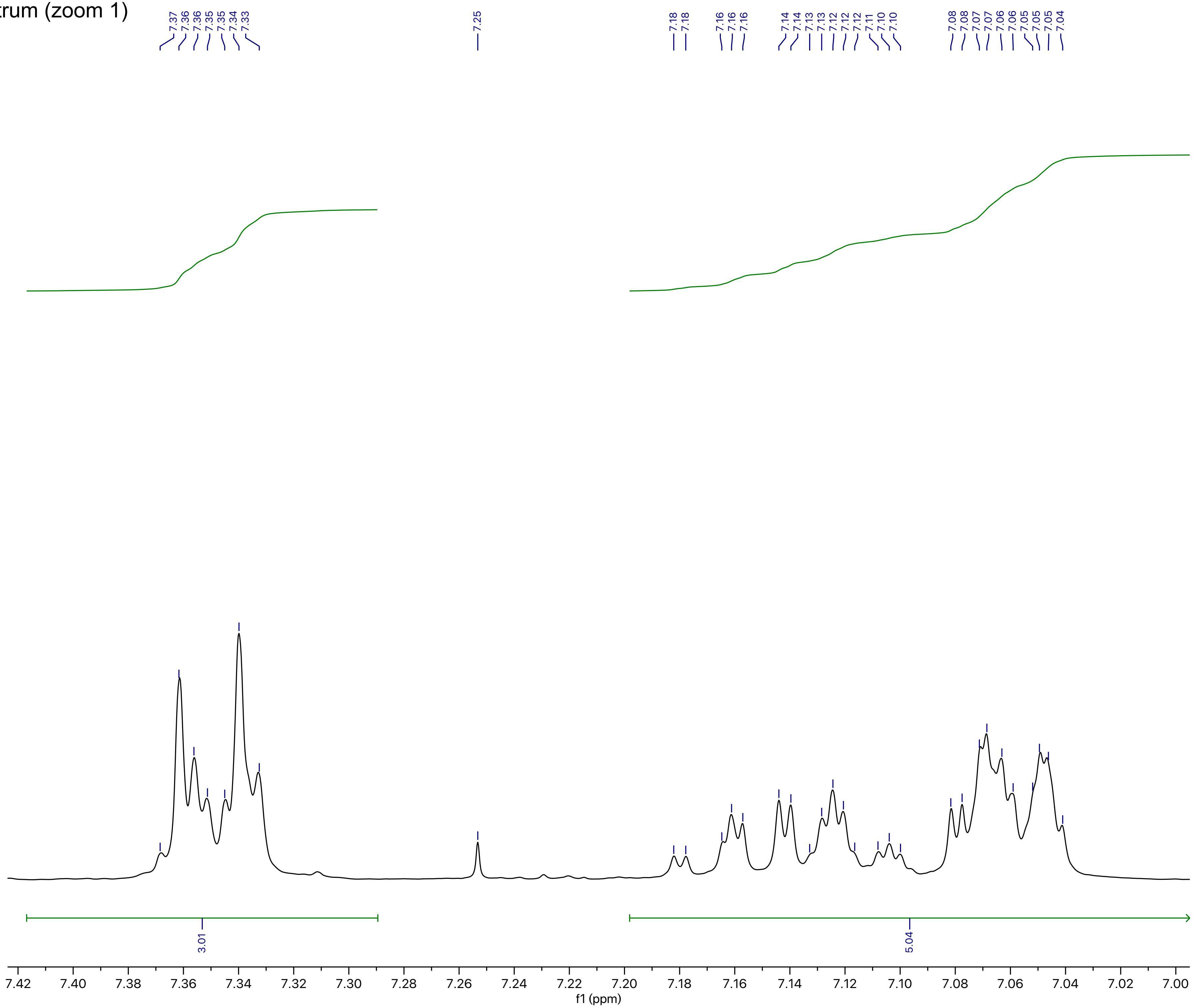

Current Data Parameters
NAME YC-GIBH-A08
EXPNO 10
PROCNO 1

F2 - Acquisition Parameters
Date_ 20190405
Time 8.20 h
INSTRUM AvanceNeo
PROBHD Z116098_0793 (
PULPROG zg30
TD 65536
SOLVENT CDCl3
NS 2
DS 0
SWH 7142.857 Hz
FIDRES 0.217983 Hz
AQ 4.5875201 sec
RG 32
DW 70.000 usec
DE 14.62 usec
TE 298.0 K
D1 2.00000000 sec
TD0 1
SFO1 400.1324008 MHz
NUC1 1H
P0 3.33 usec
P1 10.00 usec
PLW1 18.69700050 W

F2 - Processing parameters
SI 131072
SF 400.1300126 MHz
WDW EM
SSB 0
LB 0.10 Hz
GB 0
PC 1.00

**BGAz-002** $^{1}$H NMR Spectrum (zoom 2)

BGAz-002

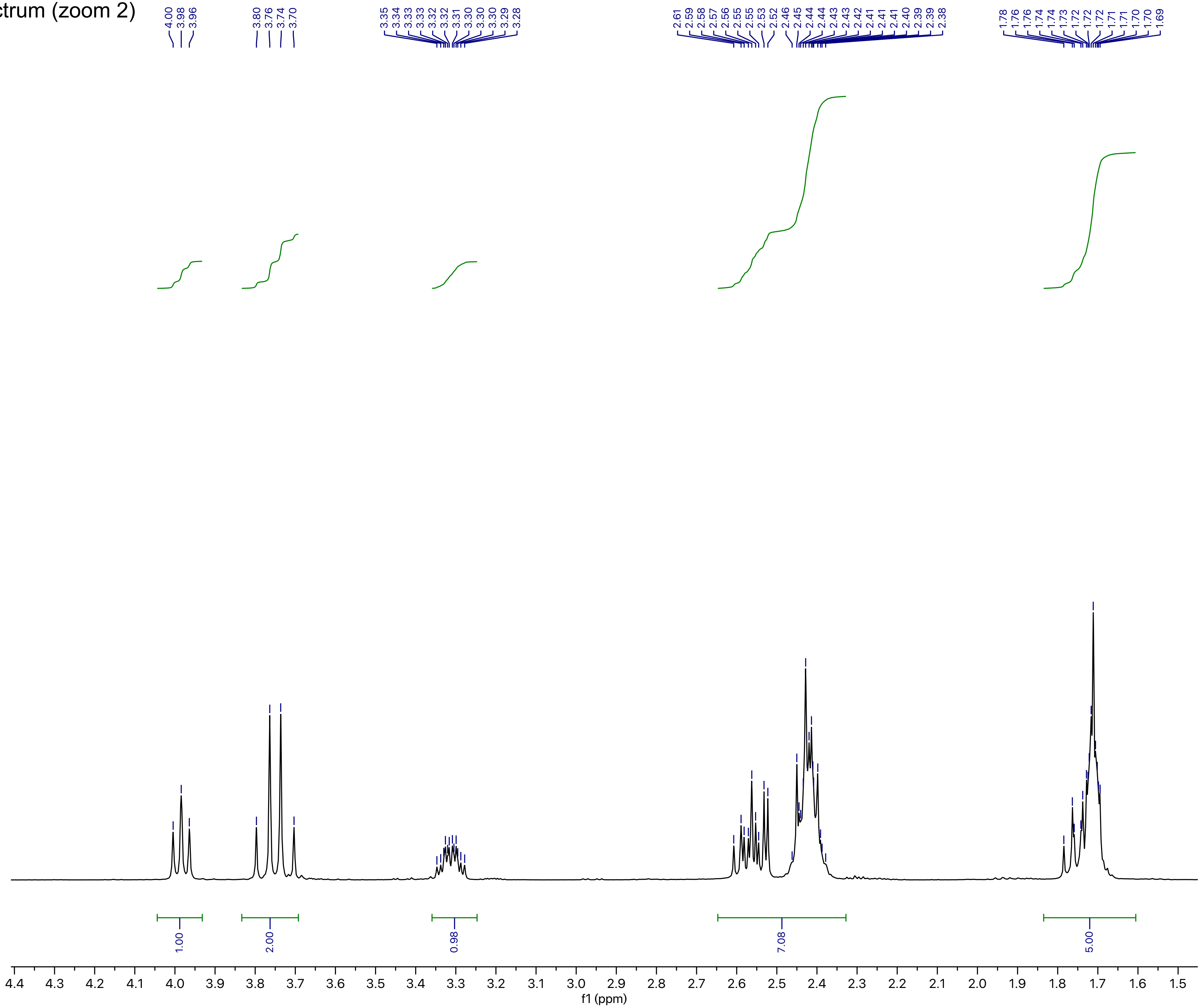

Current Data Parameters
NAME YC-GIBH-A08
EXPNO 10
PROCNO 1

F2 - Acquisition Parameters
Date_ 20190405
Time 8.20 h
INSTRUM AvanceNeo
PROBHD Z116098_0793 (
PULPROG zg30
TD 65536
SOLVENT CDCl3
NS 2
DS 0
SWH 7142.857 Hz
FIDRES 0.217983 Hz
AQ 4.5875201 sec
RG 32
DW 70.000 usec
DE 14.62 usec
TE 298.0 K
D1 2.00000000 sec
TD0 1
SFO1 400.1324008 MHz
NUC1 1H
P0 3.33 usec
P1 10.00 usec
PLW1 18.69700050 W

F2 - Processing parameters
SI 131072
SF 400.1300126 MHz
WDW EM
SSB 0
LB 0.10 Hz
GB 0
PC 1.00

**BGAz-002** $^{13}C$ NMR Spectrum

BGAz-002

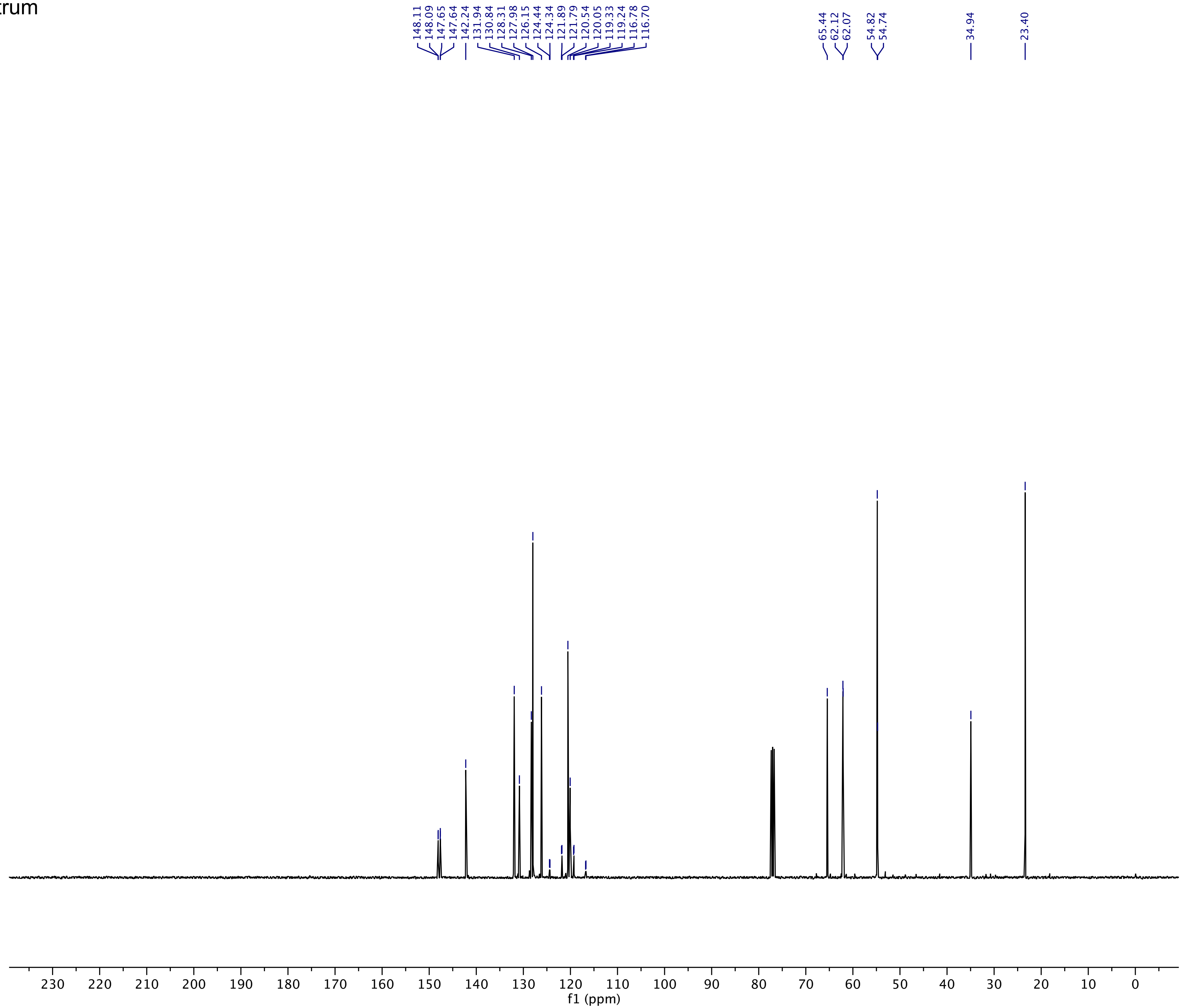

Current Data Parameters
NAME YC-GIBH-A08
EXPNO 11
PROCNO 1

F2 - Acquisition Parameters
Date_ 20190405
Time 8.50 h
INSTRUM AvanceNeo
PROBHD Z116098_0793 (
PULPROG zgpg30
TD 119044
SOLVENT CDCl3
NS 512
DS 0
SWH 25000.000 Hz
FIDRES 0.420013 Hz
AQ 2.3808801 sec
RG 31.9602
DW 20.000 usec
DE 7.12 usec
TE 298.0 K
D1 1.00000000 sec
D11 0.03000000 sec
TD0 1
SFO1 100.6243390 MHz
NUC1 13C
P0 3.33 usec
P1 10.00 usec
PLW1 83.92700195 W
SFO2 400.1318006 MHz
NUC2 1H
CPDPRG[2 waltz64
PCPD2 90.00 usec
PLW2 18.69700050 W
PLW12 0.23083000 W
PLW13 0.11611000 W

F2 - Processing parameters
SI 131072
SF 100.6127685 MHz
WDW EM
SSB 0
LB 1.00 Hz
GB 0
PC 1.40

**BGAz-002** $^{19}$F NMR Spectrum

BGAz-002

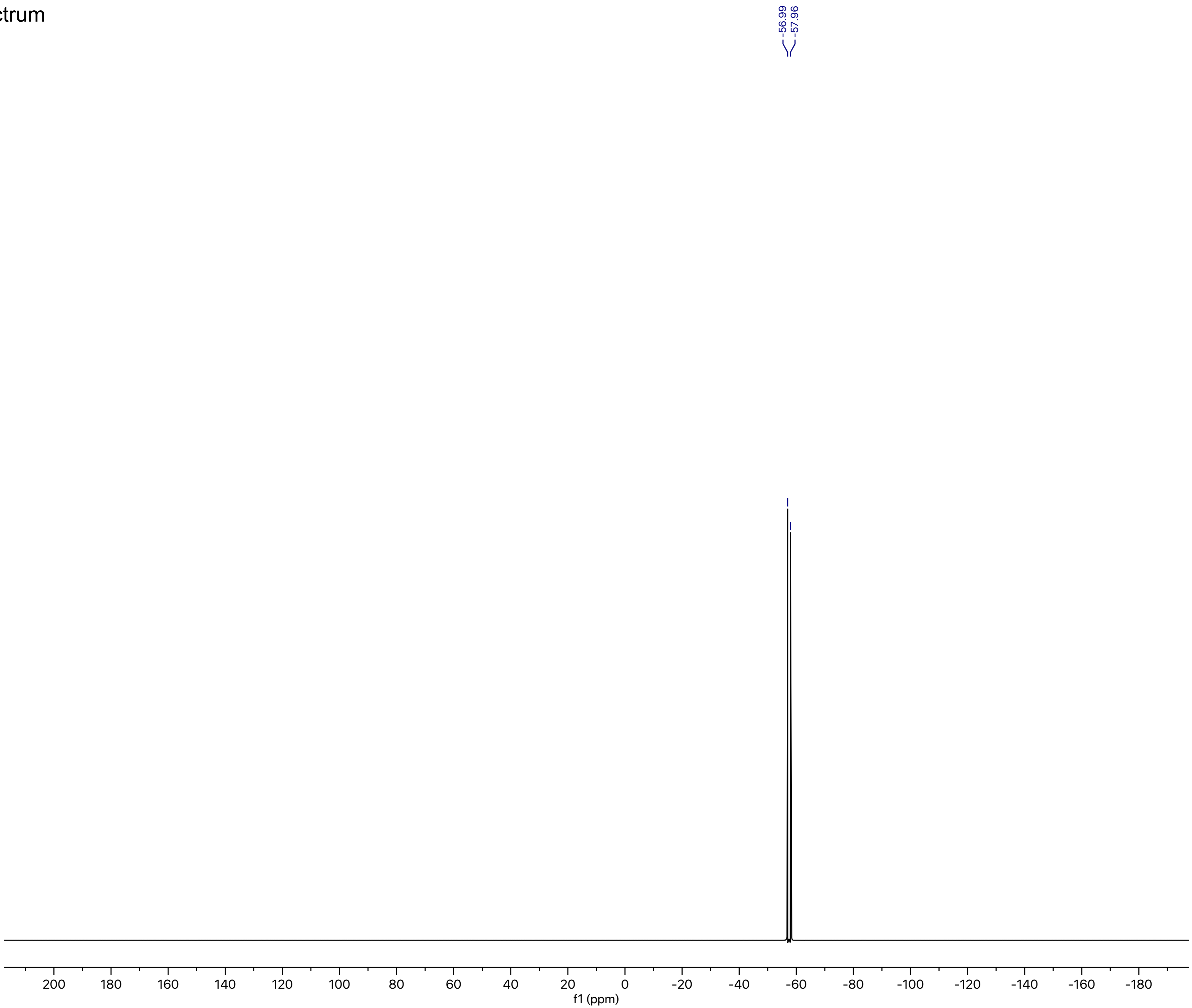

Current Data Parameters
NAME YC-GIBH-A08
EXPNO 13
PROCNO 1

F2 - Acquisition Parameters
Date_ 20190405
Time 9.12 h
INSTRUM AvanceNeo
PROBHD Z116098_0793 (
PULPROG zgig30
TD 261948
SOLVENT CDCl3
NS 16
DS 0
SWH 156250.000 Hz
FIDRES 1.192985 Hz
AQ 0.8382336 sec
RG 101
DW 3.200 usec
DE 6.82 usec
TE 298.0 K
D1 2.00000000 sec
D11 0.03000000 sec
TD0 1
SFO1 376.5021312 MHz
NUC1 19F
P0 6.00 usec
P1 18.00 usec
PLW1 18.94099998 W
SFO2 400.1318006 MHz
NUC2 1H
CPDPRG[2 waltz16
PCPD2 90.00 usec
PLW2 18.69700050 W
PLW12 0.23083000 W

F2 - Processing parameters
SI 262144
SF 376.4983662 MHz
WDW EM
SSB 0
LB 1.00 Hz
GB 0
PC 2.00

### **BGAz-003** $^{1}$H NMR Spectrum

BGAz-003

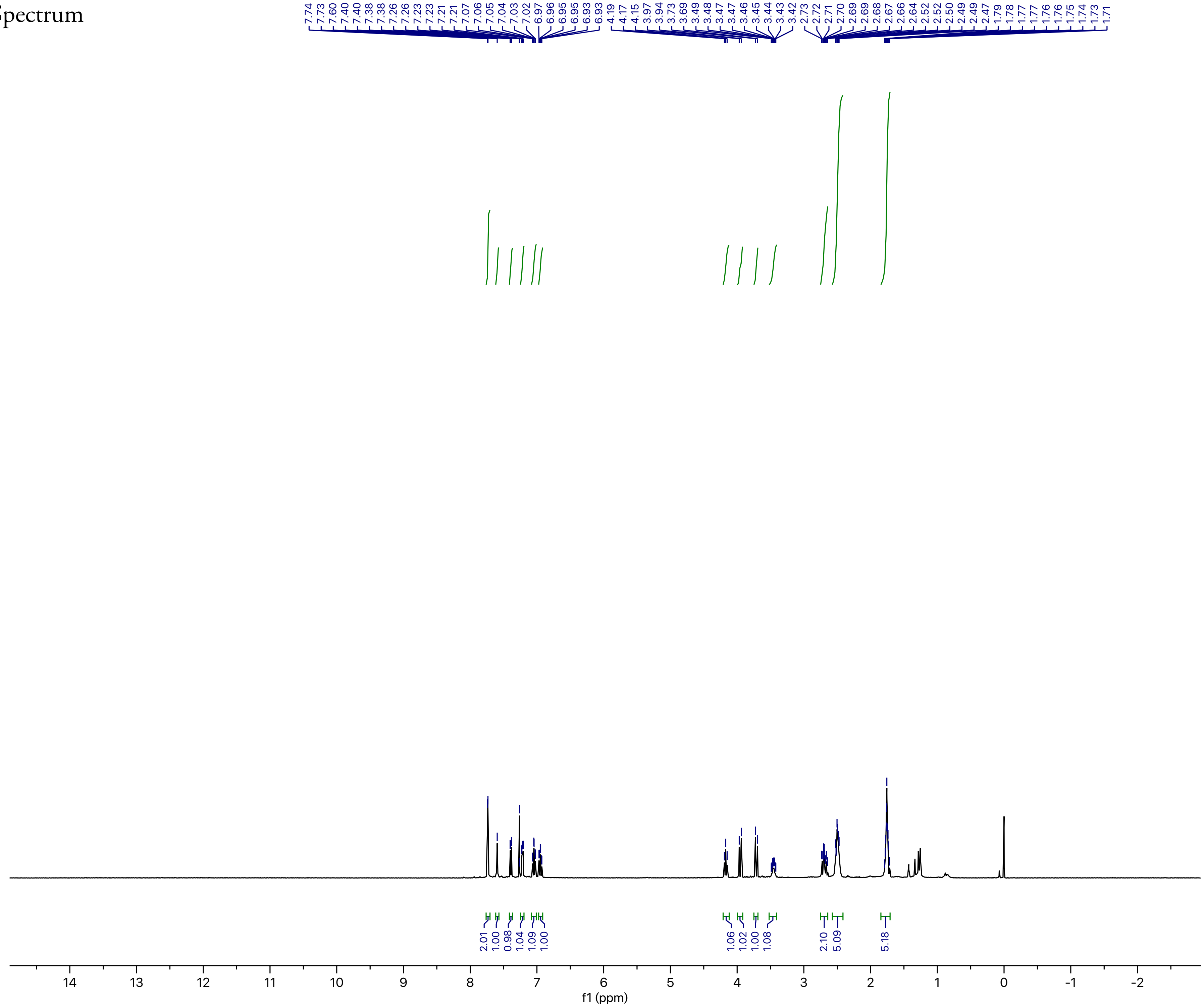

Current Data Parameters
NAME CYX91
EXPNO 10
PROCNO 1

F2 - Acquisition Parameters
Date_ 20190404
Time 1.01 h
INSTRUM AvanceNeo
PROBHD Z116098_0793 (
PULPROG zg30
TD 65536
SOLVENT CDCl3
NS 2
DS 0
SWH 7142.857 Hz
FIDRES 0.217983 Hz
AQ 4.5875201 sec
RG 92.3077
DW 70.000 usec
DE 14.62 usec
TE 298.0 K
D1 2.00000000 sec
TD0 1
SFO1 400.1324008 MHz
NUC1 1H
P0 3.33 usec
P1 10.00 usec
PLW1 18.69700050 W

F2 - Processing parameters
SI 131072
SF 400.1300093 MHz
WDW EM
SSB 0
LB 0.10 Hz
GB 0
PC 1.00

**BGAz-003** $^{1}$H NMR Spectrum (zoom 1)

BGAz-003

```
Current Data Parameters
NAME CYX91
EXPNO 10
PROCNO 1

F2 - Acquisition Parameters
Date_ 20190404
Time_ 1.01 h
INSTRUM AvanceNeo
PROBHD Z116098_0793 (
PULPROG zg30
TD 65536
SOLVENT CDCl3
NS 2
DS 0
SWH 7142.857 Hz
FIDRES 0.217983 Hz
AQ 4.5875201 sec
RG 92.3077
DW 70.000 usec
DE 14.62 usec
TE 298.0 K
D1 2.00000000 sec
TD0 1
SFO1 400.1324008 MHz
NUC1 1H
P0 3.33 usec
P1 10.00 usec
PLW1 18.69700050 W

F2 - Processing parameters
SI 131072
SF 400.1300093 MHz
WDW EM
SSB 0
LB 0.10 Hz
GB 0
PC 1.00
```

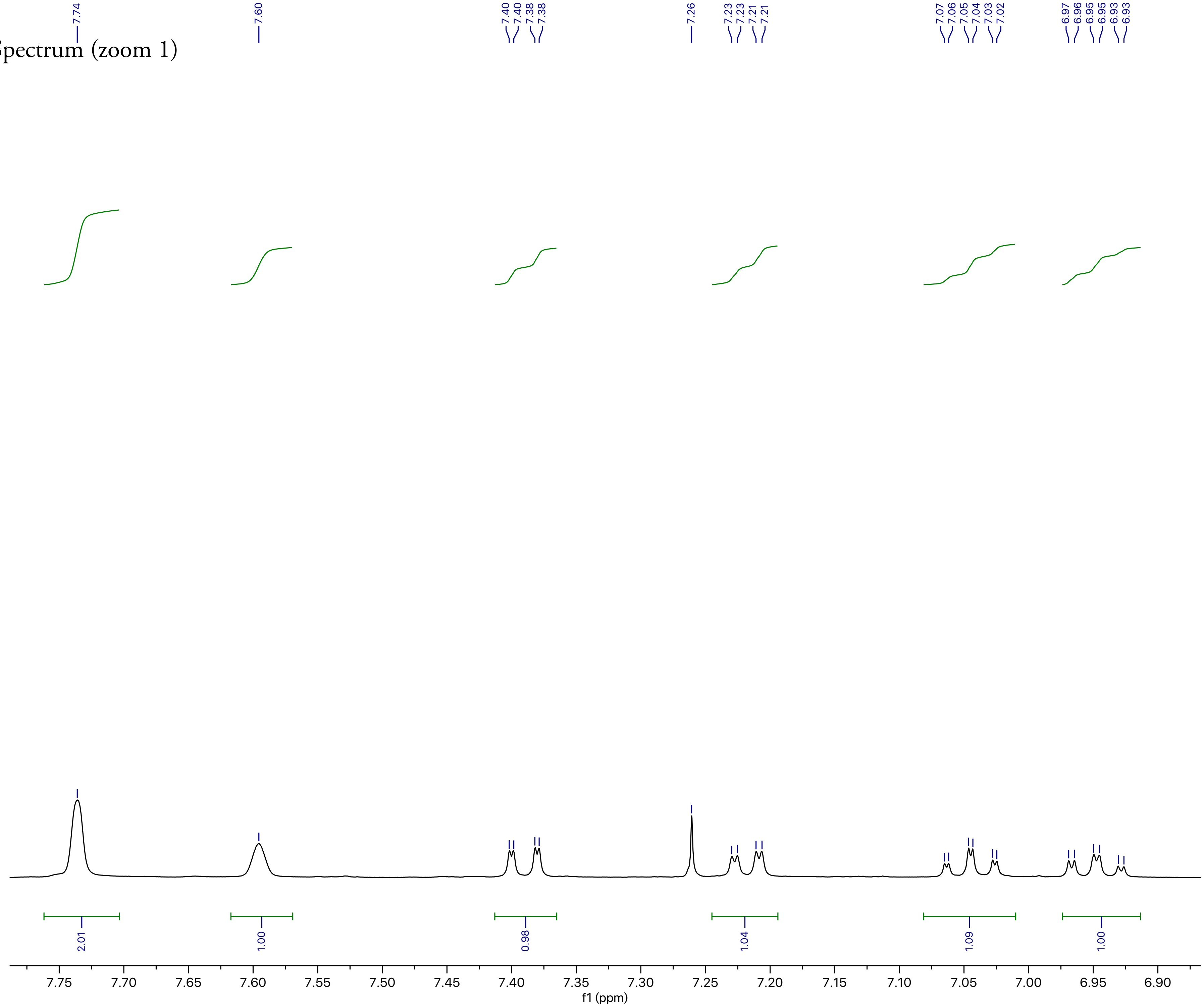

**BGAz-003** ¹H NMR Spectrum (zoom 2)

BGAz-003

Current Data Parameters
NAME CYX91
EXPNO 10
PROCNO 1

F2 - Acquisition Parameters
Date_ 20190404
Time 1.01 h
INSTRUM AvanceNeo
PROBHD Z116098_0793 (
PULPROG zg30
TD 65536
SOLVENT CDCl3
NS 2
DS 0
SWH 7142.857 Hz
FIDRES 0.217983 Hz
AQ 4.5875201 sec
RG 92.3077
DW 70.000 usec
DE 14.62 usec
TE 298.0 K
D1 2.00000000 sec
TD0 1
SFO1 400.1324008 MHz
NUC1 1H
P0 3.33 usec
P1 10.00 usec
PLW1 18.69700050 W

F2 - Processing parameters
SI 131072
SF 400.1300093 MHz
WDW EM
SSB 0
LB 0.10 Hz
GB 0
PC 1.00

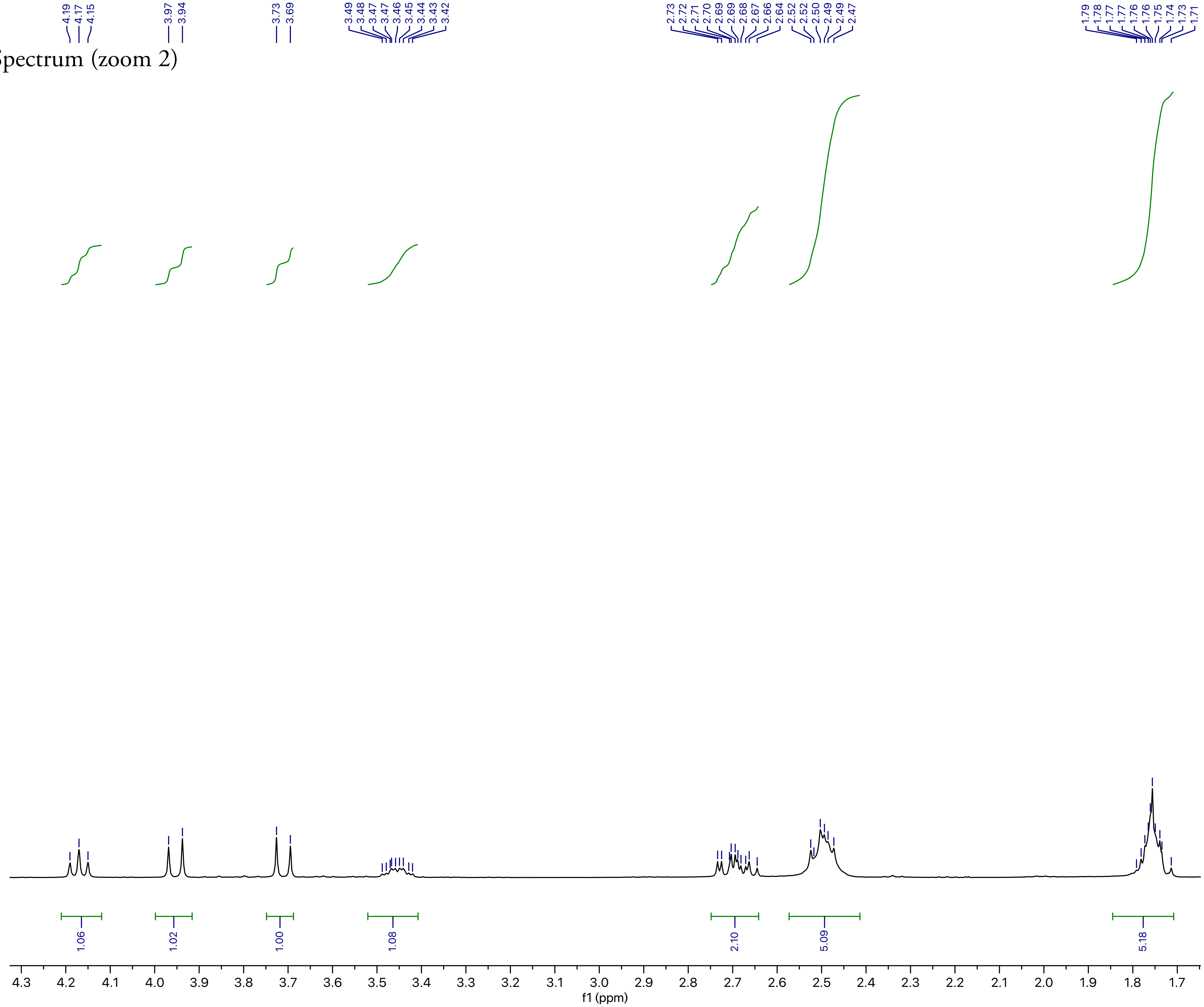

**BGAz-003** $^{13}$C NMR Spectrum

BGAz-003

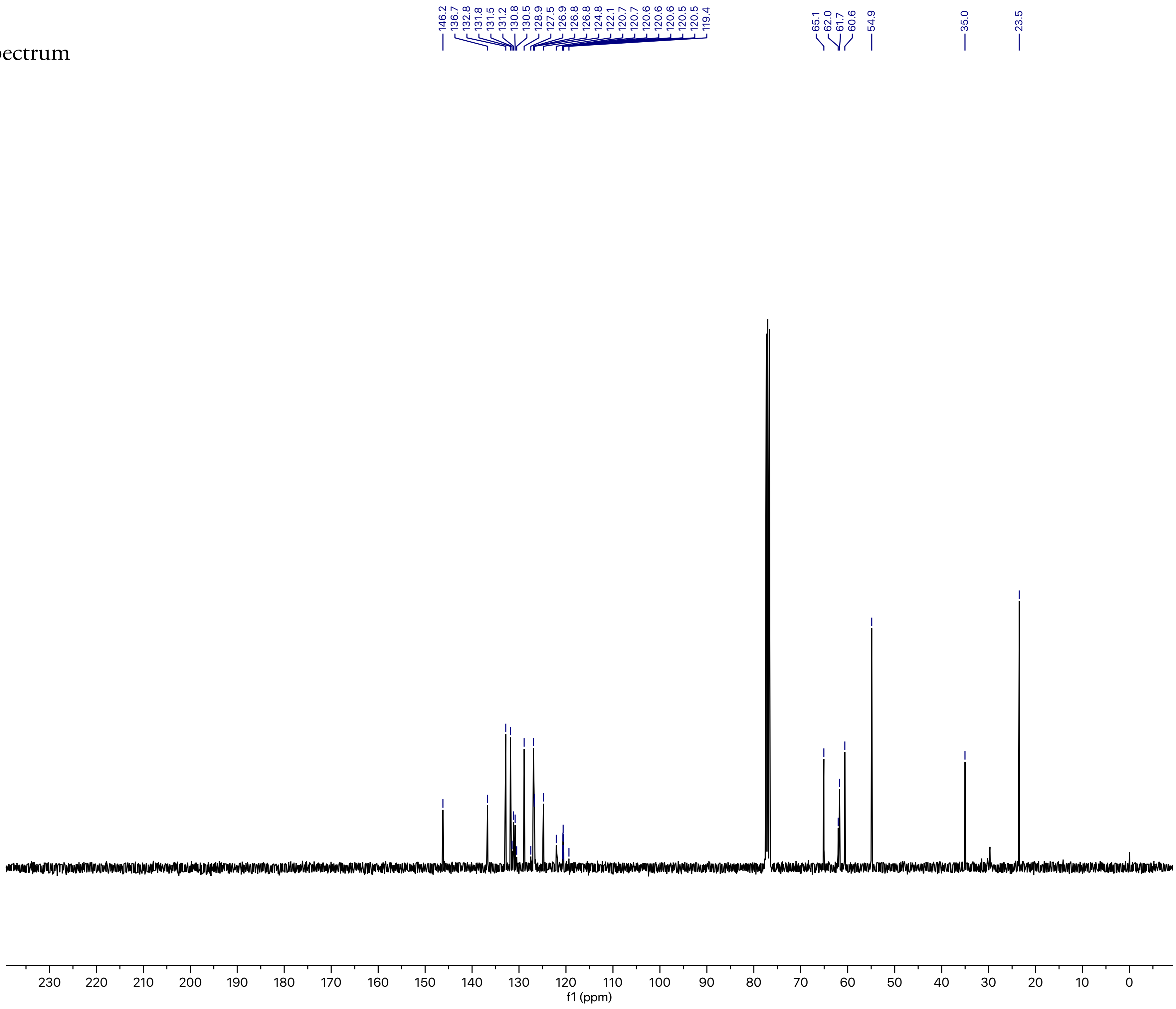

Current Data Parameters
NAME CYX91
EXPNO 12
PROCNO 1

F2 - Acquisition Parameters
Date_ 20190404
Time 1.33 h
INSTRUM AvanceNeo
PROBHD Z116098_0793 (
PULPROG zgpg30
TD 119044
SOLVENT CDCl3
NS 512
DS 0
SWH 25000.000 Hz
FIDRES 0.420013 Hz
AQ 2.3808801 sec
RG 31.9602
DW 20.000 usec
DE 7.12 usec
TE 298.0 K
D1 1.00000000 sec
D11 0.03000000 sec
TD0 1
SFO1 100.6243390 MHz
NUC1 13C
P0 3.33 usec
P1 10.00 usec
PLW1 83.92700195 W
SFO2 400.1318006 MHz
NUC2 1H
CPDPRG[2 waltz64
PCPD2 90.00 usec
PLW2 18.69700050 W
PLW12 0.23083000 W
PLW13 0.11611000 W

F2 - Processing parameters
SI 131072
SF 100.6127685 MHz
WDW EM
SSB 0
LB 1.00 Hz
GB 0
PC 1.40

**BGAz-003** $^{19}$F NMR Spectrum

BGAz-003

Current Data Parameters
NAME CYX91
EXPNO 11
PROCNO 1

F2 - Acquisition Parameters
Date_ 20190404
Time 1.02 h
INSTRUM AvanceNeo
PROBHD Z116098_0793 (
PULPROG zgig30
TD 261948
SOLVENT CDCl3
NS 16
DS 0
SWH 156250.000 Hz
FIDRES 1.192985 Hz
AQ 0.8382336 sec
RG 101
DW 3.200 usec
DE 6.82 usec
TE 298.0 K
D1 2.00000000 sec
D11 0.03000000 sec
TD0 1
SFO1 376.5021312 MHz
NUC1 19F
P0 6.00 usec
P1 18.00 usec
PLW1 18.94099998 W
SFO2 400.1318006 MHz
NUC2 1H
CPDPRG[2 waltz16
PCPD2 90.00 usec
PLW2 18.69700050 W
PLW12 0.23083000 W

F2 - Processing parameters
SI 262144
SF 376.4983662 MHz
WDW EM
SSB 0
LB 1.00 Hz
GB 0
PC 2.00

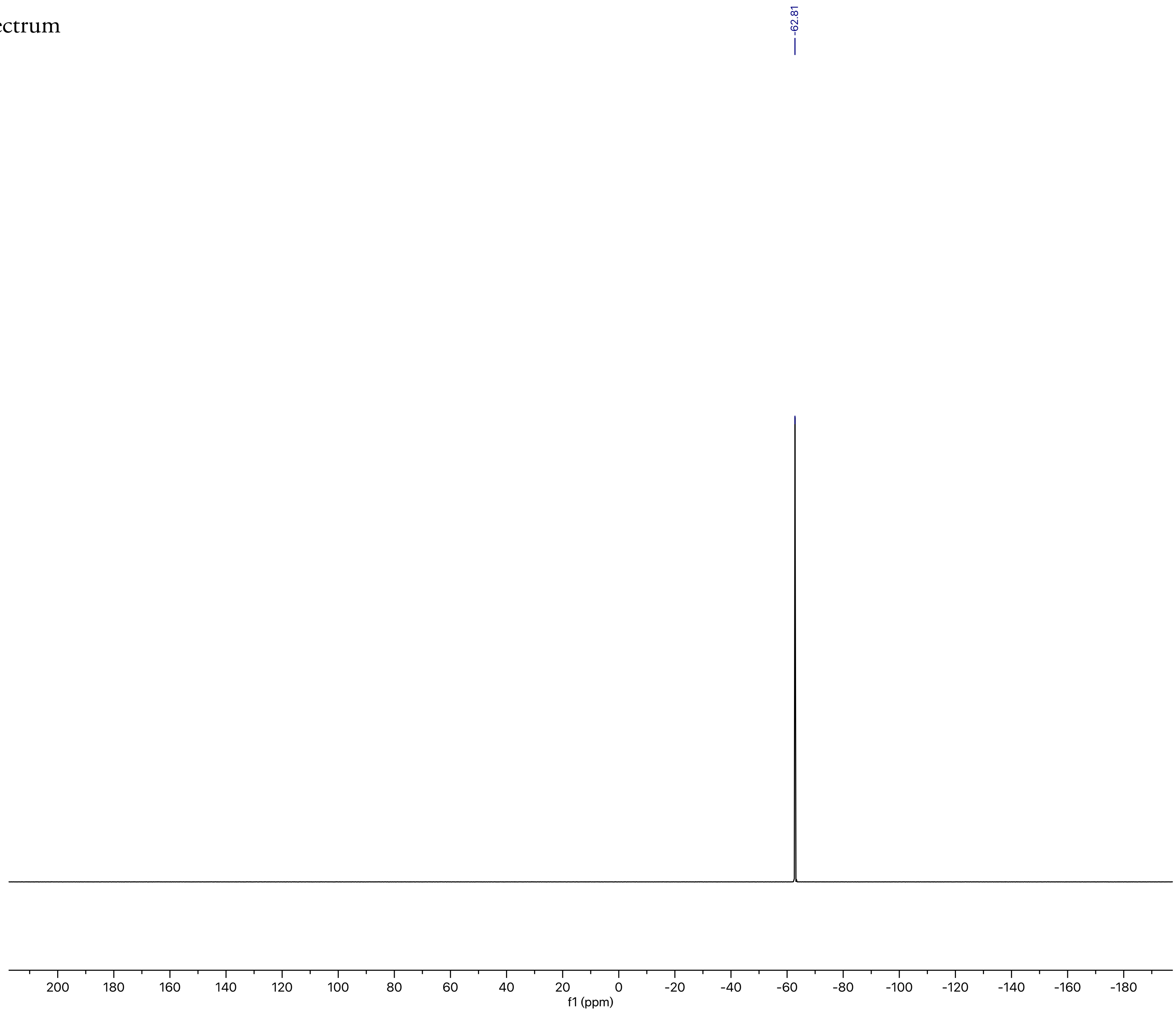

**BGAz-004** $^{1}$H NMR Spectrum

BGAz-004

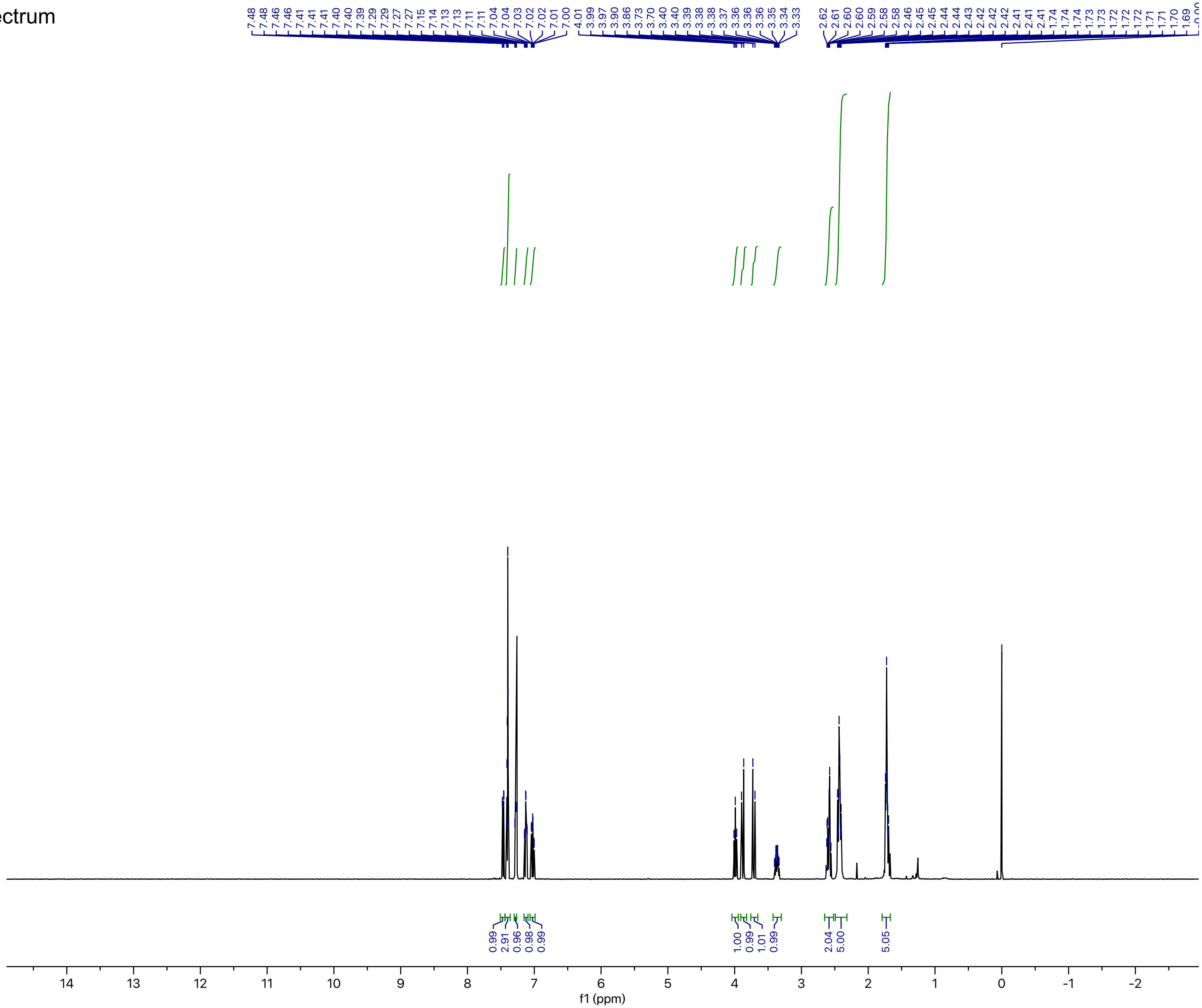

```
Current Data Parameters
NAME CYX92
EXPNO 10
PROCNO 1

F2 - Acquisition Parameters
Date_ 20190404
Time 1.40 h
INSTRUM AvanceNeo
PROBHD Z116098_0793 (
PULPROG zg30
TD 65536
SOLVENT CDCl3
NS 2
DS 0
SWH 7142.857 Hz
FIDRES 0.217983 Hz
AQ 4.5875201 sec
RG 101
DW 70.000 usec
DE 14.62 usec
TE 298.0 K
D1 2.00000000 sec
TD0 1
SFO1 400.1324008 MHz
NUC1 1H
P0 3.33 usec
P1 10.00 usec
PLW1 18.69700050 W

F2 - Processing parameters
SI 131072
SF 400.1300098 MHz
WDW EM
SSB 0
LB 0.10 Hz
GB 0
PC 1.00
```

**BGAz-004** $^{1}$H NMR Spectrum (zoom 1)

BGAz-004

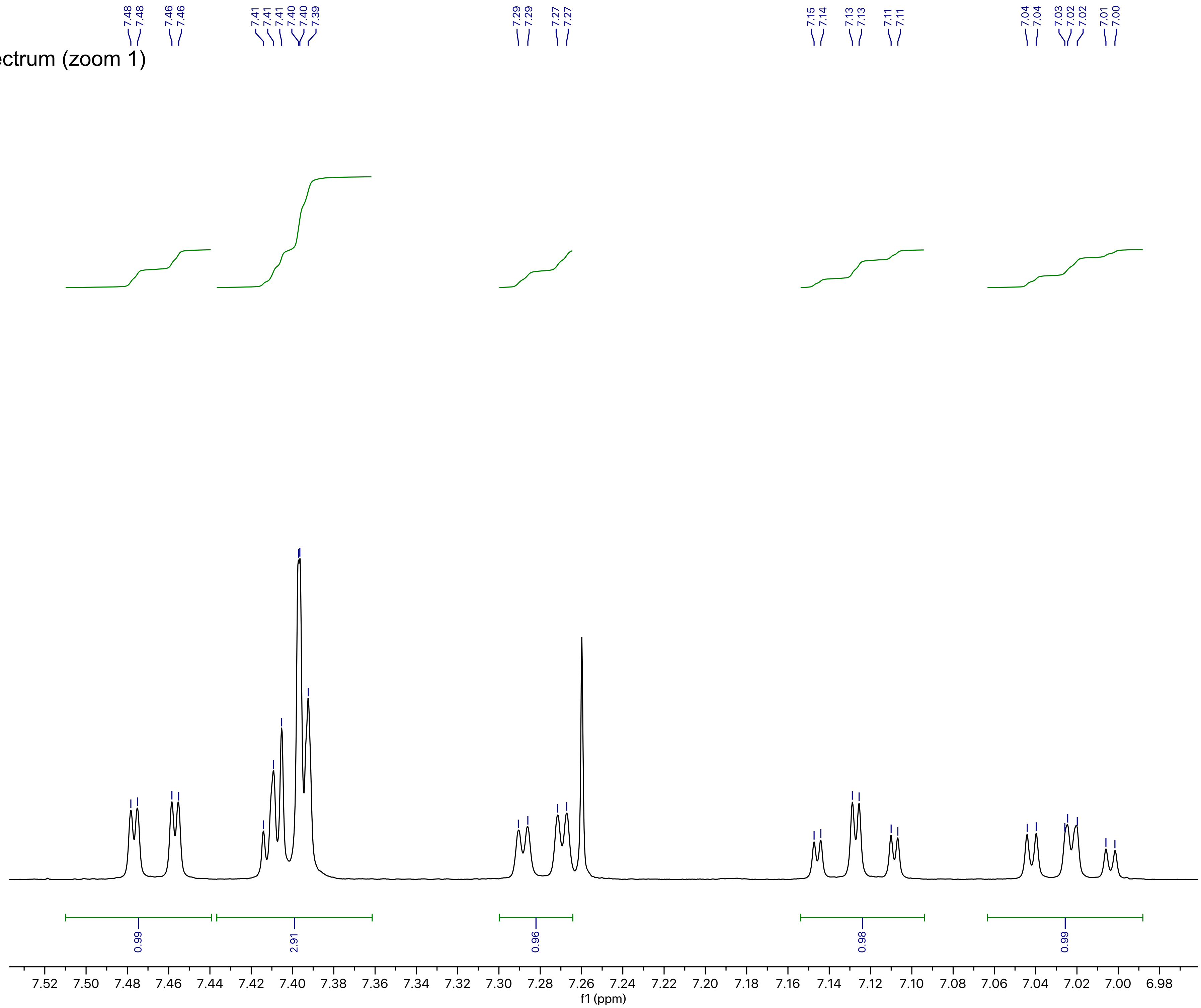

Current Data Parameters
NAME CYX92
EXPNO 10
PROCNO 1

F2 - Acquisition Parameters
Date_ 20190404
Time 1.40 h
INSTRUM AvanceNeo
PROBHD Z116098_0793 (
PULPROG zg30
TD 65536
SOLVENT CDCl3
NS 2
DS 0
SWH 7142.857 Hz
FIDRES 0.217983 Hz
AQ 4.5875201 sec
RG 101
DW 70.000 usec
DE 14.62 usec
TE 298.0 K
D1 2.00000000 sec
TD0 1
SFO1 400.1324008 MHz
NUC1 1H
P0 3.33 usec
P1 10.00 usec
PLW1 18.69700050 W

F2 - Processing parameters
SI 131072
SF 400.1300098 MHz
WDW EM
SSB 0
LB 0.10 Hz
GB 0
PC 1.00

**BGAz-004** [1]H NMR Spectrum (zoom 2)

BGAz-004

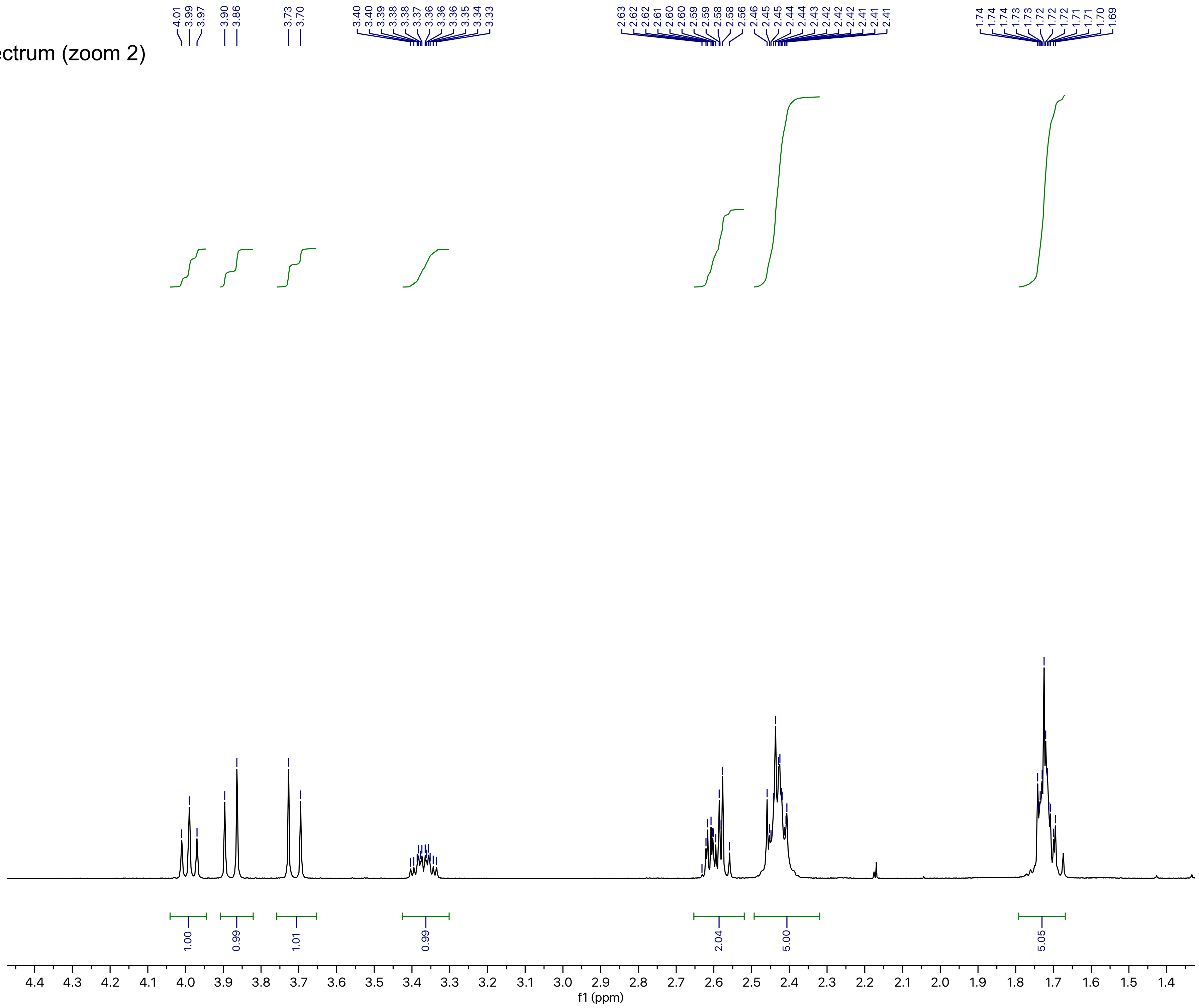

```
Current Data Parameters
NAME CYX92
EXPNO 10
PROCNO 1

F2 - Acquisition Parameters
Date_ 20190404
Time 1.40 h
INSTRUM AvanceNeo
PROBHD Z116098_0793 (
PULPROG zg30
TD 65536
SOLVENT CDCl3
NS 2
DS 0
SWH 7142.857 Hz
FIDRES 0.217983 Hz
AQ 4.5875201 sec
RG 101
DW 70.000 usec
DE 14.62 usec
TE 298.0 K
D1 2.00000000 sec
TD0 1
SFO1 400.1324008 MHz
NUC1 1H
P0 3.33 usec
P1 10.00 usec
PLW1 18.69700050 W

F2 - Processing parameters
SI 131072
SF 400.1300098 MHz
WDW EM
SSB 0
LB 0.10 Hz
GB 0
PC 1.00
```

**BGAz-004** $^{13}$C NMR Spectrum

BGAz-004

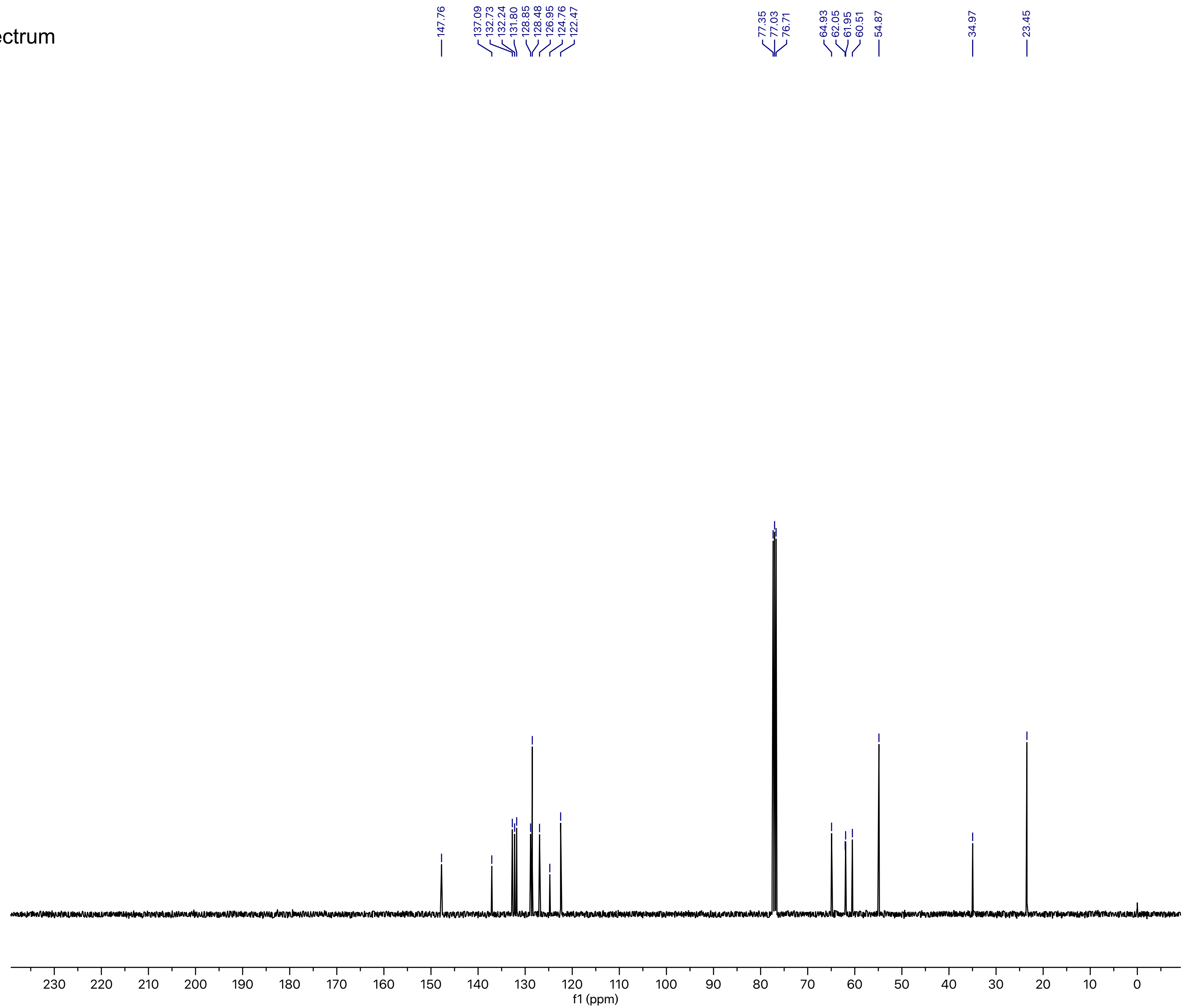

Current Data Parameters
NAME CYX92
EXPNO 11
PROCNO 1

F2 - Acquisition Parameters
Date_ 20190404
Time 2.11 h
INSTRUM AvanceNeo
PROBHD Z116098_0793 (
PULPROG zgpg30
TD 119044
SOLVENT CDCl3
NS 512
DS 0
SWH 25000.000 Hz
FIDRES 0.420013 Hz
AQ 2.3808801 sec
RG 31.9602
DW 20.000 usec
DE 7.12 usec
TE 298.0 K
D1 1.00000000 sec
D11 0.03000000 sec
TD0 1
SFO1 100.6243390 MHz
NUC1 13C
P0 3.33 usec
P1 10.00 usec
PLW1 83.92700195 W
SFO2 400.1318006 MHz
NUC2 1H
CPDPRG[2 waltz64
PCPD2 90.00 usec
PLW2 18.69700050 W
PLW12 0.23083000 W
PLW13 0.11611000 W

F2 - Processing parameters
SI 131072
SF 100.6127685 MHz
WDW EM
SSB 0
LB 1.00 Hz
GB 0
PC 1.40

**BGAz-005** $^{1}$H NMR Spectrum

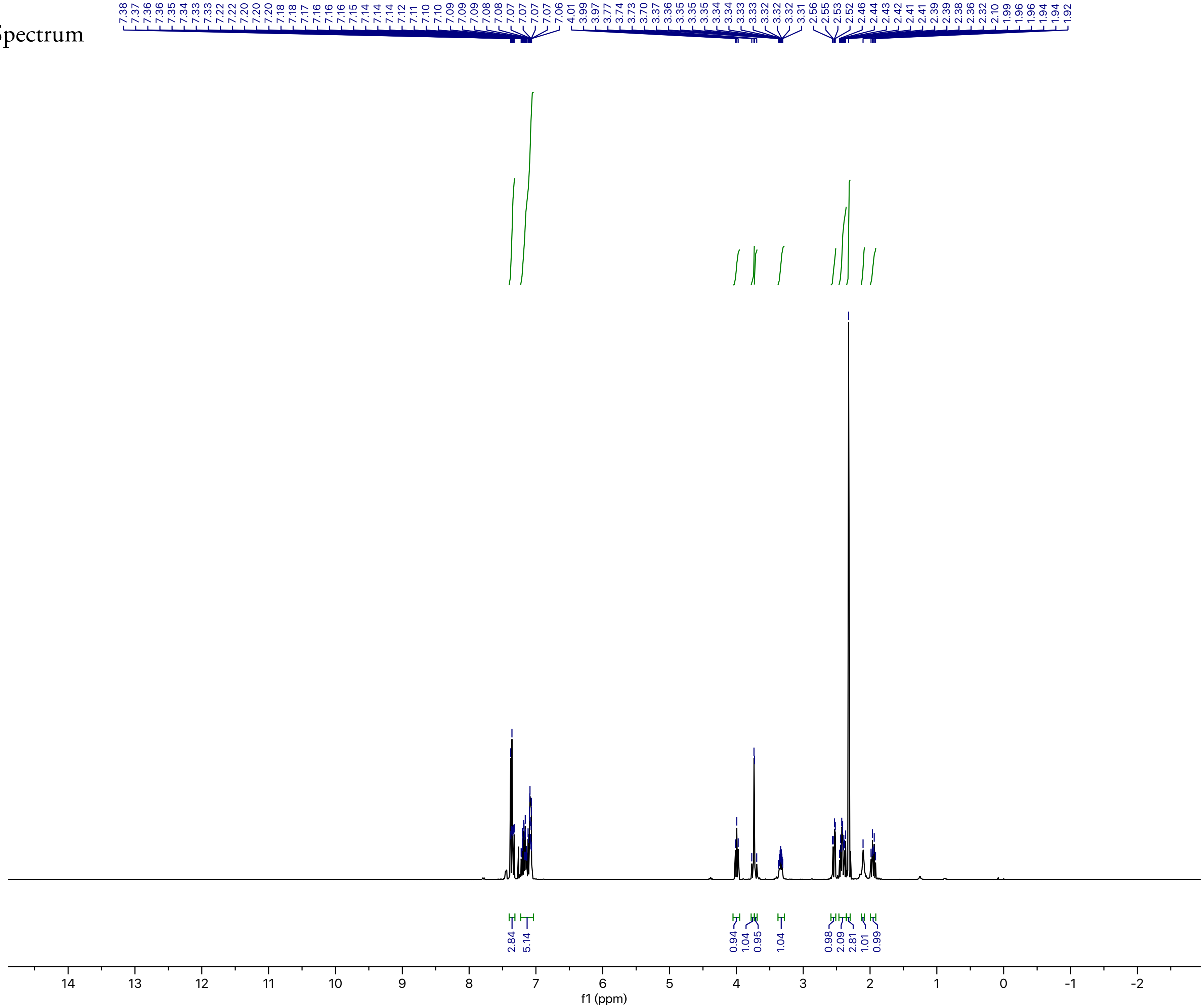

Current Data Parameters
NAME YC-GIBH-122
EXPNO 10
PROCNO 1

F2 - Acquisition Parameters
Date_ 20190615
Time 1.55 h
INSTRUM AvanceNeo
PROBHD Z116098_0793 (
PULPROG zg30
TD 65536
SOLVENT CDCl3
NS 2
DS 0
SWH 7142.857 Hz
FIDRES 0.217983 Hz
AQ 4.5875201 sec
RG 32
DW 70.000 usec
DE 14.62 usec
TE 298.0 K
D1 2.00000000 sec
TD0 1
SFO1 400.1324008 MHz
NUC1 1H
P0 3.33 usec
P1 10.00 usec
PLW1 18.69700050 W

F2 - Processing parameters
SI 131072
SF 400.1300419 MHz
WDW EM
SSB 0
LB 0.10 Hz
GB 0
PC 1.00

**BGAz-005** $^{1}$H NMR Spectrum (zoom 1)

BGAz-005

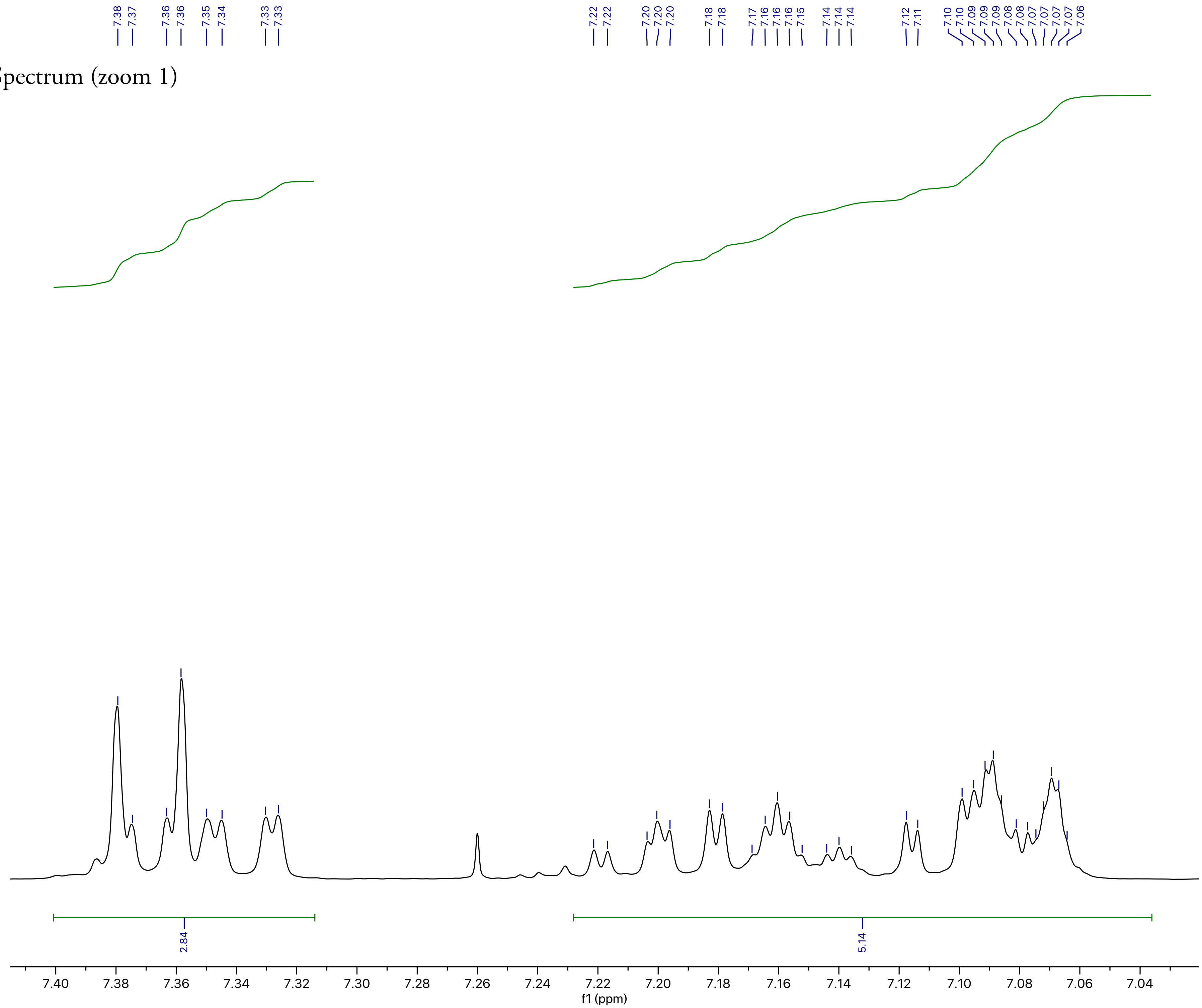

```
Current Data Parameters
NAME YC-GIBH-122
EXPNO 10
PROCNO 1

F2 - Acquisition Parameters
Date_ 20190615
Time 1.55 h
INSTRUM AvanceNeo
PROBHD Z116098_0793 (
PULPROG zg30
TD 65536
SOLVENT CDCl3
NS 2
DS 0
SWH 7142.857 Hz
FIDRES 0.217983 Hz
AQ 4.5875201 sec
RG 32
DW 70.000 usec
DE 14.62 usec
TE 298.0 K
D1 2.00000000 sec
TD0 1
SFO1 400.1324008 MHz
NUC1 1H
P0 3.33 usec
P1 10.00 usec
PLW1 18.69700050 W

F2 - Processing parameters
SI 131072
SF 400.1300419 MHz
WDW EM
SSB 0
LB 0.10 Hz
GB 0
PC 1.00
```

**BGAz-005** [1]H NMR Spectrum (zoom 2)

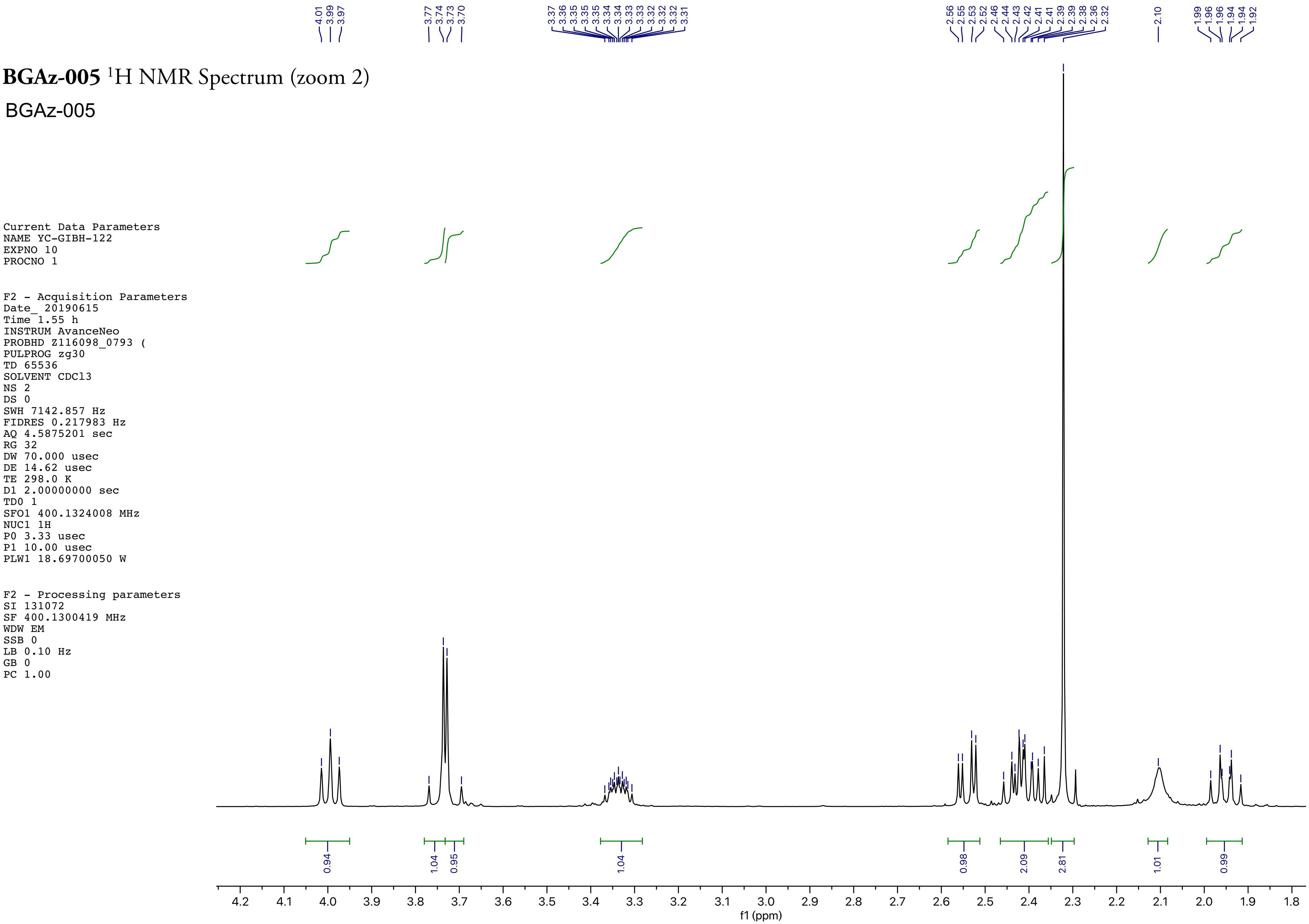

Current Data Parameters
NAME YC-GIBH-122
EXPNO 10
PROCNO 1

F2 - Acquisition Parameters
Date_ 20190615
Time 1.55 h
INSTRUM AvanceNeo
PROBHD Z116098_0793 (
PULPROG zg30
TD 65536
SOLVENT CDCl3
NS 2
DS 0
SWH 7142.857 Hz
FIDRES 0.217983 Hz
AQ 4.5875201 sec
RG 32
DW 70.000 usec
DE 14.62 usec
TE 298.0 K
D1 2.00000000 sec
TD0 1
SFO1 400.1324008 MHz
NUC1 1H
P0 3.33 usec
P1 10.00 usec
PLW1 18.69700050 W

F2 - Processing parameters
SI 131072
SF 400.1300419 MHz
WDW EM
SSB 0
LB 0.10 Hz
GB 0
PC 1.00

**BGAz-005** $^{13}$C NMR Spectrum

BGAz-005

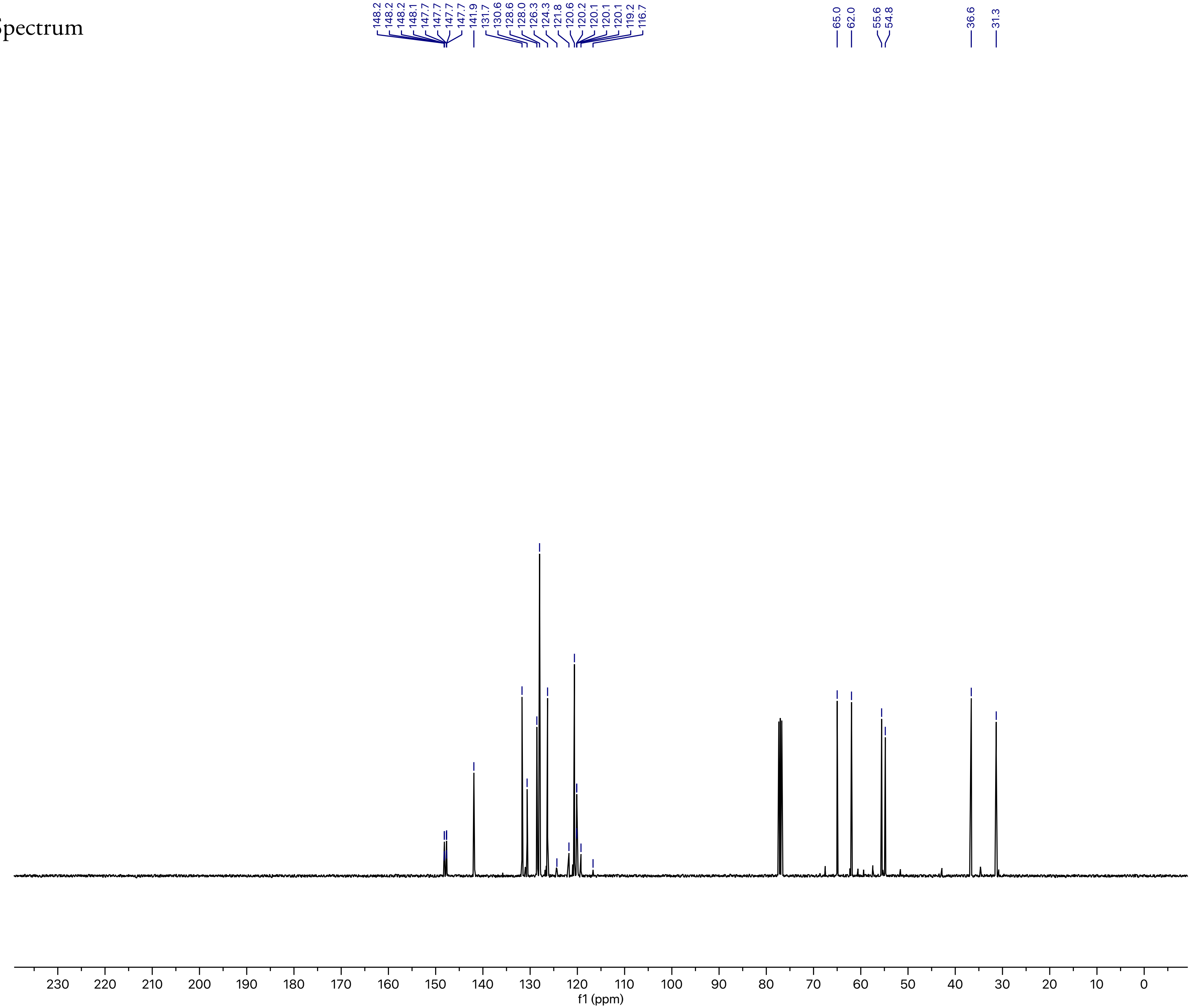

Current Data Parameters
NAME YC-GIBH-122
EXPNO 11
PROCNO 1

F2 - Acquisition Parameters
Date_ 20190615
Time 2.25 h
INSTRUM AvanceNeo
PROBHD Z116098_0793 (
PULPROG zgpg30
TD 119044
SOLVENT CDCl3
NS 512
DS 0
SWH 25000.000 Hz
FIDRES 0.420013 Hz
AQ 2.3808801 sec
RG 31.9602
DW 20.000 usec
DE 7.12 usec
TE 298.0 K
D1 1.00000000 sec
D11 0.03000000 sec
TD0 1
SFO1 100.6243390 MHz
NUC1 13C
P0 3.33 usec
P1 10.00 usec
PLW1 83.92700195 W
SFO2 400.1318006 MHz
NUC2 1H
CPDPRG[2 waltz64
PCPD2 90.00 usec
PLW2 18.69700050 W
PLW12 0.23083000 W
PLW13 0.11611000 W

F2 - Processing parameters
SI 131072
SF 100.6127685 MHz
WDW EM
SSB 0
LB 1.00 Hz
GB 0
PC 1.40

**BGAz-005** $^{19}$F NMR Spectrum

BGAz-005

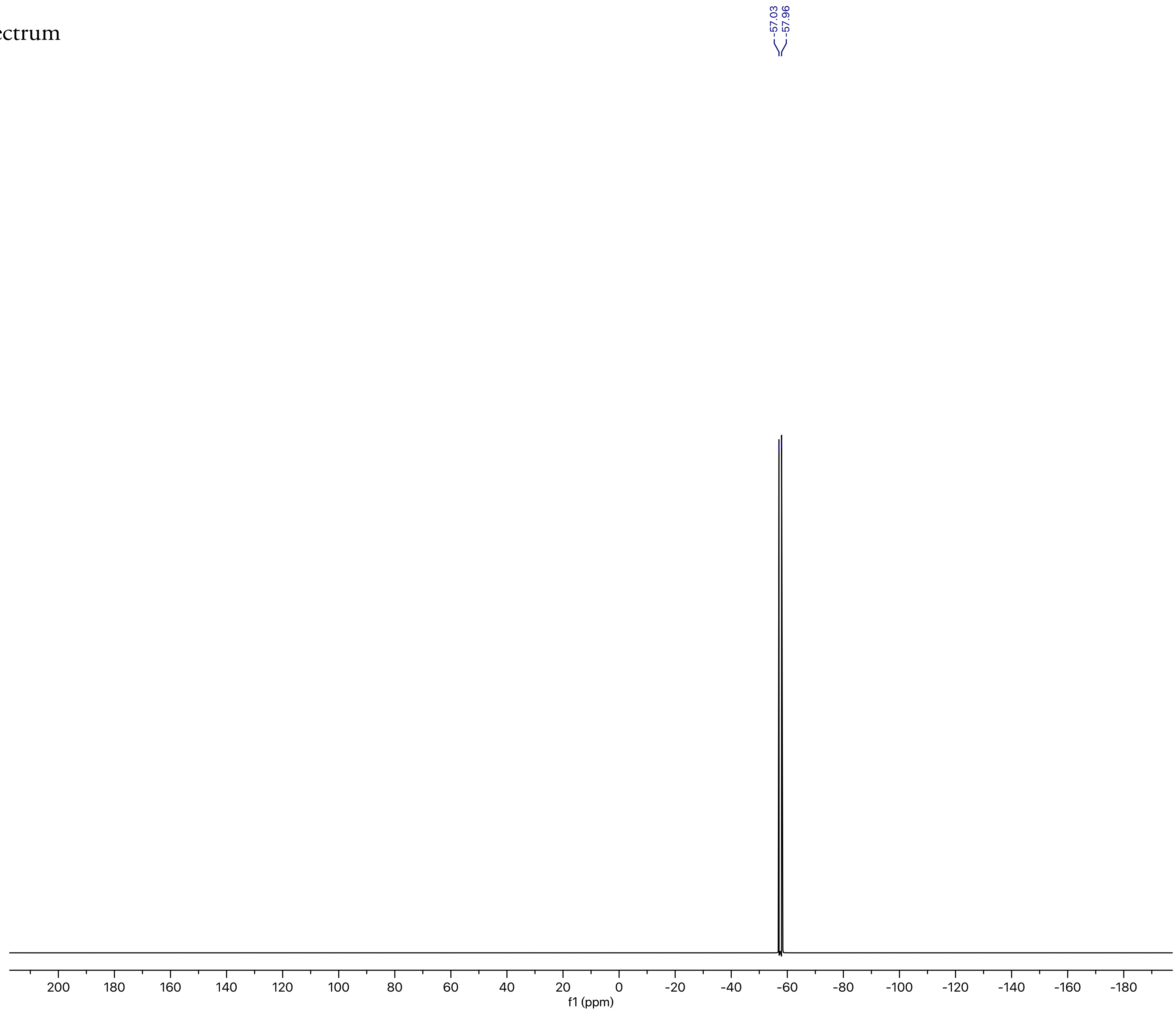

Current Data Parameters
NAME YC-GIBH-122
EXPNO 14
PROCNO 1

F2 - Acquisition Parameters
Date_ 20190615
Time 3.08 h
INSTRUM AvanceNeo
PROBHD Z116098_0793 (
PULPROG zgig30
TD 261948
SOLVENT CDCl3
NS 16
DS 0
SWH 156250.000 Hz
FIDRES 1.192985 Hz
AQ 0.8382336 sec
RG 101
DW 3.200 usec
DE 6.82 usec
TE 298.0 K
D1 2.00000000 sec
D11 0.03000000 sec
TD0 1
SFO1 376.5021312 MHz
NUC1 19F
P0 6.00 usec
P1 18.00 usec
PLW1 18.94099998 W
SFO2 400.1318006 MHz
NUC2 1H
CPDPRG[2 waltz16
PCPD2 90.00 usec
PLW2 18.69700050 W
PLW12 0.23083000 W

F2 - Processing parameters
SI 262144
SF 376.4983662 MHz
WDW EM
SSB 0
LB 1.00 Hz
GB 0
PC 2.00

**S4aa** MS

+Q1: 0.506 to 0.802 min from Sample 1472 (PXD-02370060) of DDP.wiff (Turbo Spray), subtracted (0.025 to 0.407 min) Max. 9.4e6 cps.

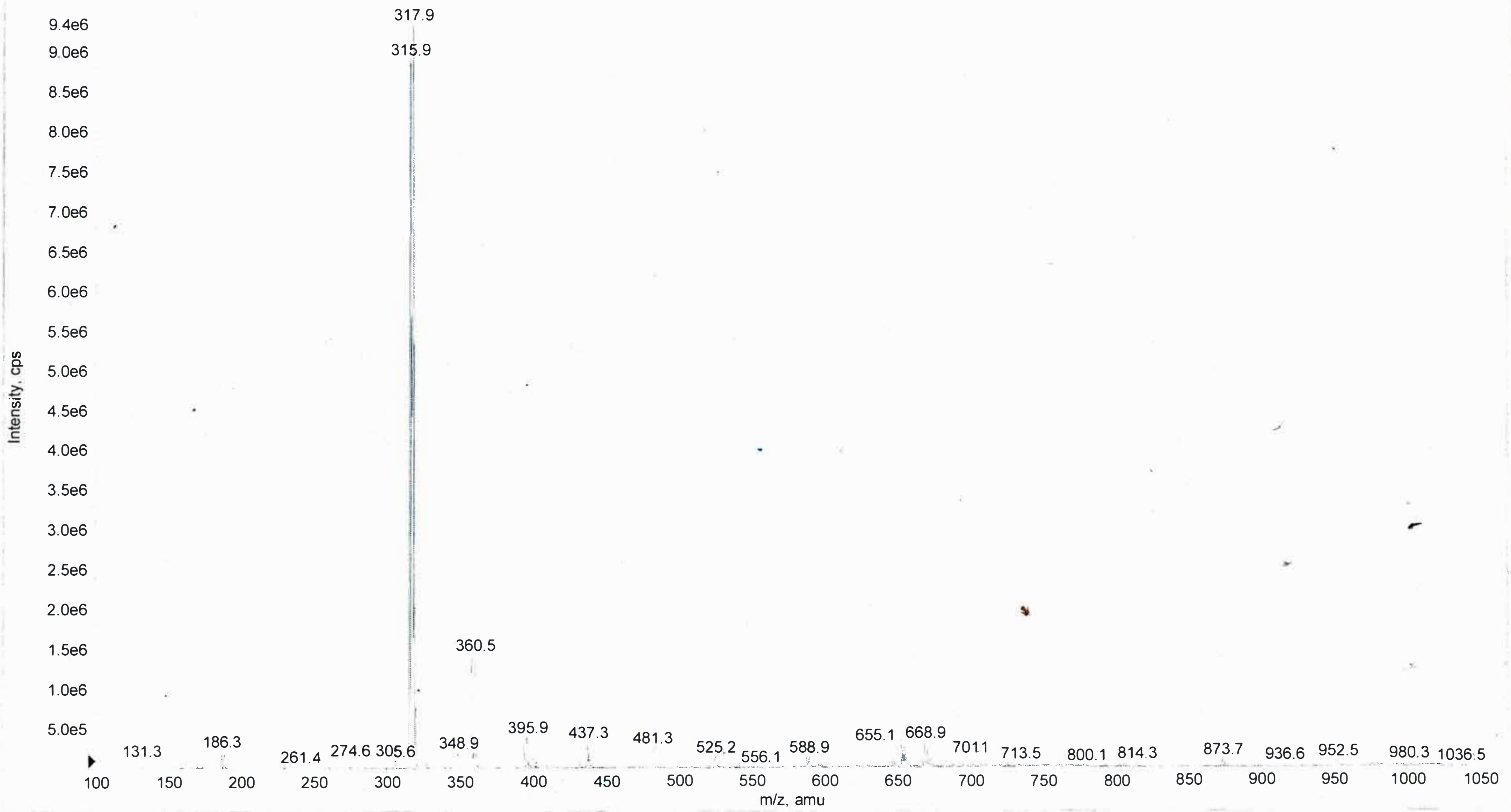

**S4bb** MS

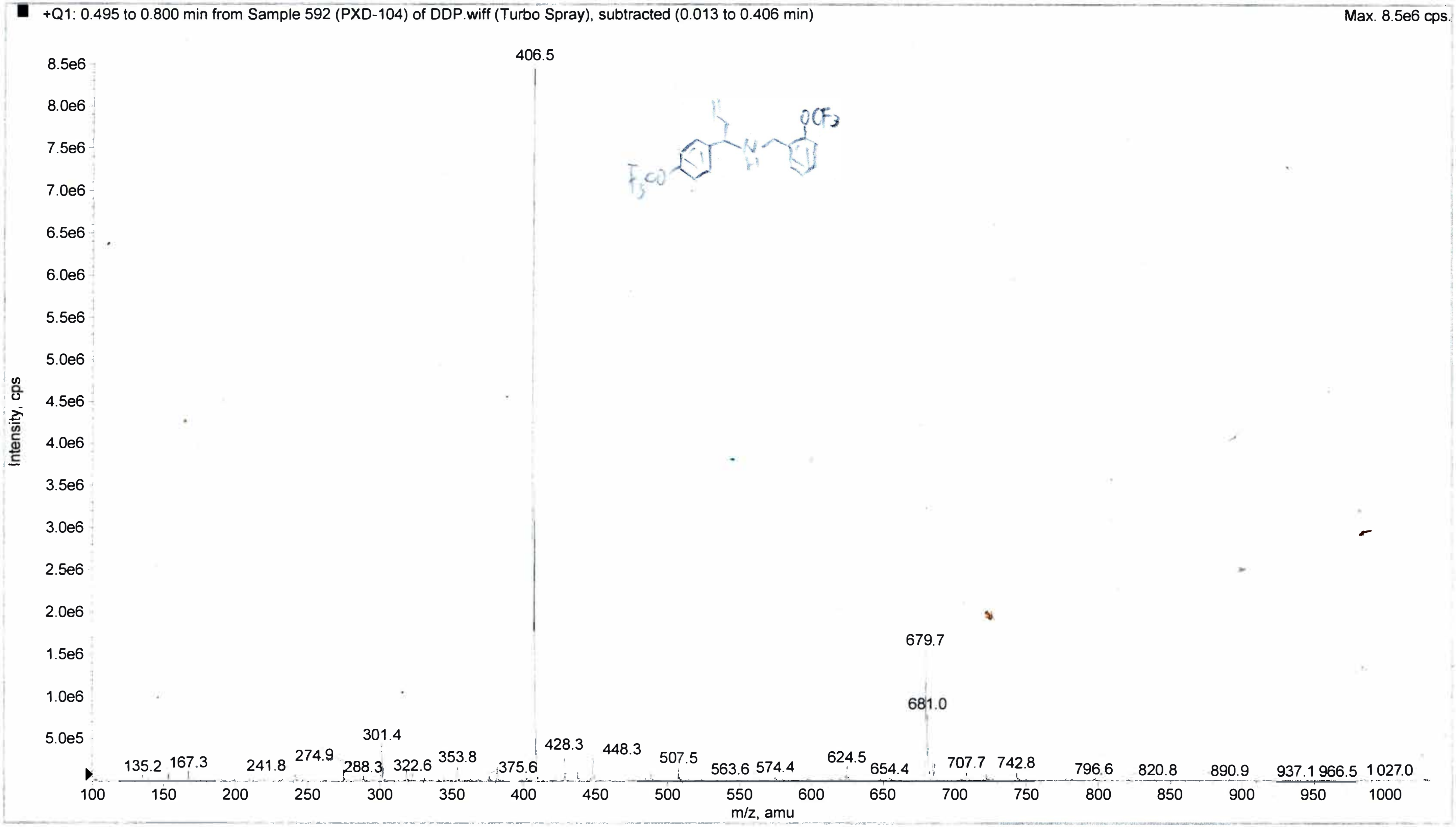
+Q1: 0.495 to 0.800 min from Sample 592 (PXD-104) of DDP.wiff (Turbo Spray), subtracted (0.013 to 0.406 min)
Max. 8.5e6 cps.
406.5
679.7
681.0
301.4
428.3
448.3
507.5
624.5
135.2
167.3
241.8
274.9
288.3
322.6
353.8
375.6
563.6
574.4
654.4
707.7
742.8
796.6
820.8
890.9
937.1
966.5
1027.0
Intensity, cps
m/z, amu

**S4ca** Mass spec (i)

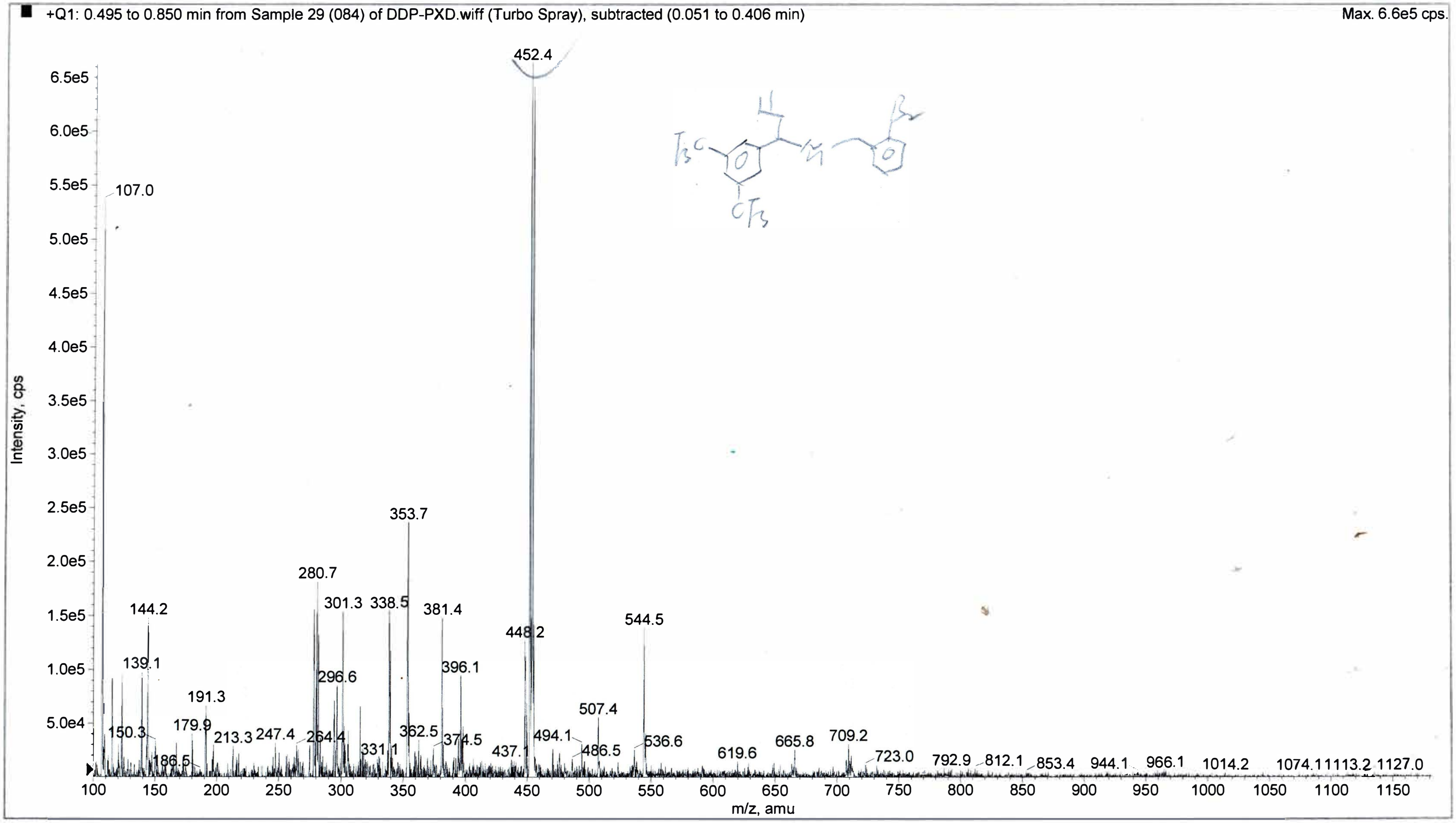

**S4ca** Mass spec (ii)

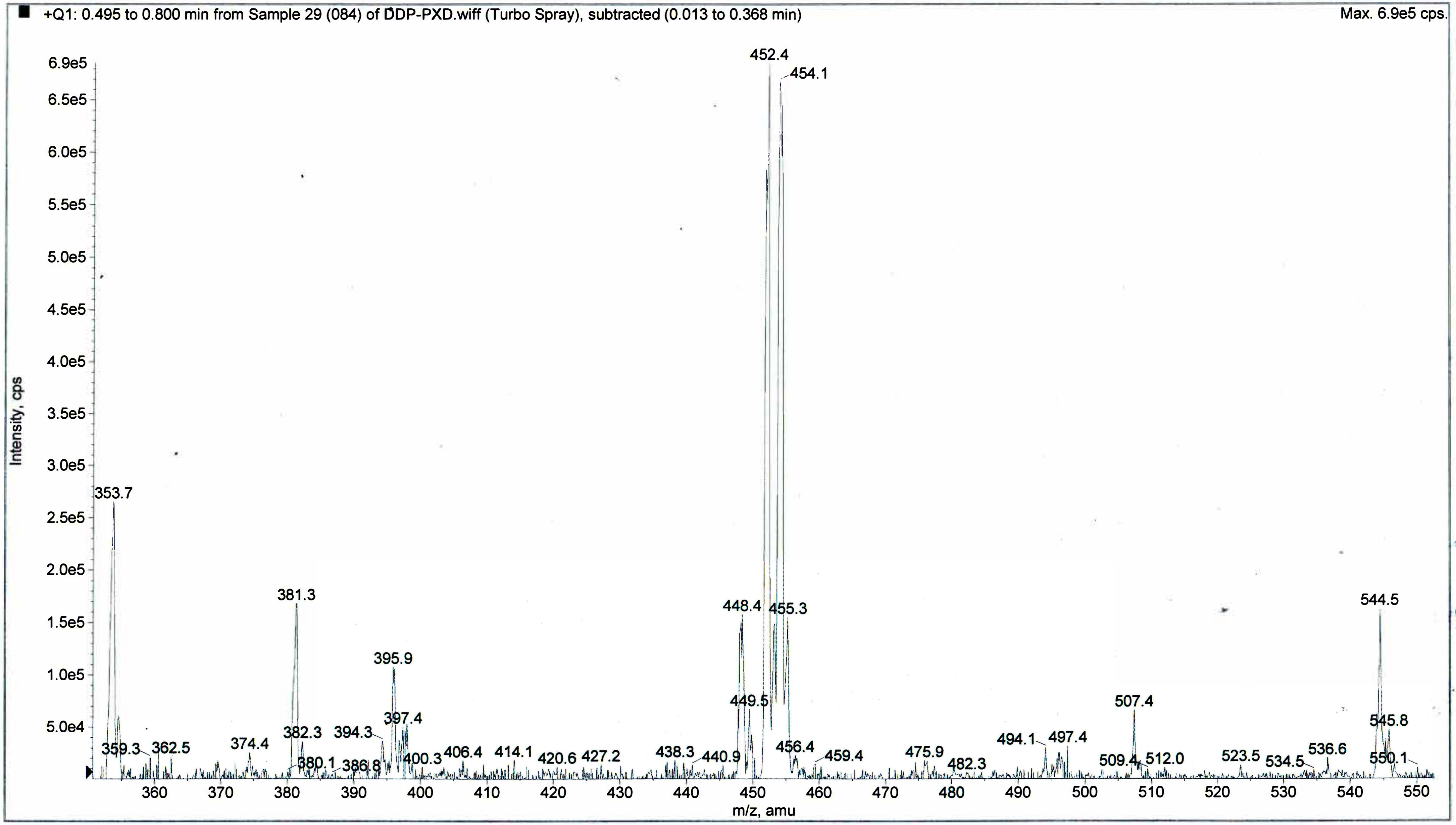

**S4da** Mass spec (i)

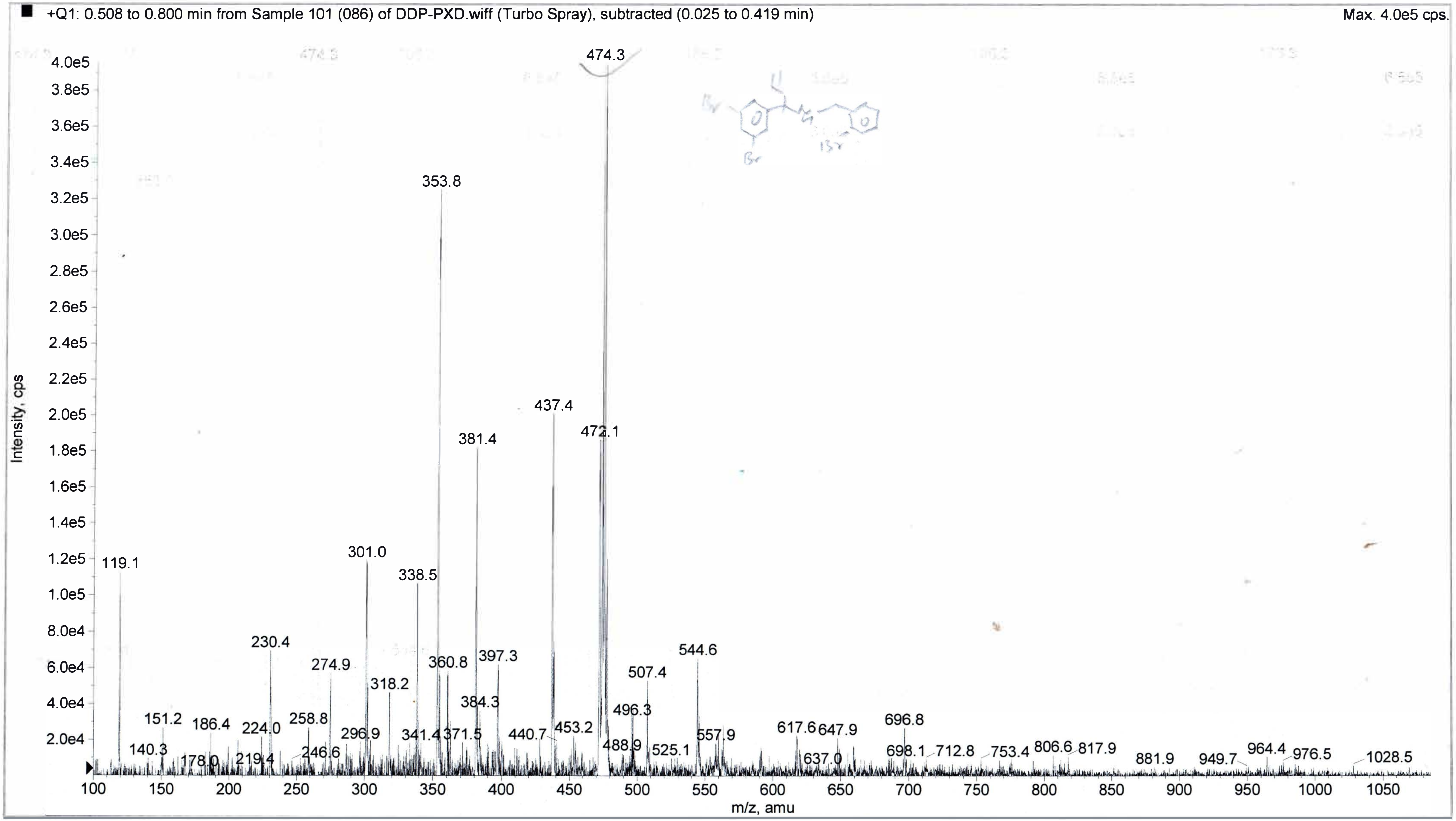

**S4da** M_qq qnca &gg'

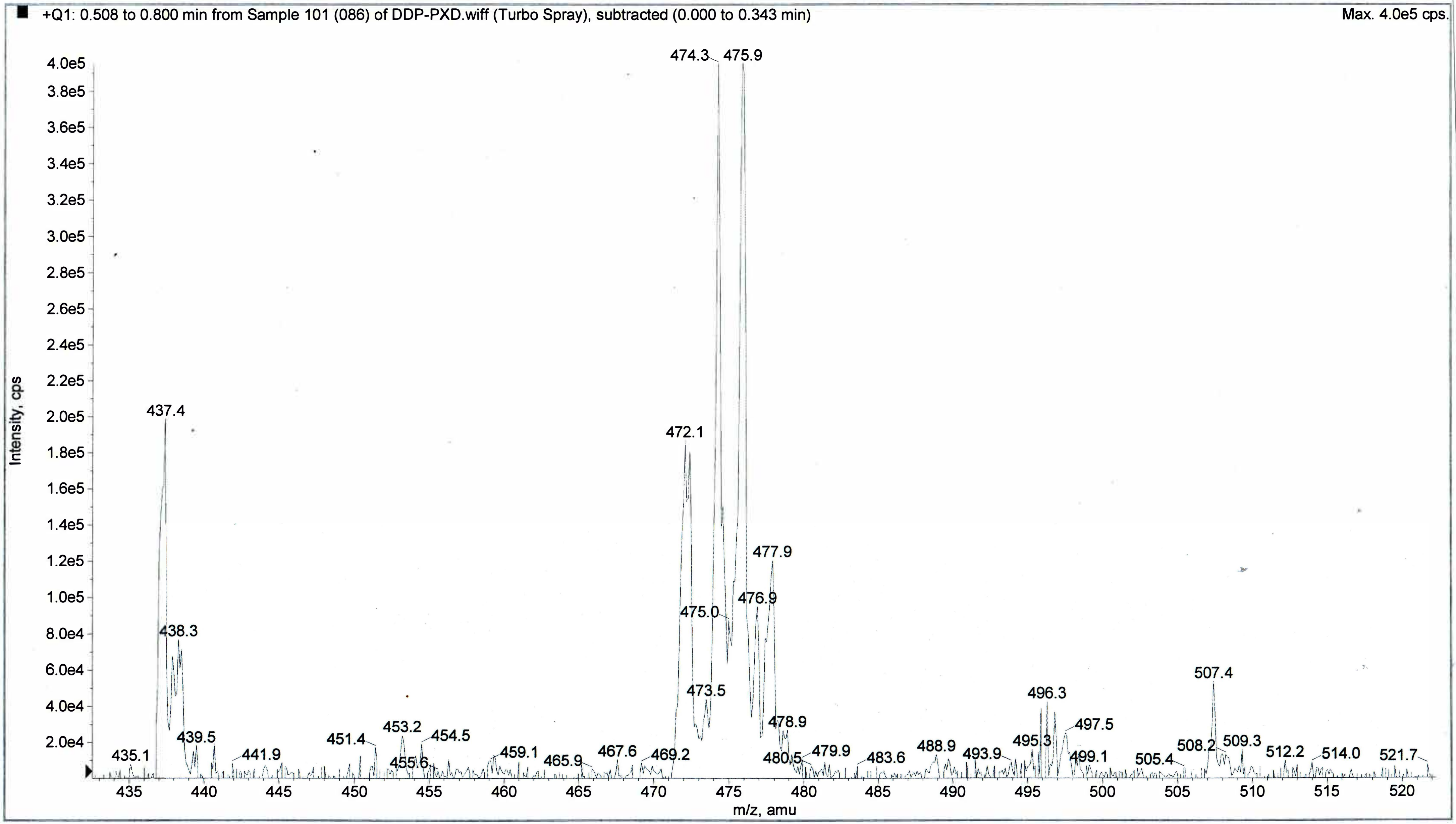

**BGAz-001** Mass spec (i)

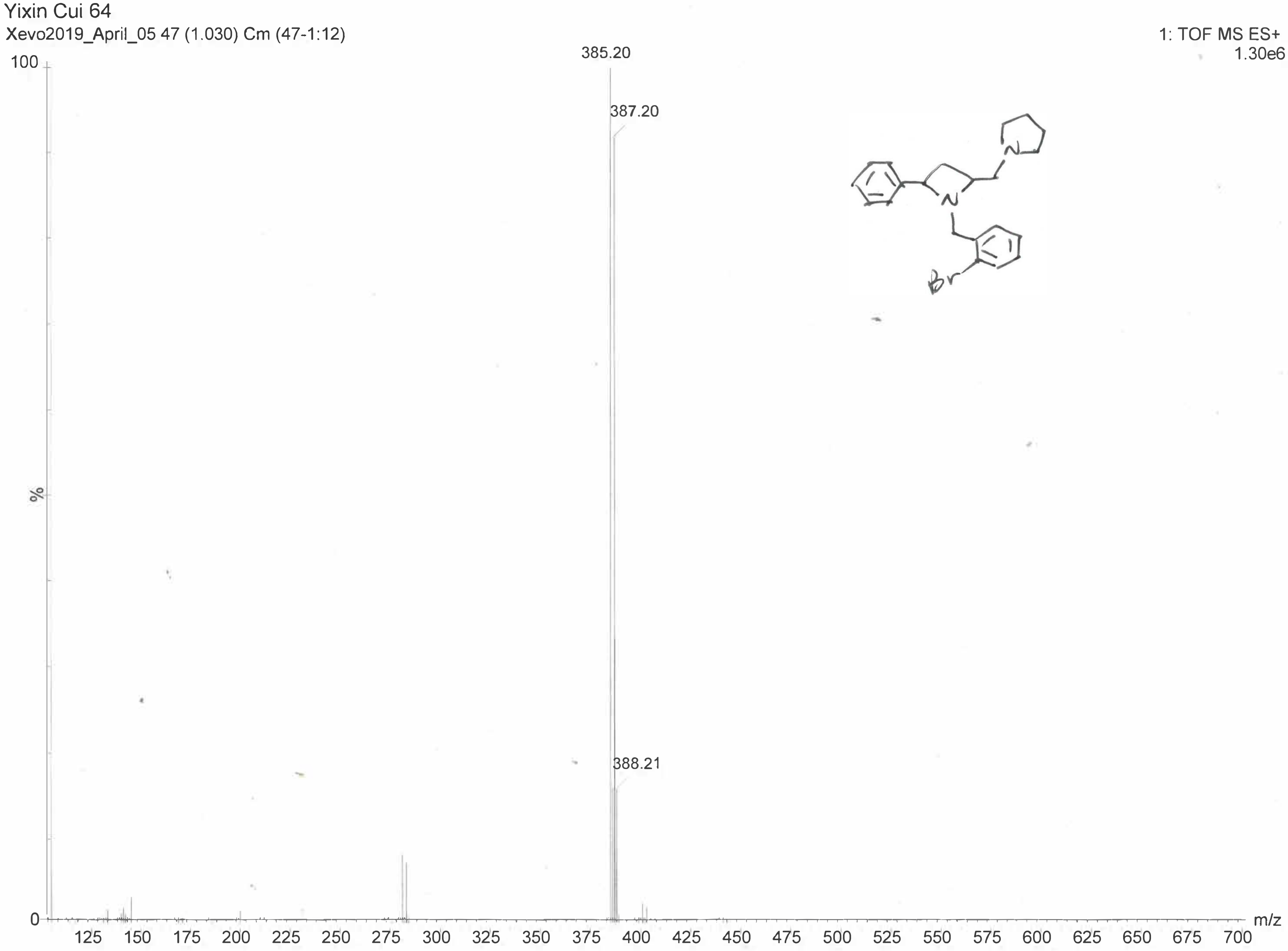

#### **BGAz-001** Mass spec (ii)

Elemental Composition Report **Yixin Cui 64**

Single Mass Analysis

Tolerance = 5.0 PPM / DBE: min = -1.5, max = 50.0

Element prediction: Off

Monoisotopic Mass, Even Electron Ions

27 formula(e) evaluated with 1 results within limits (up to 50 closest results for each mass)

Elements Used:

C: 0-100 H: 0-100 N: 0-5 79Br: 1-1

| | | | | | |
|---|---|---|---|---|---|
| Minimum: | | | -1.5 | | |
| Maximum: | 5.0 | 5.0 | 50.0 | | |
| Mass | Calc. Mass | mDa | PPM | DBE | Formula |
| **385.1277** | 385.1279 | -0.2 | **-0.5** | 9.5 | **C21 H26 N2 79Br** |

**BGAz-002** Mass Spec (i)

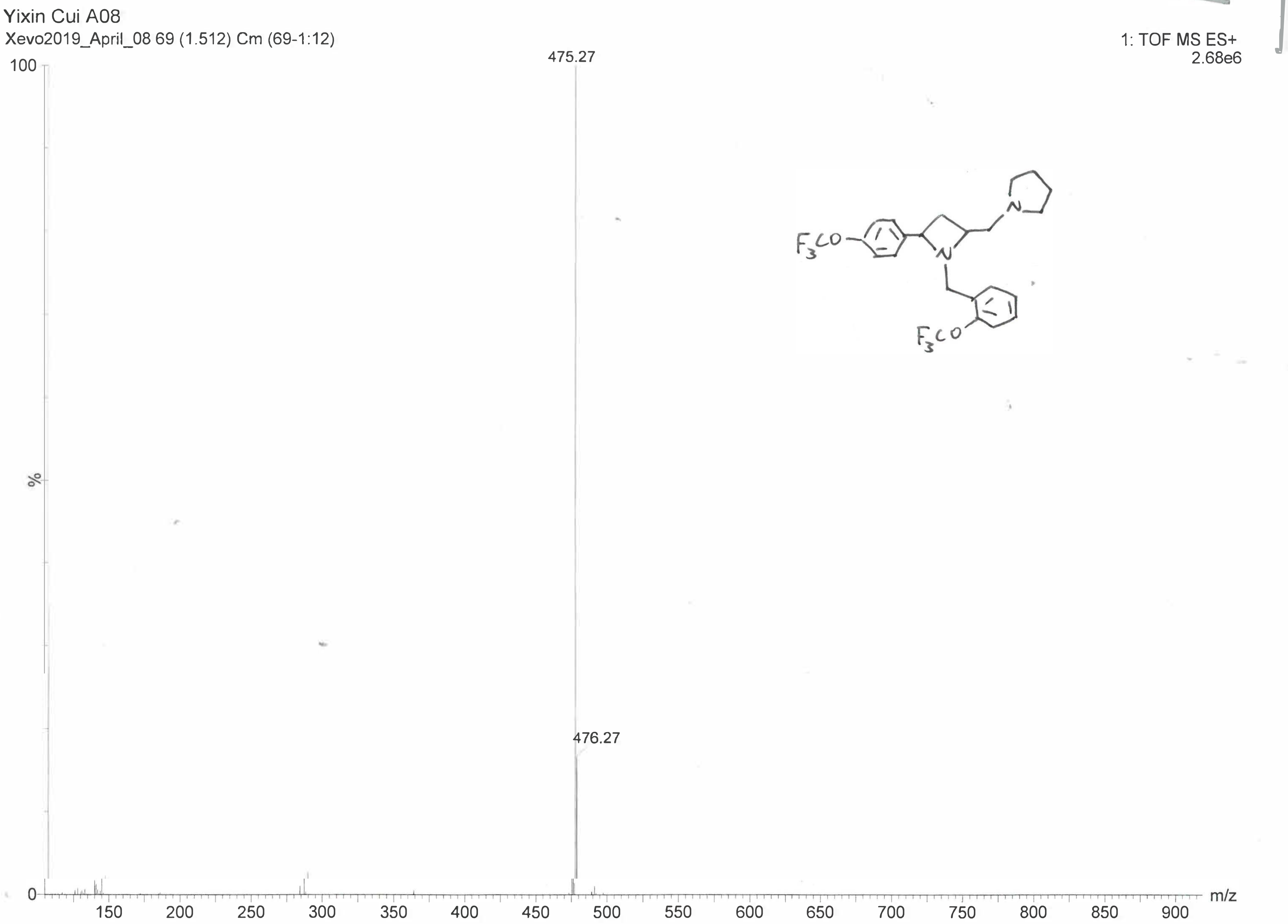

Elemental Composition Report **Yixin Cui A08**

Single Mass Analysis

Tolerance = 5.0 PPM / DBE: min = -1.5, max = 50.0

Element prediction: Off

Monoisotopic Mass, Odd and Even Electron Ions

100 formula(e) evaluated with 1 results within limits (up to 50 closest results for each mass)

Elements Used:

C: 0-100 H: 0-100 N: 0-3 O: 2-2 19F: 2-6

| | | | | | |
|---|---|---|---|---|---|
| Minimum: | | | -1.5 | | |
| Maximum: | 5.0 | 5.0 | 50.0 | | |
| Mass | Calc. Mass | mDa | PPM | DBE | Formula |
| **475.1825** | 475.1820 | 0.5 | **1.1** | 9.5 | **C23 H25 N2 O2 19F6** |

**BGAz-002** Mass Spec (iii)

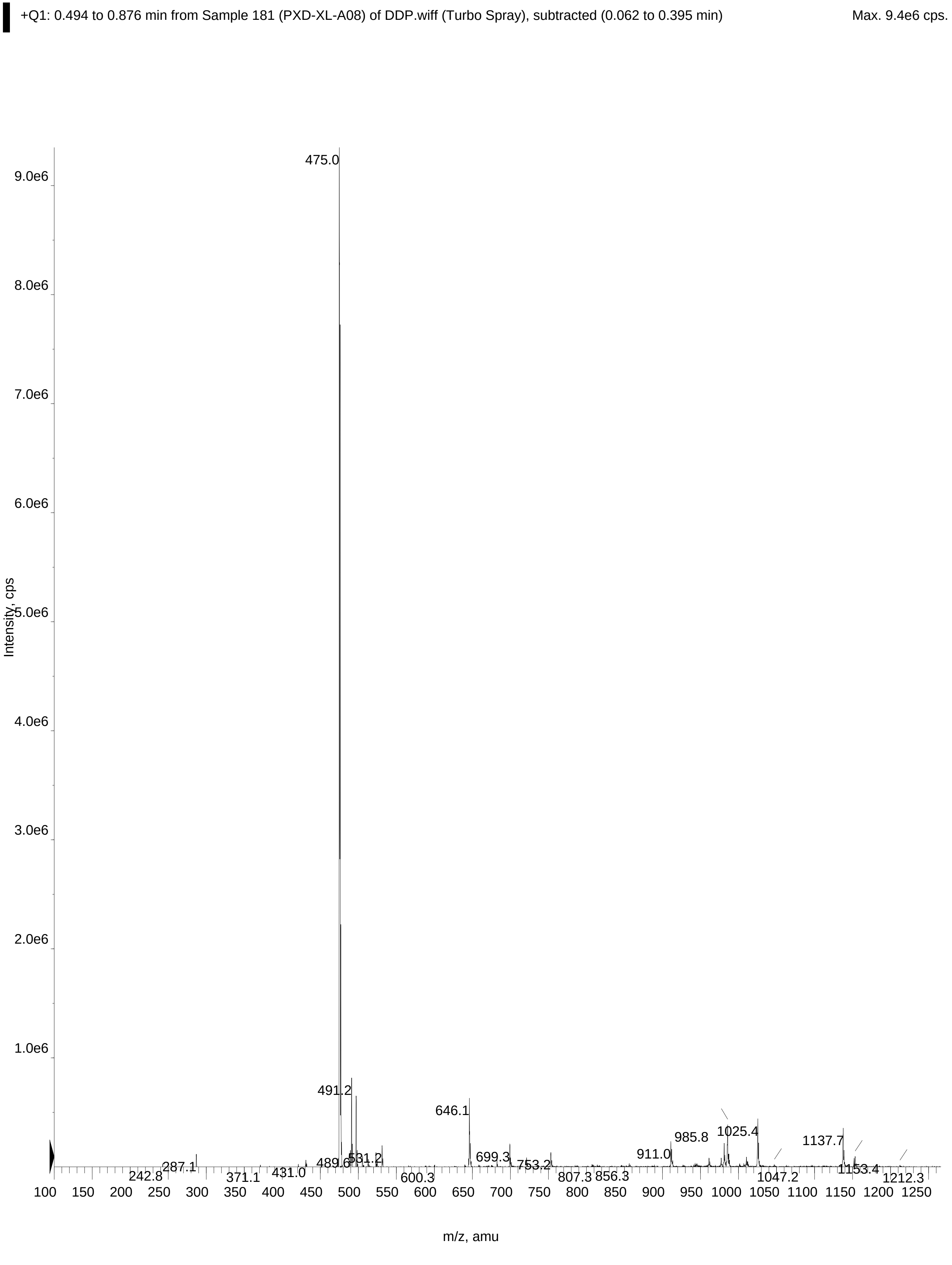

BGAz-002 Mass Spec (iv)

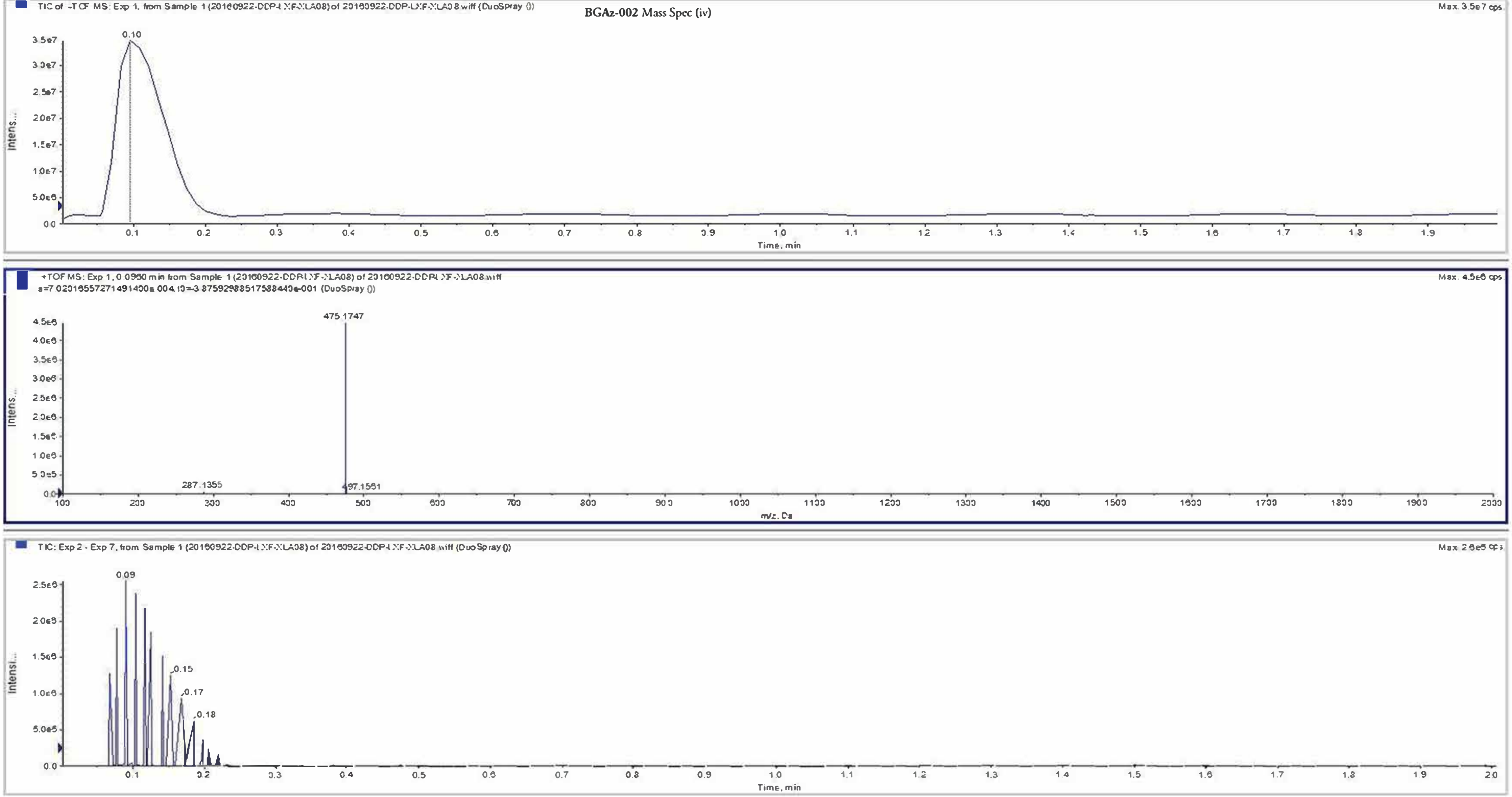

**BGAz-003** Mass spec (i)

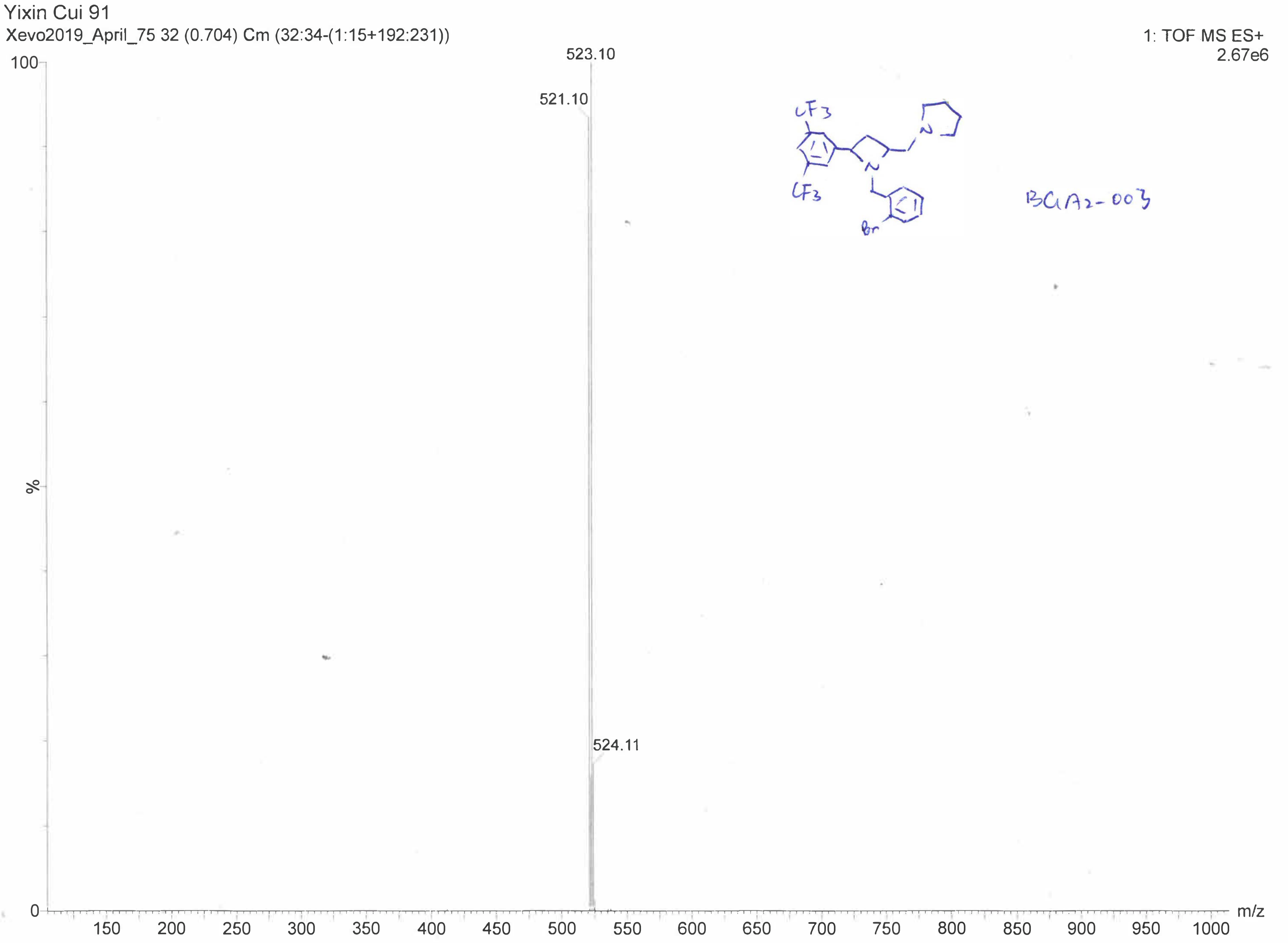

Elemental Composition Report **Yixin Cui YC91**

Single Mass Analysis

Tolerance = 5.0 PPM / DBE: min = -1.5, max = 50.0

Element prediction: Off

Monoisotopic Mass, Even Electron Ions

189 formula(e) evaluated with 1 results within limits (up to 50 closest results for each mass)

Elements Used:

C: 0-100 H: 0-100 N: 0-5 19F: 2-8 79Br: 1-1

| | | | | | |
|---|---|---|---|---|---|
| Minimum: | | | -1.5 | | |
| Maximum: | 5.0 | 5.0 | 50.0 | | |
| Mass | Calc. Mass | mDa | PPM | DBE | Formula |
| **521.1026** | 521.1027 | -0.1 | **-0.2** | 9.5 | **C23 H24 N2 19F6 79Br** |

**BGAz-003** Mass spec (iii)

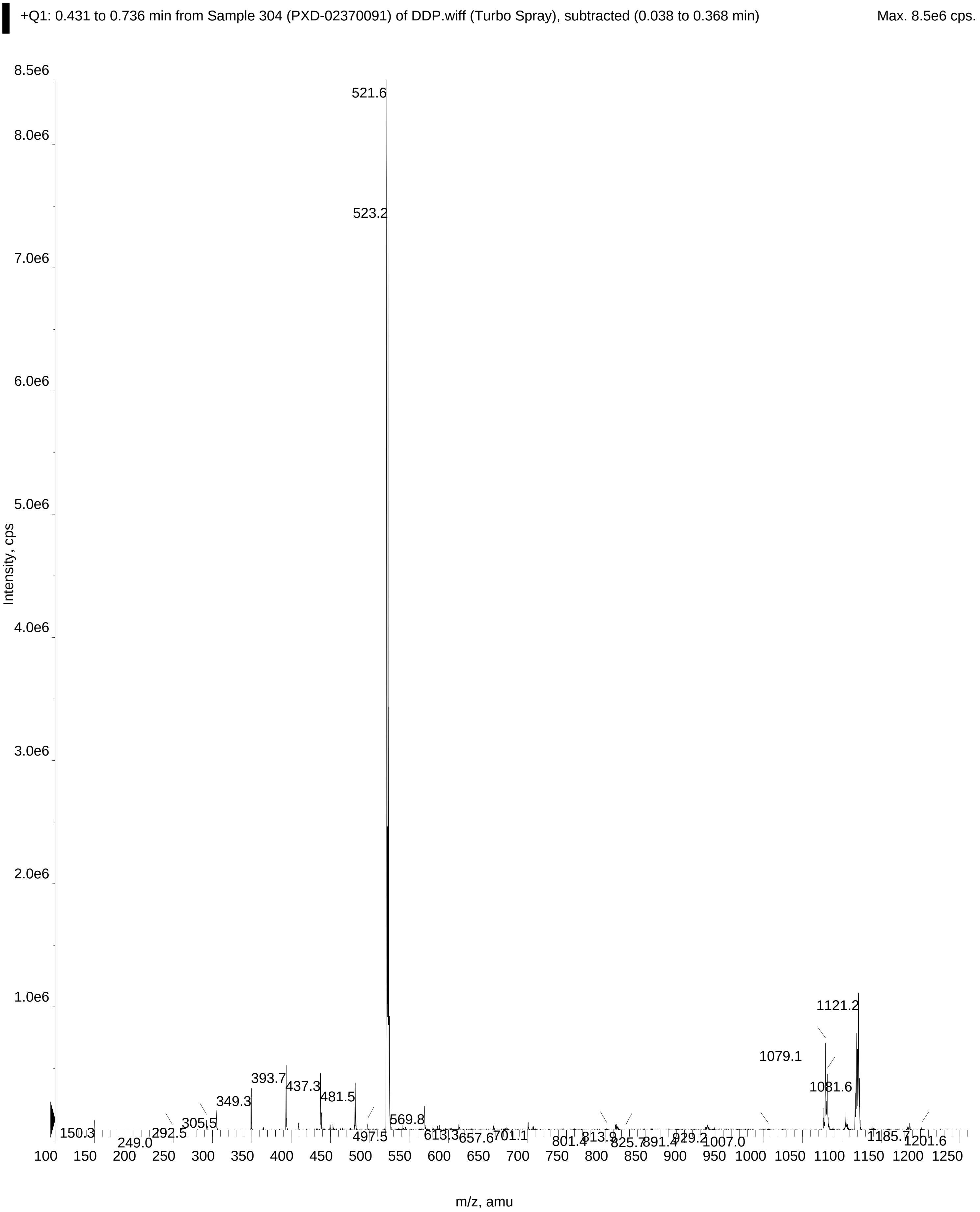

**BGAz-004** Mass spec (i)

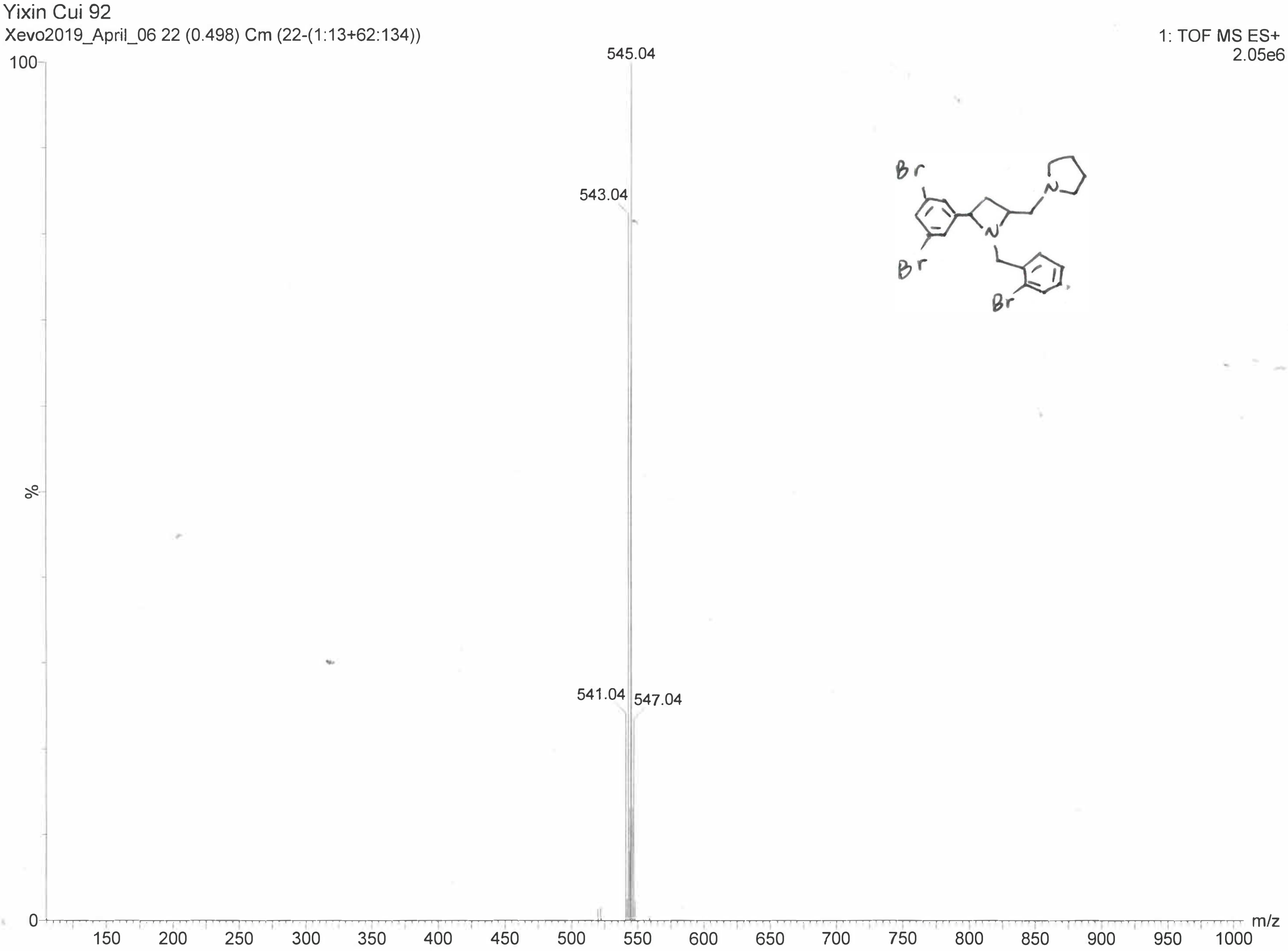

### **BGAz-004** Mass spec (ii)

Elemental Composition Report **Yixin Cui 92**

Single Mass Analysis

Tolerance = 5.0 PPM / DBE: min = -1.5, max = 50.0

Element prediction: Off

Monoisotopic Mass, Odd and Even Electron Ions

104 formula(e) evaluated with 1 results within limits (up to 50 closest results for each mass)

Elements Used:

C: 0-100 H: 0-100 N: 1-5 79Br: 1-5

| | | | | | |
|---|---|---|---|---|---|
| Minimum: | | | -1.5 | | |
| Maximum: | | 5.0 | 5.0 | 50.0 | |
| Mass | Calc. Mass | mDa | PPM | DBE | Formula |
| **540.9489** | 540.9490 | -0.1 | **-0.2** | 9.5 | **C21 H24 N2 79Br3** |

#### BGAz-005 Mass spec (i)

D:\Data\Chemistry Mass Spectrometry Project\Yixin Cui YC-GIBH-122_GE2_01_300.d 7/17/2019 4:49:49 PM

**Yixin Cui YC-GIBH-122**

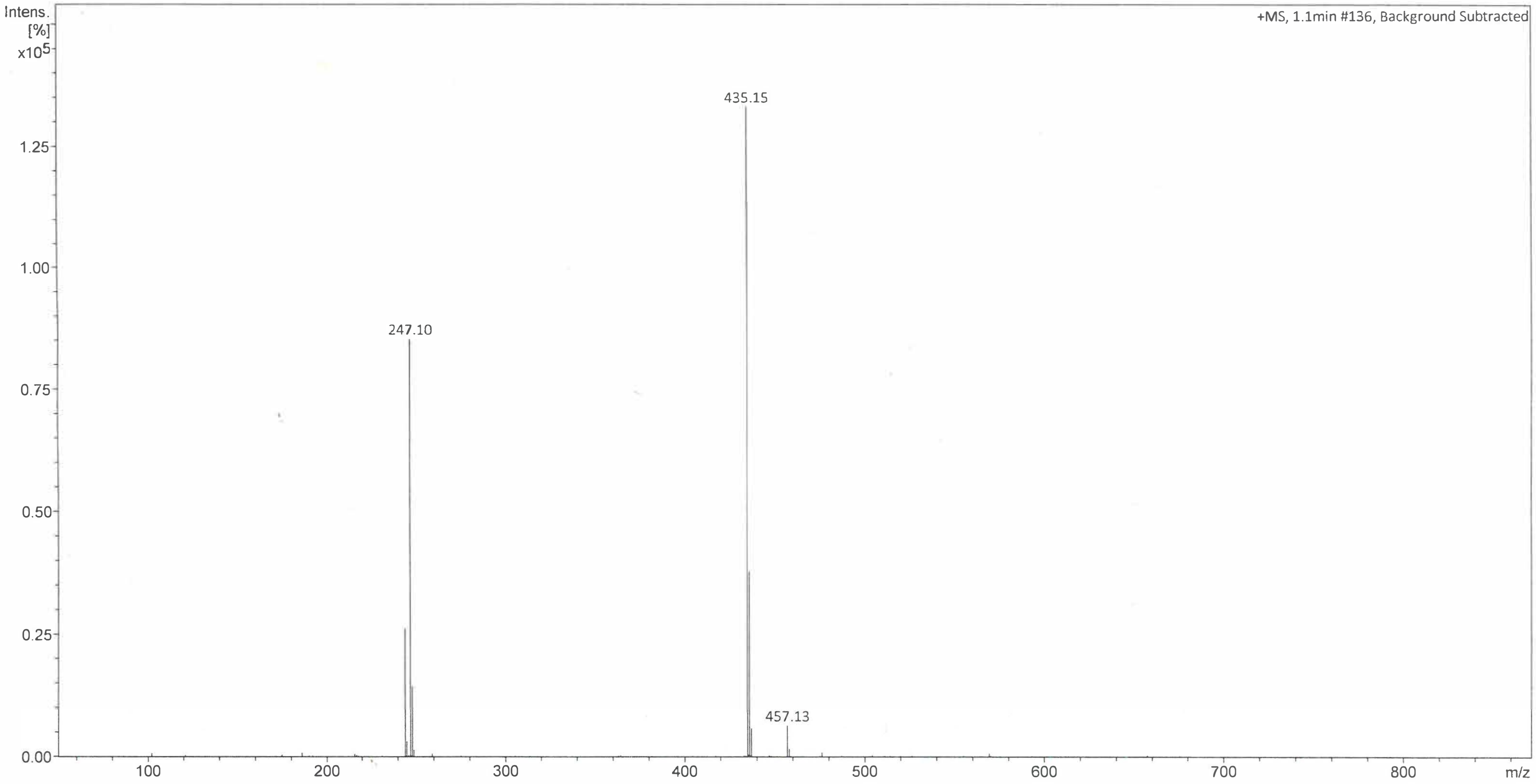

#### BGAz-005 Mass spec (ii)

D:\Data\Chemistry Mass Spectrometry Project\Yixin Cui YC-GIBH-122_GE2_01_300.d 7/17/2019 4:49:49 PM

### Yixin Cui YC-GIBH-122

| Meas. m/z | # | Ion Formula | m/z | err [ppm] | mSigma | # mSigma | Score | rdb | e⁻ Conf | N-Rule |
|---|---|---|---|---|---|---|---|---|---|---|
| 435.1505 | 1 | C20H21F6N2O2 | 435.1502 | -0.8 | 18.7 | 1 | 100.00 | 8.5 | even | ok |

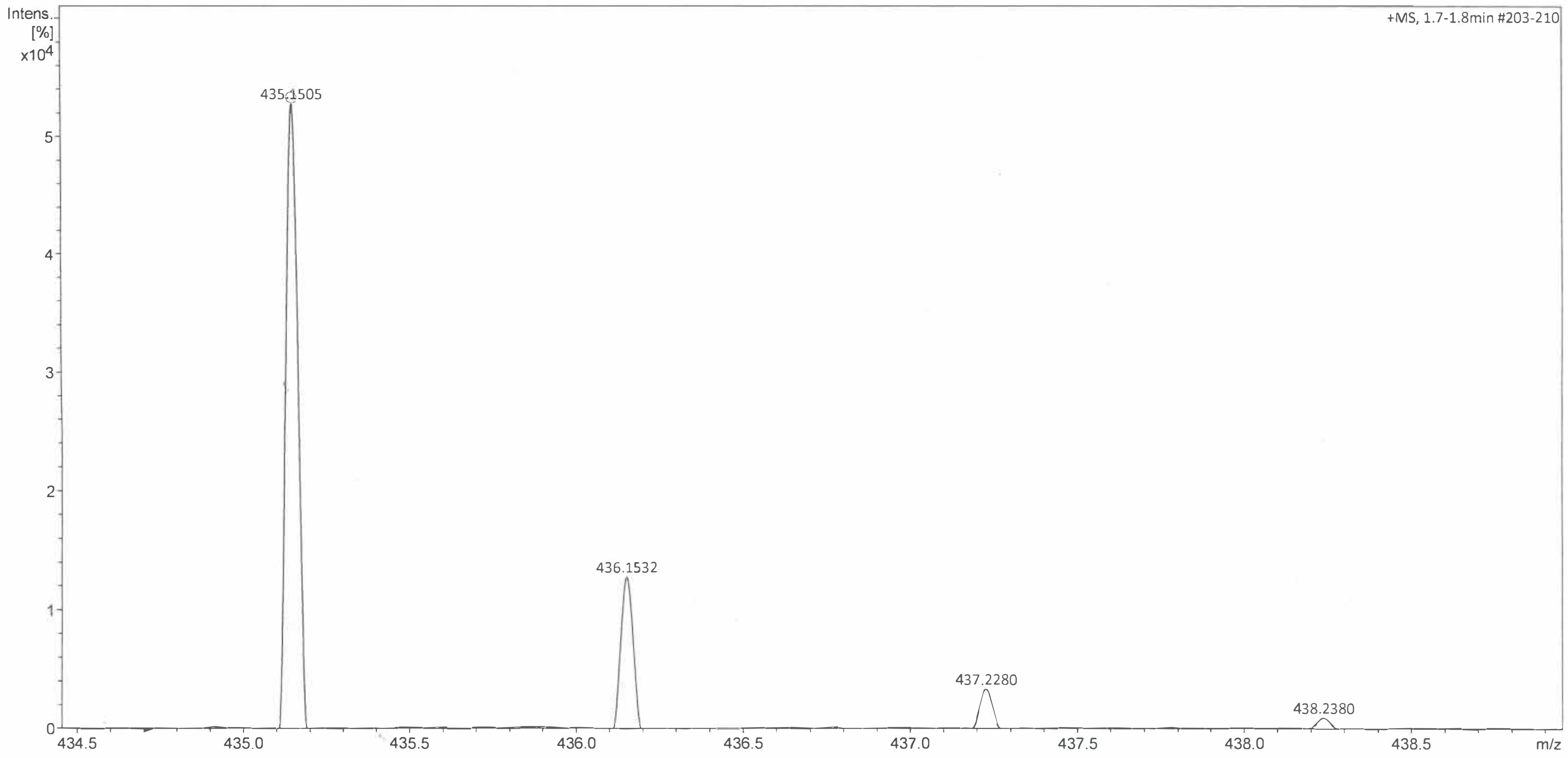

数据文件: E:\DATA\单次进样\PXD-064-YJ 2019-10-18 17-09-20.D
样品名称: PXD-064-YJ

**BGAz-001** HPLC

=====================================================================

操作者 : 系统
仪器 : 1260 位置 : P1-B-02
进样日期 : 2019-10-18 17:10:01
进样量 : 1.500 µl
采集方法 : C:\CHEM32\1\METHODS\DEF_LC.M
最后修改 : 2019-10-18 17:03:24 : 系统
(调用后修改)
分析方法 : C:\CHEM32\1\METHODS\DEF_LC.M
最后修改 : 2020-7-24 11:07:08 : 系统
(调用后修改)
附加信息: 峰被手动积分

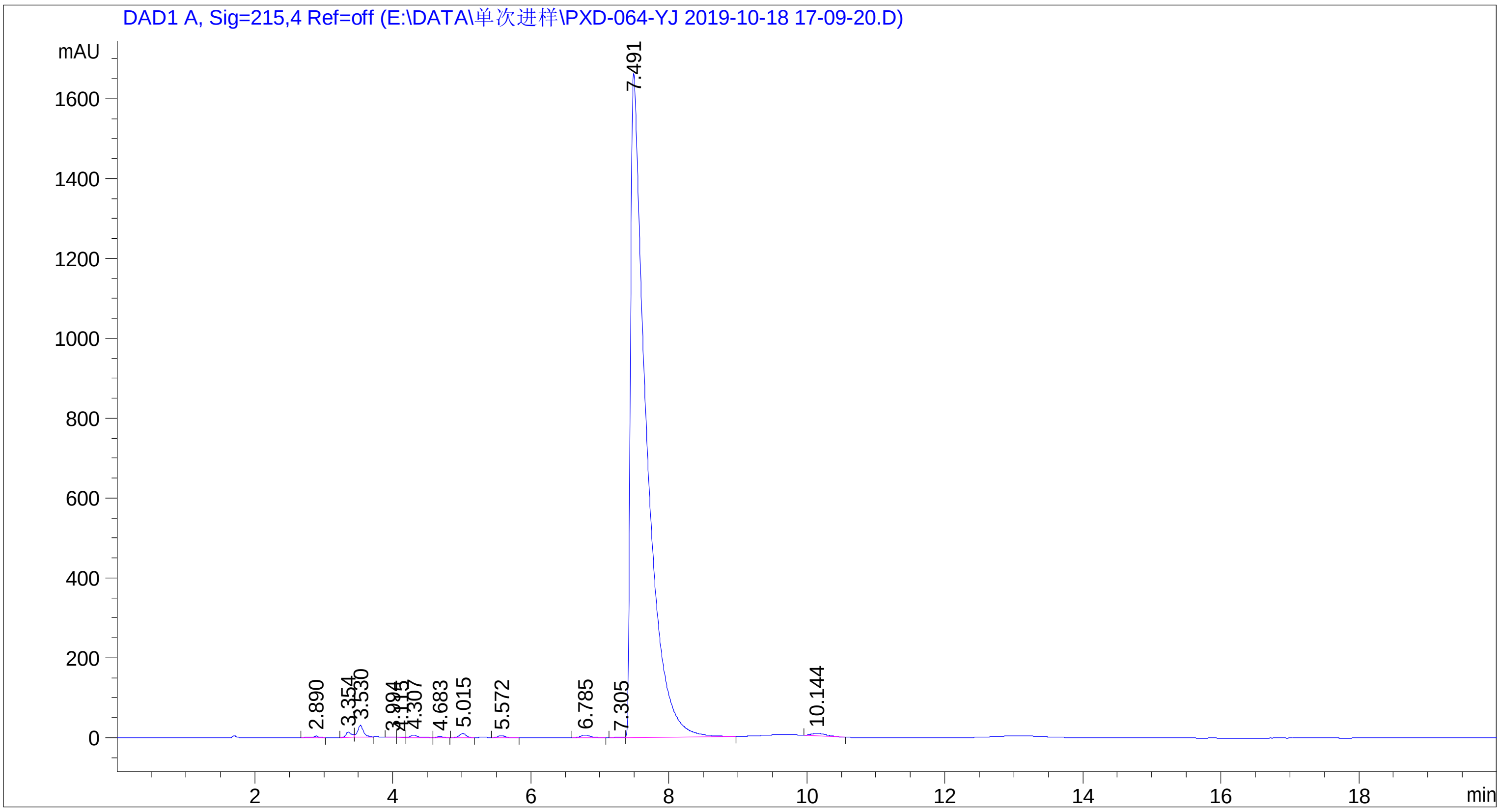

=====================================================================

面积百分比报告

=====================================================================

排序 : 信号
乘积因子 : 1.0000
稀释因子 : 1.0000
内标使用乘积因子和稀释因子

信号 1: DAD1 A, Sig=215,4 Ref=off

| 峰 # | 保留时间 [min] | 类型 | 峰宽 [min] | 峰面积 [mAU*s] | 峰高 [mAU] | 峰面积 % |
|---|---|---|---|---|---|---|
| 1 | 2.890 | BB | 0.0830 | 21.44737 | 3.62478 | 0.0792 |
| 2 | 3.354 | BV | 0.0843 | 77.17467 | 13.18584 | 0.2850 |
| 3 | 3.530 | VB | 0.0873 | 175.25566 | 29.49627 | 0.6472 |
| 4 | 3.994 | BV | 0.0872 | 7.29503 | 1.30526 | 0.0269 |
| 5 | 4.115 | VV | 0.0839 | 7.49826 | 1.36981 | 0.0277 |
| 6 | 4.307 | VB | 0.1102 | 45.70713 | 6.04725 | 0.1688 |

| 峰 # | 保留时间 [min] | 类型 | 峰宽 [min] | 峰面积 [mAU*s] | 峰高 [mAU] | 峰面积 % |
|---|---|---|---|---|---|---|
| 7 | 4.683 | BB | 0.0864 | 15.68594 | 2.84365 | 0.0579 |
| 8 | 5.015 | BB | 0.1050 | 69.37886 | 10.25137 | 0.2562 |
| 9 | 5.572 | BB | 0.1104 | 35.44512 | 5.01557 | 0.1309 |
| 10 | 6.785 | BB | 0.1397 | 63.71717 | 6.77354 | 0.2353 |
| 11 | 7.305 | BV | 0.1188 | 14.33696 | 1.92685 | 0.0529 |
| 12 | 7.491 | VB | 0.2196 | 2.64540e4 | 1662.89099 | 97.6898 |
| 13 | 10.144 | BB | 0.2361 | 92.64355 | 6.13776 | 0.3421 |

总量 : 2.70796e4 1750.86895

==============================================================

*** 报告结束 ***

数据文件: E:\DATA\单次进样\PXD-XL-A08 2019-11-18 14-29-04.D
样品名称: PXD-XL-A08

=====================================================================

操作者 : 系统
仪器 : 1260 位置 : P1-A-05
进样日期 : 2019-11-18 14:29:46
进样量 : 3.000 µl
采集方法 : C:\CHEM32\1\METHODS\DEF_LC.M
最后修改 : 2019-11-18 14:38:59 : 系统
(调用后修改)
分析方法 : C:\CHEM32\1\METHODS\DEF_LC.M
最后修改 : 2020-7-24 11:07:08 : 系统
(调用后修改)

**BGAz-002** HPLC

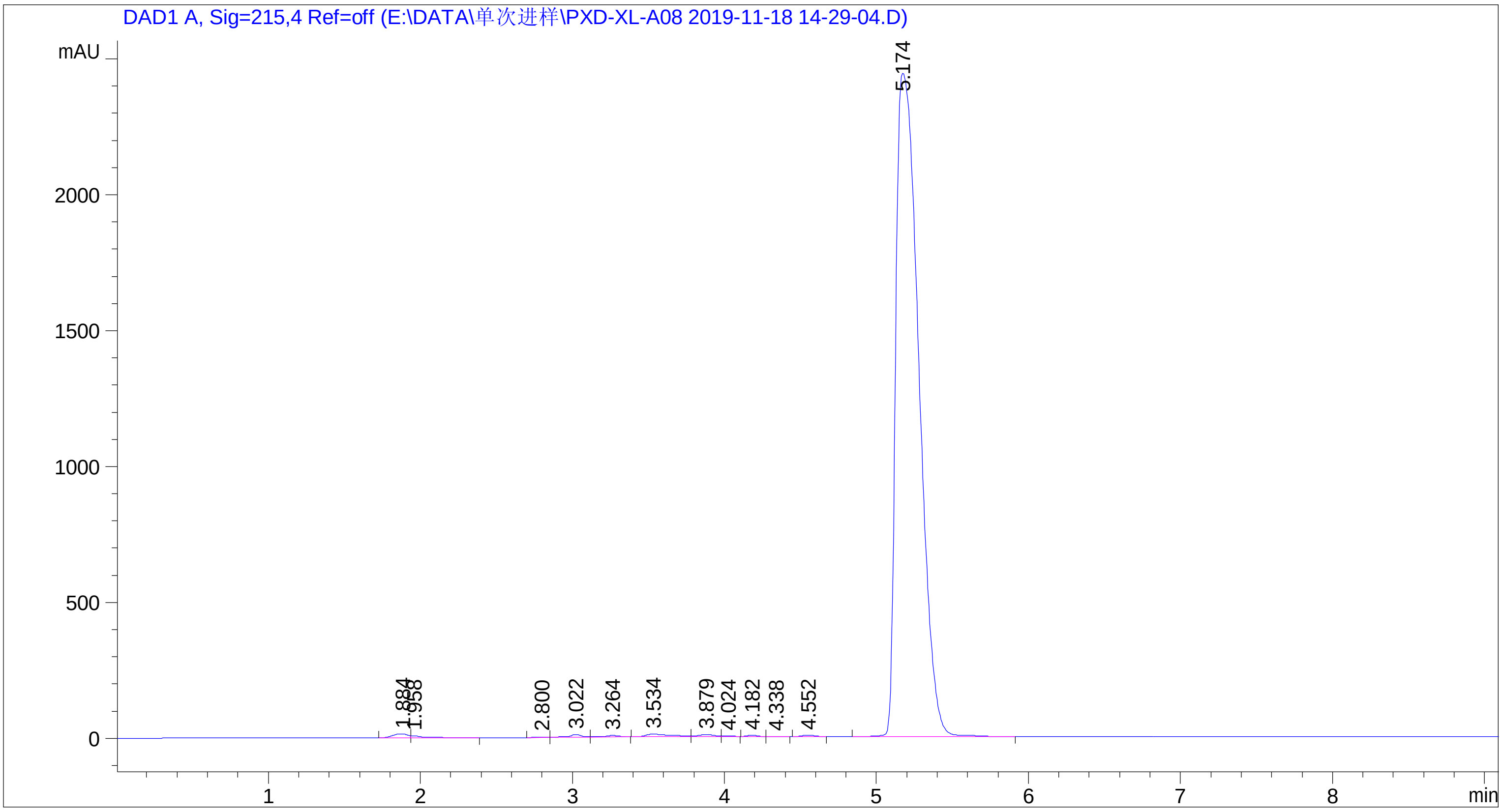

=====================================================================

面积百分比报告

=====================================================================

排序 : 信号
乘积因子 : 1.0000
稀释因子 : 1.0000
内标使用乘积因子和稀释因子

信号 1: DAD1 A, Sig=215,4 Ref=off

| 峰 # | 保留时间 [min] | 类型 | 峰宽 [min] | 峰面积 [mAU*s] | 峰高 [mAU] | 峰面积 % |
|---|---|---|---|---|---|---|
| 1 | 1.884 | BV | 0.0953 | 94.87852 | 13.28846 | 0.3642 |
| 2 | 1.958 | VB | 0.0719 | 39.90796 | 7.52465 | 0.1532 |
| 3 | 2.800 | BB | 0.0540 | 6.93734 | 2.02328 | 0.0266 |
| 4 | 3.022 | BV | 0.0805 | 55.11722 | 9.97407 | 0.2115 |
| 5 | 3.264 | VB | 0.1003 | 35.72900 | 5.32175 | 0.1371 |
| 6 | 3.534 | BV | 0.1537 | 111.60405 | 10.03849 | 0.4283 |
| 7 | 3.879 | VV | 0.1092 | 51.24211 | 7.18935 | 0.1967 |

数据文件: E:\DATA\单次进样\PXD-XL-A08 2019-11-18 14-29-04.D
样品名称: PXD-XL-A08

| 峰 # | 保留时间 [min] | 类型 | 峰宽 [min] | 峰面积 [mAU*s] | 峰高 [mAU] | 峰面积 % |
|---|---|---|---|---|---|---|
| 8 | 4.024 | VB | 0.0753 | 13.19106 | 2.68375 | 0.0506 |
| 9 | 4.182 | BB | 0.0757 | 18.94328 | 4.11529 | 0.0727 |
| 10 | 4.338 | BB | 0.0786 | 5.63545 | 1.16248 | 0.0216 |
| 11 | 4.552 | BB | 0.0956 | 30.81825 | 5.16755 | 0.1183 |
| 12 | 5.174 | BB | 0.1664 | 2.55907e4 | 2439.98096 | 98.2191 |

总量 : 2.60547e4 2508.47007

*** 报告结束 ***

数据文件: E:\DATA\新建文件夹\PXD-02370091 2017-05-10 16-13-345225.D
样品名称: pxd-02370091

**BGAz-003** HPLC

操作者 : 系统
仪器 : HPLC1260 位置 : P1-C-01
进样日期 : 2017-5-10 16:14:16
进样量 : 3.000 µl
采集方法 : C:\CHEM32\1\METHODS\DEF_LC.M
最后修改 : 2017-5-10 16:39:05 : 系统
(调用后修改)
分析方法 : C:\CHEM32\1\METHODS\DEF_LC.M
最后修改 : 2017-5-10 18:41:59 : 系统
(调用后修改)
附加信息: 峰被手动积分

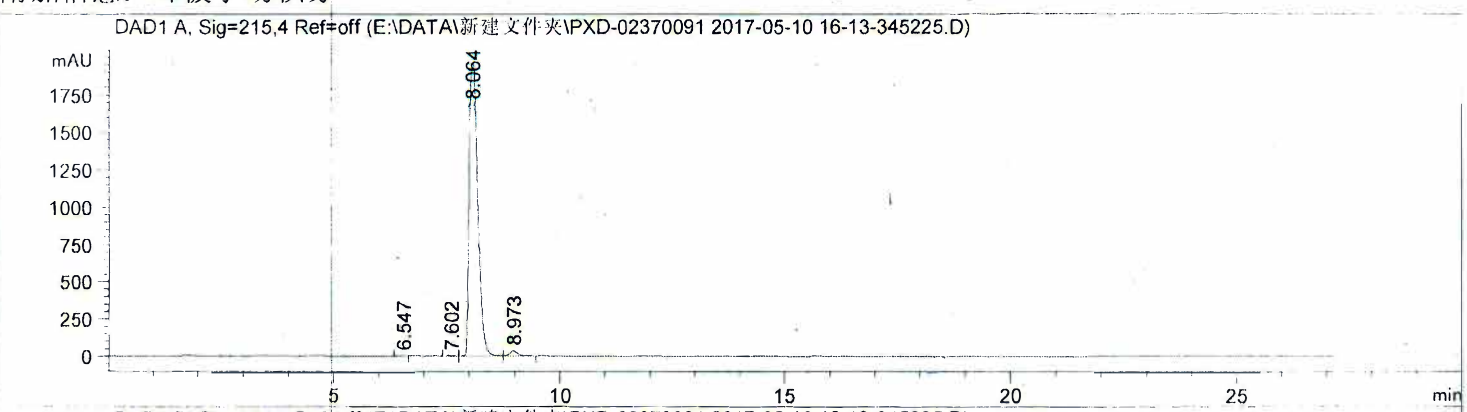

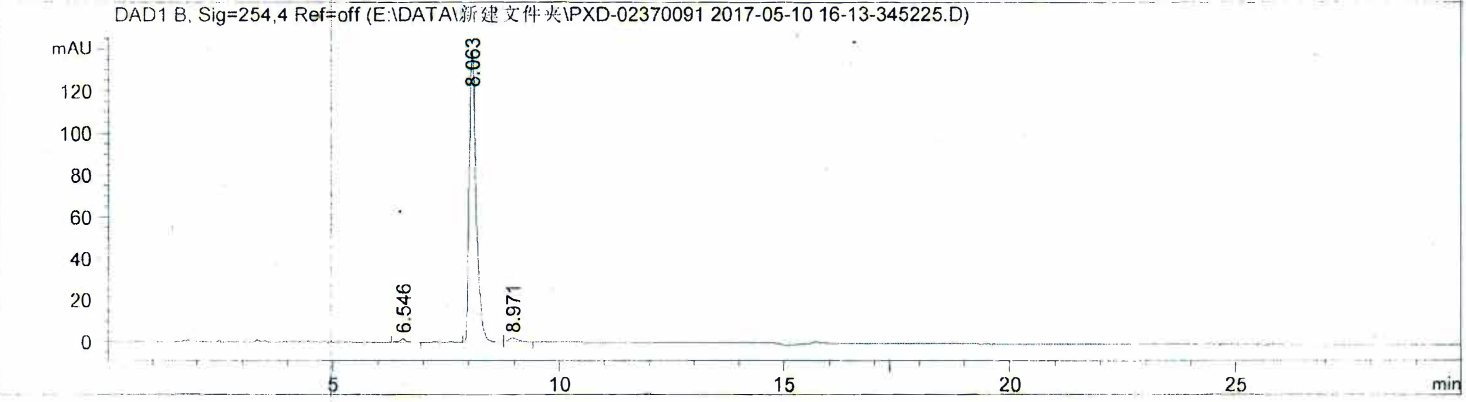

#### 面积百分比报告

排序 : 信号
乘积因子 : 1.0000
稀释因子 : 1.0000
内标使用乘积因子和稀释因子

信号 1: DAD1 A, Sig=215,4 Ref=off

| 峰 # | 保留时间 [min] | 类型 | 峰宽 [min] | 峰面积 [mAU*s] | 峰高 [mAU] | 峰面积 % |
|---|---|---|---|---|---|---|
| 1 | 6.547 | BB | 0.1058 | 16.01622 | 2.34190 | 0.0606 |
| 2 | 7.602 | BB | 0.1330 | 11.39994 | 1.26682 | 0.0431 |
| 3 | 8.064 | BV | 0.2110 | 2.59805e4 | 1956.12817 | 98.2938 |
| 4 | 8.973 | VB | 0.1916 | 423.54568 | 33.47721 | 1.6024 |

总量 : 2.64314e4 1993.21410

数据文件: E:\DATA\新建文件夹\PXD-02370091 2017-05-10 16-13-345225.D
样品名称: pxd-02370091

信号 2: DAD1 B, Sig=254,4 Ref=off

| 峰 # | 保留时间 [min] | 类型 | 峰宽 [min] | 峰面积 [mAU*s] | 峰高 [mAU] | 峰面积 % |
|---|---|---|---|---|---|---|
| 1 | 6.546 | BB | 0.1482 | 17.09118 | 1.68668 | 1.1108 |
| 2 | 8.063 | BB | 0.1638 | 1499.30359 | 139.03644 | 97.4409 |
| 3 | 8.971 | BB | 0.1811 | 22.28556 | 1.79241 | 1.4484 |

总量 : 1538.68032 142.51552

*** 报告结束 ***

数据文件: E:\DATA\单次进样\PXD-092 2020-04-07 14-01-51.D
样品名称: PXD-092

**BGAz-004** HPLC

=====================================================================

操作者 : 系统
仪器 : 1260 位置 : P1-E-02
进样日期 : 2020-4-7 14:02:34
进样量 : 2.000 µl
采集方法 : C:\CHEM32\1\METHODS\DEF_LC.M
最后修改 : 2020-4-7 14:01:43 : 系统
(调用后修改)
分析方法 : C:\CHEM32\1\METHODS\DEF_LC.M
最后修改 : 2020-7-24 11:07:08 : 系统
(调用后修改)

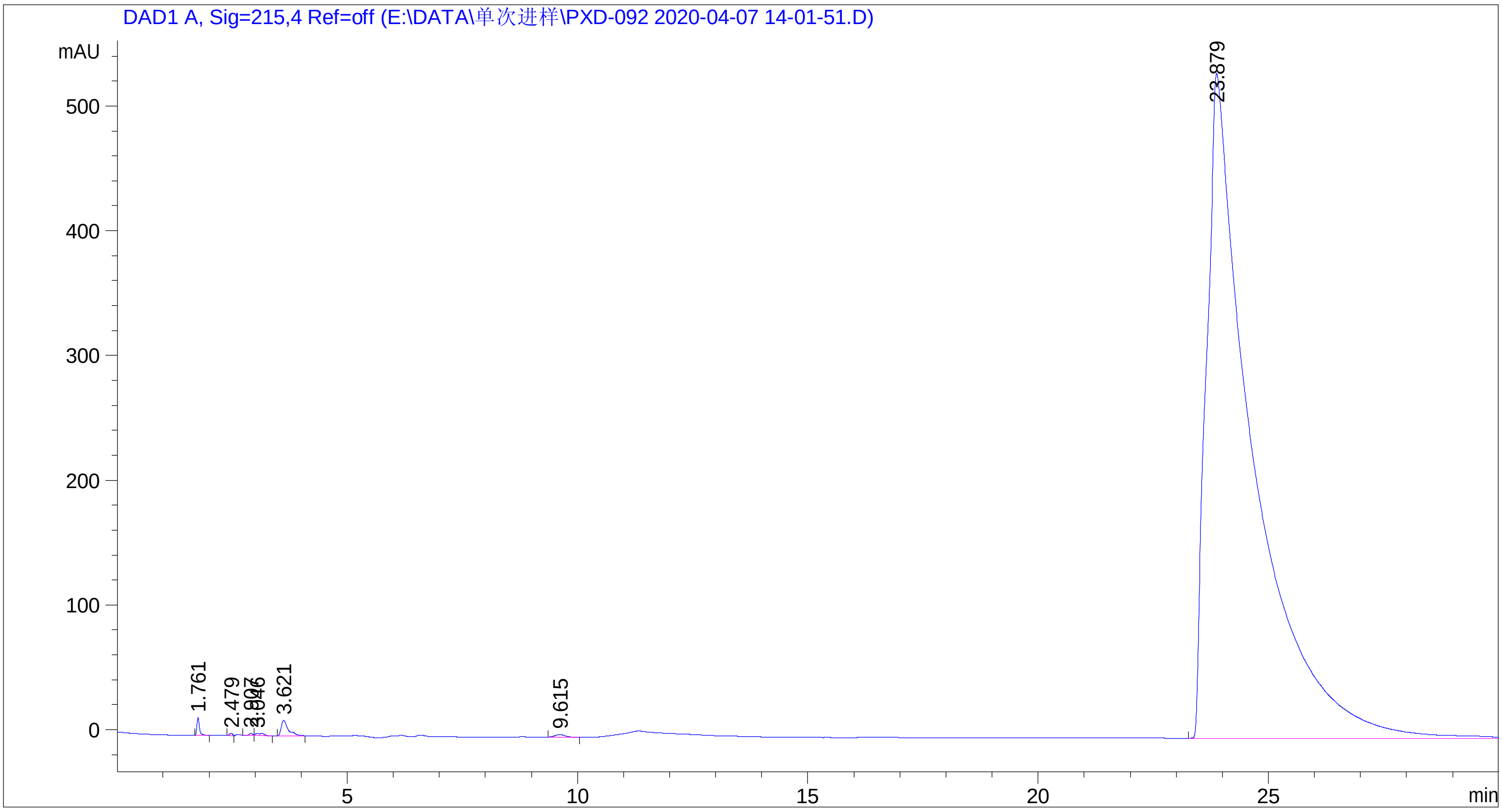

=====================================================================

面积百分比报告

=====================================================================

排序 : 信号
乘积因子 : 1.0000
稀释因子 : 1.0000
内标使用乘积因子和稀释因子

信号 1: DAD1 A, Sig=215,4 Ref=off

| 峰 # | 保留时间 [min] | 类型 | 峰宽 [min] | 峰面积 [mAU*s] | 峰高 [mAU] | 峰面积 % |
|---|---|---|---|---|---|---|
| 1 | 1.761 | BB | 0.0589 | 54.39043 | 14.12255 | 0.1452 |
| 2 | 2.479 | BB | 0.0708 | 7.95832 | 1.90398 | 0.0212 |
| 3 | 2.907 | BV | 0.0814 | 8.05163 | 1.53018 | 0.0215 |
| 4 | 3.046 | VB | 0.1642 | 21.37819 | 1.70590 | 0.0571 |
| 5 | 3.621 | BB | 0.1570 | 132.90109 | 12.40872 | 0.3548 |
| 6 | 9.615 | BB | 0.2310 | 28.47347 | 1.91995 | 0.0760 |
| 7 | 23.879 | BBA | 0.8826 | 3.72078e4 | 532.88928 | 99.3242 |

| 峰 # | 保留时间 [min] | 类型 | 峰宽 [min] | 峰面积 [mAU*s] | 峰高 [mAU] | 峰面积 % |
|---|---|---|---|---|---|---|
| 总量 : | | | | 3.74610e4 | 566.48057 | |

*** 报告结束 ***

数据文件：E:\DATA\单次进样\PXD-122 2019-11-06 14-31-15.D
样品名称：PXD-122

**BGAz-005HPLC**

=====================================================================

操作者 : 系统
仪器 : 1260 位置 : P1-D-02
进样日期 : 2019-11-6 14:31:58
进样量 : 5.000 µl
采集方法 : C:\CHEM32\1\METHODS\DEF_LC.M
最后修改 : 2019-11-6 14:45:14 : 系统
(调用后修改)
分析方法 : C:\CHEM32\1\METHODS\DEF_LC.M
最后修改 : 2019-11-6 14:34:09 : 系统
(调用后修改)
附加信息：峰被手动积分

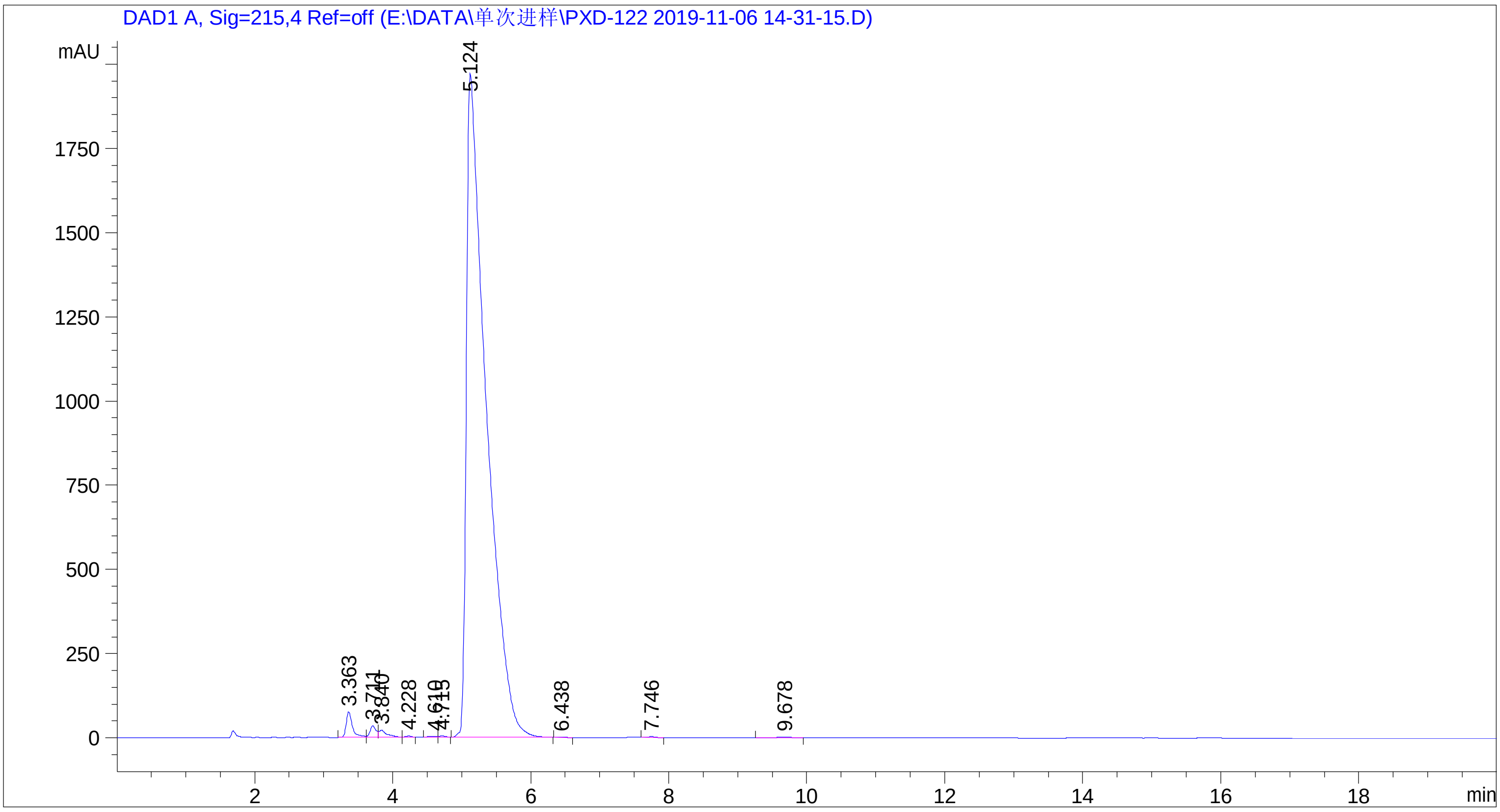

=====================================================================

面积百分比报告

=====================================================================

排序 : 信号
乘积因子 : 1.0000
稀释因子 : 1.0000
内标使用乘积因子和稀释因子

信号 1: DAD1 A, Sig=215,4 Ref=off

| 峰 # | 保留时间 [min] | 类型 | 峰宽 [min] | 峰面积 [mAU*s] | 峰高 [mAU] | 峰面积 % |
|---|---|---|---|---|---|---|
| 1 | 3.363 | BV | 0.0930 | 475.38925 | 75.93010 | 1.1842 |
| 2 | 3.711 | VV | 0.0900 | 207.26859 | 34.52227 | 0.5163 |
| 3 | 3.840 | VB | 0.1084 | 160.22266 | 20.23956 | 0.3991 |
| 4 | 4.228 | BB | 0.0737 | 13.40368 | 2.90957 | 0.0334 |
| 5 | 4.610 | BV | 0.0908 | 15.34426 | 2.45905 | 0.0382 |
| 6 | 4.715 | VB | 0.0960 | 19.88466 | 3.13596 | 0.0495 |

| 峰 # | 保留时间 [min] | 类型 | 峰宽 [min] | 峰面积 [mAU*s] | 峰高 [mAU] | 峰面积 % |
|---|---|---|---|---|---|---|
| 7 | 5.124 | BB | 0.2762 | 3.91908e4 | 1969.03772 | 97.6275 |
| 8 | 6.438 | BB | 0.1139 | 8.42819 | 1.17196 | 0.0210 |
| 9 | 7.746 | BB | 0.1323 | 21.34094 | 2.53304 | 0.0532 |
| 10 | 9.678 | BB | 0.2147 | 31.13356 | 2.17860 | 0.0776 |

总量 : 4.01432e4 2114.11783

=====================================================================

*** 报告结束 ***

### *In vivo* pharmacokinetics parameters report BGAz compounds (combination of three reports)

### Report on PK Results of Combined Dosing with BGAz-001–005

**2019.12.27**

#### 1 Materials and instrument combined dosing

An MS2 Turnover type oscillator was purchased from IKA Work's Guangzhou (China), and a 5415R Chromatographic analyses were conducted using an Agilent 1290 Infinity II high performance liquid chromatography and Agilent Technologies 6470Triple Quad LC/MS. Propranolol (≥90% in purity, internal standard, IS) was purchased from the Sigma Chemical Co. (China). Formic acid (HPLC grade) and methanol (HPLC grade) were purchased from DIKMA Co. (China). All chemicals and solvents were analytical grade. Water was purified using a Millipore (AK, USA) laboratory ultra-pure water system (0.2 μm filter).

#### 2 Animal

KM mice, weighing 25–35 g (Beijing Vital River Laboratory Animal Technology Co., Ltd., China) were utilised for the studies. The protocols were approved by the Animal Care and Use Committee, GIBH. Animals were maintained on standard animal chow and water *ad libitum*, in a climate-controlled room (23±1 °C 30–70% relative humidity, a minimum of ten exchanges of room air per hour and a 12 h light/dark cycle) for one week prior to experiments.

#### 3 Pharmacokinetic studies

Compounds **BGAz-001**，**BGAz-002**，**BGAz-003**，**BGAz-004** and **BGAz-005** were dissolved mixed in a solution containing DMSO (2%), ethanol (4%), Cremophor EL (4%) and dd$H_2O$ (90%); Pharmacokinetic properties of KM (male) were determined following IV, PO and IP administration. Animals were randomly distributed into three experimental groups (n = 4). The PO group was given mixed solution with 5 mg/kg by gastric gavage. The IV group was dosed with mixed solution 1mg/kg by injection into the tail vein. The IP group was dosed with mixed solution 1mg/kg by intraperitoneal injection. After single administration, whole blood samples (100 μL) were obtained from the orbital venous plexus at the following time points after dosing: 5, 15, 30 min and 1, 3, 5, 8, 24 hour (PO and IP groups) and at the following time points after dosing 2, 10, 30 min and 1, 3, 5, 8, 24 hour (IV group). Whole blood samples were collected in heparinised tubes. The plasma fraction was immediately separated by centrifugation. The mice were humanely euthanasia by carbon dioxide 24 hours after experiment without pain.

#### 4 Plasma sample and brain sample analysis

##### *4.1 Standard curve sample preparation*

The compounds were dissolved in DMSO (2mg/mL) and diluted with to series concentration (methanol:$H_2O$, 1:1, 10 μL) and blank plasma (50 μL) were added to 1.5 mL tubes and vortexed for 3 min, then acetonitrile-containing internal standard (150 μL) was added and vortexed for a further 5 min, and subsequently spun in a centrifuge at 13000 ×*g* for 40 min at 4 °C, the final concentrations were as follow: 5, 10, 20, 50, 100, 200, 500, 1000 ng/mL.

##### *4.2 Plasma and preparation*

Plasma samples were prepared using a protein precipitation method. Solution (10 μL, methanol $H_2O$, 1:1) and plasma samples (50 μL) were added to 1.5 mL tubes and vortex for 3 min, then acetonitrile-containing internal standard (150 μL) was added and vortex for 5 min, and subsequently spun in a centrifuge at 13000 ×*g* for 40 min at 4 °C.

#### 5 LC/MS/MS analysis

After centrifugation, supernatant (100 μL) was transfer to 96 well plates and analysed by LC-MS/MS using an Agilent 1290 Infinity II HPLC and an Agilent Technologies 6470Triple Quad LC/MS.

### 6 Results

**Table 1.** Pharmacokinetic parameters for **BGAz-001.**

| | **BGAz-001** | | |
|---|---|---|---|
| | **PO** | **IV** | **IP** |
| **Animal Number KM mice** | ♂4 | ♂4 | ♂4 |
| **Dose level mg/kg** | 5 | 1 | 1 |
| **AUC(0-∞) μg*h** | 147.069 | 143.307 | 21.656 |
| **$T_{1/2}$ (h)** | 1.482 | 1.003 | 2.128 |
| **$T_{max}$ (h)** | 0.312 | 0.033 | 0.125 |
| **$C_{max}$（μg/L）** | 54.708 | 157.627 | 21.923 |
| **BA (%)** | | 20.53 | |

**Table 2.** Pharmacokinetic parameters for **BGAz-002.**

| | **BGAz-002** | | |
|---|---|---|---|
| | **PO** | **IV** | **IP** |
| **Animal Number KM mice** | ♂4 | ♂4 | ♂4 |
| **Dose level mg/kg** | 5 | 1 | 1 |
| **AUC(0-∞) μg*h** | 1268.03 | 440.39 | 629.982 |
| **$T_{1/2}$ (h)** | 24.921 | 8.349 | 16.122 |
| **$T_{max}$ (h)** | 1.875 | 0.033 | 0.167 |
| **$C_{max}$（μg/L）** | 53.858 | 188.69 | 39.955 |
| **BA (%)** | | 57.59 | |

**Table 3.** Pharmacokinetic parameters for **BGAz-003.**

| | **BGAz-003** | | |
|---|---|---|---|
| | **PO** | **IV** | **IP** |
| **Animal Number KM mice** | ♂4 | ♂4 | ♂4 |
| **Dose level mg/kg** | 5 | 1 | 1 |
| **AUC(0-∞) μg*h** | 1403.716 | 563.438 | 849.246 |
| **$T_{1/2}$ (h)** | 27.991 | 10.834 | 21.3 |
| **$T_{max}$ (h)** | 1.375 | 0.033 | 0.167 |
| **$C_{max}$（μg/L）** | 56.034 | 254.543 | 49.137 |
| **BA (%)** | | 49.83 | |

**Table 4.** Pharmacokinetic parameters for **BGAz-004.**

| | **BGAz-004** | | |
|---|---|---|---|
| | **PO** | **IV** | **IP** |
| **Animal Number KM mice** | ♂4 | ♂4 | ♂4 |
| **Dose level mg/kg** | 5 | 1 | 1 |
| **AUC(0-∞) μg*h** | 1406.796 | 789.43 | 581.359 |
| **$T_{1/2}$ (h)** | 11.368 | 8.289 | 9.823 |
| **$T_{max}$ (h)** | 0.875 | 0.033 | 0.271 |
| **$C_{max}$（μg/L）** | 87.649 | 278.395 | 61.021 |
| **BA (%)** | | 35.64 | |

**Table 5.** Pharmacokinetic parameters for **BGAz-005.**

| | BGAz-005 | | |
|---|---|---|---|
| | **PO** | **IV** | **IP** |
| **Animal Number KM mice** | ♂4 | ♂4 | ♂4 |
| **Dose level mg/kg** | 5 | 1 | 1 |
| **AUC(0-∞) μg*h** | 1542.381 | 424.729 | 464.151 |
| **$T_{1/2}$ (h)** | 35.733 | 8.052 | 9.339 |
| **$T_{max}$ (h)** | 6 | 0.033 | 0.646 |
| **$C_{max}$（μg/L）** | 43.127 | 320.487 | 41.897 |
| **BA (%)** | | 72.63 | |

#### 6.1 *Results for* BGAz-001

**Table 6.** Concentration of **BGAz-001** for PO (μg/L).

| Time (h) | No. 1 (μg/L) | No. 2 (μg/L) | No. 3 (μg/L) | No. 4 (μg/L) | Mean (μg/L) | SD |
|---|---|---|---|---|---|---|
| 0.083 | 144.18 | 20.10 | 6.67 | 30.82 | 50.44 | 63.27 |
| 0.25 | 29.52 | 1.90 | 4.72 | 6.73 | 10.72 | 12.69 |
| 0.5 | 31.49 | 4.96 | 13.11 | 9.48 | 14.76 | 11.64 |
| 1 | 22.70 | 10.80 | 23.73 | 17.83 | 18.76 | 5.90 |
| 3 | 15.54 | 9.49 | 18.32 | 19.20 | 15.64 | 4.39 |
| 5 | 13.54 | 6.93 | death | 12.38 | 10.95 | 3.53 |
| 8 | 9.03 | 9.67 | death | 7.54 | 8.75 | 1.09 |
| 24 | 0.00 | 0.00 | death | 0.00 | 0.00 | 0.00 |

**Table 7.** Concentration of **BGAz-001** for PO (μM).

| Time (h) | No. 1 (μM) | No. 2 (μM) | No. 3 (μM) | No. 4 (μM) | Mean (μg/L) | SD |
|---|---|---|---|---|---|---|
| 0.083 | 0.38 | 0.05 | 0.02 | 0.08 | 0.13 | 0.16 |
| 0.25 | 0.08 | 0.00 | 0.01 | 0.02 | 0.03 | 0.03 |
| 0.5 | 0.08 | 0.01 | 0.03 | 0.02 | 0.04 | 0.03 |
| 1 | 0.06 | 0.03 | 0.06 | 0.05 | 0.05 | 0.02 |
| 3 | 0.04 | 0.02 | 0.05 | 0.05 | 0.04 | 0.01 |
| 5 | 0.04 | 0.02 | death | 0.03 | 0.03 | 0.01 |
| 8 | 0.02 | 0.03 | death | 0.02 | 0.02 | 0.00 |
| 24 | 0.00 | 0.00 | death | 0.00 | 0.00 | 0.00 |

**Table 8.** Pharmacokinetic parameters of **BGAz-001** for PO.

| Parameter | Unit | No. 1 | No. 2 | No. 3 | No. 4 | Mean | SD | RSD/% |
|---|---|---|---|---|---|---|---|---|
| **AUC(0-t)** | µg* | 215.047 | 146.456 | 54.715 | 172.051 | 147.067 | 67.761 | 46.1 |
| **AUC(0-∞)** | µg* | 215.049 | 146.458 | 54.715 | 172.053 | 147.069 | 67.762 | 46.1 |
| **R_AUC(t/∞)** | % | 100 | 100 | 100 | 100 | 100 | 0 | 0 |
| **AUMC(0-t)** | **µg | 985.855 | 893.394 | 87.392 | 867.533 | 708.544 | 417.204 | 58.9 |
| **AUMC(0-∞)** | **µg | 985.912 | 893.454 | 87.392 | 867.588 | 708.587 | 417.233 | 58.9 |
| **MRT(0-t)** | | 4.584 | 6.1 | 1.597 | 5.042 | 4.331 | 1.93 | 44.6 |
| **MRT(0-∞)** | | 4.585 | 6.1 | 1.597 | 5.043 | 4.331 | 1.93 | 44.6 |
| **VRT(0-t)** | ^2 | 9.89 | 7.162 | 1.028 | 7.853 | 6.483 | 3.817 | 58.9 |
| **VRT(0-∞)** | ^2 | 9.894 | 7.168 | 1.028 | 7.858 | 6.487 | 3.819 | 58.9 |
| **λz** | 1/ | 0.476 | 0.448 | 0 | 0.48 | 0.351 | 0.234 | 66.7 |
| **λz 回归尾点** | | 134 | 134 | | 134 | -- | -- | -- |
| **C_last** | µg | 0.001 | 0.001 | 1 | 0.001 | 0.251 | 0.5 | 199.2 |
| **t1/2z** | | 1.456 | 1.547 | | 1.443 | 1.482 | 0.057 | 3.8 |
| **Tmax** | | 0.083 | 0.083 | 1 | 0.083 | 0.312 | 0.459 | 147.1 |
| **Vz/F** | L/kg | 48.84 | 76.229 | | 60.517 | 61.862 | 13.744 | 22.2 |
| **CLz/F** | L//kg | 23.251 | 34.139 | 91.383 | 29.061 | 44.459 | 31.598 | 71.1 |
| **Cmax** | µg | 144.1835178 | 20.09557627 | 23.73036304 | 30.82118547 | 54.708 | 59.817 | 109.3 |

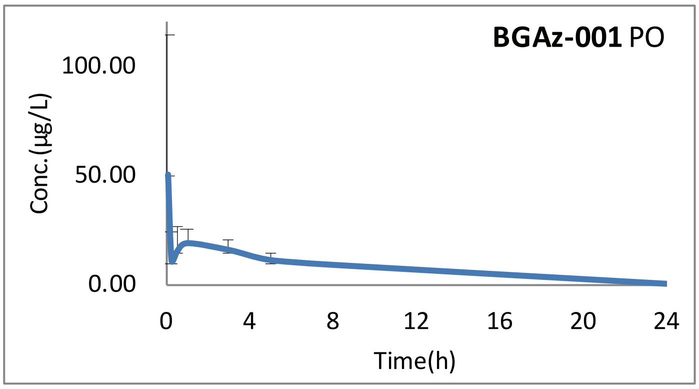

**Figure 1**

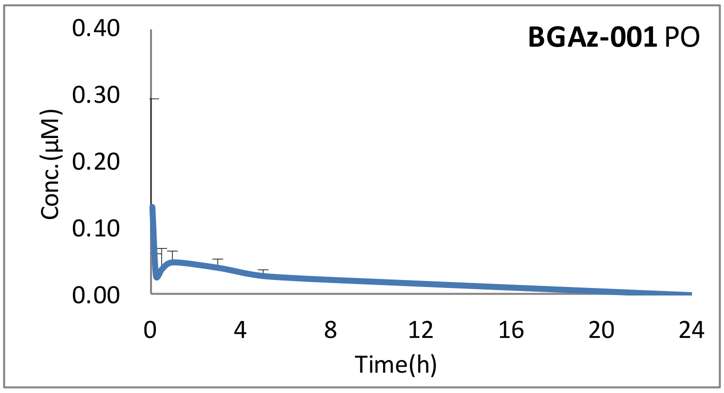

**Figure 2**

**Table 9.** Concentration of **BGAz-001** for IV (μg/L).

| Time (h) | No. 1 (μg/L) | No. 2 (μg/L) | No. 3 (μg/L) | No. 4 (μg/L) | Mean (μg/L) | SD |
|---|---|---|---|---|---|---|
| 0.033 | 180.84 | 161.14 | 222.24 | 66.28 | 157.63 | 66.00 |
| 0.167 | 85.06 | 102.64 | 124.42 | 54.81 | 91.73 | 29.41 |
| 0.5 | 35.31 | 53.80 | 58.88 | 25.46 | 43.36 | 15.65 |
| 1 | 21.78 | 66.59 | 47.23 | 18.05 | 38.41 | 22.82 |
| 3 | 4.22 | 13.07 | 12.54 | 5.18 | 8.75 | 4.70 |
| 5 | 0.79 | 15.12 | 0.00 | 0.00 | 3.98 | 7.44 |
| 8 | 0.00 | 5.14 | 0.00 | 0.00 | 1.29 | 2.57 |
| 24 | 0.00 | 0.00 | 0.00 | 0.00 | 0.00 | 0.00 |

**Table 10.** Concentration of **BGAz-001** for IV (μM).

| Time (h) | No. 1 (μM) | No. 2 (μM) | No. 3 (μM) | No. 4 (μM) | Mean (μg/L) | SD |
|---|---|---|---|---|---|---|
| 0.033 | 0.47 | 0.42 | 0.58 | 0.17 | 0.41 | 0.17 |
| 0.167 | 0.22 | 0.27 | 0.32 | 0.14 | 0.24 | 0.08 |
| 0.5 | 0.09 | 0.14 | 0.15 | 0.07 | 0.11 | 0.04 |
| 1 | 0.06 | 0.17 | 0.12 | 0.05 | 0.10 | 0.06 |
| 3 | 0.01 | 0.03 | 0.03 | 0.01 | 0.02 | 0.01 |
| 5 | 0.00 | 0.04 | 0.00 | 0.00 | 0.01 | 0.02 |
| 8 | 0.00 | 0.01 | 0.00 | 0.00 | 0.00 | 0.01 |
| 24 | 0.00 | 0.00 | 0.00 | 0.00 | 0.00 | 0.00 |

**Table 11.** Pharmacokinetic parameters of **BGAz-001** for IV.

| Parameter | Unit | No. 1 | No. 2 | No. 3 | No. 4 | Mean | SD | RSD/% |
|---|---|---|---|---|---|---|---|---|
| **AUC(0-t)** | μg* | 90.889 | 258.845 | 160.483 | 63.007 | 143.306 | 87.254 | 60.9 |
| **AUC(0-∞)** | μg* | 90.892 | 258.847 | 160.483 | 63.007 | 143.307 | 87.254 | 60.9 |
| **R_AUC(t/∞)** | % | 100 | 100 | 100 | 100 | 100 | 0 | 0 |
| **AUMC(0-t)** | **μg | 73.552 | 757.268 | 152.002 | 61.262 | 261.021 | 333.264 | 127.7 |
| **AUMC(0-∞)** | **μg | 73.619 | 757.321 | 152.002 | 61.262 | 261.051 | 333.278 | 127.7 |
| **MRT(0-t)** | | 0.809 | 2.926 | 0.947 | 0.972 | 1.414 | 1.011 | 71.5 |
| **MRT(0-∞)** | | 0.81 | 2.926 | 0.947 | 0.972 | 1.414 | 1.011 | 71.5 |
| **VRT(0-t)** | ^2 | 1.069 | 8.43 | 0.921 | 0.94 | 2.84 | 3.727 | 131.2 |
| **VRT(0-∞)** | ^2 | 1.088 | 8.434 | 0.921 | 0.94 | 2.846 | 3.726 | 130.9 |
| **λz** | 1/ | 0.379 | 0.516 | 1.738 | 1.575 | 1.052 | 0.703 | 66.8 |
| **λz 回归尾点** | | 134 | 123 | 234 | 234 | -- | -- | -- |
| **C_last** | μg | 0.001 | 0.001 | 0 | 0 | 0.001 | 0.001 | 100 |
| **t1/2z** | | 1.829 | 1.344 | 0.399 | 0.44 | 1.003 | 0.702 | 70 |
| **Tmax** | | 0.033 | 0.033 | 0.033 | 0.033 | 0.033 | 0 | 0 |
| **Vz/F** | L/kg | 29.038 | 7.491 | 3.585 | 10.074 | 12.547 | 11.313 | 90.2 |
| **CLz/F** | L//kg | 11.002 | 3.863 | 6.231 | 15.871 | 9.242 | 5.324 | 57.6 |
| **Cmax** | μg | 180.8432007 | 161.1415149 | 222.239702 | 66.28256575 | 157.627 | 66.005 | 41.9 |
| **C0** | μg | 217.759 | 180.074 | 256.369 | 69.46 | 180.916 | 80.568 | 44.5 |

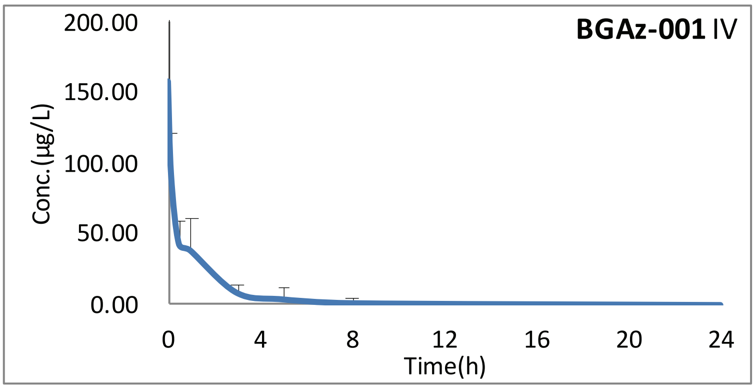

**Figure 3**

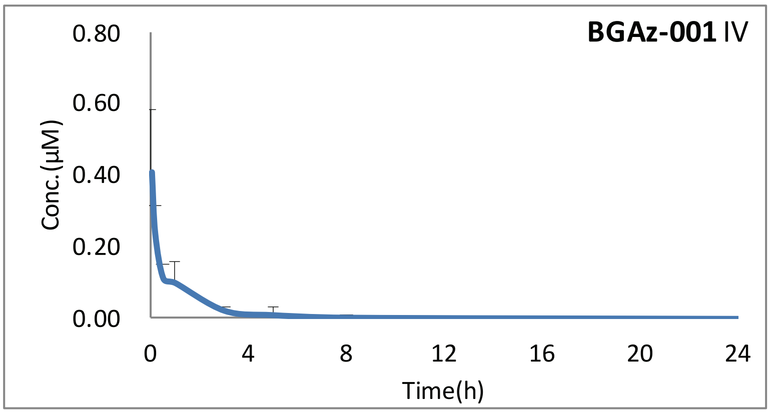

**Figure 4**

**Table 12.** Concentration of **BGAz-001** for IP (µg/L).

| Time (h) | No. 1 (µg/L) | No. 2 (µg/L) | No. 3 (µg/L) | No. 4 (µg/L) | Mean (µg/L) | SD |
|---|---|---|---|---|---|---|
| 0.083 | 36.79 | 16.68 | 11.71 | 12.46 | 19.41 | 11.79 |
| 0.25 | 17.42 | 8.00 | 9.69 | 22.50 | 14.40 | 6.78 |
| 0.5 | 12.01 | 5.99 | 9.68 | 12.89 | 10.14 | 3.08 |
| 1 | 3.14 | 3.25 | 7.60 | 7.17 | 5.29 | 2.43 |
| 3 | 0.36 | 0.00 | 4.54 | 1.45 | 1.59 | 2.06 |
| 5 | 0.00 | 0.00 | / | 0.00 | 0.00 | 0.00 |
| 8 | 0.00 | 0.39 | 0.00 | 0.00 | 0.10 | 0.19 |
| 24 | 0.00 | 0.00 | 0.00 | 0.00 | 0.00 | 0.00 |

**Table 13.** Concentration of **BGAz-001** for IP (µM).

| Time (h) | No. 1 (µM) | No. 2 (µM) | No. 3 (µM) | No. 4 (µM) | Mean (µg/L) | SD |
|---|---|---|---|---|---|---|
| 0.083 | 0.10 | 0.04 | 0.03 | 0.03 | 0.05 | 0.03 |
| 0.25 | 0.05 | 0.02 | 0.03 | 0.06 | 0.04 | 0.02 |
| 0.5 | 0.03 | 0.02 | 0.03 | 0.03 | 0.03 | 0.01 |
| 1 | 0.01 | 0.01 | 0.02 | 0.02 | 0.01 | 0.01 |
| 3 | 0.00 | 0.00 | 0.01 | 0.00 | 0.00 | 0.01 |
| 5 | 0.00 | 0.00 | / | 0.00 | 0.00 | 0.00 |
| 8 | 0.00 | 0.00 | 0.00 | 0.00 | 0.00 | 0.00 |
| 24 | 0.00 | 0.00 | 0.00 | 0.00 | 0.00 | 0.00 |

**Table 14.** Pharmacokinetic parameters of **BGAz-00x** for IP.

| Parameter | Unit | No. 1 | No. 2 | No. 3 | No. 4 | Mean | SD | RSD/% |
|---|---|---|---|---|---|---|---|---|
| **AUC(0-t)** | μg* | 17.377 | 13.769 | 32.506 | 22.953 | 21.651 | 8.163 | 37.7 |
| **AUC(0-∞)** | μg* | 17.377 | 13.783 | 32.509 | 22.953 | 21.656 | 8.16 | 37.7 |
| **R_AUC(t/∞)** | % | 100 | 99.9 | 100 | 100 | 99.975 | 0.05 | 0.1 |
| **AUMC(0-t)** | **μg | 9.629 | 35.382 | 59.605 | 21.387 | 31.501 | 21.491 | 68.2 |
| **AUMC(0-∞)** | **μg | 9.629 | 35.827 | 59.673 | 21.387 | 31.629 | 21.548 | 68.1 |
| **MRT(0-t)** | | 0.554 | 2.57 | 1.834 | 0.932 | 1.473 | 0.908 | 61.6 |
| **MRT(0-∞)** | | 0.554 | 2.599 | 1.836 | 0.932 | 1.48 | 0.92 | 62.2 |
| **VRT(0-t)** | ^2 | 0.373 | 10.94 | 1.361 | 0.725 | 3.35 | 5.077 | 151.6 |
| **VRT(0-∞)** | ^2 | 0.373 | 11.871 | 1.409 | 0.725 | 3.595 | 5.534 | 153.9 |
| **λz** | 1/ | 1.085 | 0.124 | 0.394 | 1.341 | 0.736 | 0.571 | 77.6 |
| **λz 回归尾点** | | 234 | 123 | 134 | 234 | -- | -- | -- |
| **C_last** | μg | 0 | 0.002 | 0.001 | 0 | 0.001 | 0.001 | 100 |
| **t1/2z** | | 0.639 | 5.595 | 1.761 | 0.517 | 2.128 | 2.378 | 111.7 |
| **Tmax** | | 0.083 | 0.083 | 0.083 | 0.25 | 0.125 | 0.084 | 67.2 |
| **Vz/F** | L/kg | 53.038 | 585.72 | 78.146 | 32.491 | 187.349 | 266.236 | 142.1 |
| **CLz/F** | L//kg | 57.547 | 72.553 | 30.761 | 43.568 | 51.107 | 18.002 | 35.2 |
| **Cmax** | μg | 36.79392484 | 16.67918911 | 11.7147724 | 22.5021479 | 21.923 | 10.85 | 49.5 |

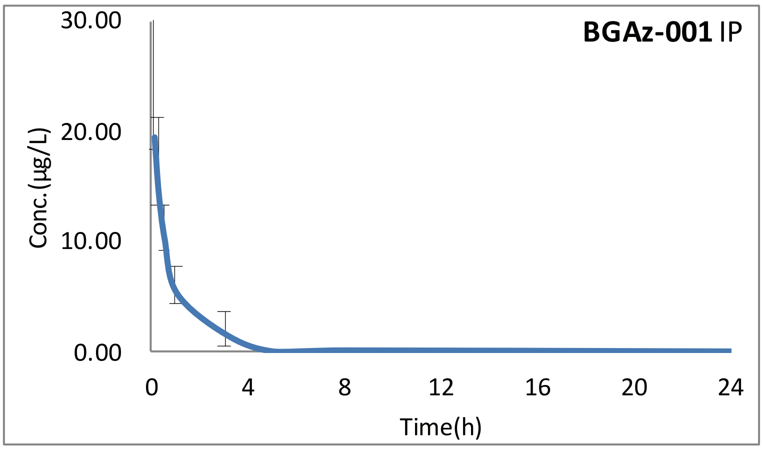

**Figure 5**

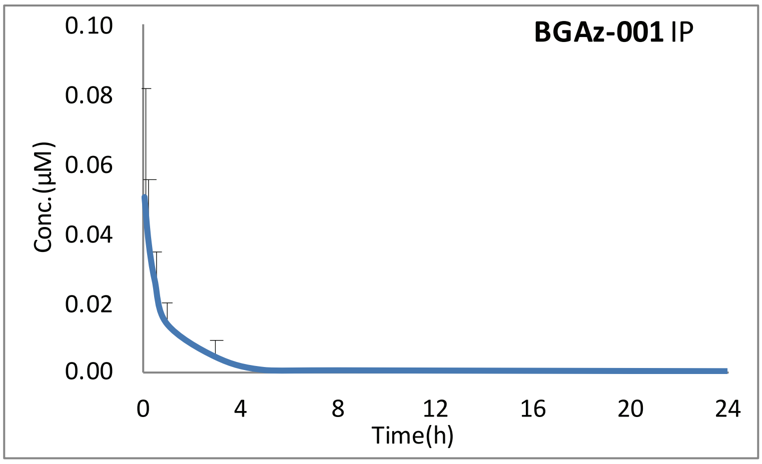

**Figure 6**

#### 6.2 ***Results for*** *BGAz-002*

**Table 15.** Concentration of **BGAz-002** for PO (μg/L).

| Time (h) | No. 1 (μg/L) | No. 2 (μg/L) | No. 3 (μg/L) | No. 4 (μg/L) | Mean (μg/L) | SD |
|---|---|---|---|---|---|---|
| 0.083 | 22.70 | 18.44 | 12.42 | 12.17 | 16.43 | 5.09 |
| 0.25 | 30.92 | 22.35 | 26.54 | 20.79 | 25.15 | 4.55 |
| 0.5 | 45.43 | 34.31 | 37.96 | 33.08 | 37.69 | 5.56 |
| 1 | 38.47 | 53.36 | 47.84 | 57.44 | 49.28 | 8.21 |
| 3 | 35.15 | 56.60 | 55.96 | 49.78 | 49.37 | 9.97 |
| 5 | 29.27 | 42.52 | death | 48.54 | 40.11 | 9.86 |
| 8 | 30.25 | 42.15 | death | 37.76 | 36.72 | 6.02 |
| 24 | 24.18 | 22.29 | death | 22.40 | 22.95 | 1.06 |

**Table 16.** Concentration of **BGAz-002** for PO (μM).

| Time (h) | No. 1 (μM) | No. 2 (μM) | No. 3 (μM) | No. 4 (μM) | Mean (μg/L) | SD |
|---|---|---|---|---|---|---|
| 0.083 | 0.05 | 0.04 | 0.03 | 0.03 | 0.03 | 0.01 |
| 0.25 | 0.07 | 0.05 | 0.06 | 0.04 | 0.05 | 0.01 |
| 0.5 | 0.10 | 0.07 | 0.08 | 0.07 | 0.08 | 0.01 |
| 1 | 0.08 | 0.11 | 0.10 | 0.12 | 0.10 | 0.02 |
| 3 | 0.07 | 0.12 | 0.12 | 0.10 | 0.10 | 0.02 |
| 5 | 0.06 | 0.09 | death | 0.10 | 0.08 | 0.02 |
| 8 | 0.06 | 0.09 | death | 0.08 | 0.08 | 0.01 |
| 24 | 0.05 | 0.05 | death | 0.05 | 0.05 | 0.00 |

**Table 17.** Pharmacokinetic parameters of **BGAz-002** for PO.

| Parameter | Unit | No. 1 | No. 2 | No. 3 | No. 4 | Mean | SD | RSD/% |
|---|---|---|---|---|---|---|---|---|
| **AUC(0-t)** | μg* | 698.688 | 884.816 | 137.083 | 848.886 | 642.368 | 346.37 | 53.9 |
| **AUC(0-∞)** | μg* | 2117.266 | 1394.629 | 137.083 | 1423.146 | 1268.031 | 824.688 | 65 |
| **R_AUC(t/∞)** | % | 33 | 63.4 | 100 | 59.6 | 64 | 27.548 | 43 |
| **AUMC(0-t)** | **μg | 7576.309 | 8429.064 | 236.316 | 8154.748 | 6099.109 | 3924.657 | 64.3 |
| **AUMC(0-∞)** | **μg | 125599.193 | 32448.81 | 236.316 | 36621.833 | 48726.538 | 53765.525 | 110.3 |
| **MRT(0-t)** | | 10.844 | 9.526 | 1.724 | 9.606 | 7.925 | 4.178 | 52.7 |
| **MRT(0-∞)** | | 59.321 | 23.267 | 1.724 | 25.733 | 27.511 | 23.791 | 86.5 |
| **VRT(0-t)** | ^2 | 71.792 | 58.534 | 1.167 | 61.059 | 48.138 | 31.837 | 66.1 |
| **VRT(0-∞)** | ^2 | 3529.144 | 560.135 | 1.167 | 684.723 | 1193.792 | 1585.023 | 132.8 |
| **λz** | 1/ | 0.017 | 0.043 | 0 | 0.039 | 0.025 | 0.02 | 80 |
| **λz 回归尾点** | | 124 | 124 | | 134 | -- | -- | -- |
| **C_last** | μg | 23.963 | 22.056 | 1 | 22.457 | 17.369 | 10.944 | 63 |
| **t1/2z** | | 41.024 | 16.019 | | 17.721 | 24.921 | 13.971 | 56.1 |
| **Tmax** | | 0.5 | 3 | 3 | 1 | 1.875 | 1.315 | 70.1 |
| **Vz/F** | L/kg | 139.798 | 82.871 | | 89.842 | 104.17 | 31.051 | 29.8 |
| **CLz/F** | L//kg | 2.362 | 3.585 | 36.474 | 3.513 | 11.484 | 16.67 | 145.2 |
| **Cmax** | μg | 45.4286889 | 56.59584787 | 55.96190526 | 57.44444395 | 53.858 | 5.652 | 10.5 |

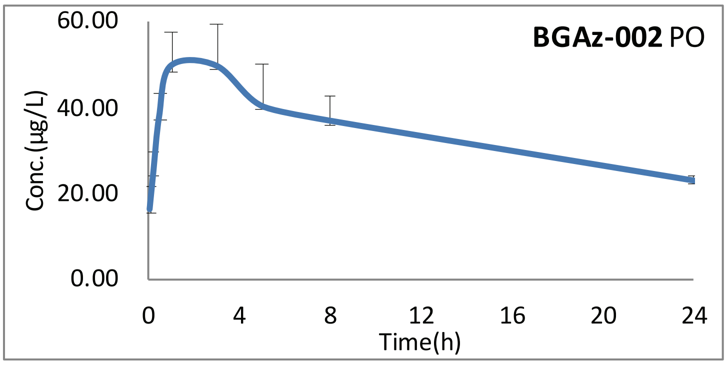

**Figure 7**

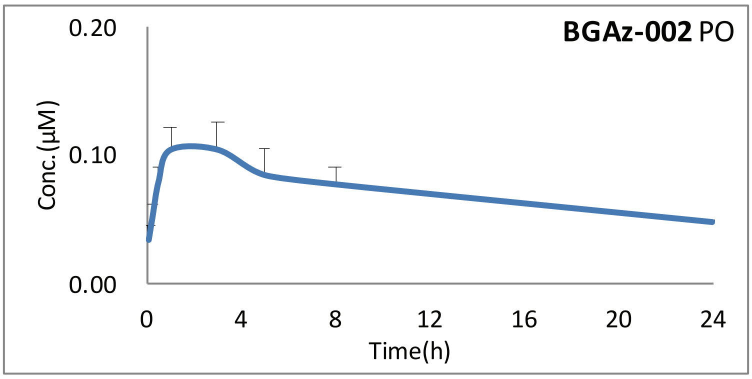

**Figure 8**

**Table 18.** Concentration of **BGAz-002** for IV (µg/L).

| Time (h) | No. 1 (µg/L) | No. 2 (µg/L) | No. 3 (µg/L) | No. 4 (µg/L) | Mean (µg/L) | SD |
|---|---|---|---|---|---|---|
| 0.033 | 220.68 | 204.88 | 235.95 | 93.24 | 188.69 | 64.89 |
| 0.167 | 113.21 | 103.79 | 107.54 | 71.07 | 98.90 | 18.95 |
| 0.5 | 63.20 | 73.71 | 74.67 | 38.53 | 62.53 | 16.82 |
| 1 | 43.14 | 54.66 | 50.75 | 33.90 | 45.61 | 9.16 |
| 3 | 26.15 | 32.42 | 30.42 | 24.06 | 28.26 | 3.83 |
| 5 | 14.69 | 24.54 | 18.33 | 15.70 | 18.32 | 4.42 |
| 8 | 10.73 | 13.38 | 13.37 | 12.48 | 12.49 | 1.25 |
| 24 | 4.80 | 12.59 | 3.91 | 3.33 | 6.16 | 4.33 |

**Table 19.** Concentration of **BGAz-002** for IV (µM).

| Time (h) | No. 1 (µM) | No. 2 (µM) | No. 3 (µM) | No. 4 (µM) | Mean (µg/L) | SD |
|---|---|---|---|---|---|---|
| 0.033 | 0.47 | 0.43 | 0.50 | 0.20 | 0.40 | 0.14 |
| 0.167 | 0.24 | 0.22 | 0.23 | 0.15 | 0.21 | 0.04 |
| 0.5 | 0.13 | 0.16 | 0.16 | 0.08 | 0.13 | 0.04 |
| 1 | 0.09 | 0.12 | 0.11 | 0.07 | 0.10 | 0.02 |
| 3 | 0.06 | 0.07 | 0.06 | 0.05 | 0.06 | 0.01 |
| 5 | 0.03 | 0.05 | 0.04 | 0.03 | 0.04 | 0.01 |
| 8 | 0.02 | 0.03 | 0.03 | 0.03 | 0.03 | 0.00 |
| 24 | 0.01 | 0.03 | 0.01 | 0.01 | 0.01 | 0.01 |

**Table 20.** Pharmacokinetic parameters of **BGAz-002** for IV.

| Parameter | Unit | No. 1 | No. 2 | No. 3 | No. 4 | Mean | SD | RSD/% |
|---|---|---|---|---|---|---|---|---|
| **AUC(0-t)** | μg* | 358.743 | 498.345 | 409.005 | 317.008 | 395.775 | 78.041 | 19.7 |
| **AUC(0-∞)** | μg* | 443.504 | 502.691 | 457.565 | 357.798 | 440.39 | 60.574 | 13.8 |
| **R_AUC(t/∞)** | % | 80.9 | 99.1 | 89.4 | 88.6 | 89.5 | 7.46 | 8.3 |
| **AUMC(0-t)** | **μg | 2150.014 | 4023.36 | 2261.527 | 1982.397 | 2604.325 | 952.954 | 36.6 |
| **AUMC(0-∞)** | **μg | 5698.919 | 4151.911 | 4034.703 | 3459.685 | 4336.305 | 957.452 | 22.1 |
| **MRT(0-t)** | | 5.993 | 8.073 | 5.529 | 6.253 | 6.462 | 1.115 | 17.3 |
| **MRT(0-∞)** | | 12.85 | 8.259 | 8.818 | 9.669 | 9.899 | 2.051 | 20.7 |
| **VRT(0-t)** | ^2 | 47.999 | 71.952 | 37.619 | 37.901 | 48.868 | 16.129 | 33 |
| **VRT(0-∞)** | ^2 | 298.828 | 75.566 | 141.33 | 141.277 | 164.25 | 94.92 | 57.8 |
| **λz** | 1/ | 0.056 | 0.179 | 0.08 | 0.082 | 0.099 | 0.054 | 54.5 |
| **λz 回归尾点** | | 123 | 234 | 123 | 123 | -- | -- | -- |
| **C_last** | μg | 4.743 | 0.778 | 3.88 | 3.339 | 3.185 | 1.706 | 53.6 |
| **t1/2z** | | 12.384 | 3.871 | 8.673 | 8.466 | 8.349 | 3.486 | 41.8 |
| **Tmax** | | 0.033 | 0.033 | 0.033 | 0.033 | 0.033 | 0 | 0 |
| **Vz/F** | L/kg | 40.291 | 11.111 | 27.351 | 34.144 | 28.224 | 12.573 | 44.5 |
| **CLz/F** | L//kg | 2.255 | 1.989 | 2.185 | 2.795 | 2.306 | 0.345 | 15 |
| **Cmax** | μg | 220.6847261 | 204.88032 | 235.9536049 | 93.24005526 | 188.69 | 64.885 | 34.4 |
| **C0** | μg | 260.116 | 242.233 | 286.328 | 99.688 | 222.091 | 83.587 | 37.6 |

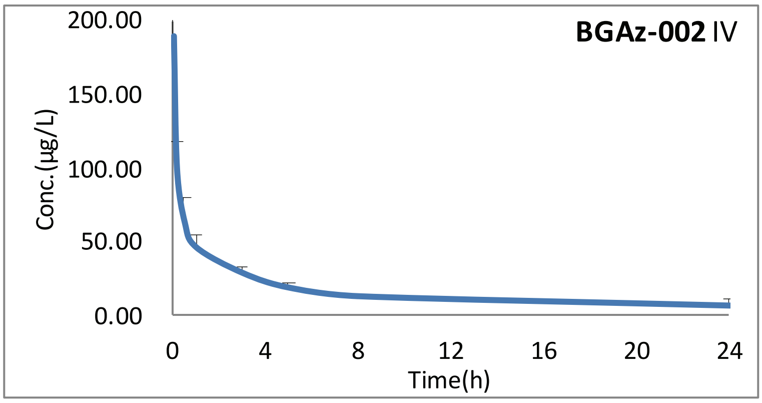

**Figure 9**

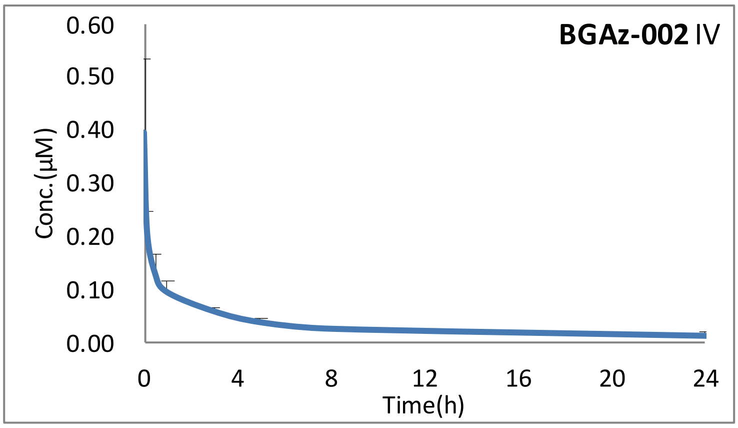

**Figure 10**

**Table 21.** Concentration of **BGAz-002** for IP (μg/L).

| Time (h) | No. 1 (μg/L) | No. 2 (μg/L) | No. 3 (μg/L) | No. 4 (μg/L) | Mean (μg/L) | SD |
|---|---|---|---|---|---|---|
| 0.083 | 52.67 | 33.27 | 27.07 | 36.75 | 37.44 | 10.91 |
| 0.25 | 54.59 | 28.96 | 26.71 | 44.89 | 38.79 | 13.29 |
| 0.5 | 41.62 | 25.98 | 25.65 | 37.33 | 32.65 | 8.08 |
| 1 | 30.90 | 22.07 | 21.44 | 28.77 | 25.79 | 4.75 |
| 3 | 29.46 | 15.73 | 25.12 | 25.84 | 24.04 | 5.85 |
| 5 | 15.26 | 13.45 | / | 22.15 | 16.96 | 4.59 |
| 8 | 19.65 | 11.65 | 11.32 | 17.23 | 14.96 | 4.14 |
| 24 | 15.40 | 14.27 | 10.09 | 9.93 | 12.42 | 2.82 |

**Table 22.** Concentration of **BGAz-002** for IP (μM).

| Time (h) | No. 1 (μM) | No. 2 (μM) | No. 3 (μM) | No. 4 (μM) | Mean (μg/L) | SD |
|---|---|---|---|---|---|---|
| 0.083 | 0.11 | 0.07 | 0.06 | 0.08 | 0.08 | 0.02 |
| 0.25 | 0.12 | 0.06 | 0.06 | 0.09 | 0.08 | 0.03 |
| 0.5 | 0.09 | 0.05 | 0.05 | 0.08 | 0.07 | 0.02 |
| 1 | 0.07 | 0.05 | 0.05 | 0.06 | 0.05 | 0.01 |
| 3 | 0.06 | 0.03 | 0.05 | 0.05 | 0.05 | 0.01 |
| 5 | 0.03 | 0.03 | / | 0.05 | 0.04 | 0.01 |
| 8 | 0.04 | 0.02 | 0.02 | 0.04 | 0.03 | 0.01 |
| 24 | 0.03 | 0.03 | 0.02 | 0.02 | 0.03 | 0.01 |

**Table 23.** Pharmacokinetic parameters of **BGAz-002** for IP.

| Parameter | Unit | No. 1 | No. 2 | No. 3 | No. 4 | Mean | SD | RSD/% |
|---|---|---|---|---|---|---|---|---|
| **AUC(0-t)** | μg* | 479.126 | 337.45 | 332.867 | 414.063 | 390.877 | 69.63 | 17.8 |
| **AUC(0-∞)** | μg* | 1027.494 | 413.093 | 607.489 | 471.851 | 629.982 | 277.229 | 44 |
| **R_AUC(t/∞)** | % | 46.6 | 81.7 | 54.8 | 87.8 | 67.725 | 20.097 | 29.7 |
| **AUMC(0-t)** | **μg | 4867.315 | 3922.348 | 3185.311 | 3693.675 | 3917.162 | 704.36 | 18 |
| **AUMC(0-∞)** | **μg | 38280.33 | 7016.073 | 17170.99 | 5791.21 | 17064.651 | 15035.3 | 88.1 |
| **MRT(0-t)** | | 10.159 | 11.623 | 9.569 | 8.921 | 10.068 | 1.153 | 11.5 |
| **MRT(0-∞)** | | 37.256 | 16.984 | 28.266 | 12.273 | 23.695 | 11.259 | 47.5 |
| **VRT(0-t)** | ^2 | 73.026 | 84.194 | 73.443 | 60.812 | 72.869 | 9.557 | 13.1 |
| **VRT(0-∞)** | ^2 | 1403.536 | 249.274 | 791.7 | 152.432 | 649.236 | 576.208 | 88.8 |
| **λz** | 1/ | 0.027 | 0.059 | 0.037 | 0.081 | 0.051 | 0.024 | 47.1 |
| **λz 回归尾点** | | 124 | 234 | 134 | 234 | -- | -- | -- |
| **C_last** | μg | 14.848 | 4.476 | 10.199 | 4.699 | 8.556 | 4.96 | 58 |
| **t1/2z** | | 25.594 | 11.711 | 18.66 | 8.522 | 16.122 | 7.602 | 47.2 |
| **Tmax** | | 0.25 | 0.083 | 0.083 | 0.25 | 0.167 | 0.096 | 57.5 |
| **Vz/F** | L/kg | 35.944 | 40.909 | 44.325 | 26.062 | 36.81 | 7.949 | 21.6 |
| **CLz/F** | L//kg | 0.973 | 2.421 | 1.646 | 2.119 | 1.79 | 0.631 | 35.3 |
| **Cmax** | μg | 54.59298631 | 33.26648936 | 27.07007106 | 44.88882257 | 39.955 | 12.239 | 30.6 |

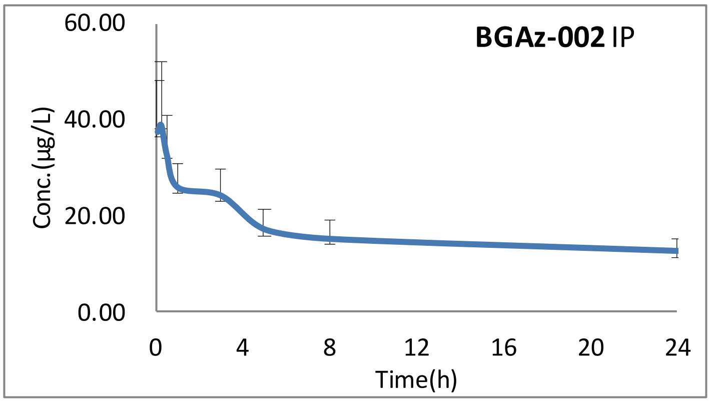

**Figure 11**

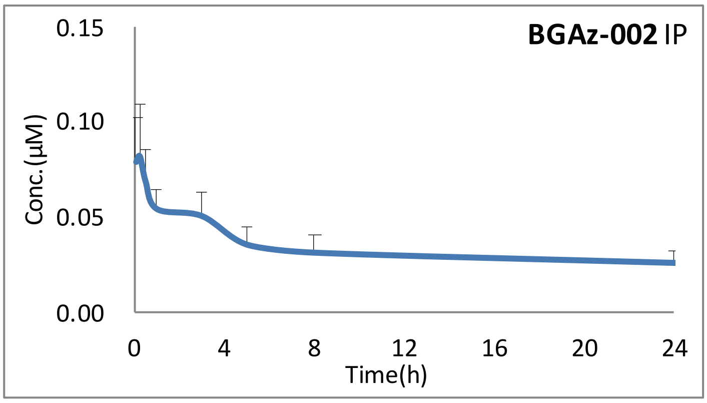

**Figure 12**

#### *6.3 Results for* *BGAz-003*

**Table 24.** Concentration of **BGAz-003** for PO (µg/L).

| Time (h) | No. 1 (µg/L) | No. 2 (µg/L) | No. 3 (µg/L) | No. 4 (µg/L) | Mean (µg/L) | SD |
|---|---|---|---|---|---|---|
| 0.083 | 15.47 | 17.39 | 11.81 | 10.58 | 13.81 | 3.17 |
| 0.25 | 34.32 | 27.35 | 30.32 | 23.35 | 28.83 | 4.64 |
| 0.5 | 51.22 | 39.33 | 41.56 | 36.72 | 42.21 | 6.32 |
| 1 | 43.03 | 59.13 | 53.16 | 59.58 | 53.72 | 7.71 |
| 3 | 36.20 | 60.18 | 50.03 | 50.81 | 49.30 | 9.88 |
| 5 | 30.57 | 43.70 | death | 45.91 | 40.06 | 8.29 |
| 8 | 31.31 | 40.51 | death | 35.15 | 35.65 | 4.62 |
| 24 | 24.56 | 25.92 | death | 23.91 | 24.80 | 1.03 |

**Table 25.** Concentration of **BGAz-003** for PO (µM).

| Time (h) | No. 1 (µM) | No. 2 (µM) | No. 3 (µM) | No. 4 (µM) | Mean (µg/L) | SD |
|---|---|---|---|---|---|---|
| 0.083 | 0.03 | 0.03 | 0.02 | 0.02 | 0.03 | 0.01 |
| 0.25 | 0.07 | 0.05 | 0.06 | 0.04 | 0.06 | 0.01 |
| 0.5 | 0.10 | 0.08 | 0.08 | 0.07 | 0.08 | 0.01 |
| 1 | 0.08 | 0.11 | 0.10 | 0.11 | 0.10 | 0.01 |
| 3 | 0.07 | 0.12 | 0.10 | 0.10 | 0.09 | 0.02 |
| 5 | 0.06 | 0.08 | death | 0.09 | 0.08 | 0.02 |
| 8 | 0.06 | 0.08 | death | 0.07 | 0.07 | 0.01 |
| 24 | 0.05 | 0.05 | death | 0.05 | 0.05 | 0.00 |

**Table 26.** Pharmacokinetic parameters of **BGAz-003** for PO.

| Parameter | Unit | No. 1 | No. 2 | No. 3 | No. 4 | Mean | SD | RSD/% |
|---|---|---|---|---|---|---|---|---|
| **AUC(0-t)** | μg* | 724.799 | 918.348 | 139.859 | 835.99 | 654.749 | 352.303 | 53.8 |
| **AUC(0-∞)** | μg* | 2104.452 | 1857.679 | 139.859 | 1512.875 | 1403.716 | 876.805 | 62.5 |
| **R_AUC(t/∞)** | % | 34.4 | 49.4 | 100 | 55.3 | 59.775 | 28.223 | 47.2 |
| **AUMC(0-t)** | **μg | 7759.603 | 9046.21 | 226.023 | 8223.172 | 6313.752 | 4093.211 | 64.8 |
| **AUMC(0-∞)** | **μg | 118968.77 | 65604.625 | 226.023 | 43660.783 | 57115.05 | 49380.611 | 86.5 |
| **MRT(0-t)** | | 10.706 | 9.851 | 1.616 | 9.836 | 8.002 | 4.277 | 53.4 |
| **MRT(0-∞)** | | 56.532 | 35.315 | 1.616 | 28.859 | 30.581 | 22.641 | 74 |
| **VRT(0-t)** | ^2 | 71.411 | 64.105 | 1.113 | 65.216 | 50.461 | 33.055 | 65.5 |
| **VRT(0-∞)** | ^2 | 3228.55 | 1328.702 | 1.113 | 842.672 | 1350.259 | 1367.026 | 101.2 |
| **λz** | 1/ | 0.018 | 0.028 | 0 | 0.035 | 0.02 | 0.015 | 75 |
| **λz 回归尾点** | | 124 | 123 | | 134 | -- | -- | -- |
| **C_last** | μg | 24.373 | 25.94 | 1 | 23.873 | 18.797 | 11.897 | 63.3 |
| **t1/2z** | | 39.228 | 25.095 | | 19.649 | 27.991 | 10.106 | 36.1 |
| **Tmax** | | 0.5 | 3 | 1 | 1 | 1.375 | 1.109 | 80.7 |
| **Vz/F** | L/kg | 134.493 | 97.464 | | 93.709 | 108.555 | 22.541 | 20.8 |
| **CLz/F** | L//kg | 2.376 | 2.692 | 35.75 | 3.305 | 11.031 | 16.484 | 149.4 |
| **Cmax** | μg | 51.21708015 | 60.17758845 | 53.16155932 | 59.58024983 | 56.034 | 4.517 | 8.1 |

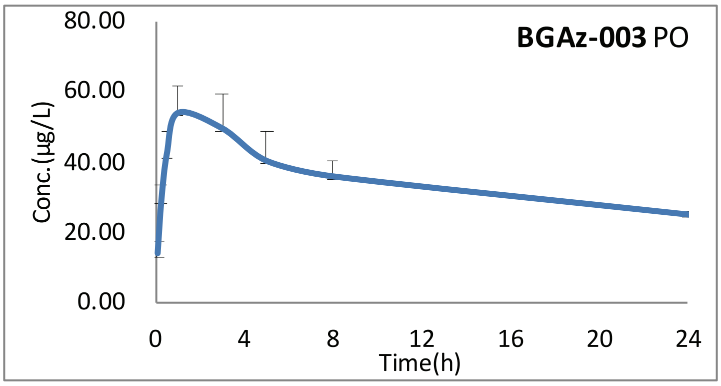

**Figure 13**

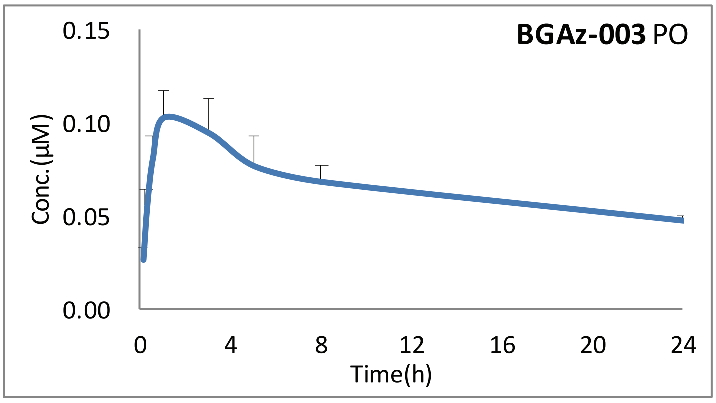

**Figure 14**

**Table 27.** Concentration of **BGAz-003** for IV (μg/L).

| Time (h) | No. 1 (μg/L) | No. 2 (μg/L) | No. 3 (μg/L) | No. 4 (μg/L) | Mean (μg/L) | SD |
|---|---|---|---|---|---|---|
| 0.033 | 296.33 | 275.38 | 314.33 | 132.14 | 254.54 | 83.14 |
| 0.167 | 141.72 | 127.01 | 126.87 | 87.93 | 120.88 | 23.05 |
| 0.5 | 73.46 | 85.29 | 87.09 | 44.34 | 72.54 | 19.75 |
| 1 | 50.44 | 67.89 | 58.02 | 39.82 | 54.04 | 11.87 |
| 3 | 28.70 | 37.64 | 35.03 | 27.22 | 32.15 | 4.99 |
| 5 | 16.34 | 27.05 | 21.69 | 18.20 | 20.82 | 4.71 |
| 8 | 12.34 | 15.90 | 15.08 | 13.80 | 14.28 | 1.56 |
| 24 | 7.04 | 15.48 | 6.10 | 5.63 | 8.56 | 4.65 |

**Table 28.** Concentration of **BGAz-003** for IV (μM).

| Time (h) | No. 1 (μM) | No. 2 (μM) | No. 3 (μM) | No. 4 (μM) | Mean (μg/L) | SD |
|---|---|---|---|---|---|---|
| 0.033 | 0.57 | 0.53 | 0.60 | 0.25 | 0.49 | 0.16 |
| 0.167 | 0.27 | 0.24 | 0.24 | 0.17 | 0.23 | 0.04 |
| 0.5 | 0.14 | 0.16 | 0.17 | 0.09 | 0.14 | 0.04 |
| 1 | 0.10 | 0.13 | 0.11 | 0.08 | 0.10 | 0.02 |
| 3 | 0.06 | 0.07 | 0.07 | 0.05 | 0.06 | 0.01 |
| 5 | 0.03 | 0.05 | 0.04 | 0.03 | 0.04 | 0.01 |
| 8 | 0.02 | 0.03 | 0.03 | 0.03 | 0.03 | 0.00 |
| 24 | 0.01 | 0.03 | 0.01 | 0.01 | 0.02 | 0.01 |

**Table 29.** Pharmacokinetic parameters of **BGAz-003** for IV.

| Parameter | Unit | No. 1 | No. 2 | No. 3 | No. 4 | Mean | SD | RSD/% |
|---|---|---|---|---|---|---|---|---|
| **AUC(0-t)** | μg* | 429.177 | 596.372 | 487.47 | 378.272 | 472.823 | 93.673 | 19.8 |
| **AUC(0-∞)** | μg* | 597.789 | 602.205 | 582.291 | 471.465 | 563.438 | 61.907 | 11 |
| **R_AUC(t/∞)** | % | 71.8 | 99 | 83.7 | 80.2 | 83.675 | 11.372 | 13.6 |
| **AUMC(0-t)** | **μg | 2751.396 | 4853.432 | 2894.712 | 2582.813 | 3270.588 | 1062.901 | 32.5 |
| **AUMC(0-∞)** | **μg | 10889.033 | 5027.179 | 6666.617 | 6375.757 | 7239.647 | 2535.593 | 35 |
| **MRT(0-t)** | | 6.411 | 8.138 | 5.938 | 6.828 | 6.829 | 0.946 | 13.9 |
| **MRT(0-∞)** | | 18.216 | 8.348 | 11.449 | 13.523 | 12.884 | 4.142 | 32.1 |
| **VRT(0-t)** | ^2 | 55.76 | 73.735 | 45.442 | 48.554 | 55.873 | 12.668 | 22.7 |
| **VRT(0-∞)** | ^2 | 560.765 | 77.842 | 234.705 | 276.043 | 287.339 | 201.286 | 70.1 |
| **λz** | 1/ | 0.041 | 0.173 | 0.063 | 0.06 | 0.084 | 0.06 | 71.4 |
| **λz 回归尾点** | | 123 | 234 | 123 | 123 | -- | -- | -- |
| **C_last** | μg | 6.95 | 1.007 | 6.009 | 5.58 | 4.887 | 2.649 | 54.2 |
| **t1/2z** | | 16.814 | 4.013 | 10.935 | 11.573 | 10.834 | 5.255 | 48.5 |
| **Tmax** | | 0.033 | 0.033 | 0.033 | 0.033 | 0.033 | 0 | 0 |
| **Vz/F** | L/kg | 40.587 | 9.616 | 27.098 | 35.422 | 28.181 | 13.567 | 48.1 |
| **CLz/F** | L//kg | 1.673 | 1.661 | 1.717 | 2.121 | 1.793 | 0.22 | 12.3 |
| **Cmax** | μg | 296.3267152 | 275.3785806 | 314.3285331 | 132.1385403 | 254.543 | 83.141 | 32.7 |
| **C0** | μg | 355.355 | 333.194 | 393.026 | 146.08 | 306.914 | 110.03 | 35.9 |

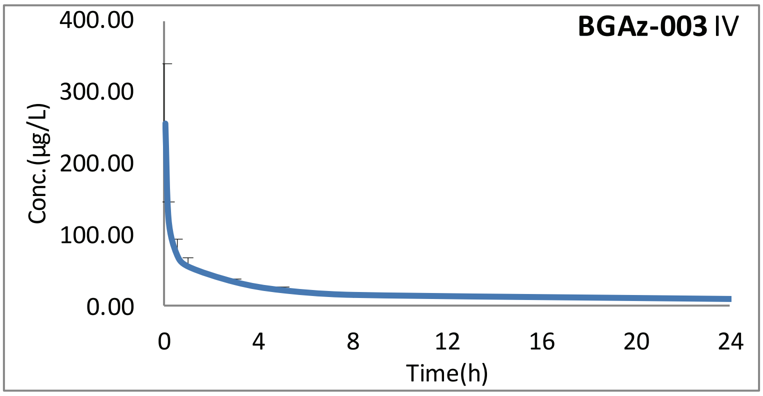

**Figure 15**

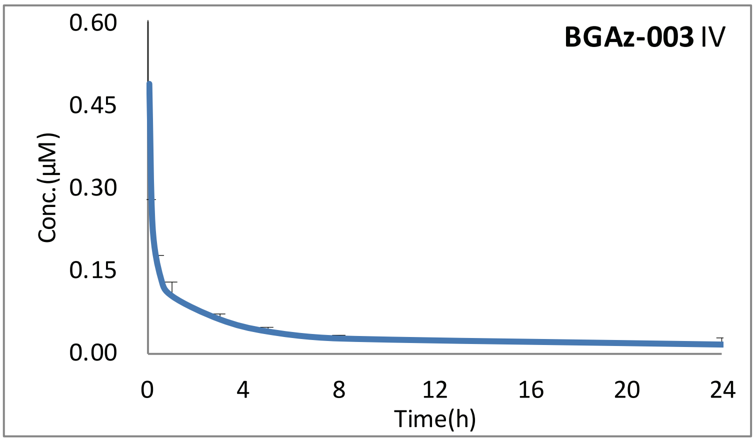

**Figure 16**

**Table 30.** Concentration of **BGAz-003** for IP (μg/L).

| Time (h) | No. 1 (μg/L) | No. 2 (μg/L) | No. 3 (μg/L) | No. 4 (μg/L) | Mean (μg/L) | SD |
|---|---|---|---|---|---|---|
| 0.083 | 59.71 | 39.66 | 35.20 | 43.85 | 44.61 | 10.67 |
| 0.25 | 66.33 | 35.29 | 33.21 | 55.35 | 47.55 | 16.02 |
| 0.5 | 48.44 | 29.41 | 29.71 | 47.54 | 38.78 | 10.65 |
| 1 | 31.43 | 22.57 | 22.02 | 31.88 | 26.97 | 5.41 |
| 3 | 29.94 | 16.81 | 24.92 | 26.99 | 24.66 | 5.63 |
| 5 | 16.39 | 14.81 | / | 23.47 | 18.22 | 4.61 |
| 8 | 20.87 | 12.79 | 13.23 | 18.01 | 16.23 | 3.90 |
| 24 | 19.42 | 16.41 | 11.70 | 12.44 | 14.99 | 3.60 |

**Table 31.** Concentration of **BGAz-003** for IP (μM).

| Time (h) | No. 1 (μM) | No. 2 (μM) | No. 3 (μM) | No. 4 (μM) | Mean (μg/L) | SD |
|---|---|---|---|---|---|---|
| 0.083 | 0.11 | 0.08 | 0.07 | 0.08 | 0.09 | 0.02 |
| 0.25 | 0.13 | 0.07 | 0.06 | 0.11 | 0.09 | 0.03 |
| 0.5 | 0.09 | 0.06 | 0.06 | 0.09 | 0.07 | 0.02 |
| 1 | 0.06 | 0.04 | 0.04 | 0.06 | 0.05 | 0.01 |
| 3 | 0.06 | 0.03 | 0.05 | 0.05 | 0.05 | 0.01 |
| 5 | 0.03 | 0.03 | / | 0.05 | 0.04 | 0.01 |
| 8 | 0.04 | 0.02 | 0.03 | 0.03 | 0.03 | 0.01 |
| 24 | 0.04 | 0.03 | 0.02 | 0.02 | 0.03 | 0.01 |

**Table 32.** Pharmacokinetic parameters of **BGAz-003** for IP.

| Parameter | Unit | No. 1 | No. 2 | No. 3 | No. 4 | Mean | SD | RSD/% |
|---|---|---|---|---|---|---|---|---|
| **AUC(0-t)** | μg* | 533.268 | 374.944 | 369.663 | 457.978 | 433.963 | 77.58 | 17.9 |
| **AUC(0-∞)** | μg* | 1654.392 | 473.609 | 750.59 | 518.391 | 849.246 | 550.322 | 64.8 |
| **R_AUC(t/∞)** | % | 32.2 | 79.2 | 49.2 | 88.3 | 62.225 | 26.072 | 41.9 |
| **AUMC(0-t)** | **μg | 5752.205 | 4443.772 | 3654.106 | 4264.861 | 4528.736 | 882.931 | 19.5 |
| **AUMC(0-∞)** | **μg | 99849.438 | 8631.896 | 25094.826 | 6456.42 | 35008.145 | 44021.111 | 125.7 |
| **MRT(0-t)** | | 10.787 | 11.852 | 9.885 | 9.312 | 10.459 | 1.11 | 10.6 |
| **MRT(0-∞)** | | 60.354 | 18.226 | 33.433 | 12.455 | 31.117 | 21.406 | 68.8 |
| **VRT(0-t)** | ^2 | 78.281 | 85.251 | 74.376 | 66.726 | 76.159 | 7.731 | 10.2 |
| **VRT(0-∞)** | ^2 | 3627.883 | 292.778 | 1103.766 | 151.371 | 1293.95 | 1611.546 | 124.5 |
| **λz** | 1/ | 0.017 | 0.054 | 0.031 | 0.081 | 0.046 | 0.028 | 60.9 |
| **λz 回归尾点** | | 124 | 234 | 134 | 234 | -- | -- | -- |
| **C_last** | μg | 18.707 | 5.348 | 11.799 | 4.921 | 10.194 | 6.489 | 63.7 |
| **t1/2z** | | 41.532 | 12.784 | 22.374 | 8.508 | 21.3 | 14.682 | 68.9 |
| **Tmax** | | 0.25 | 0.083 | 0.083 | 0.25 | 0.167 | 0.096 | 57.5 |
| **Vz/F** | L/kg | 36.225 | 38.952 | 43.014 | 23.682 | 35.468 | 8.338 | 23.5 |
| **CLz/F** | L//kg | 0.604 | 2.111 | 1.332 | 1.929 | 1.494 | 0.68 | 45.5 |
| **Cmax** | μg | 66.33443614 | 39.65805608 | 35.20115477 | 55.3538559 | 49.137 | 14.358 | 29.2 |

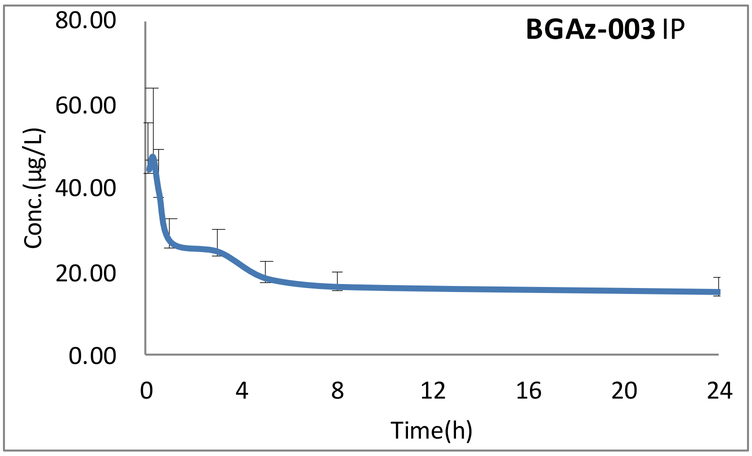

**Figure 17**

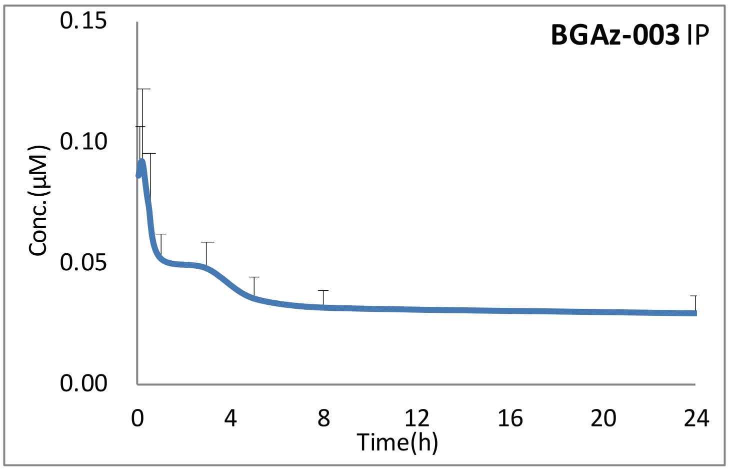

**Figure 18**

#### 6.4 ***Results for** BGAz-004*

**Table 33.** Concentration of **BGAz-00X** for PO (μg/L).

| Time (h) | No. 1 (μg/L) | No. 2 (μg/L) | No. 3 (μg/L) | No. 4 (μg/L) | Mean (μg/L) | SD |
|---|---|---|---|---|---|---|
| 0.0833 | 18.93 | 20.62 | 17.03 | 12.67 | 17.31 | 3.42 |
| 0.25 | 47.52 | 41.91 | 46.72 | 34.36 | 42.63 | 6.05 |
| 0.5 | 71.61 | 60.84 | 70.38 | 57.60 | 65.11 | 6.94 |
| 1 | 58.47 | 88.31 | 96.10 | 94.58 | 84.36 | 17.59 |
| 3 | 53.58 | 79.62 | 60.20 | 80.36 | 68.44 | 13.61 |
| 5 | 55.17 | 62.93 | death | 71.81 | 63.31 | 8.33 |
| 8 | 53.73 | 69.78 | death | 57.99 | 60.50 | 8.32 |
| 24 | 23.54 | 17.68 | death | 25.27 | 22.16 | 3.98 |

**Table 34.** Concentration of BGAz-004 for PO (μM).

| Time (h) | No. 1 (μM) | No. 2 (μM) | No. 3 (μM) | No. 4 (μM) | Mean (μg/L) | SD |
|---|---|---|---|---|---|---|
| 0.083 | 0.04 | 0.04 | 0.03 | 0.02 | 0.03 | 0.01 |
| 0.25 | 0.09 | 0.08 | 0.09 | 0.06 | 0.08 | 0.01 |
| 0.5 | 0.13 | 0.11 | 0.13 | 0.11 | 0.12 | 0.01 |
| 1 | 0.11 | 0.16 | 0.18 | 0.18 | 0.16 | 0.03 |
| 3 | 0.10 | 0.15 | 0.11 | 0.15 | 0.13 | 0.03 |
| 5 | 0.10 | 0.12 | death | 0.13 | 0.12 | 0.02 |
| 8 | 0.10 | 0.13 | death | 0.11 | 0.11 | 0.02 |
| 24 | 0.04 | 0.03 | death | 0.05 | 0.04 | 0.01 |

**Table 35.** Pharmacokinetic parameters of **BGAz-004** for PO.

| Parameter | Unit | No. 1 | No. 2 | No. 3 | No. 4 | Mean | SD | RSD/% |
|---|---|---|---|---|---|---|---|---|
| **AUC(0-t)** | μg* | 1056.005 | 1265.482 | 218.572 | 1241.851 | 945.478 | 493.575 | 52.2 |
| **AUC(0-∞)** | μg* | 1627.205 | 1517.952 | 781.063 | 1700.962 | 1406.796 | 423.875 | 30.1 |
| **R_AUC(t/∞)** | % | 64.9 | 83.4 | 28 | 73 | 62.325 | 24.104 | 38.7 |
| **AUMC(0-t)** | **μg | 9702.775 | 10087.367 | 316.518 | 10769.797 | 7719.114 | 4954.749 | 64.2 |
| **AUMC(0-∞)** | **μg | 37195.654 | 19768.43 | 7049.909 | 30130.148 | 23536.035 | 13115.572 | 55.7 |
| **MRT(0-t)** | | 9.188 | 7.971 | 1.448 | 8.672 | 6.82 | 3.616 | 53 |
| **MRT(0-∞)** | | 22.859 | 13.023 | 9.026 | 17.714 | 15.656 | 5.972 | 38.1 |
| **VRT(0-t)** | ^2 | 53.474 | 38.713 | 0.964 | 51.831 | 36.246 | 24.431 | 67.4 |
| **VRT(0-∞)** | ^2 | 584.618 | 194.427 | 80.537 | 348.053 | 301.909 | 218.03 | 72.2 |
| **λz** | 1/ | 0.041 | 0.07 | 0.111 | 0.055 | 0.069 | 0.03 | 43.5 |
| **λz 回归尾点** | | 134 | 134 | 123 | 134 | -- | -- | -- |
| **C_last** | μg | 23.67 | 17.6 | 62.703 | 25.269 | 32.311 | 20.529 | 63.5 |
| **t1/2z** | | 16.723 | 9.941 | 6.217 | 12.591 | 11.368 | 4.425 | 38.9 |
| **Tmax** | | 0.5 | 1 | 1 | 1 | 0.875 | 0.25 | 28.6 |
| **Vz/F** | L/kg | 74.151 | 47.252 | 57.426 | 53.409 | 58.06 | 11.515 | 19.8 |
| **CLz/F** | L//kg | 3.073 | 3.294 | 6.402 | 2.94 | 3.927 | 1.656 | 42.2 |
| **Cmax** | μg | 71.61139092 | 88.30540751 | 96.09674961 | 94.58256818 | 87.649 | 11.211 | 12.8 |

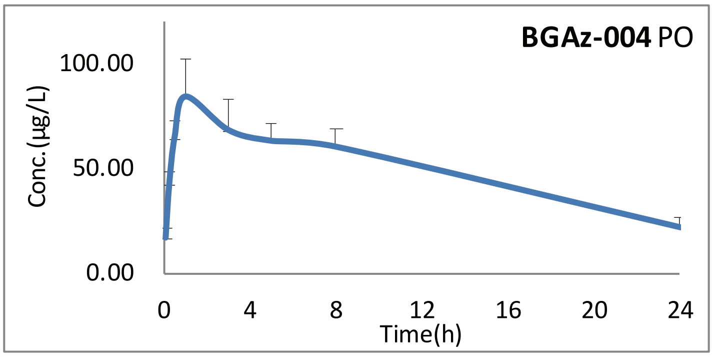

**Figure 19**

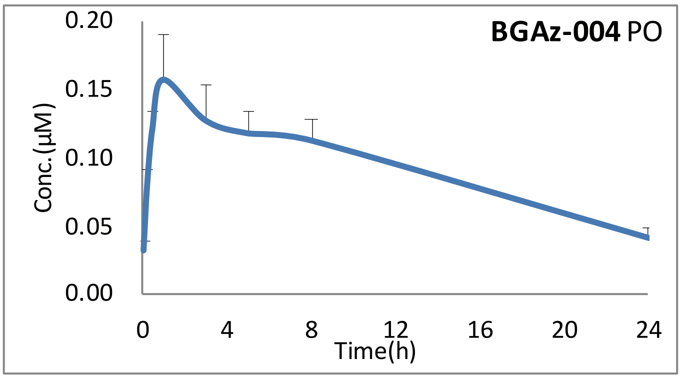

**Figure 20**

**Table 36.** Concentration of **BGAz-004** for IV (μg/L).

| Time (h) | No. 1 (μg/L) | No. 2 (μg/L) | No. 3 (μg/L) | No. 4 (μg/L) | Mean (μg/L) | SD |
|---|---|---|---|---|---|---|
| 0.033 | 312.46 | 299.55 | 348.67 | 152.89 | 278.39 | 86.21 |
| 0.167 | 154.96 | 167.65 | 170.71 | 97.46 | 147.69 | 34.18 |
| 0.5 | 76.49 | 105.49 | 101.34 | 55.23 | 84.64 | 23.42 |
| 1 | 69.82 | 153.90 | 87.34 | 47.23 | 89.57 | 45.92 |
| 3 | 34.42 | 53.91 | 43.69 | 30.09 | 40.53 | 10.57 |
| 5 | 20.54 | 44.93 | 26.66 | 21.15 | 28.32 | 11.41 |
| 8 | 13.46 | 40.83 | 17.31 | 14.90 | 21.62 | 12.90 |
| 24 | 2.96 | 50.95 | 3.09 | 3.23 | 15.06 | 23.93 |

**Table 37.** Concentration of **BGAz-004** for IV (μM).

| Time (h) | No. 1 (μM) | No. 2 (μM) | No. 3 (μM) | No. 4 (μM) | Mean (μg/L) | SD |
|---|---|---|---|---|---|---|
| 0.033 | 0.58 | 0.55 | 0.65 | 0.28 | 0.52 | 0.16 |
| 0.167 | 0.29 | 0.31 | 0.32 | 0.18 | 0.27 | 0.06 |
| 0.5 | 0.14 | 0.20 | 0.19 | 0.10 | 0.16 | 0.04 |
| 1 | 0.13 | 0.29 | 0.16 | 0.09 | 0.17 | 0.09 |
| 3 | 0.06 | 0.10 | 0.08 | 0.06 | 0.08 | 0.02 |
| 5 | 0.04 | 0.08 | 0.05 | 0.04 | 0.05 | 0.02 |
| 8 | 0.02 | 0.08 | 0.03 | 0.03 | 0.04 | 0.02 |
| 24 | 0.01 | 0.09 | 0.01 | 0.01 | 0.03 | 0.04 |

**Table 38.** Pharmacokinetic parameters of **BGAz-004** for IV.

| Parameter | Unit | No. 1 | No. 2 | No. 3 | No. 4 | Mean | SD | RSD/% |
|---|---|---|---|---|---|---|---|---|
| AUC(0-t) | μg* | 459.31 | 1321.748 | 570.433 | 400.795 | 688.072 | 428.27 | 62.2 |
| AUC(0-∞) | μg* | 488.727 | 1637.476 | 597.917 | 433.6 | 789.43 | 569.472 | 72.1 |
| R_AUC(t/∞) | % | 94 | 80.7 | 95.4 | 92.4 | 90.625 | 6.729 | 7.4 |
| AUMC(0-t) | **μg | 2164.698 | 13991.502 | 2642.241 | 2271.474 | 5267.479 | 5819.615 | 110.5 |
| AUMC(0-∞) | **μg | 3166.206 | 27445.659 | 3548.323 | 3394.024 | 9388.553 | 12039.094 | 128.2 |
| MRT(0-t) | | 4.713 | 10.586 | 4.632 | 5.667 | 6.4 | 2.83 | 44.2 |
| MRT(0-∞) | | 6.478 | 16.761 | 5.934 | 7.828 | 9.25 | 5.07 | 54.8 |
| VRT(0-t) | ^2 | 29.652 | 87.357 | 26.472 | 32.386 | 43.967 | 29.028 | 66 |
| VRT(0-∞) | ^2 | 82.613 | 296.959 | 64.162 | 94.844 | 134.645 | 108.942 | 80.9 |
| λz | 1/ | 0.1 | 0.054 | 0.112 | 0.098 | 0.091 | 0.025 | 27.5 |
| λz 回归尾点 | | 123 | 234 | 123 | 123 | -- | -- | -- |
| C_last | μg | 2.928 | 16.963 | 3.065 | 3.21 | 6.542 | 6.949 | 106.2 |
| t1/2z | | 6.962 | 12.899 | 6.215 | 7.081 | 8.289 | 3.097 | 37.4 |
| Tmax | | 0.033333333 | 0.033333333 | 0.033333333 | 0.033333333 | 0.033 | 0 | 0 |
| Vz/F | L/kg | 20.555 | 11.367 | 14.998 | 23.567 | 17.622 | 5.476 | 31.1 |
| CLz/F | L//kg | 2.046 | 0.611 | 1.672 | 2.306 | 1.659 | 0.745 | 44.9 |
| Cmax | μg | 312.4616133 | 299.5545268 | 348.6708335 | 152.8928136 | 278.395 | 86.212 | 31 |
| C0 | μg | 372.341 | 346.335 | 416.829 | 171.112 | 326.654 | 107.703 | 33 |

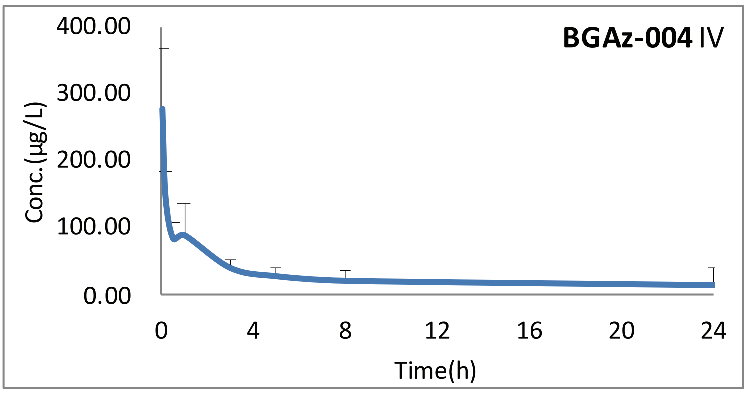

**Figure 21**

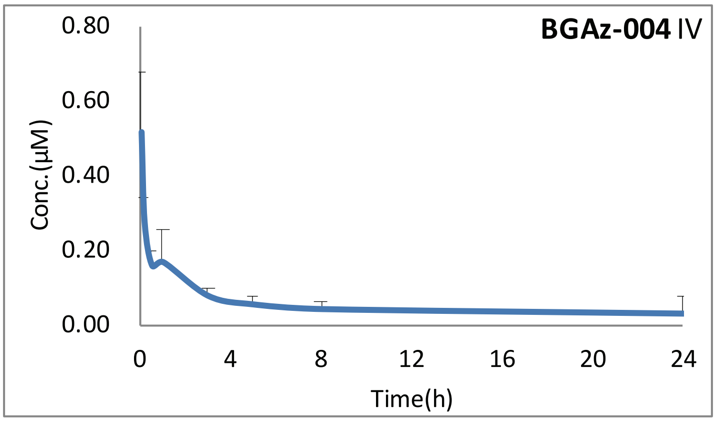

**Figure 22**

**Table 39.** Concentration of **BGAz-004** for IP (μg/L).

| Time (h) | No. 1 (μg/L) | No. 2 (μg/L) | No. 3 (μg/L) | No. 4 (μg/L) | Mean (μg/L) | SD |
|---|---|---|---|---|---|---|
| 0.083 | 65.88 | 48.41 | 45.50 | 61.29 | 55.27 | 9.85 |
| 0.25 | 71.52 | 46.22 | 45.43 | 77.07 | 60.06 | 16.59 |
| 0.5 | 52.15 | 41.72 | 47.09 | 60.20 | 50.29 | 7.86 |
| 1 | 37.03 | 38.73 | 37.78 | 44.09 | 39.41 | 3.20 |
| 3 | 31.92 | 28.16 | 33.48 | 30.35 | 30.98 | 2.27 |
| 5 | 21.17 | 27.36 | / | 24.54 | 24.36 | 3.10 |
| 8 | 24.39 | 22.52 | 23.22 | 17.09 | 21.81 | 3.24 |
| 24 | 7.45 | 8.48 | 7.51 | 5.18 | 7.15 | 1.40 |

**Table 40.** Concentration of **BGAz-004** for IP (μM).

| Time (h) | No. 1 (μM) | No. 2 (μM) | No. 3 (μM) | No. 4 (μM) | Mean (μg/L) | SD |
|---|---|---|---|---|---|---|
| 0.083 | 0.12 | 0.09 | 0.08 | 0.11 | 0.10 | 0.02 |
| 0.25 | 0.13 | 0.09 | 0.08 | 0.14 | 0.11 | 0.03 |
| 0.5 | 0.10 | 0.08 | 0.09 | 0.11 | 0.09 | 0.01 |
| 1 | 0.07 | 0.07 | 0.07 | 0.08 | 0.07 | 0.01 |
| 3 | 0.06 | 0.05 | 0.06 | 0.06 | 0.06 | 0.00 |
| 5 | 0.04 | 0.05 | / | 0.05 | 0.05 | 0.01 |
| 8 | 0.05 | 0.04 | 0.04 | 0.03 | 0.04 | 0.01 |
| 24 | 0.01 | 0.02 | 0.01 | 0.01 | 0.01 | 0.00 |

**Table 41.** Pharmacokinetic parameters of **BGAz-004** for IP.

| Parameter | Unit | No. 1 | No. 2 | No. 3 | No. 4 | Mean | SD | RSD/% |
|---|---|---|---|---|---|---|---|---|
| **AUC(0-t)** | μg* | 497.01 | 486.239 | 501.186 | 427.24 | 477.919 | 34.368 | 7.2 |
| **AUC(0-∞)** | μg* | 603.895 | 624.016 | 608.203 | 489.32 | 581.359 | 61.966 | 10.7 |
| **R_AUC(t/∞)** | % | 82.3 | 77.9 | 82.4 | 87.3 | 82.475 | 3.84 | 4.7 |
| **AUMC(0-t)** | **μg | 3800.137 | 3909.467 | 3803.862 | 2853.255 | 3591.68 | 494.886 | 13.8 |
| **AUMC(0-∞)** | **μg | 7882.021 | 9457.334 | 7895.112 | 5088.798 | 7580.816 | 1818.514 | 24 |
| **MRT(0-t)** | | 7.646 | 8.04 | 7.59 | 6.678 | 7.489 | 0.576 | 7.7 |
| **MRT(0-∞)** | | 13.052 | 15.156 | 12.981 | 10.4 | 12.897 | 1.947 | 15.1 |
| **VRT(0-t)** | ^2 | 44.353 | 48.522 | 44.823 | 40.606 | 44.576 | 3.237 | 7.3 |
| **VRT(0-∞)** | ^2 | 208.032 | 274.912 | 208.691 | 149.064 | 210.175 | 51.421 | 24.5 |
| **λz** | 1/ | 0.07 | 0.061 | 0.07 | 0.083 | 0.071 | 0.009 | 12.7 |
| **λz 回归尾点** | | 124 | 123 | 124 | 134 | -- | -- | -- |
| **C_last** | μg | 7.533 | 8.47 | 7.521 | 5.169 | 7.173 | 1.408 | 19.6 |
| **t1/2z** | | 9.833 | 11.273 | 9.861 | 8.323 | 9.823 | 1.205 | 12.3 |
| **Tmax** | | 0.25 | 0.083333333 | 0.5 | 0.25 | 0.271 | 0.172 | 63.5 |
| **Vz/F** | L/kg | 23.497 | 26.068 | 23.397 | 24.545 | 24.377 | 1.241 | 5.1 |
| **CLz/F** | L//kg | 1.656 | 1.603 | 1.644 | 2.044 | 1.737 | 0.206 | 11.9 |
| **Cmax** | μg | 71.52326579 | 48.41040301 | 47.08688711 | 77.06535763 | 61.021 | 15.502 | 25.4 |

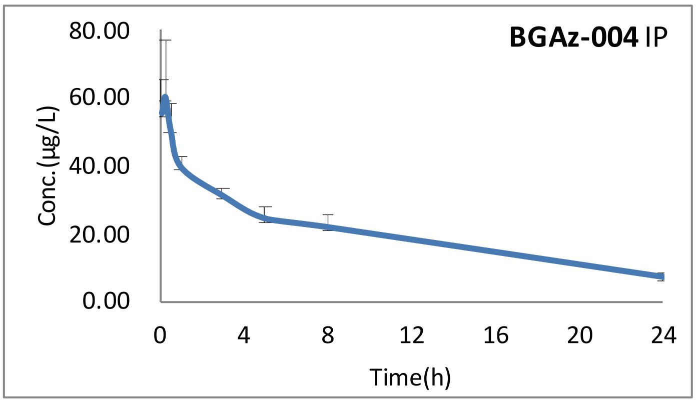

**Figure 23**

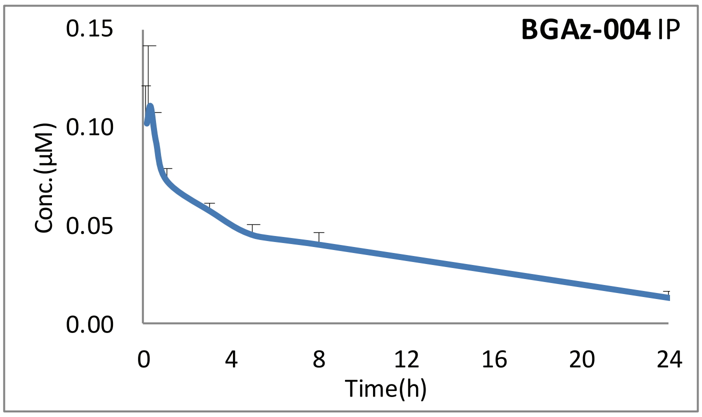

**Figure 24**

#### *6.5 Results for BGAz-005*

**Table 42.** Concentration of **BGAz-005** for PO (μg/L).

| Time (h) | No. 1 (μg/L) | No. 2 (μg/L) | No. 3 (μg/L) | No. 4 (μg/L) | Mean (μg/L) | SD |
|---|---|---|---|---|---|---|
| 0.083 | 11.85 | 9.55 | 3.03 | 3.87 | 7.08 | 4.30 |
| 0.25 | 15.24 | 8.96 | 11.40 | 9.61 | 11.30 | 2.82 |
| 0.5 | 26.93 | 17.11 | 19.69 | 18.05 | 20.45 | 4.45 |
| 1 | 24.17 | 35.42 | 31.56 | 39.03 | 32.55 | 6.36 |
| 3 | 25.26 | 46.01 | 46.93 | 38.34 | 39.13 | 10.02 |
| 5 | 22.52 | 43.17 | death | 47.25 | 37.65 | 13.26 |
| 8 | 28.02 | 50.31 | death | 42.42 | 40.25 | 11.30 |
| 24 | 18.13 | 31.32 | death | 23.47 | 24.31 | 6.63 |

**Table 43.** Concentration of **BGAz-005** for PO (μM).

| Time (h) | No. 1 (μM) | No. 2 (μM) | No. 3 (μM) | No. 4 (μM) | Mean (μg/L) | SD |
|---|---|---|---|---|---|---|
| 0.083 | 0.03 | 0.02 | 0.01 | 0.01 | 0.02 | 0.01 |
| 0.25 | 0.04 | 0.02 | 0.03 | 0.02 | 0.03 | 0.01 |
| 0.5 | 0.06 | 0.04 | 0.05 | 0.04 | 0.05 | 0.01 |
| 1 | 0.06 | 0.08 | 0.07 | 0.09 | 0.07 | 0.01 |
| 3 | 0.06 | 0.11 | 0.11 | 0.09 | 0.09 | 0.02 |
| 5 | 0.05 | 0.10 | death | 0.11 | 0.09 | 0.03 |
| 8 | 0.06 | 0.12 | death | 0.10 | 0.09 | 0.03 |
| 24 | 0.04 | 0.07 | death | 0.05 | 0.06 | 0.02 |

**Table 44.** Pharmacokinetic parameters of **BGAz-005** for PO.

| Parameter | Unit | No. 1 | No. 2 | No. 3 | No. 4 | Mean | SD | RSD/% |
|---|---|---|---|---|---|---|---|---|
| **AUC(0-t)** | µg* | 563.054 | 982.143 | 96.511 | 843.549 | 621.314 | 390.896 | 62.9 |
| **AUC(0-∞)** | µg* | 1848.857 | 2744.193 | 96.511 | 1479.962 | 1542.381 | 1100.411 | 71.3 |
| **R_AUC(t/∞)** | % | 30.5 | 35.8 | 100 | 57 | 55.825 | 31.597 | 56.6 |
| **AUMC(0-t)** | **µg | 6079.921 | 10699.775 | 184.544 | 8603.037 | 6391.819 | 4548.828 | 71.2 |
| **AUMC(0-∞)** | **µg | 128529.145 | 152271.287 | 184.544 | 41130.722 | 80528.925 | 71783.066 | 89.1 |
| **MRT(0-t)** | | 10.798 | 10.894 | 1.912 | 10.199 | 8.451 | 4.37 | 51.7 |
| **MRT(0-∞)** | | 69.518 | 55.489 | 1.912 | 27.792 | 38.678 | 30.022 | 77.6 |
| **VRT(0-t)** | ^2 | 65.399 | 63.023 | 1.149 | 59.142 | 47.178 | 30.794 | 65.3 |
| **VRT(0-∞)** | ^2 | 5058.543 | 3169.501 | 1.149 | 760.034 | 2247.307 | 2310.173 | 102.8 |
| **λz** | 1/ | 0.014 | 0.018 | 0 | 0.037 | 0.017 | 0.015 | 88.2 |
| **λz 回归尾点** | | 134 | 134 | | 123 | -- | -- | -- |
| **C_last** | µg | 18.051 | 31.273 | 1 | 23.474 | 18.45 | 12.837 | 69.6 |
| **t1/2z** | | 49.364 | 39.047 | | 18.788 | 35.733 | 15.555 | 43.5 |
| **Tmax** | | 8 | 8 | 3 | 5 | 6 | 2.449 | 40.8 |
| **Vz/F** | L/kg | 192.637 | 102.662 | | 91.593 | 128.964 | 55.419 | 43 |
| **CLz/F** | L//kg | 2.704 | 1.822 | 51.808 | 3.378 | 14.928 | 24.595 | 164.8 |
| **Cmax** | µg | 28.02378861 | 50.30584346 | 46.92633289 | 47.25369321 | 43.127 | 10.183 | 23.6 |

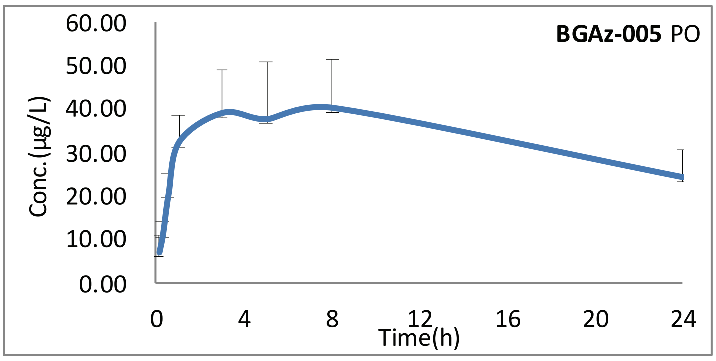

**Figure 25**

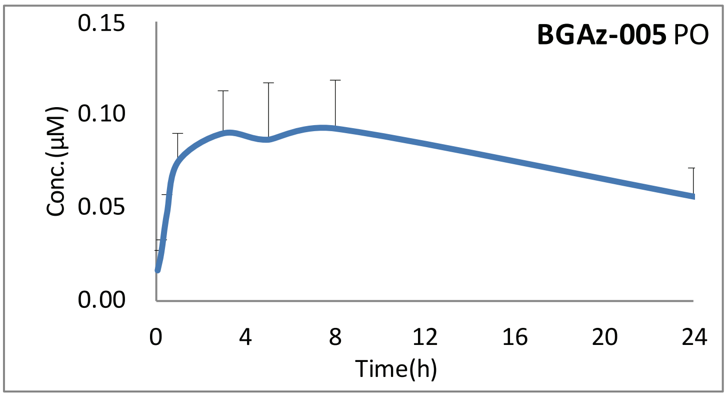

**Figure 26**

**Table 45.** Concentration of **BGAz-005** for IV (µg/L).

| Time (h) | No. 1 (µg/L) | No. 2 (µg/L) | No. 3 (µg/L) | No. 4 (µg/L) | Mean (µg/L) | SD |
|---|---|---|---|---|---|---|
| 0.033 | 333.20 | 356.84 | 403.52 | 188.39 | 320.49 | 92.78 |
| 0.167 | 144.37 | 160.92 | 156.16 | 119.07 | 145.13 | 18.71 |
| 0.5 | 63.00 | 81.74 | 78.03 | 78.03 | 75.20 | 8.32 |
| 1 | 38.37 | 56.47 | 52.87 | 52.87 | 50.15 | 8.03 |
| 3 | 20.15 | 28.18 | 26.64 | 26.64 | 25.40 | 3.58 |
| 5 | 12.02 | 21.68 | 15.37 | 15.37 | 16.11 | 4.03 |
| 8 | 8.24 | 12.87 | 10.08 | 10.08 | 10.32 | 1.91 |
| 24 | 3.70 | 11.57 | 2.76 | 2.76 | 5.20 | 4.27 |

**Table 46.** Concentration of **BGAz-005** for IV (µM).

| Time (h) | No. 1 (µM) | No. 2 (µM) | No. 3 (µM) | No. 4 (µM) | Mean (µg/L) | SD |
|---|---|---|---|---|---|---|
| 0.033 | 0.77 | 0.82 | 0.93 | 0.43 | 0.74 | 0.21 |
| 0.167 | 0.33 | 0.37 | 0.36 | 0.27 | 0.33 | 0.04 |
| 0.5 | 0.15 | 0.19 | 0.18 | 0.18 | 0.17 | 0.02 |
| 1 | 0.09 | 0.13 | 0.12 | 0.12 | 0.12 | 0.02 |
| 3 | 0.05 | 0.06 | 0.06 | 0.06 | 0.06 | 0.01 |
| 5 | 0.03 | 0.05 | 0.04 | 0.04 | 0.04 | 0.01 |
| 8 | 0.02 | 0.03 | 0.02 | 0.02 | 0.02 | 0.00 |
| 24 | 0.01 | 0.03 | 0.01 | 0.01 | 0.01 | 0.01 |

**Table 47.** Pharmacokinetic parameters of **BGAz-005** for IV.

| Parameter | Unit | No. 1 | No. 2 | No. 3 | No. 4 | Mean | SD | RSD/% |
|---|---|---|---|---|---|---|---|---|
| **AUC(0-t)** | µg* | 320.753 | 504.628 | 386.76 | 355.171 | 391.828 | 79.885 | 20.4 |
| **AUC(0-∞)** | µg* | 383.317 | 511.214 | 417.987 | 386.399 | 424.729 | 59.747 | 14.1 |
| **R_AUC(t/∞)** | % | 83.7 | 98.7 | 92.5 | 91.9 | 91.7 | 6.156 | 6.7 |
| **AUMC(0-t)** | **µg | 1675.205 | 3735.11 | 1737.886 | 1735.846 | 2221.012 | 1009.818 | 45.5 |
| **AUMC(0-∞)** | **µg | 4253.915 | 3934.839 | 2845.377 | 2843.337 | 3469.367 | 733.361 | 21.1 |
| **MRT(0-t)** | | 5.223 | 7.402 | 4.493 | 4.887 | 5.501 | 1.302 | 23.7 |
| **MRT(0-∞)** | | 11.098 | 7.697 | 6.807 | 7.359 | 8.24 | 1.94 | 23.5 |
| **VRT(0-t)** | ^2 | 45.113 | 70.24 | 32.475 | 33.463 | 45.323 | 17.575 | 38.8 |
| **VRT(0-∞)** | ^2 | 263.087 | 76.534 | 106.18 | 110.84 | 139.16 | 84.003 | 60.4 |
| **λz** | 1/ | 0.058 | 0.158 | 0.087 | 0.087 | 0.098 | 0.043 | 43.9 |
| **λz 回归尾点** | | 123 | 234 | 123 | 123 | -- | -- | -- |
| **C_last** | µg | 3.634 | 1.041 | 2.724 | 2.724 | 2.531 | 1.082 | 42.7 |
| **t1/2z** | | 11.932 | 4.385 | 7.945 | 7.945 | 8.052 | 3.084 | 38.3 |
| **Tmax** | | 0.033333333 | 0.033333333 | 0.033333333 | 0.033333333 | 0.033 | 0 | 0 |
| **Vz/F** | L/kg | 44.917 | 12.377 | 27.43 | 29.672 | 28.599 | 13.316 | 46.6 |
| **CLz/F** | L//kg | 2.609 | 1.956 | 2.392 | 2.588 | 2.386 | 0.303 | 12.7 |
| **Cmax** | µg | 333.1965586 | 356.8355463 | 403.522902 | 188.3926665 | 320.487 | 92.784 | 29 |
| **C0** | µg | 410.685 | 435.447 | 511.611 | 211.29 | 392.258 | 128.062 | 32.6 |

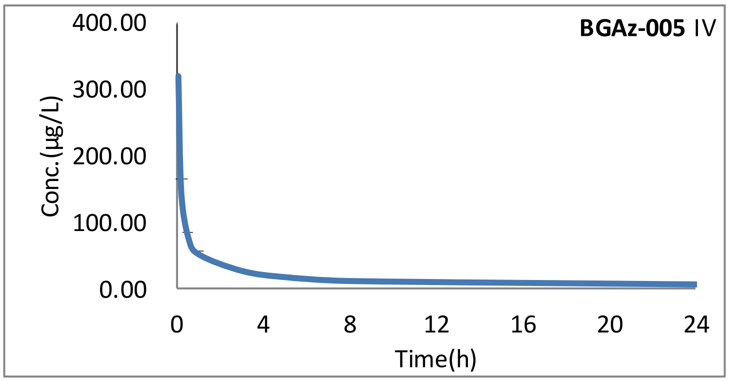

**Figure 27**

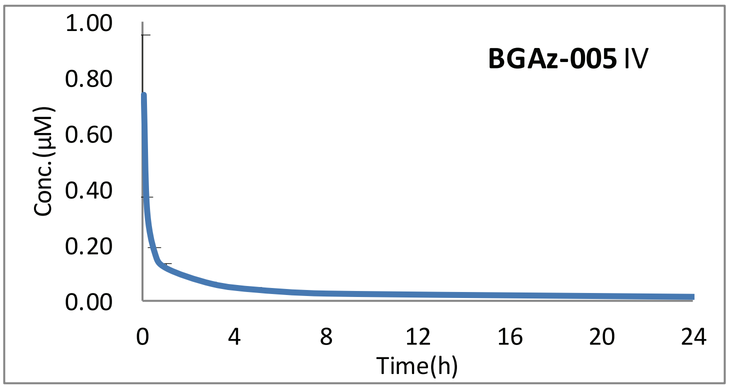

**Figure 28**

**Table 48.** Concentration of **BGAz-005** for IP (µg/L).

| Time (h) | No. 1 (µg/L) | No. 2 (µg/L) | No. 3 (µg/L) | No. 4 (µg/L) | Mean (µg/L) | SD |
|---|---|---|---|---|---|---|
| 0.083 | 20.53 | 14.50 | 19.62 | 53.21 | 26.96 | 17.70 |
| 0.25 | 39.41 | 20.97 | 31.29 | 27.59 | 29.82 | 7.69 |
| 0.5 | 42.67 | 28.16 | 40.95 | 33.32 | 36.27 | 6.76 |
| 1 | 43.88 | 29.55 | 39.49 | 46.14 | 39.76 | 7.35 |
| 3 | 34.52 | 22.40 | 39.02 | 37.48 | 33.35 | 7.54 |
| 5 | 18.30 | 19.40 | / | 26.18 | 21.29 | 4.27 |
| 8 | 16.77 | 15.15 | 12.75 | 17.08 | 15.44 | 1.03 |
| 24 | 5.58 | 6.67 | 5.89 | 3.64 | 5.44 | 1.53 |

**Table 49.** Concentration of **BGAz-005** for IP (µM).

| Time (h) | No. 1 (µM) | No. 2 (µM) | No. 3 (µM) | No. 4 (µM) | Mean (µg/L) | SD |
|---|---|---|---|---|---|---|
| 0.083 | 0.05 | 0.03 | 0.05 | 0.12 | 0.06 | 0.04 |
| 0.25 | 0.09 | 0.05 | 0.07 | 0.06 | 0.07 | 0.02 |
| 0.5 | 0.10 | 0.06 | 0.09 | 0.08 | 0.08 | 0.02 |
| 1 | 0.10 | 0.07 | 0.09 | 0.11 | 0.09 | 0.02 |
| 3 | 0.08 | 0.05 | 0.09 | 0.09 | 0.08 | 0.02 |
| 5 | 0.04 | 0.04 | / | 0.06 | 0.05 | 0.01 |
| 8 | 0.04 | 0.03 | 0.03 | 0.04 | 0.04 | 0.00 |
| 24 | 0.01 | 0.02 | 0.01 | 0.01 | 0.01 | 0.00 |

**Table 50.** Pharmacokinetic parameters of **BGAz-005** for IP.

| Parameter | Unit | No. 1 | No. 2 | No. 3 | No. 4 | Mean | SD | RSD/% |
|---|---|---|---|---|---|---|---|---|
| **AUC(0-t)** | μg* | 400.345 | 344.276 | 391.262 | 414.319 | 387.551 | 30.368 | 7.8 |
| **AUC(0-∞)** | μg* | 487.462 | 460.9 | 460.52 | 447.721 | 464.151 | 16.704 | 3.6 |
| **R_AUC(t/∞)** | % | 82.1 | 74.7 | 85 | 92.5 | 83.575 | 7.363 | 8.8 |
| **AUMC(0-t)** | **μg | 2846.397 | 2852.293 | 2670.84 | 2614.567 | 2746.024 | 121.52 | 4.4 |
| **AUMC(0-∞)** | **μg | 6285.492 | 7693.972 | 5141.16 | 3725.026 | 5711.413 | 1686.32 | 29.5 |
| **MRT(0-t)** | | 7.11 | 8.285 | 6.826 | 6.311 | 7.133 | 0.836 | 11.7 |
| **MRT(0-∞)** | | 12.894 | 16.693 | 11.164 | 8.32 | 12.268 | 3.501 | 28.5 |
| **VRT(0-t)** | ^2 | 43.674 | 52.184 | 47.976 | 31.434 | 43.817 | 8.957 | 20.4 |
| **VRT(0-∞)** | ^2 | 232.444 | 325.316 | 167.527 | 85.551 | 202.71 | 101.457 | 50.1 |
| **λz** | 1/ | 0.065 | 0.057 | 0.086 | 0.108 | 0.079 | 0.023 | 29.1 |
| **λz 回归尾点** | | 123 | 134 | 134 | 134 | -- | -- | -- |
| **C_last** | μg | 5.629 | 6.659 | 5.936 | 3.613 | 5.459 | 1.304 | 23.9 |
| **t1/2z** | | 10.726 | 12.138 | 8.086 | 6.407 | 9.339 | 2.577 | 27.6 |
| **Tmax** | | 1 | 1 | 0.5 | 0.083333333 | 0.646 | 0.443 | 68.6 |
| **Vz/F** | L/kg | 31.75 | 38.002 | 25.337 | 20.649 | 28.935 | 7.566 | 26.1 |
| **CLz/F** | L//kg | 2.051 | 2.17 | 2.171 | 2.234 | 2.157 | 0.076 | 3.5 |
| **Cmax** | μg | 43.87528447 | 29.55363799 | 40.94640377 | 53.21341977 | 41.897 | 9.751 | 23.3 |

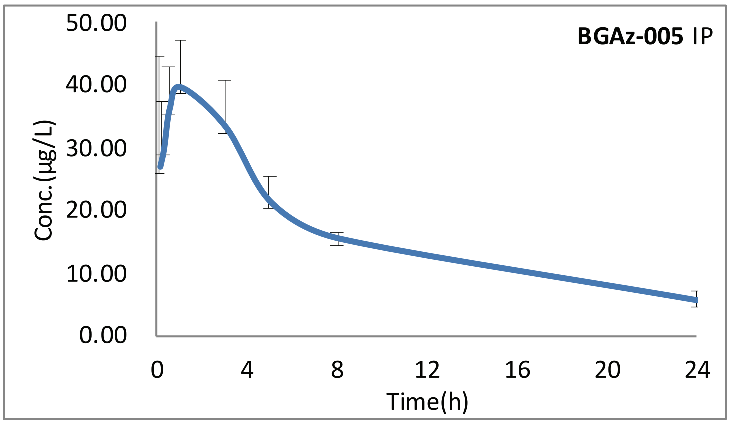

**Figure 29**

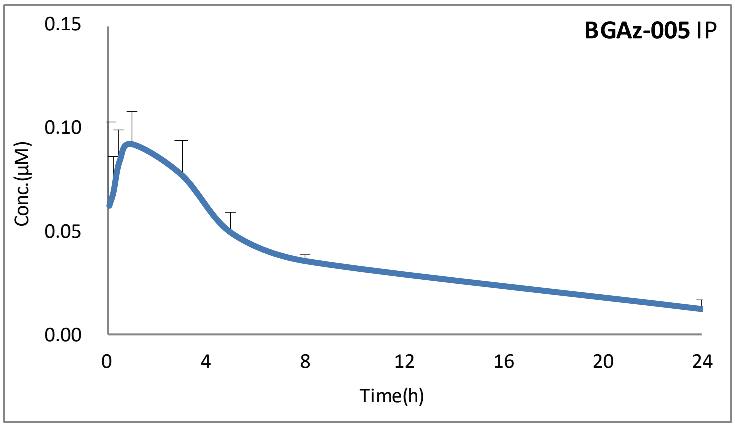

**Figure 30**

### Report on PK results of BGAz-004 by multiple oral dosing

**2020.04.08**

#### 7 Materials and instrument combined dosing

An MS2 Turnover type oscillator was purchased from IKA Work's Guangzhou (China), and a 5415R Chromatographic analyses were conducted using an Agilent 1290 Infinity II high performance liquid chromatography and Agilent Technologies 6470Triple Quad LC/MS. Propranolol (≥90% in purity, internal standard, IS) was purchased from the Sigma Chemical Co. (China). Formic acid (HPLC grade) and methanol (HPLC grade) were purchased from DIKMA Co. (China). All chemicals and solvents were analytical grade. Water was purified using a Millipore (AK, USA) laboratory ultra-pure water system (0.2 µm filter).

#### 8 Animal

KM mice, weighing 25–35 g (Beijing Vital River Laboratory Animal Technology Co., Ltd., China) were utilised for the studies. The protocols were approved by the Animal Care and Use Committee, GIBH. Animals were maintained on standard animal chow and water *ad libitum*, in a climate-controlled room (23±1 °C 30–70% relative humidity, a minimum of ten exchanges of room air per hour and a 12 h light/dark cycle) for one week prior to experiments.

#### 9 Pharmacokinetic studies

Compound **BGAz-004** were dissolved in the solution containing DMSO (2%), ethanol (4%), Cremophor EL (4%) and dd$H_2O$ (90%). The mice were given solution with 30 mg/kg by gastric gavage for 3 days. Whole blood samples (100 µL) were obtained from the orbital venous plexus at the following time points after the third day dosing: 5, 15, 30 min and 1, 3, 5, 8, 24 hour. Whole blood samples were collected in heparinised tubes. The plasma fraction was immediately separated by centrifugation. The mice were humanely euthanasia by carbon dioxide 24 hours after experiment without pain.

#### 10 Plasma sample analysis

##### *10.1 Standard curve sample preparation*

The compounds were dissolved in DMSO (2mg/mL) and diluted with to series concentration (methanol:$H_2O$, 1:1, 10 µL) and blank plasma (50 µL) were added to 1.5 mL tubes and vortexed for 3 min, then acetonitrile-containing internal standard (150 µL) was added and vortexed for a further 5 min, and subsequently spun in a centrifuge at 13000 ×*g* for 40 min at 4 °C, the final concentrations were as follow: 5, 10, 20, 50, 100, 200, 500 ng/mL.

##### *10.2 Plasma and preparation*

Plasma samples were prepared using a protein precipitation method. Solution (10 µL, methanol $H_2O$, 1:1) and plasma samples (50 µL) were added to 1.5 mL tubes and vortex for 3 min, then acetonitrile-containing internal standard (150 µL) was added and vortex for 5 min, and subsequently spun in a centrifuge at 13000 ×*g* for 40 min at 4 °C.

#### 11 LC/MS/MS analysis

After centrifugation, supernatant (100 µL) was transfer to 96 well plates and analysed by LC-MS/MS using an Agilent 1290 Infinity II HPLC and an Agilent Technologies 6470Triple Quad LC/MS.

### 12 Results

**Table 51.** Pharmacokinetic parameters of multiple oral dosing for **BGAz-004**.

| BGAz-004 | |
|---|---|
| **Dosing method** | Multiple PO |
| **Times** | 4 |
| **Animal Number KM mice** | ♂4 |
| **Dose level mg/kg** | 30 |
| **AUC(0-∞) μg*h** | 5420.987 |
| **$T_{1/2}$ (h)** | 8.072 |
| **$T_{max}$ (h)** | 1 |
| **$C_{max}$(μg/L)** | 363.85 |

**Table 52.** Concentration of **BGAz -004** for PO (μg/L).

| Time (h) | No. 1 (μg/L) | No. 2 (μg/L) | No. 3 (μg/L) | No. 4 (μg/L) | Mean (μg/L) | SD (μg/L) |
|---|---|---|---|---|---|---|
| 0.083 | 96.71 | 101.68 | 138.93 | 123.99 | 115.33 | 19.71 |
| 0.25 | 176.35 | 170.28 | 252.32 | 180.64 | 194.90 | 38.52 |
| 0.5 | 236.08 | 351.51 | 392.71 | 293.87 | 318.54 | 68.30 |
| 1 | 265.84 | 421.49 | 413.82 | 354.25 | 363.85 | 71.92 |
| 3 | 245.29 | 393.56 | 354.44 | 290.26 | 320.88 | 65.98 |
| 5 | 195.41 | 337.53 | 302.89 | 236.49 | 268.08 | 64.07 |
| 8 | 155.47 | 404.46 | 251.36 | 161.49 | 243.20 | 116.11 |
| 24 | 66.06 | 11.72 | 82.40 | 94.62 | 63.70 | 36.57 |

**Table 53.** Concentration of **BGAz -004** for PO (μM).

| Time (h) | No. 1 (μM) | No. 2 (μM) | No. 3 (μM) | No. 4 (μM) | Mean (μM) | SD (μM) |
|---|---|---|---|---|---|---|
| 0.083 | 0.179 | 0.188 | 0.257 | 0.230 | 0.214 | 0.036 |
| 0.25 | 0.327 | 0.315 | 0.467 | 0.335 | 0.361 | 0.071 |
| 0.5 | 0.437 | 0.651 | 0.727 | 0.544 | 0.590 | 0.127 |
| 1 | 0.492 | 0.781 | 0.766 | 0.656 | 0.674 | 0.133 |
| 3 | 0.454 | 0.729 | 0.656 | 0.538 | 0.594 | 0.122 |
| 5 | 0.362 | 0.625 | 0.561 | 0.438 | 0.496 | 0.119 |
| 8 | 0.288 | 0.749 | 0.466 | 0.299 | 0.450 | 0.215 |
| 24 | 0.122 | 0.022 | 0.153 | 0.175 | 0.118 | 0.068 |

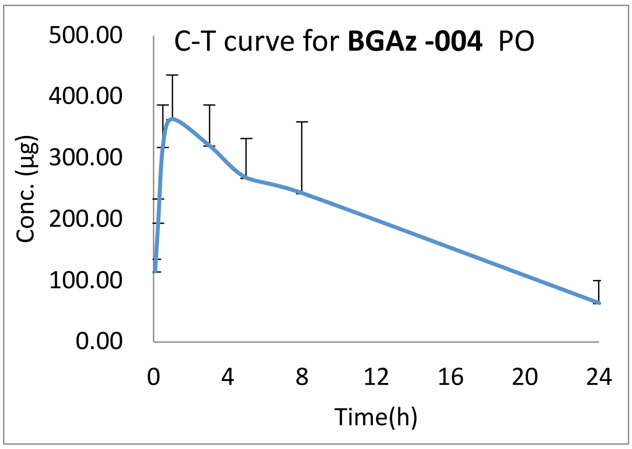

**Figure 31**

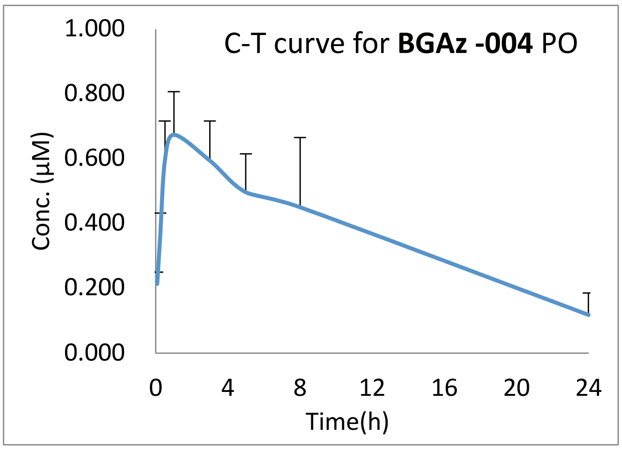

**Figure 32**

**Table 54.** Pharmacokinetic parameters of **BGAz-004** for PO.

| Parameter | Unit | No. 1 | No.2 | No.3 | No.4 | Mean | SD | RSD/% |
|---|---|---|---|---|---|---|---|---|
| **AUC(0-t)** | μg*h | 3454.221 | 6273.905 | 5247.724 | 4069.043 | 4761.223 | 1253.297 | 26.3 |
| **AUC(0-∞)** | μg*h | 4629.924 | 6343.027 | 6433.361 | 4277.637 | 5420.987 | 1126.66 | 20.8 |
| **R_AUC(t/∞)** | % | 74.6 | 98.9 | 81.6 | 95.1 | 87.55 | 11.386 | 13 |
| **AUMC(0-t)** | h*h*μg | 28799.836 | 40171.862 | 41441.784 | 35646.495 | 36514.994 | 5713.256 | 15.6 |
| **AUMC(0-∞)** | h*h*μg | 78051.986 | 42234.646 | 86948.542 | 42419.904 | 62413.77 | 23476.659 | 37.6 |
| **MRT(0-t)** | h | 8.338 | 6.403 | 7.897 | 8.76 | 7.85 | 1.027 | 13.1 |
| **MRT(0-∞)** | h | 16.858 | 6.658 | 13.515 | 9.917 | 11.737 | 4.416 | 37.6 |
| **VRT(0-t)** | h^2 | 50.888 | 11.386 | 44.039 | 59.569 | 41.471 | 21.039 | 50.7 |
| **VRT(0-∞)** | h^2 | 332.554 | 17.556 | 213.745 | 86.244 | 162.525 | 139.485 | 85.8 |
| **λz** | 1/h | 0.056 | 0.171 | 0.07 | 0.118 | 0.104 | 0.052 | 50 |
| **λz 回归尾点** | | 123 | 134 | 124 | 234 | -- | -- | -- |
| **C_last** | μg | 65.712 | 11.83 | 82.441 | 24.622 | 46.151 | 33.372 | 72.3 |
| **t1/2z** | h | 12.399 | 4.049 | 9.967 | 5.871 | 8.072 | 3.801 | 47.1 |
| **Tmax** | h | 1 | 1 | 1 | 1 | 1 | 0 | 0 |
| **Vz/F** | L/kg | 115.931 | 27.634 | 67.065 | 59.414 | 67.511 | 36.517 | 54.1 |
| **CLz/F** | L/h/kg | 6.48 | 4.73 | 4.663 | 7.013 | 5.722 | 1.204 | 21 |
| **Cmax** | μg | 265.8381411 | 421.4879417 | 413.8210272 | 354.252085 | 363.85 | 71.921 | 19.8 |

### Report on PK results of BGAz-005 by multiple oral dosing

**2020.05.22**

#### 13 Materials and instrument combined dosing

An MS2 Turnover type oscillator was purchased from IKA Work's Guangzhou (China), and a 5415R Chromatographic analyses were conducted using an Agilent 1290 Infinity II high performance liquid chromatography and Agilent Technologies 6470Triple Quad LC/MS. Propranolol (≥90% in purity, internal standard, IS) was purchased from the Sigma Chemical Co. (China). Formic acid (HPLC grade) and methanol (HPLC grade) were purchased from DIKMA Co. (China). All chemicals and solvents were analytical grade. Water was purified using a Millipore (AK, USA) laboratory ultra-pure water system (0.2 μm filter).

#### 14 Animal

KM mice, weighing 25–35 g (Beijing Vital River Laboratory Animal Technology Co., Ltd., China) were utilised for the studies. The protocols were approved by the Animal Care and Use Committee, GIBH. Animals were maintained on standard animal chow and water *ad libitum*, in a climate-controlled room (23±1 °C 30–70% relative humidity, a minimum of ten exchanges of room air per hour and a 12 h light/dark cycle) for one week prior to experiments.

#### 15 Pharmacokinetic studies

Compound **BGAz-005** were dissolved in the solution containing DMSO (2%), ethanol (4%), Cremophor EL (4%) and dd$H_2O$ (90%). The mice were given solution with 30 mg/kg by gastric gavage for 3 days. Whole blood samples (100 μL) were obtained from the orbital venous plexus at the following time points after the third day dosing: 5, 15, 30 min and 1, 3, 5, 8, 24 hour. Whole blood samples were collected in heparinised tubes. The plasma fraction was immediately separated by centrifugation. The mice were humanely euthanasia by carbon dioxide 24 hours after experiment without pain.

#### 16 Plasma sample analysis

##### *16.1 Standard curve sample preparation*

The compounds were dissolved in DMSO (2mg/mL) and diluted with to series concentration (methanol:$H_2O$, 1:1, 10 μL) and blank plasma (50 μL) were added to 1.5 mL tubes and vortexed for 3 min, then acetonitrile-containing internal standard (150 μL) was added and vortexed for a further 5 min, and subsequently spun in a centrifuge at 13000 ×*g* for 40 min at 4 °C, the final concentrations were as follow: 5, 10, 20, 50, 100, 200, 500, 2000 ng/mL.

##### *16.2 Plasma and preparation*

Plasma samples were prepared using a protein precipitation method. Solution (10 μL, methanol $H_2O$, 1:1) and plasma samples (50 μL) were added to 1.5 mL tubes and vortex for 3 min, then acetonitrile-containing internal standard (150 μL) was added and vortex for 5 min, and subsequently spun in a centrifuge at 13000 ×*g* for 40 min at 4 °C.

#### 17 LC/MS/MS analysis

After centrifugation, supernatant (100 μL) was transfer to 96 well plates and analysed by LC-MS/MS using an Agilent 1290 Infinity II HPLC and an Agilent Technologies 6470Triple Quad LC/MS.

**Table 55.** Pharmacokinetic parameters of multiple oral dosing for **BGAz-005**.

| **BGAz-005** | |
|---|---|
| **Dosing method** | Multiple PO |
| **Times** | 3 |
| **Animal Number KM mice** | ♂4 |
| **Dose level mg/kg** | 30 |
| **AUC(0-∞) μg/L*h** | 82247.729 |
| **$T_{1/2}$ (h)** | 80.437 |
| **$T_{max}$ (h)** | 0.875 |
| **$C_{max}$(μg/L)** | 1712.54 |

**Table 56.** Concentration of **BGAz-005** for PO (μg/L).

| Time (h) | No. 1 (μg/L) | No. 2 (μg/L) | No. 3 (μg/L) | No. 4 (μg/L) | Mean (μg/L) | SD (μg/L) |
|---|---|---|---|---|---|---|
| 0.083 | 513.99 | 1123.99 | 826.27 | 968.72 | 858.24 | 259.72 |
| 0.25 | 656.54 | 935.75 | 1196.01 | 1887.19 | 1168.87 | 527.11 |
| 0.5 | 702.36 | 1539.13 | 1591.63 | 2314.39 | 1536.88 | 659.28 |
| 1 | 979.43 | 1928.73 | 1276.73 | 2350.26 | 1633.79 | 620.75 |
| 3 | 535.60 | 1377.36 | 631.57 | 875.45 | 854.99 | 376.48 |
| 5 | 614.08 | 831.63 | 654.72 | 754.98 | 713.85 | 98.34 |
| 8 | 507.07 | 594.94 | 570.05 | 759.27 | 607.83 | 107.52 |
| 24 | 383.27 | 744.63 | 605.57 | 546.05 | 569.88 | 149.67 |

**Table 57.** Concentration of **BGAz-005** for PO (μM).

| Time (h) | No. 1 (μM) | No. 2 (μM) | No. 3 (μM) | No. 4 (μM) | Mean (μM) | SD (μM) |
|---|---|---|---|---|---|---|
| 0.083 | 1.181 | 2.582 | 1.898 | 2.225 | 1.971 | 0.597 |
| 0.25 | 1.508 | 2.149 | 2.747 | 4.334 | 2.685 | 1.211 |
| 0.5 | 1.613 | 3.535 | 3.656 | 5.316 | 3.530 | 1.514 |
| 1 | 2.250 | 4.430 | 2.932 | 5.398 | 3.752 | 1.426 |
| 3 | 1.230 | 3.163 | 1.451 | 2.011 | 1.964 | 0.865 |
| 5 | 1.410 | 1.910 | 1.504 | 1.734 | 1.640 | 0.226 |
| 8 | 1.165 | 1.366 | 1.309 | 1.744 | 1.396 | 0.247 |
| 24 | 0.880 | 1.710 | 1.391 | 1.254 | 1.309 | 0.344 |

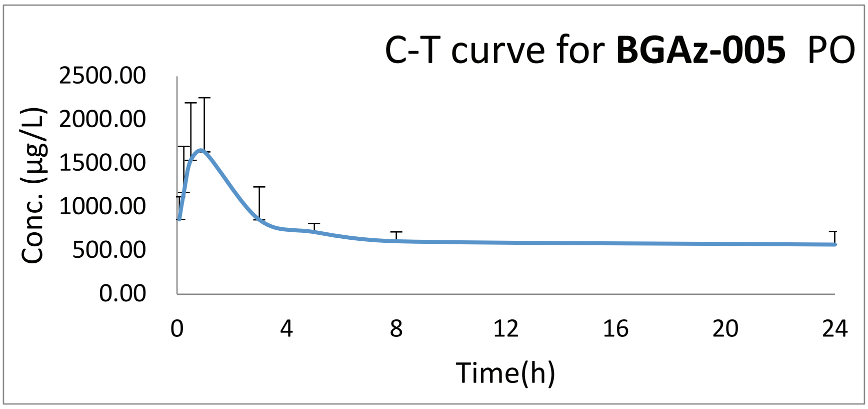

**Figure 33**

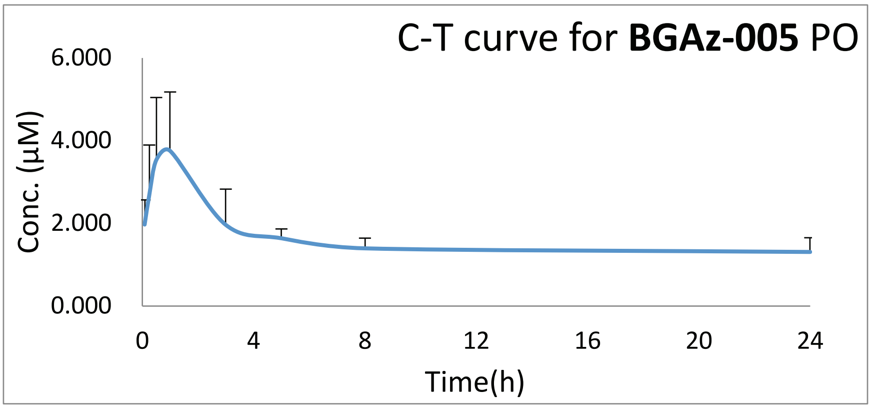

**Figure 34**

**Table 58.** Pharmacokinetic parameters of **BGAz-005** for PO.

| Parameter | Unit | No. 1 | No.2 | No.3 | No.4 | Mean | SD | RSD/% |
|---|---|---|---|---|---|---|---|---|
| **AUCss** | μg/L*h | 12178.507 | 19766.468 | 15705.449 | 19540.058 | 16797.621 | 3599.262 | 21.4 |
| **AUC(0-t)** | μg/L*h | 12178.507 | 19766.468 | 15705.449 | 19540.058 | 16797.621 | 3599.262 | 21.4 |
| **AUC(0-∞)** | μg/L*h | 35753.919 | 20021.166 | 228986.832 | 44229 | 82247.729 | 98338.883 | 119.6 |
| **R_AUC(t/∞)** | % | 34.1 | 98.7 | 6.9 | 44.2 | 45.975 | 38.518 | 83.8 |
| **AUMC(0-t)** | h*h*μg/L | 124409.562 | 209604.037 | 173532.842 | 180715.464 | 172065.476 | 35389.288 | 20.6 |
| **AUMC(0-∞)** | h*h*μg/L | 2135136.115 | 217275.027 | 80252935.26 | 1894415.043 | 21124940.36 | 39427892.07 | 186.6 |
| **MRT(0-t)** | h | 10.216 | 10.604 | 11.049 | 9.248 | 10.279 | 0.767 | 7.5 |
| **MRT(0-∞)** | h | 59.718 | 10.852 | 350.47 | 42.832 | 115.968 | 157.643 | 135.9 |
| **VRT(0-t)** | h^2 | 70.025 | 83.458 | 81.101 | 70.246 | 76.208 | 7.078 | 9.3 |
| **VRT(0-∞)** | h^2 | 3766.565 | 87.655 | 123543.426 | 2074.815 | 32368.115 | 60802.133 | 187.8 |
| **λz** | 1/h | 0.016 | 0.163 | 0.003 | 0.022 | 0.051 | 0.075 | 147.1 |
| **λz 回归尾点** | | 124 | 234 | 134 | 124 | -- | -- | -- |
| **C_last** | μg/L | 384.659 | 41.631 | 606.838 | 543.67 | 394.2 | 252.951 | 64.2 |
| **t1/2z** | h | 42.473 | 4.24 | 243.564 | 31.47 | 80.437 | 109.932 | 136.7 |
| **Tmax** | h | 1 | 1 | 0.5 | 1 | 0.875 | 0.25 | 28.6 |
| **Vz/F** | L/kg | 51.426 | 9.167 | 46.046 | 30.802 | 34.36 | 18.931 | 55.1 |
| **Vss/F** | L/kg | 50.107 | 16.261 | 45.916 | 29.052 | | | |
| **CLz/F** | L/h/kg | 0.839 | 1.498 | 0.131 | 0.678 | 0.787 | 0.563 | 71.5 |
| **Cmax** | μg/L | 979.4331092 | 1928.734817 | 1591.628366 | 2350.260294 | 1712.514 | 578.934 | 33.8 |
| **Cav** | μg/L | 507.438 | 823.603 | 654.394 | 814.169 | 699.901 | 149.969 | 21.4 |
| **DF** | | 1.93 | 2.342 | 2.432 | 2.887 | 2.398 | 0.393 | 16.4 |
